# Tumor-tropic *E. coli* engineered as living T and NK cell engagers

**DOI:** 10.64898/2026.08.18.745642

**Authors:** Shaobo Yang, Anna Clara Bader, Stephanie Sendker, Ashley Hu, Daniel Chuyang Chen, Hetal Nath, Alice Chen, Eden Bobilev, Michal Sheffer, Veronica W. Hui, Tereza E. Kochs, Andreia Maia, Jiting Tang, Fuguo Liu, Xingyu Deng, Maily Nguyen, Mila Stanojevic, Mubin Tarannum, Castilleja L. Albert, Alaa Kassim Ali, Roman Shapiro, Yun Wei, Ke Zhang, Zongqi Wang, Young Rock Chung, Erin Parry, Marco Campisi, David Barbie, Andrew A. Lane, Hongyu Li, Keith L. Ligon, Kun Huang, Kai W. Wucherpfennig, Seema Chugh, Evelyn Ullrich, Hermann Einsele, Jianzhu Chen, John Koreth, Vanessa S. Silveira, Robert Soiffer, Jessica S. Little, Catherine J. Wu, Jerome Ritz, Jiahe Li, Andrew J. Aguirre, Rizwan Romee

## Abstract

Despite advances in immunotherapy, most solid tumors remain resistant to treatment. Immune cell engagers redirect cytotoxic lymphocytes against cancer, but limited tumor access, immunosuppressive microenvironments and systemic immune activation limit efficacy. Here we develop live immune modulating engagers (LIME), a modular platform where non-pathogenic, tumor-tropic *Escherichia coli* display tandem single-chain variable fragments targeting a tumor-associated antigen and an activating receptor on T or natural killer cells. LIME bridged effector and tumor cells, induced transcriptional programs of T cell activation, metabolism and proliferation, and enhanced cytotoxicity across cancer cell lines and patient-derived organoids. In mouse models, LIME safely accumulated in tumors, outperformed tarlatamab in small cell lung cancer, and induced durable immunity in lymphoma. RAS inhibition and PD-L1 blockade enhanced LIME activity in pancreatic cancer and induced humoral responses. Multi-lineage immune modulation remained tumor-confined, without organ toxicity. These findings establish LIME as a versatile living therapeutic platform for programmable, tumor-restricted immune orchestration.

## Main

Harnessing the immune system to recognize and eliminate malignant cells has transformed cancer therapy^1, 2^. Immune cell engagers represent one of the most direct implementations of this principle: these multi-specific proteins, most commonly bispecific antibodies, simultaneously bind a tumor-associated antigen and an activating receptor on an immune effector cell. By bringing tumor and immune cells into close proximity, these agents redirect cytotoxicity independent of the endogenous antigen specificity of the effector cell. CD3 directed T cell engagers have produced substantial clinical activity in hematologic malignancies, including CD19^+^ B cell precursor acute lymphoblastic leukemia and BCMA^+^ multiple myeloma^3, 4^. Progress in solid tumors has been more limited, tebentafusp, a soluble gp100-targeted T-cell receptor–anti-CD3 fusion protein, improved survival in metastatic uveal melanoma, and the DLL3-CD3 bispecific T cell engager tarlatamab was approved for previously treated extensive stage small cell lung cancer^5–7^. Together, these successes establish the therapeutic potential of immune-cell redirection while highlighting the need for platforms that can extend this activity across a broader range of solid tumors.

Several features of solid tumors limit the efficacy and therapeutic window of conventional immune engagers. Abnormal vasculature, dense extracellular matrix and cancer-associated fibroblast networks produce heterogeneous antibody delivery and restrict access to malignant cells^8–10^. In human pancreatic ductal adenocarcinoma (PDAC), for example, recent spatial pharmacology studies have linked periostin-rich extracellular matrix and FAP^+^ fibroblast neighborhoods to reduced therapeutic antibody delivery^11^. Engager activity is further constrained by heterogeneous antigen expression, insufficient effector cell infiltration and local immunosuppression^12^. Conversely, systemic exposure to CD3 directed agents can activate T cells in the peripheral blood and cause cytokine release syndrome and neurologic toxicity, as reflected in the boxed warning for tarlatamab^6, 13^. Pharmacokinetic and manufacturing constraints also vary by molecular formats: compact Fc-free BiTEs such as blinatumomab have short serum half-lives and require continuous infusion, whereas newer Fc-containing molecules achieve longer exposure at the cost of introducing additional engineering, assembly and purification demand^14, 15^. A strategy that simplifies manufacturing and concentrates immune engager activity within tumors could therefore improve local efficacy while limiting systemic immune activation.

Microbial cancer therapy offers a complementary approach to spatially restricted treatment. Association between bacterial infection and tumor regression has been recognized for more than a century, and subsequent studies have shown that selected non-pathogenic strains of *Escherichia*, *Salmonella*, *Listeria*, *Clostridium* and *Bifidobacterium* preferentially accumulate in tumors^16–18^. Tumor colonization is supported by hallmarks of tumor microenvironment; hypoxia, necrosis, abnormal vasculature, limited nutrient availability and relative immune privilege, that create ecological niches largely inaccessible to most conventional therapies. Once established, bacteria can amplify locally, disrupt tumor tissue, activate innate and adaptive immunity and serve as programmable vehicles for therapeutic proteins^16–18^. These properties make living bacteria attractive for treating poorly perfused and potentially disseminated tumors.

Clinical experience has begun to define both the promise and the challenges of this approach. Intravenous administration of attenuated *Salmonella Typhimurium* VNP20009 demonstrated that live bacteria could be safely administered in patients with advanced cancer, though no objective responses were observed^19^. Intratumoral *Clostridium novyi*-NT subsequently showed selective germination and tumor lysis in treatment-refractory solid tumors^20^. More recently, Saltikva, an orally administered attenuated *Salmonella Typhimurium* engineered to express human IL-2, was well tolerated and increased circulating natural killer (NK) and natural killer T (NKT) cells in a phase I trial^21^. A non-randomized phase II study of repeated Saltikva dosing with FOLFIRINOX in metastatic PDAC reported encouraging progression-free and overall survival signals, though randomized testing is required to establish efficacy^22^. A phase I study of the engineered *Escherichia coli* (*E. coli*) Nissle therapeutic SYNB1891 has likewise supported the clinical feasibility of genetically programmed bacterial immunotherapies^23^. Collectively, these studies provide an emerging translational foundation for living microbial therapeutics but also indicate that bacterial tropism alone is unlikely to yield consistent therapeutic benefit. Advances in synthetic biology now offer an opportunity to convert tumor colonizing bacteria into programmable medicines with defined therapeutic payloads, regulated activity and engineered biocontainment.

Non-pathogenic *E. coli* is a particularly attractive living therapeutic chassis: it is genetically tractable, compatible with scalable fermentation and, depending on the strain, sensitive to a broad array of clinically available antibiotics^24, 25^. As a facultative anaerobe, *E. coli* can grow across oxygen gradients and preferentially colonizes hypoxic and necrotic tumor regions^26^, and its outer membrane can be engineered to display functional proteins directly at the bacterial surface. We previously developed a surface display toolbox for non-pathogenic *E. coli* and showed that tumor tropic bacteria displaying a decoy-resistant IL-18 mutein activated intratumoral T and NK cells and enhanced CAR NK cell trafficking and efficacy^27^. We subsequently extended this strategy beyond cancer immunotherapy by displaying the colibactin self-resistance protein ClbS on engineered bacteria, thereby preventing colibactin-mediated genotoxicity in human cell and organoid systems and suppressing intestinal injury and tumorigenesis in mouse models^28^. Together, these studies establish bacterial surface as a versatile platform for presenting biologically active proteins. Unlike bacterial secretion or lysis, surface anchoring keeps the displayed molecule physically associated with the living chassis and enables multivalent interactions with neighboring cells.

Here, we introduce Live Immune Modulating Engagers (LIME), which, to our knowledge, represent the first modular living bacterial immune engager platform. LIME consists of non-pathogenic, tumor tropic *E. coli* engineered to display tandem single-chain variable fragments (scFvs) directed against a tumor-associated surface antigen and an immune effector receptor (**Fig. 1a**). The modularity of LIME’s two antigen binding domains enables each arm to be independently exchanged, reprogramming the platform to recruit T or NK cells against diverse tumor antigens. We demonstrate that LIME bridges effector and tumor cells, induces distinct activation and proliferative programs, and enhances cytotoxicity across cancer cell lines and patient derived PDAC organoids. In vivo, LIME preferentially accumulates in tumors, concentrates effector cell activation and inflammatory cytokine production within the tumor microenvironment, controls xenograft and syngeneic tumors, and outperforms tarlatamab in DLL3^+^ small cell lung cancer models. Combining LIME with immune checkpoint blockade or RAS inhibition further enhances antitumor activity in immunocompetent models. Together, these findings establish a living, modular strategy that couples the precision of immune cell engagers with the tumor-homing and locally amplifying properties of engineered bacteria.

**Fig. 1.**
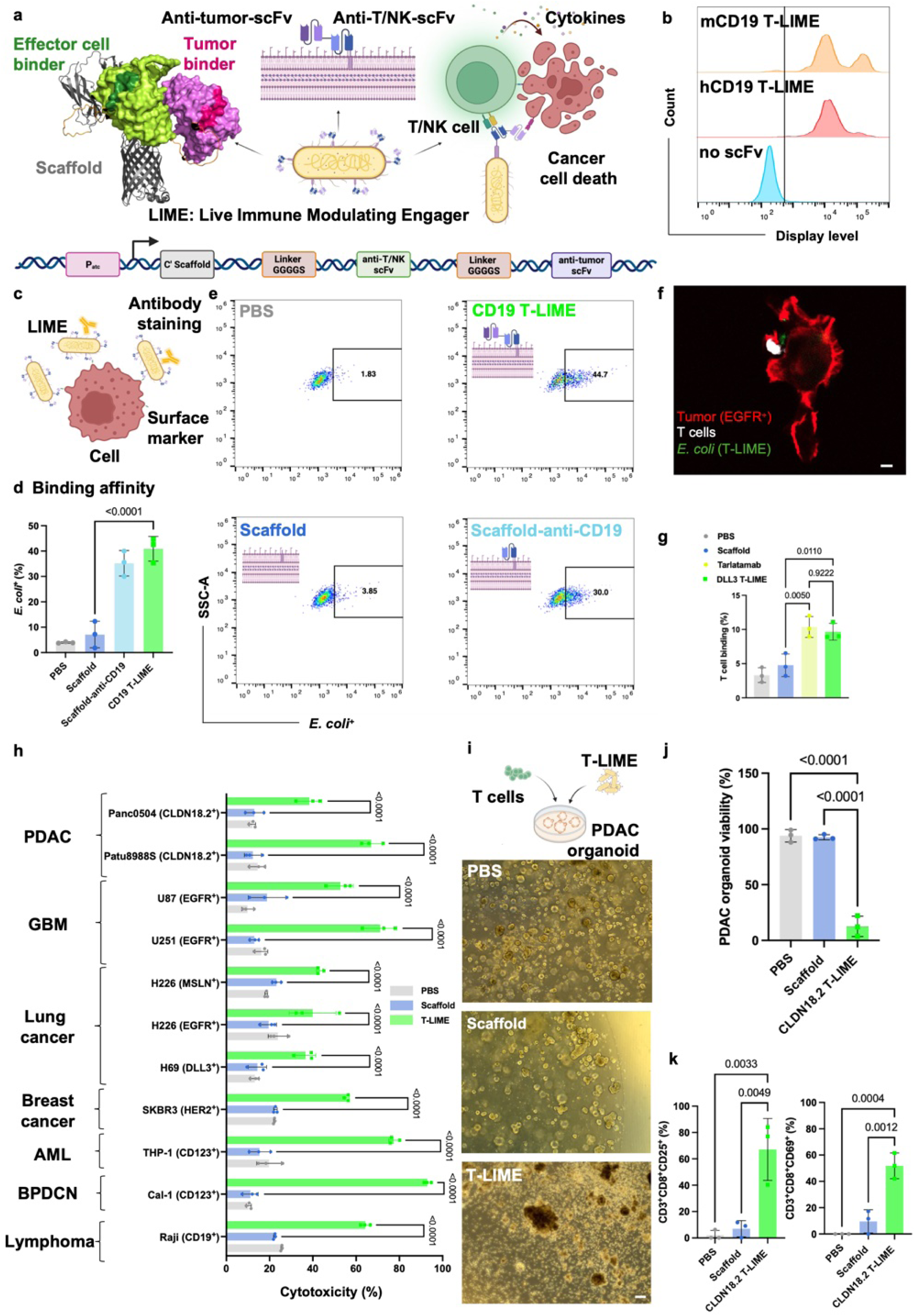
LIME bridge effector and target cells to induce potent antitumor responses in vitro. **a**, Schematic illustration and construct design of Live Immune Modulating Engagers (LIME). Briefly, non-pathogenic *E. coli* K-12 were engineered to display two single-chain variable fragments (scFvs) that bridge effector (T or NK) and cancer cells, thereby triggering effector cell activation and mediating tumor cell killing. **b**, Histograms showing the surface display levels of control bacteria (no scFv; engineered *E. coli* displaying only the C′ scaffold, as indicated in Fig. 1a) and two representative LIMEs: engineered *E. coli* displaying anti-murine CD3 and CD19 scFvs (mCD19 T-LIME) or anti-human CD3 and CD19 scFvs (hCD19 T-LIME). **c**, Schematic of the binding assay used to evaluate the affinity of LIME for cells expressing the corresponding surface marker. **d**, **e**, Bar graph (**d**) and representative flow cytometry dot plot (**e**) illustrating the binding of hCD19 T-LIME to CD19-positive Raji cells. **f**, **g**, Immune synapse formation, as described in Methods, between CD3^+^ T cells and tumor cells mediated by T-LIME or control (PBS, tarlatamab or scaffold control consisting of engineered *E. coli* displaying only the C′ scaffold), analyzed by confocal microscopy (**f**; T cells and the EGFR^+^ glioblastoma multiforme cell line U87; scale bar = 5 μm) or flow cytometry (**g**; T cells and the DLL3^+^ small cell lung cancer [SCLC] cell line H69). **h**, Cytotoxicity of T cells against various tumor cell lines mediated by the corresponding T-LIME or controls (PBS or scaffold). Tumor cell lines included PDAC (Panc0504 and Patu8988S, CLDN18.2^+^), glioblastoma multiforme (GBM; U87 and U251, EGFR^+^), lung cancer (H69, DLL3^+^; H226, EGFR^+^MSLN^+^), breast cancer (SKBR3, HER2^+^), acute myeloid leukemia (AML; THP-1, CD123^+^), blastic plasmacytoid dendritic cell neoplasm (BPDCN; Cal-1, CD123^+^) and lymphoma (Raji, CD19^+^). **i**-**k**, Schematic, representative images (**i**), quantification of T cell-mediated killing efficacy (**j**) and CD8^+^ T cell activation (**k**) in CLDN18.2^+^ PDAC patient-derived organoids, analyzed by flow cytometry. Statistical significance was determined using one-way ANOVA in **d**, **g**, **h**, **j** and **k**. Data are presented as mean ± s.d.

## Results

### Non-pathogenic, tumor-tropic *E. coli* engineered as potent immune engagers

To identify an optimal bacterial surface-display scaffold for single-chain variable fragments (scFvs), we fused an anti-CD19 scFv to five candidate scaffolds: lipoprotein-outer membrane protein A (OmpA), the C-terminal domain of IgA protease (C-IgAP), the N-terminal domain of intimin (Neae), and two N-terminal fragments of YiaT comprising amino acids 1-232 (YiaT232) or 1-181 (YiaT181). Surface display was quantified by flow cytometry (**Extended Data Fig. 1a, b**). All five scaffolds supported detectable surface expression of the anti-CD19 scFv (**Extended Data Fig. 1c**). We next assessed the functionality of the displayed scFvs by measuring bacterial binding to the CD19^+^ Raji lymphoma cells and among the scaffolds tested, OmpA and Neae mediated the strongest binding (**Extended Data Fig. 1d**). Because activation of effector cells is essential for immune-engager activity, we cocultured primary human CD3^+^ T cells with *E. coli* displaying an anti-CD3 scFv from each scaffold. Neae-based display promoted robust bacterial association with both CD4^+^ and CD8^+^ T cells and induced expression of the activation markers CD25 and CD69 (**Extended Data Fig. 1e**). Based on these results with the combined display and optimal functional performance, we selected Neae for the subsequent studies.

We next engineered live immune-modulating engagers (**LIME**) by using Neae to display two scFvs in tandem, separated by four G_4_S repeats: one directed against an effector-cell receptor and the other against a tumor-associated surface antigen (**Fig. 1a** and **Supplemental Fig. 1a**). Effector-cell-targeting scFvs recognized CD3 to engage T cells, and CD16, NKG2D or NKp46 (two independent clones) to engage NK cells, whereas tumor-targeting scFvs recognized Claudin 18.2 (CLDN18.2), Delta-like ligand 3 (DLL3), epidermal growth factor receptor (EGFR), CD123, CD19, mesothelin (MSLN) or human epidermal growth factor receptor 2 (HER2). Flow cytometry confirmed surface display of diverse human and murine scFv combinations (**Fig. 1b** and **Supplemental Fig. 1b, c**). We refer to constructs engaging T cells (CD3) as T-LIME and those engaging NK-cell-abundant receptors (NKG2D, NKp46 and CD16) as N-LIME; individual constructs are designated by their tumor target, such as CD19 T-LIME.

To determine whether our tandem scFv display preserved the activity of both binding domains, we first assessed the binding of murine CD19 T-LIME to CD19^+^ A20 cells. CD19 T-LIME associated efficiently with A20 cells, at a level comparable to bacteria displaying the corresponding monospecific anti-CD19 scFv (**Fig. 1c-e**). Across a broader panel of constructs, both T-LIME and N-LIME also associated with their cognate effector cells, increased expression of CD25 and CD69, and promoted proliferation, as indicated by Ki67 expression (**Supplemental Fig. 2a-e**). These findings indicate that both targeting domains remain functional in the tandem LIME architecture.

We then tested whether T-LIME could physically bridge effector and tumor cells. Coculture assays were performed using primary human CD3^+^ T cells and tumor cell lines expressing EGFR (U87, glioblastoma; GBM), CLDN18.2 (Patu8988s, pancreatic ductal adenocarcinoma; PDAC) or DLL3 (H69, small-cell lung cancer; SCLC). Confocal microscopy and flow cytometry demonstrated increased formation of T cell-tumor cell conjugates in the presence of the corresponding T-LIME (**Fig. 1f, g** and **Supplemental Fig. 3a-e**). DLL3 T-LIME mediated conjugate formation between T cells and DLL3-positive SCLC cells at a level comparable to that achieved with the FDA approved DLL3-targeting T-cell engager tarlatamab (**Fig. 1g** and **Supplemental Fig. 3c-e**), consistent with the formation of engager-mediated immune synapses.

We next evaluated whether LIME-mediated cell bridging translated into tumor-cell killing *in vitro*. In Annexin V and 7-AAD-based cytotoxicity assays, cognate T-LIME constructs significantly increased T-cell-mediated killing across tumor cell lines representing PDAC, glioblastoma, SCLC, non-small cell lung cancer, breast cancer, acute myeloid leukemia (AML) and B cell lymphoma (**Fig. 1h**). Phenotypic analysis showed that T-LIME treatment increased bacterial association with CD4^+^ and CD8^+^ T cells and enhanced their activation and proliferation in presence of tumor cells (**Supplemental Fig. 4a-c**). Increased bacterial association was also detected on tumor cells exposed to the corresponding T-LIME relative to PBS and scaffold controls (**Supplemental Fig. 5**), further supporting dual engagement of effector and target cells.

N-LIME similarly enhanced NK-cell-mediated cytotoxicity across multiple tumor cell lines (**Extended Data Fig. 2a-c**). In tumor cell cocultures, N-LIME associated with both NK cells and cognate tumor cells and increased NK cell activation and proliferation, as measured by CD25, CD69 and Ki67 expression (**Extended Data Fig. 3a-f** and **Supplemental Fig. 6**). The magnitude of these responses varied among NK cell-targeting binders: constructs engaging CD16 or NKp46 clone 1 generally produced the strongest cytotoxic and activation responses (**Extended Data Fig**. **2a-c** and **Extended Data Fig. 3a-f**). These differences highlight binder selection as an important parameter for optimizing N-LIME activity.

Finally, we evaluated T-LIME activity in a more physiologically relevant patient-derived model. CLDN18.2 T-LIME increased human T cell-mediated killing of CLDN18.2-positive PDAC organoids (**Fig. 1i, j**). Flow cytometric profiling further showed activation of both CD8^+^ and CD4^+^ T cells in these cocultures (**Fig. 1k** and **Supplemental Fig. 7a, b**).

Together, these results establish LIME as a plug and play modular bacterial immune engager platform that can be adapted by exchanging tumor- and effector-cell-targeting scFvs. LIME constructs retained functional engagement of both target populations, promoted effector-target cell conjugation, activated T or NK cells, and enhanced cytotoxicity across diverse cancer cell lines and patient derived PDAC organoids.

### LIME reshapes the transcriptional landscape of human T cells

To determine how LIME engages and activates effector cells, we next sought to define transcriptional programs induced by LIME. We compared primary T cells exposed to PBS, scaffold bacteria, the soluble DLL3-targeting T-cell engager (tarlatamab) or DLL3 T-LIME for 12 hours, followed by bulk RNA sequencing. The principal component analysis, after correcting for donor-specific effects, revealed that DLL3 T-LIME treated cells occupied a distinct transcriptional space compared to PBS, scaffold, and tarlatamab controls (**Fig. 2a**). We compared DLL3 T-LIME with scaffold bacteria to isolate the transcriptional consequences of LIME-mediated engagement. DLL3 T-LIME altered the expression of genes associated with T cell activation and inflammatory signaling, including *IRF4, IL2, IFNG, CD69, CCL4, CCL4L2, TNFRSF9, MIR155HG* and TNFAIP3 (**Fig. 2b, c**). Gene set enrichment analysis (GSEA) further demonstrated positive enrichment of programs associated with mTORC1 signaling, IL-2-STAT5 signaling, MYC targets, the G2M checkpoint, interferon-gamma responses and glycolysis in DLL3 T-LIME-treated cells relative to scaffold-treated cells (p < 0.05, q < 0.25, |NES| > 1, **Fig. 2d**). Enrichment plots highlighted strong activation of the IL-2-STAT5 pathway and positive enrichment of TNF signaling through NF-kB (**Fig. 2e**). These results indicate that DLL3 T-LIME induce a T cell transcriptional state distinct from that elicited by scaffold bacteria, characterized by enrichment of cytokine signaling, metabolic and cell-cycle programs associated with T cell activation.

**Fig. 2.**
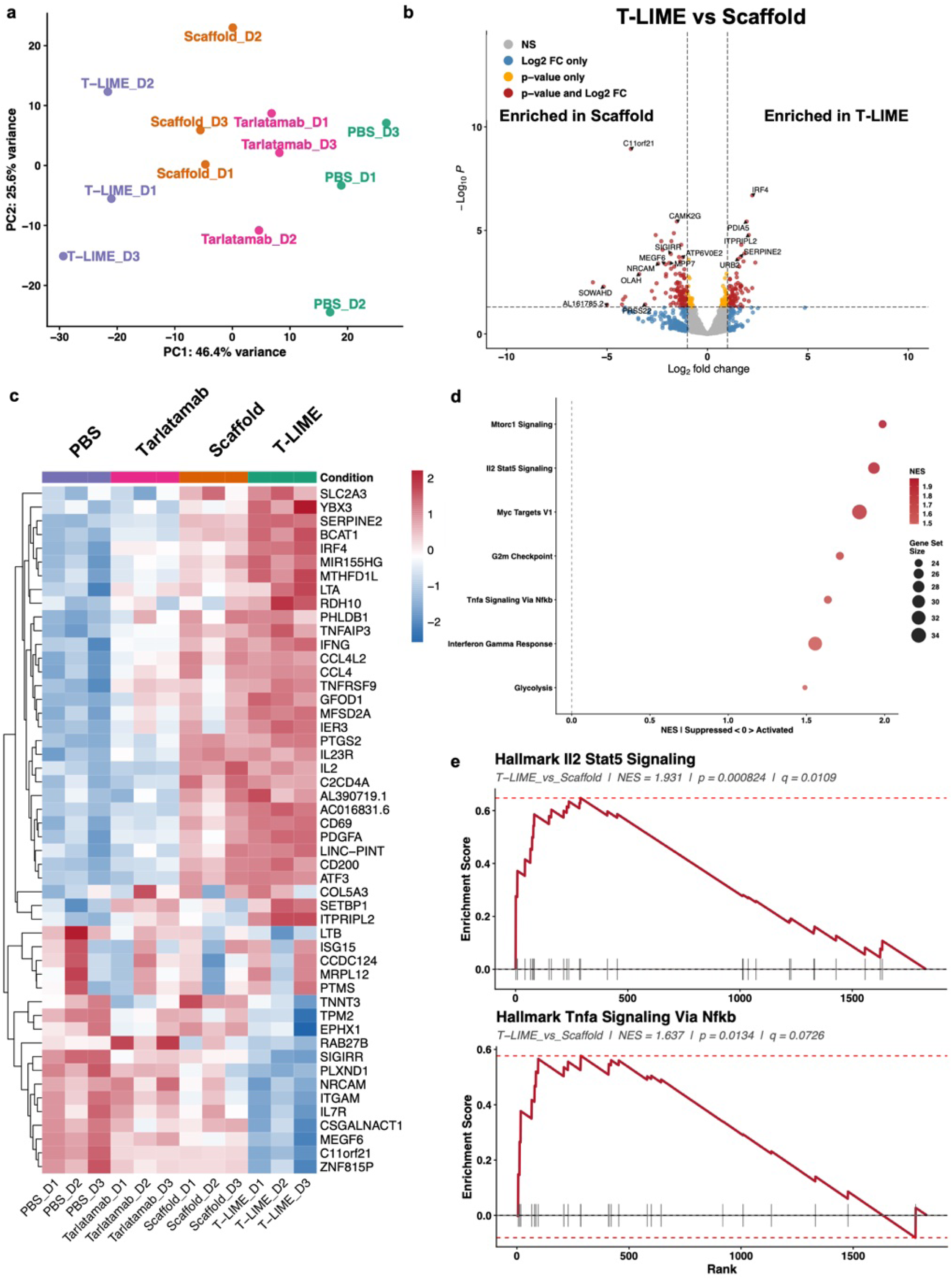
T-LIME transcriptionally activates human T cells and induces inflammatory antitumor signaling. **a**, Principal component analysis (PCA) of RNA-seq profiles from human primary T cells (three donors: D1, D2, D3) following coculture with PBS, Scaffold, tarlatamab, or DLL3 T-LIME after donor correction. **b**, Volcano plot showing differentially expressed genes in T cells treated with DLL3 T-LIME compared with Scaffold control. **c**, Heatmap showing the top differentially expressed genes across treatment groups after donor correction. **d**, Gene set enrichment analysis (GSEA) of MSigDB Hallmark pathways enriched in DLL3 T-LIME-treated T cells compared with Scaffold-treated T cells. Dot size indicates gene set size and color indicates normalized enrichment score (NES). **e**, Representative GSEA enrichment plots showing activation of Hallmark IL2–STAT5 signaling and Hallmark TNFα signaling via NF-κB in DLL3 T-LIME-treated T cells compared with Scaffold control. Differential gene expression analysis was performed using DESeq2. Pathway enrichment was assessed using GSEA. Significance thresholds are indicated in the plots.

Pairwise comparisons with PBS further resolved the contributions of the soluble engager and bacterial scaffold. Relative to PBS, tarlatamab induced genes associated with T cell activation, including *IRF4, TNFRSF9, CCL4, CCL4L2 and PTGS2,* together with enrichment of IL-2-STAT5, interferon gamma, TNF-NF-kB and inflammatory response pathways (**Extended Data Fig. 4a-d**). Scaffold bacteria also produced a broad transcriptional response relative to PBS, marked by increased expression of *IL2, IFNG, CD69, TNFRSF9, CCL4, CCL4L2, NFKB1, TNFAIP3 and PTGS2*. This response was accompanied by enrichment of inflammatory, TNF-NF-kB, interferon gamma and IL-2-STAT5 programs, whereas the G2M-checkpoint program was negatively enriched (**Extended Data Fig. 5a-d**). These results show that the bacterial scaffold control on its own elicits a broad inflammatory transcriptional response under these culture conditions.

DLL3 T-LIME elicited a broader response relative to PBS, with pronounced differential expression of genes involved in cytokine signaling, activation and inflammatory regulation, including *IL2, IFNG, CD69, CCL4, CCL4L2, IRF4, TNFRSF9, TNFAIP3 and PTGS2* (**Supplemental Fig. 8a-d**). Consistent with these gene level changes, DLL3 T-LIME enriched TNF-NF-kB, interferon gamma, IL-2-STAT5, mTORC1, inflammatory-response, apoptosis and hypoxia-associated programs. Direct comparison with tarlatamab showed that DLL3 T-LIME induced a distinct transcriptional state characterized by stronger mTORC1 and TNF-NF-kB pathway activity, together with positive enrichment of IL-2-STAT5 and interferon gamma response programs (**Supplemental Fig. 9a-d**). Conversely, comparison of tarlatamab with scaffold bacteria showed preferential enrichment of E2F targets, the G2M checkpoint and mitotic-spindle programs following tarlatamab treatment, whereas inflammatory response genes such as *IL2, PTGS2, SERPINE2, CD200* and *ATF3* were more highly expressed following scaffold treatment (**Supplemental Fig. 10a-d**). These comparisons suggest that soluble T-cell engagement and bacterial stimulation make distinct contributions to the transcriptional response, which are integrated in DLL3 T-LIME-treated cells.

To assess pathway activity across all treatment groups without relying on individual pairwise comparisons, we performed donor corrected gene set variation analysis (GSVA). Hallmark pathway analysis identified treatment-associated differences in IL-2-STAT5 signaling, TNF signaling through NF-kB, apoptosis, hypoxia and p53 signaling, as well as mitotic-spindle and apical-junction programs (**Supplemental Fig. 11**). Analysis of KEGG pathways independently identified differential activity in immune signaling networks, including JAK-STAT, T-cell receptor and MAPK signaling pathways (**Supplemental Fig. 12**). Together, these analyses show that DLL3 T-LIME establishes a transcriptional state that integrates inflammatory signaling driven by the bacterial chassis with cytokine, metabolic and cell-cycle programs associated with targeted T cell engagement.

### DLL3 T-LIME drives localized T-cell activation and tumor control in small cell lung cancer xenografts

To evaluate the in vivo antitumor activity and pharmacodynamic profile of LIME and to benchmark against FDA approved DLL3 T-cell engager (tarlatamab), we established a human small cell lung cancer (SCLC) xenograft model by subcutaneously implanting DLL3^+^ H69 cells into immunocompromised NSG mice. Human primary CD3^+^ T cells were administered intravenously on day 11 after tumor implantation. Beginning on day 12, mice received three intravenous doses of DLL3 T-LIME, scaffold bacteria (2 × 10^8^ CFU per dose), or PBS, on days 12, 17, and 21. A separate group received tarlatamab intraperitoneally (3 mg/kg) on the same schedule to provide a clinically relevant comparison (**Fig. 3a**).

**Fig. 3.**
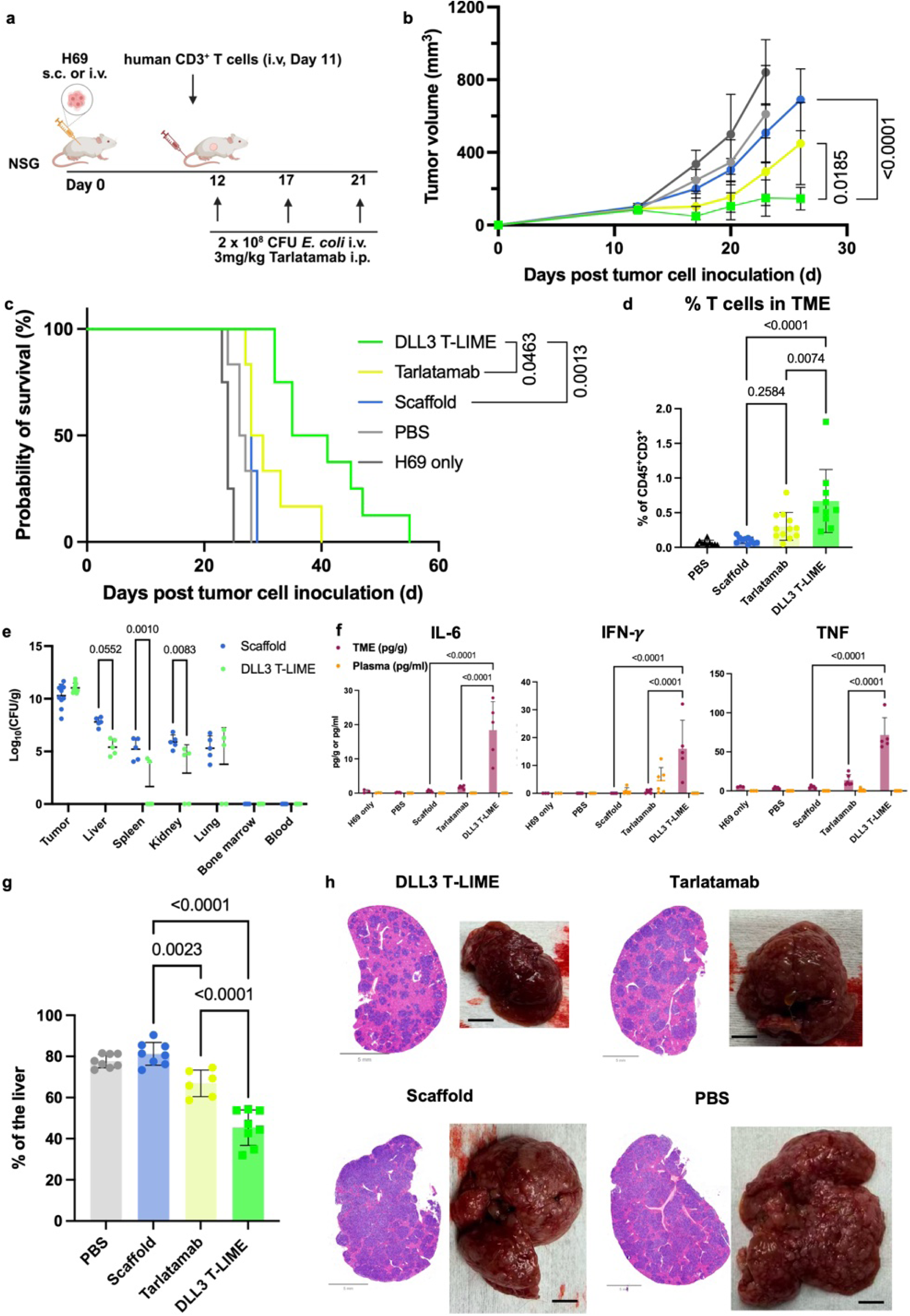
DLL3 T-LIME drives antitumor activity and localized immune activation in small-cell lung cancer xenograft models. **a**, Schematic of the small cell lung cancer (H69) xenograft model. H69 tumor cells were implanted subcutaneously (**b**, **c** for survival studies, **d**, **e**, **f** for tumor or organ profiling) or injected intravenously (**g**, **h** for metastasis), and human CD3^+^ T cells were administered intravenously (i.v., tail vein) on day 11. On day 12, mice were randomized to receive PBS (n = 6), scaffold bacteria (n = 6), DLL3 T-LIME (n = 8) or DLL3-targeting T-cell engager tarlatamab (n = 6). Bacterial treatments were administered i.v. at 2 × 10⁸ colony-forming units (CFU) per dose, and tarlatamab was administered intraperitoneally at 3 mg/kg on the indicated treatment days. **b**, **c**, Tumor-growth kinetics (**b**) and survival analysis (**c**) of H69 subcutaneous xenograft-bearing mice treated as in **a**. **d**–**f**, in a separate parallel cohort, mice were treated as in a and sacrificed on day 24 for mechanistic analyses. **d**, Frequency of human T cells in the tumor microenvironment after treatment. **e**, Biodistribution of scaffold bacteria and DLL3 T-LIME across tumor and major organs, quantified as CFU per gram of tissue. **f**, Cytokine analysis of tumor tissue (pg/g) and plasma (pg/ml), including IFN-γ, IL-6 and TNF, showing local and systemic cytokine responses after treatment. **g**, Quantification of liver metastatic tumor burden across treatment groups. **h**, Representative H&E-stained liver sections (scale bar = 5 mm) and gross liver images (scale bar = 5 mm). Data are presented as mean ± s.d. (**d**-**g**). Statistical significance was determined by two-way ANOVA (**b**), log-rank test (**c**) and one-way ANOVA (**e**-**g**).

DLL3 T-LIME markedly inhibited tumor growth relative to scaffold bacteria (p < 0.0001) and tarlatamab (p = 0.0185; **Fig. 3b**). Treatment also prolonged survival relative to scaffold and PBS controls and significantly extended survival compared with tarlatamab (p = 0.0463; **Fig. 3c**). Thus, DLL3 T-LIME achieved superior control of subcutaneous tumor growth and prolonged survival compared with the soluble engager.

To investigate the immune response associated with tumor control, we analyzed a parallel cohort treated with the same schedule and collected tumors and organs on day 24. DLL3 T-LIME increased the frequency of human T cells within the tumor microenvironment relative to both scaffold bacteria and tarlatamab (**Fig. 3d**). The distribution of naive, central memory and effector memory subsets was not significantly altered among intratumoral CD4^+^ or CD8^+^ T cells. However, DLL3 T-LIME increased the frequency of CD25^+^CD69^+^ co-expression in CD4^+^ T cells relative to all control groups. Among CD8^+^ T cells, CD25^+^CD69^+^ co-expression was increased relative to PBS and scaffold controls and was comparable to that induced by tarlatamab. Critically, DLL3 T-LIME reduced PD-1 expression in both intratumoral CD4^+^ and CD8^+^ T cells relative to tarlatamab (**Supplemental Fig. 13a-c** and **Supplemental Fig. 14a, b**). Together, these findings demonstrate that DLL3 T-LIME drives T cell accumulation and activation within the tumor while limiting an increase in the inhibitory PD-1 expression.

We next compared human T cell abundance and phenotype in peripheral blood, bone marrow, liver, spleen and lung. DLL3 T-LIME increased human T cell frequencies across several peripheral tissues, including blood, bone marrow, liver, spleen and lung. In contrast to the tumor, however, CD25 and CD69 expression was generally lower following DLL3 T-LIME than following tarlatamab in non-tumor tissues, particularly in the bone marrow, liver and lung (**Supplemental Fig. 15a-d**, **Supplemental Fig. 16a-e**, **Supplemental Fig. 17a-e** and **Supplemental Fig. 18a-e**). This compartment-specific pattern suggests that DLL3 T-LIME concentrates productive T cell activation within the tumor while limiting activation in peripheral organs.

Bacterial biodistribution provided a potential basis for this localized immune activity. Both scaffold bacteria and DLL3 T-LIME accumulated to more than 10^10^ colony-forming units (CFU) per gram of tumor, several orders (10^5^ - 10^6^) of magnitude above the burdens detected in normal tissues (**Fig. 3e**). Tumor bacterial burdens were comparable between DLL3 T-LIME and scaffold treated mice. However, DLL3 T-LIME burdens were significantly lower in the spleen and kidney, with a similar trend in the liver, conferring greater tumor-to-normal-tissue selectivity than scaffold bacteria alone (**Fig. 3e**). Thus, scFv display preserved bacterial tumor tropism and was associated with a more tumor-selective distribution of viable bacteria.

We measured soluble immune factors in tumor lysates and plasma. DLL3 T-LIME elicited marked intratumoral accumulation of pro-inflammatory cytokines IL-6, IFN-γ and TNF relative to the control treatment groups (**Fig. 3f**). Despite this local inflammatory response, the corresponding plasma cytokine levels remained low. By comparison, tarlatamab treatment was associated with higher circulating IFN-γ levels despite lower intratumoral cytokine concentrations. Broader cytokine profiling further demonstrated the distinct local and systemic response patterns across the treatment groups (**Supplemental Fig. 19a**). Together, these results show that DLL3 T-LIME induces robust intratumoral inflammatory responses with minimal systemic exposure, providing a mechanistic basis for its superior anti-tumor activity and potentially improved safety profile compared with conventional soluble engagers.

We also assessed DLL3 T-LIME in a metastatic H69 model established by intravenous tumor cell injection, in which metastatic burden developed predominantly in the liver (**Fig. 3a**). Mice were treated using the same schedule and analyzed on day 35. While DLL3 T-LIME and tarlatamab produced similar reductions in liver weight relative to scaffold treated mice (**Supplemental Fig. 20**), quantitative analysis showed significantly less tumor involvement following DLL3 T-LIME than either tarlatamab or control treatments (**Fig. 3g, h** and **Supplemental Figs. 21-24**). These findings show that DLL3 T-LIME achieves robust antitumor activity in a disseminated SCLC model and reduces hepatic metastatic burden more effectively than tarlatamab.

Together, these results demonstrate that DLL3 T-LIME effectively controls both subcutaneous and metastatic SCLC xenografts while concentrating bacterial accumulation, T cell activation and pro-inflammatory cytokine production within the tumor microenvironment. This spatially restricted immune response distinguishes DLL3 T-LIME from conventional soluble T cell engagers and supports the use of tumor-tropic bacteria to localize potent antitumor immunity.

### LIME drive durable, localized anti-tumor immunity in syngeneic lymphoma and pancreatic tumors

To extend LIME evaluation to immunocompetent syngeneic models, we first developed constructs capable of engaging murine effector cells. *E. coli* displaying an anti-murine CD3 scFv, either alone or in tandem with anti-murine CD19 or anti-murine MSLN scFvs, bound murine T cells and increased expression of the activation markers CD25 and CD69 in splenocyte cocultures (**Supplemental Fig. 25a-c**). Because few murine NK-cell-targeting scFvs were available, we screened human CD16-, NKG2D- and NKp46-targeting scFvs for cross-reactivity with murine NK cells (CD45^+^CD3^-^NK1.1^+^ cells). The NKG2D-targeting scFv and NKp46-targeting clone 2 showed the strongest cross-reactive binding, and N-LIME constructs incorporating these binders induced CD25 and CD69 expression in murine NK cells (**Supplemental Fig. 25d-f**). In three-cell cocultures containing murine splenocytes and either MSLN^+^ KPC pancreatic cancer cells or CD19^+^ A20 lymphoma, cognate T-LIME and N-LIME constructs increased tumor cell killing. Among the N-LIME designs, the NKG2D-targeting construct produced the most consistent cytotoxic activity and was selected for subsequent in vivo studies (**Supplemental Fig. 25g-i**).

We first evaluated CD19-targeting LIME in the syngeneic A20 B-cell lymphoma model. Immunocompetent BALB/c mice bearing subcutaneous A20 cells were treated intravenously with CD19 T-LIME, CD19 N-LIME or scaffold bacteria at 1 × 10^8^ CFU per dose, or with PBS, on days 10 and 14, when tumors had reached approximately 80-150 mm^3^. Anti-PD-1 (8 mg/kg, RMP1-14) was administered intraperitoneally on the same schedule, either alone or in combination with T-LIME or N-LIME (**Fig. 4a**). Both CD19 T-LIME and CD19 N-None of the animals demonstrated any evidence of obvious toxicity including weight loss. LIME markedly inhibited tumor growth relative to scaffold bacteria and PBS control (p < 0.0001; **Fig. 4b**) and conferred durable tumor-free survival in a subset of mice. CD19 T-LIME and CD19 N-LIME resulted in tumor-free survival in 3 of 6 and 2 of 6 mice, respectively, compared with 1 of 6 mice treated with anti-PD-1 alone. Combining anti-PD-1 with CD19 T-LIME or CD19 N-LIME further increased the tumor-free fraction to 6 of 7 and 5 of 7 mice, respectively (**Fig. 4c**).

**Fig. 4.**
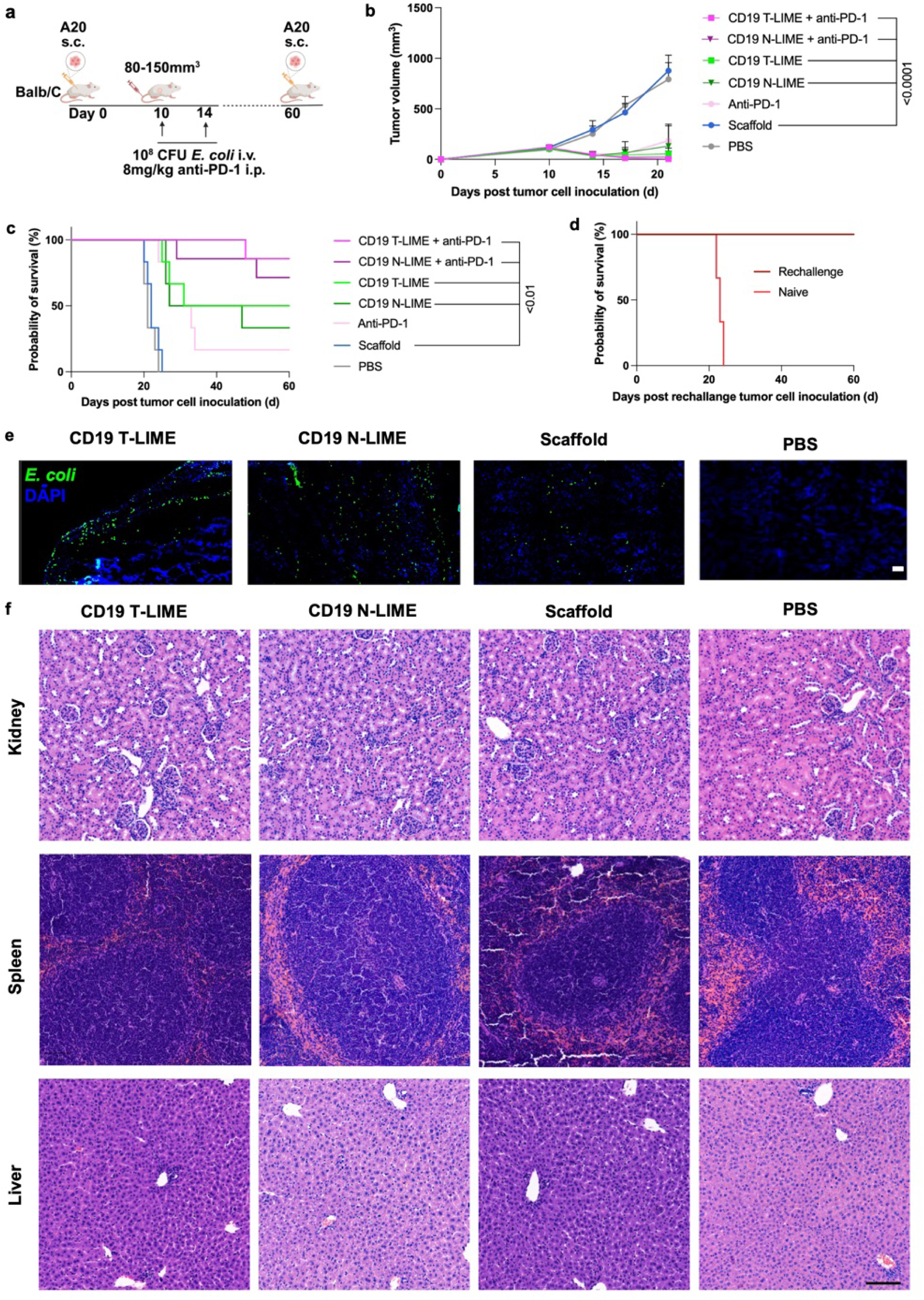
LIME combined with anti-PD-1 treatment induces durable antitumor responses and immunological memory with a favorable safety profile in a syngeneic lymphoma model. **a**, BALB/c mice were subcutaneously implanted with 1 × 10⁶ CD19^+^ A20 B-cell lymphoma cells. Beginning on day 10, mice received i.v. injections of PBS (n = 6), Scaffold (n = 6), anti-PD-1 (n = 6), CD19 T-LIME (n = 6), CD19 N-LIME (n = 6), CD19 T-LIME + anti-PD-1 (n = 7), or CD19 N-LIME + anti-PD-1 (n = 7) on days 10 and 14. On day 60, 17 tumor-free mice were rechallenged with 1 × 10⁶ CD19^+^ A20 B-cell lymphoma cells in the contralateral flank; the control naïve group included the age and gender matched mice without tumor injections. **b**, Tumor growth kinetics across the indicated treatment groups. **c**, Kaplan–Meier survival analysis of the indicated treatment groups. **d**, Kaplan–Meier survival analysis following tumor rechallenge. **e**, Representative fluorescence microscopy images showing intratumoral distribution of bacteria across treatment groups. Tumor cryosections were stained with antibodies against *E. coli* (green) and with DAPI (blue), pseudocolor applied. Images were acquired using a Leica THUNDER Imager. Scale bar represents 100 μm. **f**, Representative hematoxylin and eosin (H&E)-stained sections of kidney, spleen and liver from the indicated treatment groups. Scale bar represents 100 μm. Statistical significance was determined using two-way ANOVA with Tukey’s post hoc test in **b** and log-rank test in **c** and **d**. Data shown are representative of (**e**, **f**) or combined from two independent biological replicates (**b**-**d**) and are presented as mean ± s.d. in **b**.

To determine whether tumor clearance generated protective immunological memory, all 17 tumor-free mice were rechallenged on day 60 with A20 cells implanted in the contralateral flank. Every previously treated mouse remained tumor-free, whereas tumors grew in all treatment-naive controls (**Fig. 4d**). Thus, LIME treatment, alone or with anti-PD-1, induced durable protection against tumor rechallenge.

We next assessed bacterial localization and tissue histology in a parallel cohort collected on day 21. *E. coli* signal was readily detected within A20 tumors from mice treated with CD19 T-LIME, CD19 N-LIME or scaffold bacteria, confirming bacterial accumulation in the lymphoma microenvironment (**Fig. 4e** and **Supplemental Fig. 26**). H&E-stained kidney, spleen and liver sections showed no overt treatment-associated histopathological abnormalities relative to PBS-treated controls (**Fig. 4f** and **Supplemental Figs. 27-29**). Together, these findings demonstrate durable antitumor activity in an immunocompetent lymphoma model without evident injury to the organs examined.

We then investigated MSLN-targeting LIME in the poorly immunogenic KPC pancreatic ductal adenocarcinoma (PDAC) model. C57BL/6 mice bearing established subcutaneous KPC tumors (100-150 mm^3^) received four intravenous doses of MSLN T-LIME, MSLN N-LIME or scaffold bacteria at 2 × 10^8^ CFU per dose, or PBS, on days 10, 13, 16 and 20. Tumors and peripheral tissues were collected on day 23 for immune profiling (**Fig. 5a**). Both MSLN T-LIME and MSLN N-LIME strongly suppressed tumor growth relative to scaffold and PBS controls (p < 0.0001; **Fig. 5b**). Biodistribution analysis showed high viable bacterial colonization in tumors, whereas burdens in the liver, spleen and kidney were several orders of magnitude lower (100 – 1000-fold, **Fig. 5c**). Confocal imaging further demonstrated intratumoral colocalization of *E. coli*, CD3^+^ T cells and MSLN^+^ tumor regions following LIME treatment (**Fig. 5d** and **Supplemental Fig. 30**). These results show that MSLN-targeting LIME retains the tumor-selective distribution of the bacterial chassis in an immunocompetent PDAC model.

**Fig. 5.**
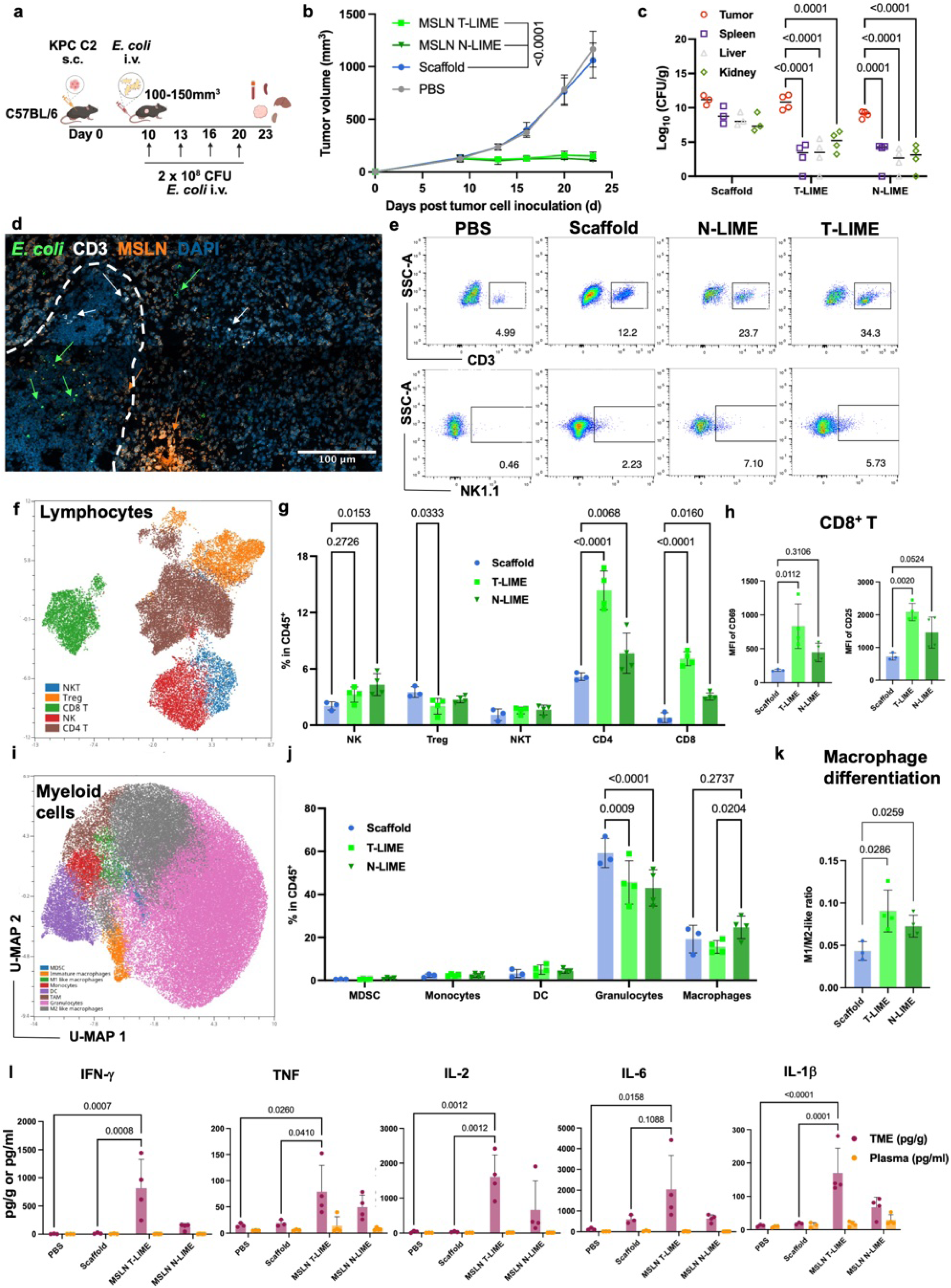
Tumor homing LIME remodel the tumor microenvironment and drive coordinated innate and adaptive immune activation with minimal systemic inflammation. **a**, Experimental schema. C57BL/6 mice were subcutaneously (s.c.) implanted with 1 × 10⁶ KPC C2 cells in the flank. Beginning on day 10, mice received four intravenous (i.v.) injections of PBS (n = 3), Scaffold (n = 3), MSLN T-LIME (n = 4), or MSLN N-LIME (n = 4) on days 10, 13, 16, and 20. Mice were euthanized on day 23 for tissue collection and downstream analyses. **b**, Tumor growth kinetics of the indicated treatment groups. **c**, Quantification of bacterial biodistribution in tumor, liver, kidney, and spleen tissues. Colony-Forming Units (CFU g⁻¹) indicate the number of bacteria per gram of indicated tissue. **d**, Representative confocal images showing intratumoral localization of bacteria, T cells, and tumor cells following T-LIME treatment. The white dash curve indicates the broader of the necrotic region. Tumor cryosections were stained with antibodies for *E. coli* (green), CD3 (white), mesothelin (MSLN, orange), and with DAPI (blue), pseudocolor was applied. Images were acquired using a ZEISS LSM 980 confocal microscope. Scale bar represents 100 μm. **e**, Representative flow cytometry plots demonstrating intratumoral infiltration of CD3^+^ T cells and NK1.1^+^ cells across treatment groups. **f**, **g**, Uniform Manifold Approximation and Projection (UMAP) visualization (**f**) and quantitative analysis (**g**) of lymphocyte subpopulations (excluding B cells) among CD45^+^ intratumoral immune cells. **h**, Activation status of intratumoral CD8^+^ T cells, quantified by CD69 and CD25 expression. **i**,**j**, UMAP visualization (**i**) and quantification (**j**) of myeloid cell subpopulations within CD45^+^ tumor-infiltrating immune cells. **k**, Macrophage polarization quantified as the ratio of M1-like (CD45^+^CD11b^+^F4/80^+^MHC-II^+^CD206^-^) to M2-like (CD45^+^CD11b^+^F4/80^+^MHC-II^-^CD206^+^) populations. **l**, Luminex-based immune relevant soluble marker profiling of tumor lysates and plasma collected from the KPC C2 tumor-bearing mice at the study end point (day 23). Statistical significance was determined using one-way or two-way ANOVA with Tukey’s post hoc test in **g**, **h**, **j**, **k** and **l**. Data are presented as mean ± s.d. in **g**, **h**, **j**, **k** and **l.**

Flow cytometry revealed increased intratumoral accumulation of CD3^+^ T cells and NK1.1^+^ cells following LIME treatment (**Fig. 5e** and **Supplemental Fig. 31**). Immune profiling showed that MSLN N-LIME increased the frequency of NK cells, whereas both MSLN T-LIME and MSLN N-LIME increased CD4^+^ T cell and CD8^+^ T cell populations. MSLN T-LIME also reduced the frequency of regulatory T cells (Treg; **Fig. 5f, g**). Phenotypic analysis showed that MSLN T-LIME strongly increased CD25 and CD69 expression on intratumoral CD8^+^ T cells (**Fig. 5h**). Across CD4^+^ T cell, CD8^+^ T cell and NK cell compartments, LIME also enhanced selected activation and proliferation markers; PD-1 expression was reduced in intratumoral T cell subsets relative to scaffold controls (**Extended Data Fig. 6a-c**). By contrast, the overall frequency and composition of the intratumoral B cell lineage were not detectably altered (**Supplemental Fig. 32a-c**).

LIME treatment also remodeled the intratumoral myeloid compartment. Both MSLN T-LIME and MSLN N-LIME treatment reduced granulocytes frequencies (**Fig. 5i, j** and **Supplemental Fig. 33a, b**). Although total macrophage abundance was not consistently changed, both treatments increased the ratio of M1-like to M2-like macrophages, consistent with a shift toward a pro-inflammatory phenotype (**Fig. 5k**).

Immune profiling of the spleen, tumor draining lymph nodes (TDLN) and peripheral blood (PBMC) revealed selective peripheral responses rather than broad systemic activation. Overall splenic lymphocyte composition remained largely stable, although LIME increased CD4^+^ T cell proliferation and altered selected splenic B cell, myeloid and dendritic cell subsets. In tumor draining lymph nodes, the major lymphocyte and B cell compartments were generally preserved, while proliferative responses were detected in CD4^+^ T cells, CD8^+^ T cells and NK cells together with modest changes in macrophage and dendritic cell (DC) populations. In PBMC, LIME increased CD4^+^ and CD8^+^ T cell frequencies, whereas activation and inhibitory marker profiles remained broadly comparable across treatment groups. Circulating myeloid cell composition was also largely preserved, with an increase in cDC1 frequency but no significant change in the cDC1-to-cDC2 ratio **(Supplemental Figs. 34-41**). These findings indicate that LIME induces coordinated but compartment specific immune remodeling, with the strongest activation concentrated in the tumor microenvironment.

Finally, Luminex profiling revealed marked increases in IFN-γ, TNF, IL-2, IL-6 and IL-1β within tumors following MSLN T-LIME treatment, with more moderate induction following MSLN N-LIME (**Fig. 5l**). Additional cytokine and chemokine analysis identified treatment-associated changes in intratumoral G-CSF, IL-5, IL-7, IL-9, IL-10, IL-12p70, VEGF, CXCL10, CXCL1 and other immune mediators. For most analytes, corresponding plasma concentrations remained substantially lower than those in tumor lysates (**Supplemental Fig. 42**). Thus, LIME generated a robust local inflammatory response without comparably broad systemic cytokine induction.

Together, these results establish that T-LIME and N-LIME are active in immunocompetent syngeneic tumor models including otherwise highly immunosuppressive PDAC model. LIME produced durable tumor control and immunological memory in lymphoma, suppressed established PDAC tumors, and coordinated lymphoid and myeloid remodeling within the tumor microenvironment. Tumor selective bacterial distribution limited systemic cytokine induction and the absence of overt histopathological injury in the examined organs further support a favorable preclinical safety profile.

### Pan-RAS inhibition enhances LIME efficacy and reveals PD-L1-associated myeloid remodeling in pancreatic cancer

We next investigated whether LIME could be combined with complementary therapies to improve control of pancreatic ductal adenocarcinoma (PDAC). Although anti-PD-1 enhanced LIME activity in the A20 lymphoma model, adding anti-PD-1 to MSLN T-LIME or MSLN N-LIME did not further improve tumor control or survival in mice bearing KPC tumors (**Extended Data Fig. 7a-c**). We therefore evaluated RMC-7977, a multi-selective RAS(ON) inhibitor, as an alternative combination partner.

C57BL/6 mice bearing subcutaneous KPC C2 tumors were treated with MSLN T-LIME, MSLN N-LIME, scaffold bacteria or PBS on days 10, 14, 17 and 21. RMC-7977 (RASi; 25 mg/kg) was administered orally on the same schedule, either alone or in combination with T-LIME or N-LIME (**Fig. 6a**). Both combination regimens produced stronger tumor control than either LIME or RASi monotherapy (**Fig. 6b**). This improvement was accompanied by prolonged survival, with median survival increasing to 52.5-56 days following combination treatment, compared with 29-32 days following the corresponding monotherapies (MSLN T-LIME + RASi vs Scaffold, p = 0.0006) (**Fig. 6c**). Thus, RAS inhibition enhanced the antitumor efficacy of both T cell- and NK/T cell-engaging LIME.

**Fig. 6.**
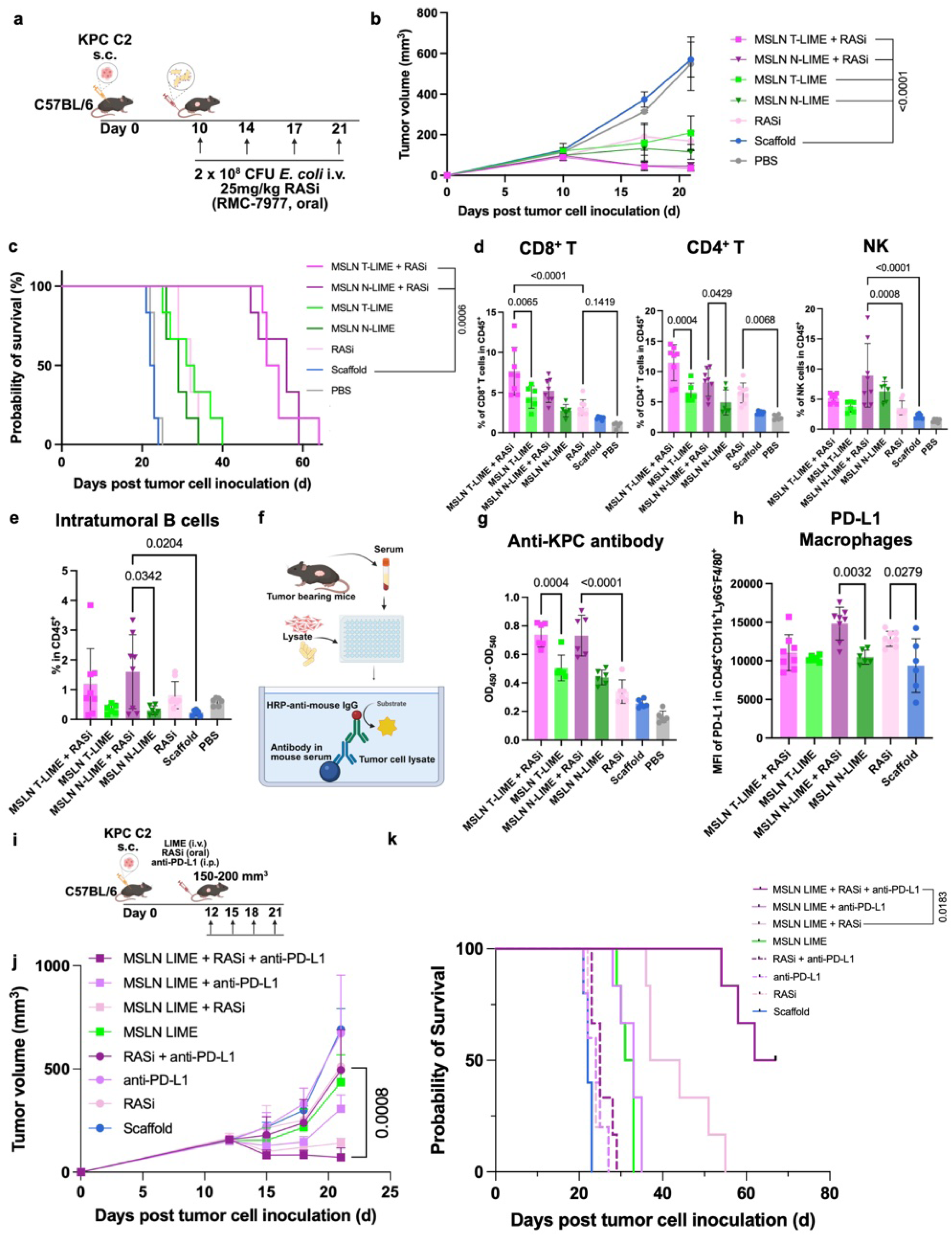
RAS inhibition and anti-PD-L1 therapy enhance MSLN-targeting LIME therapy in a syngeneic pancreatic cancer model. **a**, Experimental schema. C57BL/6 mice were subcutaneously implanted with KPC C2 pancreatic cancer cells and treated intravenously with 2 × 10⁸ CFU E. coli displaying MSLN-targeting T-LIME or N-LIME, scaffold control, or PBS, with or without oral RAS inhibitor treatment (RASi; RMC-7977, 25 mg/kg) on the indicated days. **b**, Tumor growth kinetics of KPC C2 tumor-bearing mice treated with PBS, Scaffold, RAS inhibitor, MSLN T-LIME, MSLN N-LIME, MSLN T-LIME + RAS inhibitor, or MSLN N-LIME + RAS inhibitor. **c**, Kaplan–Meier survival analysis of the indicated treatment groups. **d**, Frequencies of intratumoral CD8^+^ T cells, CD4^+^ T cells, and NK cells among CD45^+^ immune cells following treatment. **e**, Frequency of intratumoral B cells among CD45^+^ immune cells following treatment. **f**, Schematic of the serum antibody-binding assay used to detect host antibody responses against bacterial lysate or KPC tumor-cell lysate. **g**, Serum anti-KPC antibody responses in the indicated treatment groups, quantified by ELISA. **h**, Expression of PD-L1 on intratumoral CD45^+^CD11b^+^Ly6G^−^F4/80^+^ myeloid cells (macrophages) following treatment, quantified as mean fluorescence intensity (MFI). **i**, Experimental schema for combination therapy with LIME (MSLN N-LIME + MSLN T-LIME), RAS inhibitor, and anti-PD-L1 (clone 10F.9G2) treatment in KPC C2 tumor-bearing mice. Mice were treated with intravenous LIME, oral RAS inhibitor, intraperitoneal anti-PD-L1, or the indicated combinations after tumors reached 150– 200 mm³. **j**, Tumor growth kinetics of mice treated as in **i**. **k**, Kaplan–Meier survival analysis of mice treated as in **i**. Data are presented as mean ± s.d. (**b**, **d**, **e**, **g**, **h**, **j**).

To define the mechanisms underlying this improved activity, we analyzed tumors, blood and peripheral organs from a parallel cohort collected on day 24. Viable bacteria remained strongly enriched in tumors, whereas bacterial burden was substantially lower in the liver, spleen and kidney and were undetectable in TDLNs, bone marrow and blood. RMC-7977 did not measurably alter this distribution, indicating that RAS inhibition preserved the tumor selective localization of LIME (**Supplemental Fig. 43**).

Flow cytometric analysis revealed that RAS inhibition further remodeled the intratumoral lymphocyte compartment. Adding RASi to MSLN T-LIME increased both CD8^+^ and CD4^+^ T cell frequencies relative to T-LIME alone, whereas adding RASi to MSLN N-LIME increased CD4^+^ T cell abundance and maintained the elevated NK cell population induced by N-LIME (**Fig. 6d**). Across the T cell-engaging and NK cell-engaging regimens, LIME reduced intratumoral regulatory T cell abundance and increased effector memory T cell (central memory: CD44^hi^CD62L^hi^; effector memory: CD44^hi^CD62L^lo^) populations. LIME-containing treatments also increased selected activation, proliferation and cytotoxicity markers, including CD25, CD69, Ki67 and granzyme B, while reducing PD-1 expression in intratumoral T cell subsets relative to scaffold or PBS controls. NK cells similarly displayed increased Ki67, CD25, CD69 and granzyme B following LIME treatment, although the magnitude of these changes varied between T-LIME and N-LIME and with RASi coadministration (**Extended Data Figs. 8, 9** and **Supplemental Fig. 44**). These findings indicate that RAS inhibition complements LIME by increasing intratumoral lymphocyte accumulation while preserving the activated effector phenotype induced by LIME.

Immune changes outside the tumor were more limited. The overall frequencies of CD8^+^ T cells, CD4^+^ T cells, NK cells, NKT cells and regulatory T cells in TDLNs were broadly similar across groups. Nevertheless, LIME-containing regimens increased selected proliferative and activation-associated features, including Ki67 expression in CD8^+^ T cells and NK cells and CD69, Ki67 and effector memory differentiation in CD4^+^ T cells (**Supplemental Figs. 45-48**). In peripheral blood, T-LIME-containing groups showed increased CD8^+^ and CD4^+^ T cell frequencies, whereas lymphocyte composition and phenotypic markers in the spleen and bone marrow remained largely unchanged (**Supplemental Figs. 49-60**). Together, these data show that the most pronounced lymphocyte remodeling was concentrated within the tumor microenvironment rather than broadly distributed across peripheral immune compartments.

The combination regimens also altered humoral immune features. Intratumoral B cell abundance was higher following MSLN T-LIME plus RASi and MSLN N-LIME plus RASi than following the corresponding LIME monotherapies (**Fig. 6e**). To determine whether this increase was associated with tumor-reactive antibody production, we measured serum IgG binding to KPC tumor cell lysate by ELISA (**Fig. 6f**). Both combination groups exhibited significantly higher titers of KPC lysate-reactive IgG than the corresponding LIME monotherapy groups (**Fig. 6g**). In the bone marrow, conventional B cells were reduced across bacteria treated groups, whereas the scaffold group showed a selective reduction in plasma cells. Major B cell populations in the blood, TDLNs and spleen otherwise remained broadly comparable across treatments (**Supplemental Fig. 61a-d**). These results suggest that combined LIME and RAS inhibition promotes a tumor reactive humoral response, although the functional contribution of these antibodies to tumor control remains to be established.

We next examined the myeloid compartment. Total intratumoral monocyte, dendritic cell, macrophage and granulocyte frequencies were not uniformly altered by the combination treatments. However, RASi containing LIME regimens increased PD-L1 expression across several intratumoral myeloid populations, including dendritic cells, macrophages, granulocytes and monocytes (**Fig. 6h**, **Extended Data Fig. 10** and **Supplemental Figs. 62**). These changes were accompanied by context-dependent reductions in CD80/CD86 co-expression and Ki67, particularly within macrophage, granulocyte and monocyte populations. By contrast, myeloid composition and phenotype in the spleen, peripheral blood, bone marrow and TDLNs showed no consistent treatment wide pattern. (**Supplemental Figs. 63-69**). Thus, the combination generated a predominantly intratumoral myeloid response characterized by increased PD-L1, suggesting a potential adaptive resistance mechanism.

Multiplex cytokine analysis further revealed a tumor predominant inflammatory response. LIME-containing regimens, particularly MSLN T-LIME plus RASi, increased multiple inflammatory mediators within tumors, including IFN-γ, IL-2, IL-6, IL-12p70, IL-15, IL-28B and CXCL10. For most analytes, corresponding plasma concentrations remained markedly lower than concentrations in tumor lysates (**Supplemental Figs. 70-72**). These data indicate that combining LIME with RAS inhibition maintains localized immune activation without comparably broad systemic cytokine induction.

The induction of PD-L1 on intratumoral myeloid cells seen above provided a rationale for evaluating PD-L1 blockade with our treatment regimens. For this, we treated mice bearing larger established KPC tumors of 150-200 mm^3^ with LIME, formulated as an equal mixture of MSLN T-LIME and MSLN N-LIME (10^8^ CFU each), RMC-7977 and anti-PD-L1 (clone 10F.9G2), either individually, in pairwise combinations or as a three-agent regimen (**Fig. 6i**). The triple combination produced the strongest tumor-growth inhibition and longest survival among the regimens tested, with 66.6% of mice demonstrating survival beyond day 60 (**Fig. 6j, k**). This result is consistent with PD-L1 blockade countering a resistance program associated with combined LIME and RAS inhibition and further improving activity against large established PDAC tumors.

Together, these results demonstrate that RAS inhibition enhances LIME efficacy through coordinated remodeling of adaptive immunity, including increased intratumoral T cell accumulation and elevated tumor-reactive IgG. At the same time, the combination induces a PD-L1-high intratumoral myeloid state that may constrain therapeutic activity. Addition of anti-PD-L1 further improved tumor control, supporting a three-component strategy that couples tumor-localized immune engagement and oncogenic RAS inhibition with blockade of a candidate adaptive myeloid checkpoint response.

## Discussions

Here, we developed live immune modulating engagers (LIME), a modular platform in which non-pathogenic, tumor-tropic *E. coli* display tandem scFvs that bind a tumor-associated antigen and an immune effector receptor. Rather than functioning solely as carriers for a secreted therapeutic payload, LIME bacteria provide a multivalent cell-binding surface that directly recruits T cells or NK cells to tumor cells. Across cell lines, patient-derived PDAC organoids, xenografts and immunocompetent syngeneic models, LIME promoted effector cell activation, tumor cell killing and tumor control while concentrating bacterial accumulation and inflammatory activity within tumors. To our knowledge, these findings establish the first broadly modular living bacterial platform that functions as a surface displayed immune engager.

A central feature of LIME is its modularity. Exchange of either scFv arm enabled targeting of diverse tumor antigens and engagement of CD3, CD16, NKG2D or NKp46. Both T-LIME and N-LIME were active across multiple tumor types, and CLDN18.2 T-LIME retained activity in patient derived PDAC organoids. The differences observed among NK cell targeting receptors and even between independent clones directed against the same receptor emphasize, however, that the platform is not entirely target agnostic. Binder affinity, epitope location, surface expression level, orientation and linker geometry are all likely to influence bacterial attachment, receptor clustering and cytotoxic activity. Thus, the plug-and-play architecture enables rapid substitution and screening, but individual constructs will still require empirical optimization. Although this study used scFvs, the same display framework could in principle accommodate nanobodies^29^, minibinders^30^ or other compact binding domains (e.g. CD70^31^), potentially expanding the accessible antigen and effector receptor space.

The transcriptional analyses further distinguish LIME from either component of the system alone. Scaffold bacteria induced a broad inflammatory response in human T cells, including interferon-gamma, TNF-NF-kB and IL-2-STAT5 programs, but this response was accompanied by negative enrichment of the G2M-checkpoint program (**Extended Data Fig. 5a-d**). This pattern suggests that bacterial sensing alone can activate inflammatory pathways without fully supporting a proliferative effector state. DLL3 T-LIME integrated this bacteria-derived inflammation with stronger cytokine, metabolic and cell-cycle programs, including mTORC1 signaling, glycolysis, MYC targets and G2M-checkpoint activity (**Fig. 2d, e**). LIME therefore did not simply amplify bacterial inflammation; it converted that response into a transcriptional state more consistent with productive T cell engagement and expansion. The multivalent display of many binding molecules on each bacterium may contribute to this activity by increasing avidity and receptor clustering^32^.

The DLL3^+^ SCLC models provided a direct comparison between LIME and the approved soluble T cell engager tarlatamab^13, 33^. DLL3 T-LIME produced stronger control of both subcutaneous and metastatic H69 tumors and prolonged survival compared with tarlatamab. SCLC commonly exhibits impaired MHC-I antigen presentation, including defects described in H69 cells^34^. Because CD3-directed engagers recruit T cells independently of peptide-MHC recognition, the activity of DLL3 T-LIME in this setting supports its potential to bypass one important mechanism of resistance to conventional antigen-specific T cell responses^12, 35, 36^.

The spatial distribution of the response may be as important as its magnitude. DLL3 T-LIME increased intratumoral T cell accumulation and inflammatory cytokines, whereas activation markers and cytokine concentrations remained lower in most peripheral tissues than after tarlatamab treatment. Similar tumor-predominant immune remodeling and cytokine production were observed in immunocompetent KPC tumors. These findings are relevant because CRS is a recognized toxicity of systemic T cell engagers, including tarlatamab^12, 13, 33^. Therefore, we also establish that LIME has minimal risk of causing CRS or sepsis. In summary, the results support that bacterial tumor tropism can widen the therapeutic window by spatially restricting potent immune engagement with safety profile.

The syngeneic models showed that LIME can operate within an intact immune system and engage immune mechanisms beyond the directly targeted effector population. In A20 lymphoma, CD19 T-LIME and CD19 N-LIME generated durable tumor-free survival in a subset of mice, and every tumor-free mouse resisted subsequent A20 rechallenge. Combining LIME with anti-PD-1 further increased the tumor-free fraction, demonstrating that localized engagement can cooperate with checkpoint blockade in an immunotherapy responsive setting. In the poorly immunogenic KPC model, MSLN T-LIME and MSLN N-LIME increased intratumoral T cell and NK cell populations, reduced regulatory T cells and granulocytes, shifted macrophages toward an M1-like phenotype and induced tumor localized inflammatory cytokines. Together, these changes are consistent with conversion of an immune poor PDAC microenvironment toward a more inflamed state.

The comparable efficacy of T-LIME and N-LIME in several syngeneic experiments also broadens the biological scope of the platform. NKG2D is expressed not only by NK cells but also by NKT cells, γ-δ T cells and subsets of activated CD8^+^ T cells^37^. NKG2D directed N-LIME may therefore recruit a mixed population of cytotoxic lymphocytes rather than functioning as a strictly NK cell specific engager. Similarly, the reduction in intratumoral granulocytes following LIME treatment may favor T cell accumulation, as tumor-associated neutrophils can suppress T cell function in some settings^38^.

Combining LIME with RAS inhibition further improved control of established KPC tumors and approximately doubled median survival relative to the corresponding monotherapies. This combination is clinically relevant because RAS alterations dominate PDAC biology and the RAS(ON) multi-selective inhibitor daraxonrasib has demonstrated clinical activity in RAS-mutant PDAC^39^. RMC-7977 increased intratumoral T cell abundance while preserving the activated and proliferative phenotypes associated with LIME. The combination also increased intratumoral B cells and serum IgG reactive to KPC cell lysate, suggesting that direct effector cell engagement can propagate into a broader adaptive response.

RAS inhibition also exposed a candidate resistance mechanism. LIME plus RMC-7977 increased PD-L1 expression across several intratumoral myeloid populations, motivating addition of PD-L1 blockade. The three-agent regimen produced the strongest tumor control and longest survival in mice bearing large established KPC tumors. Although a formal interaction analysis will be needed before classifying these effects as pharmacological synergy, the findings support a model in which RAS inhibition expands or recruits antitumor lymphocytes while simultaneously inducing a PD-L1-associated adaptive myeloid response. A recent study similarly found that oncogenic RAS inhibition increased T cell infiltration in PDAC and created a therapeutic window for checkpoint modulation; in that setting, anti-CTLA-4, but not anti-PD-1, promoted durable immunity and functional tertiary lymphoid structures^40^. Together with our data, these observations suggest that the optimal checkpoint partner for RAS directed therapy is likely to depend on the cellular source and anatomical organization of the inhibitory response. Direct comparison of PD-L1, CTLA-4 and other myeloid- or Treg-directed interventions will be informative.

LIME may also address a major limitation of conventional macromolecular therapies in solid tumors: heterogeneous intratumoral delivery. Spatial pharmacology studies in human PDAC have identified periostin-rich extracellular matrix and FAP-positive fibroblast neighborhoods associated with reduced antibody delivery^11^. In contrast, tumor-tropic bacteria can enter permissive tumor regions, proliferate locally and sustain a high intratumoral burden^27, 41–44^. Across the models examined here, LIME retained robust tumor colonization and, in several settings, showed reduced recovery from normal organs relative to scaffold bacteria. Surface displayed binders may increase tumor retention by attaching bacteria to antigen positive cells, an interpretation consistent with recent ligand-directed bacterial surface engineering studies^45^.

The physical mechanism by which a micrometer-scale bacterium supports engager mediated cytotoxicity remains an important open question. Conventional T cell engagers are thought to favor close membrane apposition, with productive synaptic distances on the order of approximately 13 nm^46^, whereas an *E. coli* cell is roughly 1-2 micrometers in size. LIME may therefore not form a canonical tripartite synapse in which the entire bacterium occupies the T cell-tumor cell interface. Instead, surface tethered scFvs may create local receptor clusters, stabilize multicellular contacts or recruit effector and tumor cells to distinct regions of the bacterial surface, after which nanoscale cell-cell contacts form adjacent to the bacterium. Super resolution or electron microscopy, live cell membrane imaging and receptor proximal signaling measurements will be required to define whether LIME induces a conventional cytolytic synapse, an adjacent synapse or a distinct multivalent signaling structure.

In summary, LIME combines the programmability of a bispecific immune engager with the tumor-seeking and locally amplifying behavior of a living bacterial therapeutic. The platform is modular across tumor antigens and effector-cell receptors, induces a transcriptionally distinct effector state and supports localized antitumor immunity in both xenograft and immunocompetent models. Its activity with RAS inhibition and checkpoint blockade further illustrates how living engagers can be integrated with targeted and immunomodulatory therapies.

With additional mechanistic, safety and translational development, LIME offers a strategy for concentrating otherwise systemically toxic immune engager activity within solid tumors while coordinating T cell, NK cell, myeloid and humoral antitumor responses.

## Methods

This study complied with all relevant ethical regulations. All animal studies were performed in accordance with federal, state and institutional guidelines and were approved by the Institutional Animal Care and Use Committee of Dana-Farber Cancer Institute (DFCI; protocol 24-009). De-identified leukapheresis collars from anonymous healthy donors were obtained from the Crimson Core at Brigham and Women’s Hospital (resource identifier T0197) and used to isolate primary human T and natural killer (NK) cells.

### Cell culture

Cell lines, sources and culture media are listed in **Supplementary Table S1**. Unless otherwise indicated, cells were maintained at 37 °C in a humidified incubator with 5% CO_2_ and were used between passages 2 and 10.

Patient-derived PDAC organoids were maintained as three-dimensional cultures in Matrigel (BD Biosciences; #354234) and organoid PDAC (OPAC) medium. Basal medium consisted of Advanced DMEM/F12 supplemented with 10 mM HEPES, 2 mM GlutaMAX, 100 U ml^-1^ penicillin-streptomycin and 0.1 mg ml^-1^ Primocin. Complete OPAC medium additionally contained 1x R-spondin conditioned medium, 1x Wnt3A conditioned medium, 50 ng ml^-1^ EGF, 100 ng ml^-1^ Noggin, 100 ng ml^-1^ FGF10, 500 nM A83-01, 1x B27 supplement, 10 mM nicotinamide, 10 nM gastrin and 1.25 mM N-acetylcysteine. Organoids were passaged at 80-90% confluence. Matrigel domes were washed once with PBS, disrupted in pre-warmed TrypLE and incubated at 37 °C until dissociated. TrypLE was quenched with cold wash medium, and cells were collected by centrifugation at 1,500 r.p.m. for 5 min. Organoids were resuspended in Matrigel, plated as 50 ul domes and allowed to polymerize for 20-30 min at 37 °C before addition of 500 ul complete OPAC medium per well of a 24-well plate.

### T and NK cell isolation and culture

Peripheral blood mononuclear cells (PBMCs) were isolated from healthy donor leukapheresis collars by Ficoll-Paque density-gradient centrifugation. T cells were enriched using the EasySep Human CD3 Positive Selection Kit II (STEMCELL Technologies, 17851) according to the manufacturer’s instructions. For activation, 5 × 10^6^ T cells were cultured in 5 ml TexMACS medium containing 50 U ml^-1^ recombinant human IL-2 (Miltenyi Biotec, 130-114-429) and 50 ul TransAct reagent (Miltenyi Biotec, 130-111-160). After 3 days, cells were washed with sterile PBS and maintained in TexMACS medium supplemented with 50 U ml^-1^ recombinant human IL-2 unless otherwise indicated.

NK cells were enriched from PBMCs using the RosetteSep Human NK Cell Enrichment Kit (STEMCELL Technologies, 15065). Purified NK cells were cultured in NK MACS medium (Miltenyi Biotec) supplemented with 5% human serum and 500 U ml^-1^ recombinant human IL-2.

### Preparation of murine splenocytes

Spleens were collected from untreated BALB/c or C57BL/6J mice, mechanically dissociated and passed through a 70-um cell strainer. Red blood cells were lysed with ammonium-chloride-potassium (ACK) buffer (Gibco, A1049201). Splenocytes were washed and resuspended in complete RPMI medium before coculture. For murine immune cell binding and activation assays, splenocytes were incubated with the indicated bacteria at a bacteria-to-immune-cell ratio of 100:1 for 48 h. For cytotoxicity assays, A20 or KPC C2 tumor cells, splenocytes and the indicated LIME or control bacteria were cocultured at a ratio of 1:1:100 for 48 h.

### Bacterial strains and culture

Non-pathogenic *E. coli* K-12 NEB 5-alpha carrying the indicated surface display plasmids was maintained in lysogeny broth (LB) under aerobic conditions. Kanamycin was added at 50 ug ml^-1^ for plasmids containing the corresponding resistance cassette. For all cell culture and animal experiments, overnight bacterial cultures were diluted tenfold into fresh selective LB and induced as described below. Bacterial concentrations were estimated from optical density at 600 nm (OD600), using 1 OD600 unit = 8 × 10^8^ colony-forming units (CFU) ml^-1^, and confirmed by plating when indicated.

### Plasmid construction and bacterial surface display

For the surface display of LIME, we used the following plasmids: pLyGo-Ec-7 (Addgene, 163135)^47^, pDSG323 (Addgene, 115594)^48^ and pDS861 (from Quintara Biosciences) with Lpp-OmpA (OmpA)^27, 28, 47^, YiaT232 or YiaT181^49^. DNA sequences for the encoding cytokines including 3× GGGGS linker between scaffolds and the cytokines with DYKDDDDK-tag (FLAG-tag) in the N terminus and Myc-tag in the C terminus were inserted between two SapI sites for pLyGo-Ec-7, between SpeI and PstI sites for pDSG323 and between NotI and BamHI sites for pDS861-RhaR-RhaS-P_rha_ by NEBuilder HiFi DNA Assembly Master Mix (New England Biolabs (NEB), M5520AVIAL). *E. coli* NEB 5-alpha with the correct plasmid were inoculated in fresh LB medium with 50 μg mL^−1^ kanamycin. After overnight culture in a shaker (37°C, 250 rpm unless otherwise indicated), bacterial suspensions were diluted by 10-fold in the fresh LB with 50 μg mL^−1^ kanamycin and 10 mM L-rhamnose (pLyGo-Ec-7 and pDSG861 derived plasmids) and induced at 25 °C, 200rpm for 16-24h or or 100 ng mL^−1^ anhydrotetracycline (aTc; pDSG323 derived) and induced at 25 °C, 200rpm for 16-24h. For bacterial surface display verification, 50 μl of bacterial suspension was collected, washed once with PBS and then incubated with anti-DYKDDDDK Tag Antibody (BioLegend, 637315) in PBS with a concentration of 500:1 for 15 min at room temperature, washed two times and then suspended in PBS for flow cytometry analysis. All plasmids were validated by nanopore based whole plasmid sequencing. The structures of various fusion proteins were predicted by ColabFold^50^. The structure images were analyzed with PyMOL (version 3.1.6.1).

### Preparation of bacteria for coculture

Bacteria were inoculated approximately 48 h before coculture and induced 16-24 h before use. On the day of the experiment, cultures were diluted to an OD600 of 0.2-1.0 for spectrophotometric measurement. Bacterial concentrations were calculated using the conversion described above, and the required CFU were determined from the multiplicity of infection (MOI, cell: bacteria ratio) indicated for each assay. Bacteria were washed and resuspended in the complete culture medium used for the corresponding mammalian cells.

### Cell binding assays

For assays involving adherent tumor cells, 10,000-20,000 cells per well were seeded in 96-well plates. Cells were cocultured with the indicated bacteria for 2 h, washed and detached by incubation with PBS containing 2 mM EDTA for 30 min at 37 degrees C. Cell associated bacteria were detected using an *anti-E. coli* polyclonal antibody (1000:1) listed in **Supplementary Table S2** and quantified by flow cytometry. Suspension tumor cells were processed in parallel without the detachment step.

### Immune cell binding and activation assays

Primary human T or NK cells were incubated with the indicated LIME or scaffold control bacteria at a bacteria-to-immune-cell ratio of 100:1 for 48 h. Cells were then collected and analyzed by flow cytometry for cell-associated *E. coli* and for the activation and proliferation markers specified in **Supplementary Table S2**. Where tumor cells were included, tumor cells, effector cells and bacteria were combined at a ratio of 1:1:100 and incubated for 48 h.

### Immune-cell-tumor-cell bridging assays

For flow cytometric measurement of LIME-mediated cellular bridging, H69 cells were labeled with CellTrace Violet (Invitrogen, C34557). Primary human T cells, labeled H69 cells and DLL3 T-LIME or the indicated controls were cocultured at a ratio of 5:1:50 for 30 min. Samples were fixed with 4% paraformaldehyde (PFA), stained with anti-human CD3 antibody (1000:1) and analyzed by flow cytometry. T-cell-tumor-cell conjugates were defined as CellTrace Violet-positive events carrying a CD3 signal.

For fluorescence imaging, Patu8988S or U87 tumor cells were labeled with Alexa Fluor 647 NHS ester (Invitrogen, A20006) before seeding at 5,000 cells per well in ibiTreat eight-well chamber slides (ibidi, 80806). Primary human T cells were labeled with Alexa Fluor 568 NHS ester (Invitrogen, A20003). T cells, tumor cells and bacteria were cocultured at a ratio of 5:1:50 for 30 min, fixed with 4% PFA and stained with anti-*E. coli* antibody (1000:1). Slides were mounted with ProLong Diamond Antifade Mountant (Invitrogen, P36970) and imaged using a Leica THUNDER Imager equipped with an HC PL APO 20x/0.80 objective and the 475 nm, 555 nm and 635 nm channels.

### In vitro tumor cell cytotoxicity assays

Tumor cells, primary human T or NK cells, and the indicated LIME or control bacteria were cocultured for 48 h at a ratio of 1:1:100 in the complete medium used for the tumor cell line, supplemented with 20 U ml^-1^ recombinant human IL-2. At the end of the coculture, cells were divided for analysis of tumor cell death and effector cell phenotype. Tumor cells were distinguished from immune cells by CD45 staining (1000:1) and evaluated using Annexin V and 7-aminoactinomycin D (7-AAD; BioLegend) according to the manufacturer’s instructions. Parallel samples were stained with a fixable viability dye and the immune-phenotyping antibodies (1000:1) listed in **Supplementary Table S2**.

### PDAC organoid coculture

PDAC organoid derived cells, primary human T cells and the indicated bacteria were combined at a ratio of 1:1:100. Approximately 20,000 organoid derived cells were embedded with T cells and bacteria in Matrigel and cultured in complete OPAC medium supplemented with 20 U ml^-1^ recombinant human IL-2 for 48 h. Cultures were recovered from Matrigel and dissociated with TrypLE. Tumor cell death and T cell phenotype were assessed by flow cytometry as described for cell line cocultures.

### Bulk RNA sequencing and analysis

In total, 2–3 million T cells from three different donors (D1, D2 and D3) were co-cultured with engineered bacteria (bacteria to immune cell ratio 100:1), PBS, Tarlatamab (10nM) or scaffold bacteria for 12h at one humidified incubator with 5% CO_2_, 37 °C, after which cells were harvested for RNA extraction and RNA-seq. Libraries were sequenced on a NovaSeq X Plus platform (paired-end, non-directional). Raw reads were filtered to remove adapter contamination, reads with >10% uncertain nucleotides, and low-quality reads. Clean reads were aligned to the human reference genome (hg38) using HISAT2 v2.2.1^51^ with default parameters, and gene level read counts were generated using featureCounts v2.0.6^52^.

Differential expression analysis was performed with DESeq2^53^ in R (v4.6.0, Bioconductor v3.23). Because samples were derived from three independent donors, donor was included as a blocking factor in the design formula (∼ donor + condition), such that donor-to-donor variation was accounted for in the statistical model. Genes with at least 10 counts in a minimum of three samples were retained, yielding 3,490 genes for analysis. All six pairwise comparisons among the four conditions were tested (T-LIME versus PBS, Tarlatamab versus PBS, scaffold versus PBS, T-LIME versus scaffold, Tarlatamab versus scaffold and T-LIME versus Tarlatamab). Genes with Benjamini–Hochberg adjusted P value < 0.05 and |log2 fold change (FC)| > 1 were considered differentially expressed. For volcano plot visualization, log2FC estimates were shrunk using the adaptive shrinkage method (ashr)^54^ implemented in DESeq2’s lfcShrink function; significance thresholds were applied to unshrunken adjusted P values.

For visualization, counts were transformed using the variance-stabilizing transformation (VST), and the estimated donor effect was removed using removeBatchEffect from limma^55^. This correction was applied to principal component analysis, hierarchical clustering, heatmaps and gene set variation analysis only; all statistical testing was performed on uncorrected data using the donor-aware design formula described above.

Pathway analysis was performed using gene set enrichment analysis (GSEA)^56^ on pre-ranked gene lists via the fgsea package^57^, with 1,000 permutations and gene set size limits of 15–500 genes. Genes were ranked by signed –log10(adjusted P value) for significantly changed genes and by log2FC otherwise. Gene sets were obtained from MSigDB (Hallmark, KEGG, GO Biological Process and Reactome collections) using msigdbr^58^. Pathways were considered significantly enriched at nominal P < 0.05, false discovery rate (FDR) q value < 0.25 and |normalized enrichment score (NES)| > 1, consistent with standard GSEA methodology.

Gene set variation analysis (GSVA)^59^ was performed on the donor-corrected VST expression matrix using Hallmark and KEGG gene sets (minimum 10, maximum 500 genes per set). Differences in pathway activity scores across the four conditions were assessed by Kruskal–Wallis test, with pairwise comparisons by two-sided Wilcoxon rank-sum test.

### Animal experiment

Female C57BL/6J (The Jackson Laboratory, 000664), BALB/c (The Jackson Laboratory, 000651) and NOD.Cg-Prkdcscid Il2rgtm1Wjl/SzJ (NSG; The Jackson Laboratory, 005557) mice were maintained in a specific-pathogen-free facility at DFCI. Mice were housed under a 12-h light-dark cycle at 22 degrees C and 40-70% relative humidity, with standard chow and water provided ad libitum. Female C57BL/6J and BALB/c mice were 6-8 weeks old at tumor implantation; Both male and female NSG mice were 4-6 weeks old for the subcutaneous H69 model and the metastatic H69 model.

Tumors were measured two times per week using digital calipers, and volume was calculated as 0.5 x length x width^2^. Mice were randomized among treatment groups after tumors reached the prespecified size range. Investigators were not blinded to treatment allocation or outcome assessment. Mice were euthanized when tumor or health-related humane endpoints specified in the approved protocol were reached, including severe ulceration, a hunched posture or loss of more than 15% of initial body weight. Humane euthanasia was recorded as an event in survival analyses.

For A20 B cell lymphoma model, female BALB/c mice were injected subcutaneously in one flank with 1 × 10^6^ A20 cells in 100 ul sterile PBS. On day 10, when tumors measured 80-150 mm^3^, mice were randomized to receive PBS, scaffold bacteria, CD19 T-LIME, CD19 N-LIME, anti-PD-1, CD19 T-LIME plus anti-PD-1 or CD19 N-LIME plus anti-PD-1. Bacteria were administered intravenously at 1 × 10^8^ CFU in 100 ul PBS on days 10 and 14.

Anti-PD-1 (InVivoMAb anti-mouse PD-1, clone RMP1-14; Bio X Cell, BE0146) was administered intraperitoneally at 8 mg kg^-1^ on the same days. Survival was the primary endpoint.

On day 60, mice that remained tumor-free after the initial challenge were injected subcutaneously with 1 × 10^6^ A20 cells in the contralateral flank. Age-matched, tumor-naive BALB/c mice received the same challenge as controls. Mice were monitored for tumor development and survival without further treatment.

For KPC pancreatic cancer model, female C57BL/6J mice were injected subcutaneously in one flank with 1 × 10^6^ KPC C2 cells in 100 ul sterile PBS. For the tumor profiling study, mice with tumors measuring 150-200 mm^3^ on day 10 were randomized to PBS, scaffold bacteria, MSLN T-LIME or MSLN N-LIME. Bacteria were administered intravenously at 2 × 10^8^ CFU in 100 ul PBS on days 10, 13, 16 and 20. Mice were euthanized on day 23, and tumors, blood, tumor draining lymph nodes (TDLNs), spleen, liver, lungs, kidneys and bone marrow were collected for biodistribution, histology, flow cytometry and soluble factor analysis.

For the anti-PD-1 combination study, tumor-bearing mice were treated with PBS, scaffold bacteria, MSLN T-LIME or MSLN N-LIME, with or without anti-PD-1 (clone RMP1-14; 8 mg kg^-1^ intraperitoneally), on the schedule shown in **Extended Data Fig. 7**. Tumor growth and survival were monitored.

For studies of RAS inhibition, mice bearing KPC C2 tumors were randomized on day 10 to PBS, scaffold bacteria, MSLN T-LIME, MSLN N-LIME, RMC-7977, MSLN T-LIME plus RMC-7977 or MSLN N-LIME plus RMC-7977. Bacteria were administered intravenously at 2 × 10^8^ CFU per dose, and RMC-7977 (Selleckchem, E1858) was administered by oral gavage at 25 mg kg^-1^ on days 10, 14, 17 and 21. The formulation and vehicle used for RMC-7977 are 10/20/10/60 (%v/v/v/v) DMSO/PEG400/Solutol HS15/water (MedChemExpress) as described previously^60^. Mice in the efficacy cohort were monitored for tumor growth and survival. A parallel mechanistic cohort was euthanized on day 24 for collection of tumors, blood and peripheral organs.

For the three agent study, treatment was initiated when KPC C2 tumors reached 150-200 mm^3^. LIME consisted of an equal CFU mixture of MSLN T-LIME and MSLN N-LIME and was administered intravenously; RMC-7977 was administered orally at 25 mg kg^-1^; and anti-PD-L1 (InVivoMAb anti-mouse PD-L1, clone 10F.9G2™; Bio X Cell, BE0101, 8 mg kg^-1^) was administered intraperitoneally on days 12, 15, 18 and 21. Mice received each monotherapy, the indicated pairwise combinations or the three agent combination. Tumor growth and survival were monitored.

For Subcutaneous H69 small-cell lung cancer model, NSG mice (male and female) were injected subcutaneously in one flank with 5 × 10^6^ H69 cells in 100 ul sterile PBS. On day 11, mice received 5 × 10^6^ activated primary human CD3^+^ T cells intravenously in 100 ul PBS. On day 12, when tumors measured approximately 100-120 mm^3^, mice were randomized to PBS, scaffold bacteria, DLL3 T-LIME or tarlatamab (MedChemExpress, 2307488-83-9). Scaffold and DLL3 T-LIME were administered intravenously at 2 × 10^8^ CFU in 100 ul PBS on days 12, 17 and 21. Tarlatamab was administered intraperitoneally at 3 mg kg^-1^ on the same schedule. An H69 only group did not receive human T cells or study treatment. Mice in the efficacy cohort were monitored for tumor growth and survival. A parallel cohort was treated using the same schedule and euthanized on day 24 for immune profiling, bacterial biodistribution and soluble factor analysis.

For metastatic H69 model, NSG mice received 5 × 10^6^ H69 cells intravenously through the tail vein in 100 ul sterile PBS. Human CD3^+^ T cells were administered intravenously on day 11, and PBS, scaffold bacteria, DLL3 T-LIME or tarlatamab was administered on days 12, 17 and 21 using the doses and routes described for the subcutaneous H69 model. Mice were euthanized on day 35. Livers were weighed, photographed, fixed and processed for hematoxylin and eosin (H&E) staining. Metastatic burden was quantified as the percentage of the whole-liver section occupied by tumor by Qupath v0.6.0.

### Bacterial biodistribution

Tumors and the indicated organs were collected aseptically, weighed and homogenized in digestion buffer (RPMI-10 with 5% FBS, 10 mM HEPES, 100 μg ml^−1^ P/S and 1 mg ml^−1^ Collagenase IV). Serial dilutions of tissue homogenates were plated on selective LB agar containing 50 ug ml^-1^ kanamycin and incubated at 37 °C. Colonies were counted and normalized to tissue mass as CFU g^-1^. Blood and bone marrow (flushed by 500μl RPMI-10 media) were processed similarly and reported per volume or tissue mass as indicated.

### Preparation of tissues for flow cytometry

Tumors and solid organs were minced and digested for 1h at 37 °C with agitation in RPMI containing 5% FBS, 10 mM HEPES, 100 ug ml^-1^ penicillin-streptomycin and 1 mg ml^-1^ collagenase IV. Digested tissues were passed through 70 um strainers. Red blood cells were lysed using ACK buffer. Spleens and TDLNs were mechanically dissociated and passed through 70 um strainers. Bone marrow was flushed from the hindlimbs, filtered and treated with ACK buffer. Peripheral blood was collected into anticoagulant tubes (heparin coated) and subjected to red blood cell lysis before staining.

### Flow cytometry

Antibodies, fluorophores, clones and manufacturers are listed in **Supplementary Table 2**. All the dilutions follow the instructions provided on vendor webpage unless mentioned otherwise. Single cell suspensions were washed with PBS and incubated with Fc-blocking antibody (clone 2.4G2; BD Biosciences) for 10 min at 4 °C when murine cells were analyzed. Cells were stained with a fixable viability dye in PBS for 15-20 min at room temperature in the dark, washed with FACS buffer (PBS containing 5 mM EDTA, 2% FBS and 0.1% sodium azide) and incubated with surface antibody cocktails for 30 min at room temperature in the dark.

For intracellular or intranuclear staining, cells were fixed and permeabilized using the True-Nuclear Transcription Factor Buffer Set (BioLegend, 424401) according to the manufacturer’s instructions. Intracellular antibodies were diluted in BD Perm/Wash Buffer (BD Biosciences, 554723) and incubated with cells at 4 °C overnight. Samples were washed and resuspended in PBS before acquisition. Annexin V/7-AAD samples were acquired without fixation. Data were acquired using a Sony ID7000 spectral flow cytometer and analyzed in FlowJo and OMIQ.

### Immunofluorescence microscopy

Freshly collected tumors were embedded in optimal cutting temperature compound, frozen at −80 °C and cryosectioned by iHisto. Frozen sections were washed three times with PBS, fixed in 1% PFA for 2 min in the dark and washed again. Sections were permeabilized in PBS containing 1% Triton X-100 for 30 min on ice, washed with PBS containing 0.2% Triton X-100, and blocked for 2h on ice in PBS containing 0.2% Triton X-100, 2 mg ml^-1^ BSA and 10% mouse serum. Primary antibodies were diluted in blocking buffer as recommended by the vendor and incubated overnight at 4 °C. After washing, sections were incubated with the appropriate secondary antibodies for 1 h on ice. Sections were washed, post-fixed in 1% PFA for 10 min and mounted using EverBrite Mounting Medium with DAPI (Biotium, 23002).

Images were acquired using a Zeiss LSM 980 confocal microscope or a Leica THUNDER Imager, as indicated in the figure legends. ZEN Blue v.2.6 or the Leica acquisition software was used for acquisition and processing. Pseudocolors were applied uniformly within each experiment. Image processing settings used for quantitative comparisons were held constant across treatment groups.

### Histology

Liver, spleen and kidney samples were fixed by 4% PFA in 4 °C for 24h, embedded in paraffin, sectioned and stained with H&E by iHisto. Whole-slide images or representative fields were acquired at the magnifications indicated in the figure legends. For the metastatic H69 model, tumor area was quantified across whole liver sections and expressed as a percentage of total liver section area.

### Luminex analysis

Blood was collected before euthanasia into lithium-heparin tubes (Greiner Bio-One, 450479). Plasma was isolated by centrifugation at 4,500g for 30 min at 4 °C and stored frozen until analysis. Tumor tissue was homogenized in tissue protein extraction reagent (T-PERTM, Thermo Fisher Scientific, 78510) in the presence of 1% proteinase and phosphatase inhibitors (Thermo Fisher Scientific, 78442) at 4 °C for 30 min with slow rotation then centrifuged to remove debris. Plasma and tumor lysates were analyzed by Eve Technologies using human or mouse multiplex Luminex panels, as appropriate. Analytes are listed in the corresponding figure legends. Plasma concentrations are reported as pg ml^-1^; tumor concentrations are reported as pg g^-1^ tissue.

### Detection of serum antibodies against KPC C2 lysate

5 × 10^6^ KPC C2 cells were lysed in 1ml RIPA Lysis and Extraction Buffer (Thermo Fisher Scientific, 89900) in the presence of 1% proteinase and phosphatase inhibitors at 4 °C for 30 min with slow rotation then centrifuged to remove debris. 96-well plates were coated with lysate (50ul/well) overnight at 4 degrees C. Plates were washed and blocked with PBS with 1% BSA. Serum collected from treated mice diluted 100 times was added to each well (50ul/well) and followed by horseradish peroxidase-conjugated anti-mouse IgG (Invitrogen, 31430, 1000:1). Signal was developed using Substrate Reagent Pack (R&D, DY999B) and absorbance was measured at 450 nm with background correction at 540 nm. Results are reported as OD450-OD540.

### Statistical analysis and reproducibility

Statistical analyses were performed using GraphPad Prism v.11.1.1. Data are presented as mean +/- s.d., with individual biological replicates shown where possible. Exact sample sizes and definitions of replicates are provided in the figure legends. No statistical method was used to predetermine sample size. Mice were randomized to treatment groups after tumor establishment; investigators were not blinded to treatment allocation or outcome assessment.

Two-group comparisons were performed using two-sided Student’s t-tests when indicated. Comparisons among three or more groups were performed using one-way analysis of variance (ANOVA) followed by Tukey’s multiple-comparison test. Tumor growth curves and experiments containing two independent factors were analyzed by two-way ANOVA with Tukey’s multiple comparison test. Survival distributions were estimated by the Kaplan-Meier method and compared using the two-sided log-rank (Mantel-Cox) test. For RNA sequencing, differential-expression P values were calculated using DESeq2 and corrected for multiple testing by the Benjamini-Hochberg method. GSEA results were evaluated using nominal P values and FDR q values as described above. All tests were two-sided unless otherwise specified. Statistical significance thresholds and exact P values are reported in the figures and legends.

## Acknowledgements

This work was supported by Clinical Investigator Award (R.R.), and S.Y. and R.R. received funding from the Parker Institute for Cancer Immunotherapy. This work has further been supported by German Cancer Aid (Deutsche Krebshilfe) in frame of the CAR FACTORY consortium (70115200 to EU) as part of the preclinical cancer drug development network (preCDD) (E.U.). R.R. is a recipient of the Career Development Award from Blood Cancer Unitied (formerly the Leukemia and Lymphoma Society (LLS)). R. R and J. L. acknowledge funding from the National Cancer Institute R01CA299949. S.S. was supported by the Deutsche Forschungsgemeinschaft (DFG, German Research Foundation) through a Walter Benjamin Fellowship. We would like to thank Flow cytometry core, Molecular imaging core and Animal research facility at DFCI for their support. The cartoons were created using BioRender (https://biorender.com/). Figures were generated by RStudio (v 2026.05.0+218), Graph Pad Prism (v 10.1.1) and Adobe Illustrator (v 27.5).

## Author contributions

R.R., S.Y. and A.C.B. conceptualized the study. R.R., S.Y. and A.C.B. wrote manuscripts. S.Y., R.R. and A.C.B. designed the experiments, performed analysis and data interpretation. S.Y., A.C.B., S.S., A.H., D.C.C., H.N., A.C., E.B., A.M., J.T., F.L., X.D., M.N., M.S., V.W.H., T.E.K., C.L.A., Y.W., J.T., H.L., performed the experiments. M.S., M.T., A.K.A., R.S., K.Z., Y.R.C., Z.W., M.C., J.L., E.P., S.D., D.B., A.A.L., S.C., E.U., H.E., K.L., K.H., K.W.W., J.C., J.K., V.S.S., R.S., J.L., C.J.W., J.R., A.J.A. assisted during the optimization, assisted during the experimental procedures and analysis.

## Competing interests

R.R., S.Y., and A.C.B. are named inventors on a patent application that describes the surface display of engineered bacteria.

R.R and J.C are co-founders of the InnDura Therapeutics. J.R. received research funding from Kite/Gilead, Novartis and Oncternal Therapeutics and serves on advisory boards for Akron Biotech, Clade Therapeutics, Garuda Therapeutics, LifeVault Bio, Novartis and Smart Immune. J.L. involved as an investigator in clinical trials conducted in collaboration with Moderna Inc., SNIPR Biome, and Elio, receives research funding from Merck, is a member of adjudication committee for Basilea, a member of advisory board for Aicuris Inc and Pfizer, and consulting for Melinta and Shionogi. A.A.L. received research funding from Abbvie and Stemline Therapeutics, is a consultant for Qiagen and Stelexis Biosciences, serves on a steering committee for Stemline Therapeutics, and has equity as an advisor for Medzown and as a co-founder of Stelexis BioSciences. K.W.W. serves on the scientific advisory boards of DEM BioPharma, Solu Therapeutics, D2M Biotherapeutics, DoriNano, Inc., and Nextechinvest. He is a co-founder of Immunitas Therapeutics and receives sponsored research funding from Fate Therapeutics. He holds equity in TScan Therapeutics. J.K. serves on scientific advisory board and/or consults for Biolojic Design, Cue Biotherapeutics, CSL Behring, Cugene, Equillium, Gentibio, Mallinckrodt/Therakos, Sonoma Bio, Visterra, and Biopharm Communications LLC; and has received grants/research support from BMS, Equillium, Iovance, Miltenyi, and Regeneron. These activities are not related to the research reported in this publication.

The remaining authors declare no competing interests.

**Correspondence and requests for materials** should be addressed to Rizwan Romee, MD.

## Data Availability

All data generated during this study are available within the paper. The bulk RNA-seq data will be deposited to the Gene Expression Omnibus before publication. Source data will be provided with this paper before publication.

## Supplementary materials

**Supplementary table S1.**
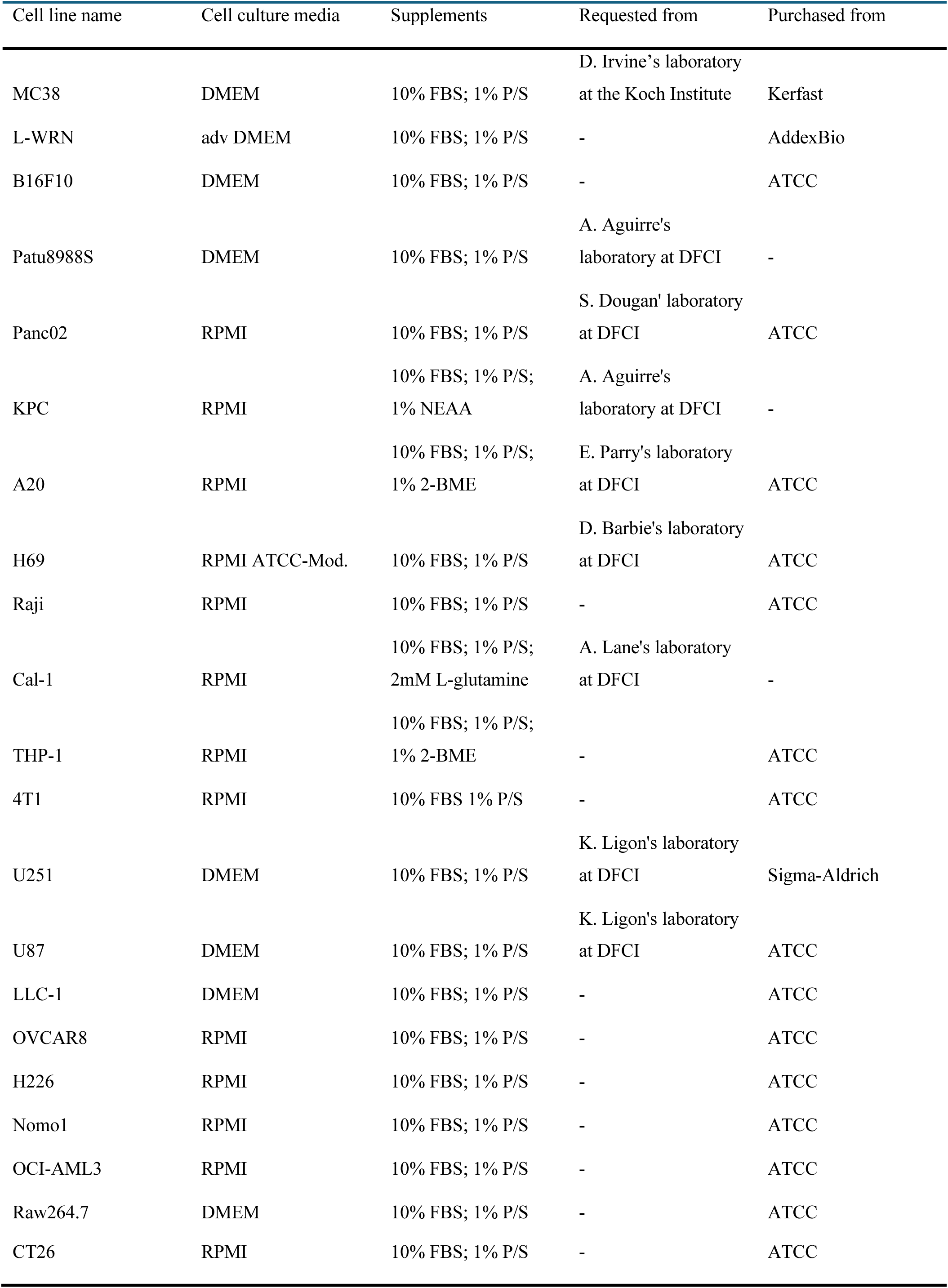
Cell names, media and source of cell lines used in this study.

**Supplementary table S2.**
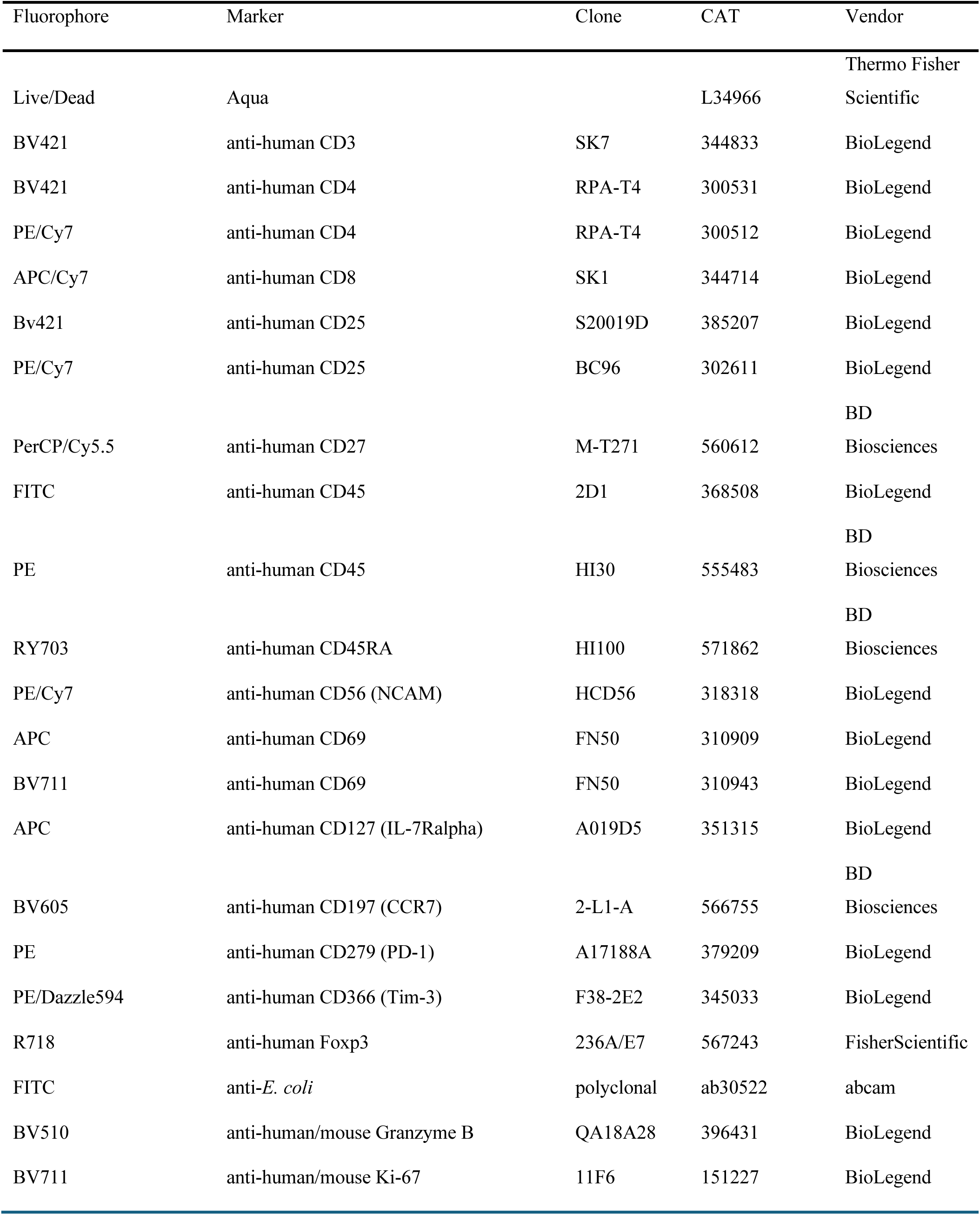
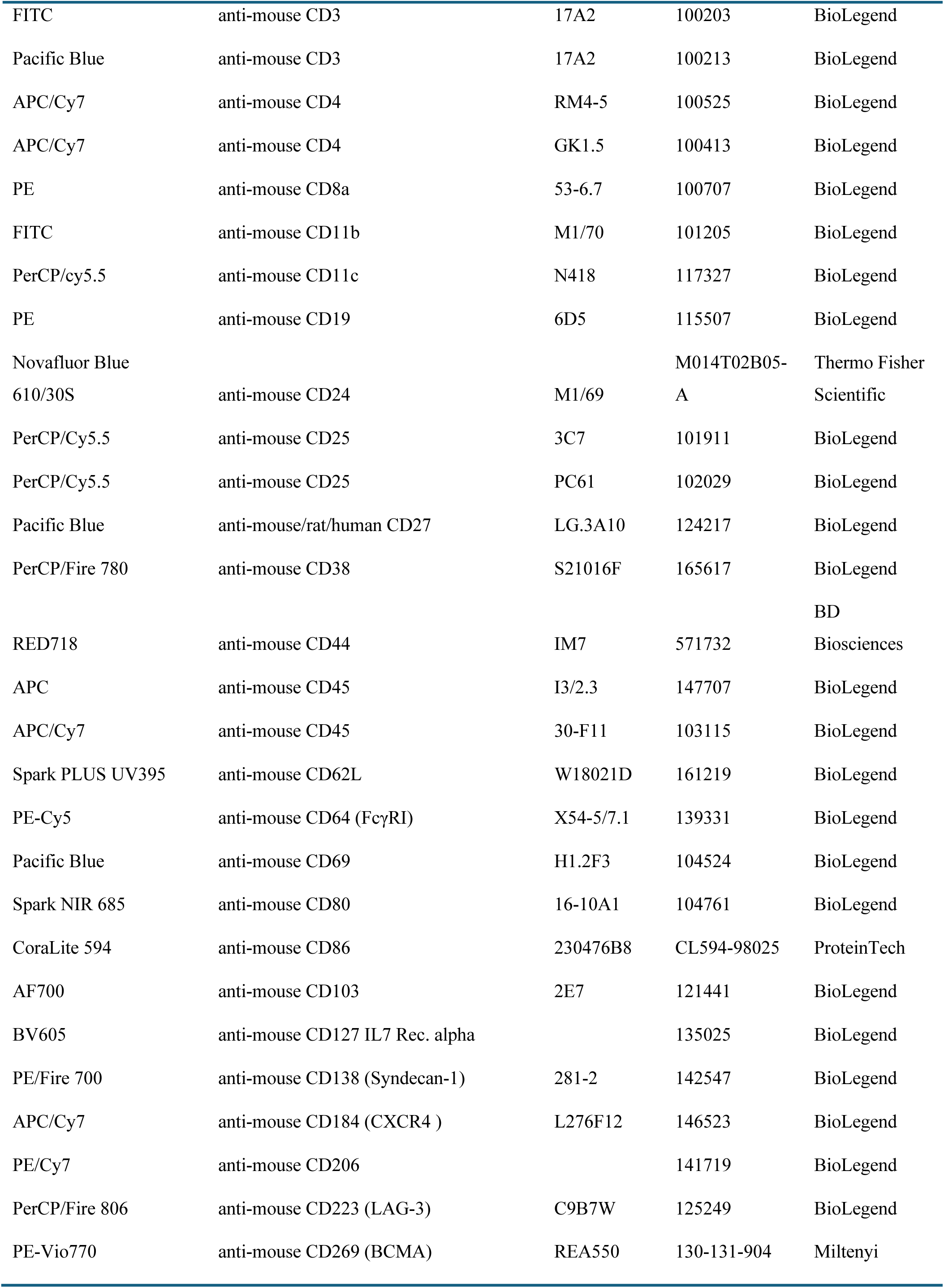
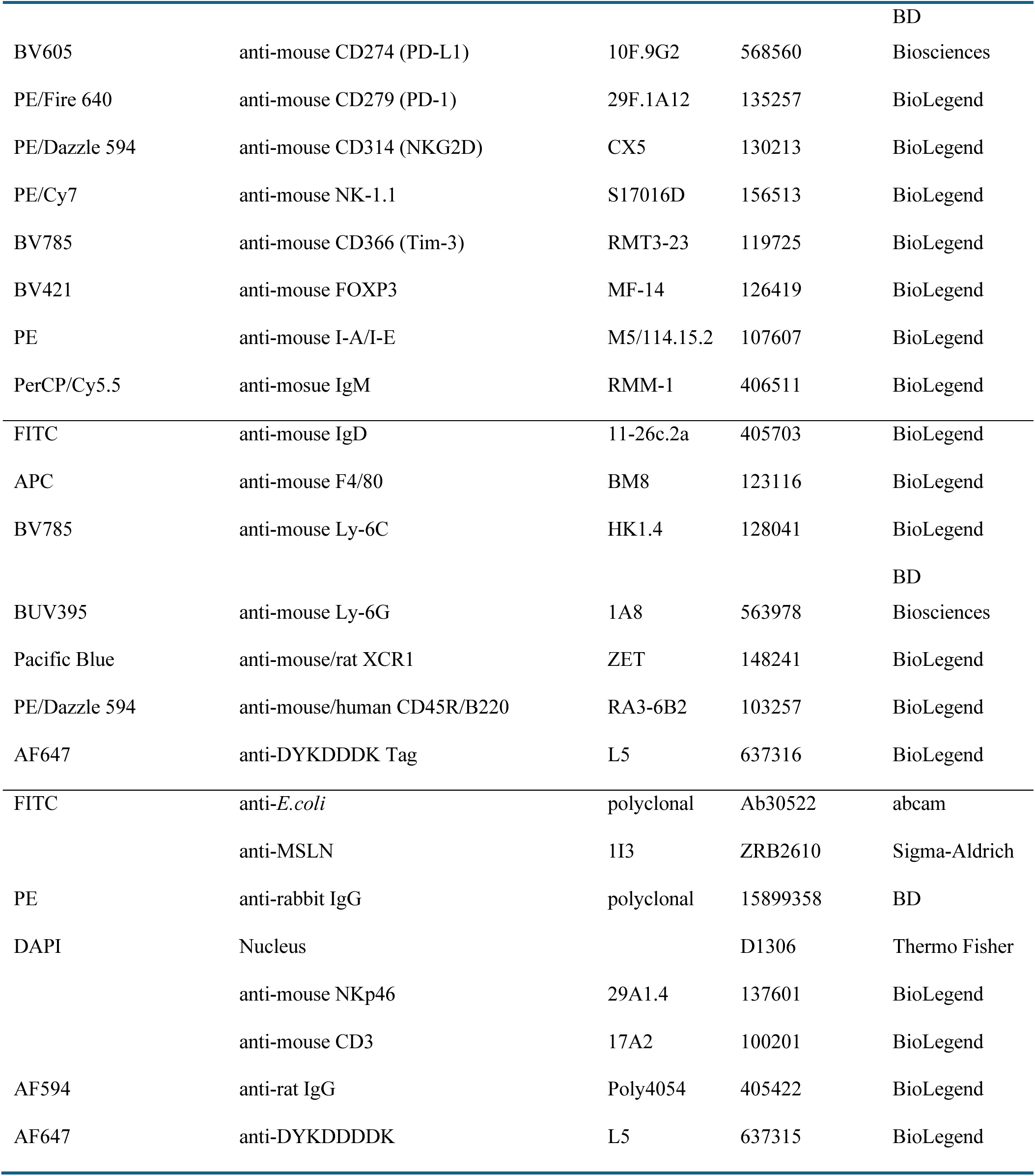
Antibodies and Live/Dead dye used in this study.

## Extended Data Figures and Supplemental Figures

**Extended Data Fig. 1.**
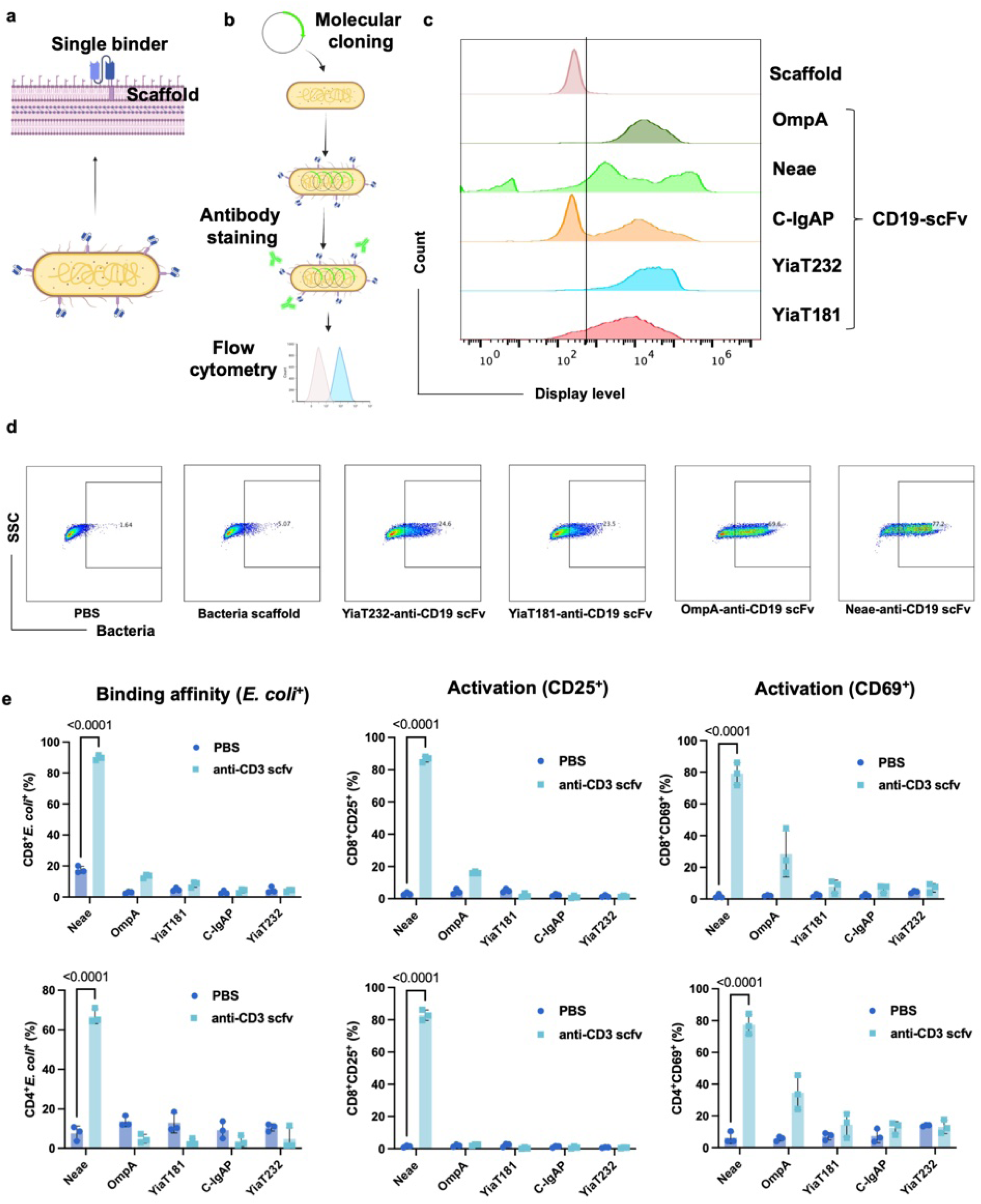
Neae is the optimal scaffold for surface displaying of single-chain variable fragments (scFvs) *E. coli*. **a,** Schematic figure of scFv surface expression in non-pathogenic *E. coli*. **b,** Workflow for construct design and evaluation of scFv surface expression by flow cytometry. **c,** Comparative analysis of surface display levels of anti-CD19 scFv across multiple scaffolds (C-IgAP, OmpA, YiaT232, YiaT181, and Neae). **d,** Representative dot plots showing the binding affinity of *E. coli* displaying anti-CD19 scFv to CD19^+^ Raji cells. The binding affinity was quantified by flow cytometric detection of bacterial signal on the Raji cell surface, using a FITC-conjugated anti-*E. coli* polyclonal antibody. **e,** Binding affinity (*E. coli*^+^) and activation (CD25^+^ and CD69^+^) of human primary CD3^+^ T cells (CD8^+^ T and CD4^+^ T) following coculture with *E. coli* displaying anti-human CD3 scFv or scaffold-only controls, assessed by flow cytometry. Statistical significance was determined using one-way ANOVA test (**e**). Data represent mean ± s.d.

**Supplemental Fig. 1.**
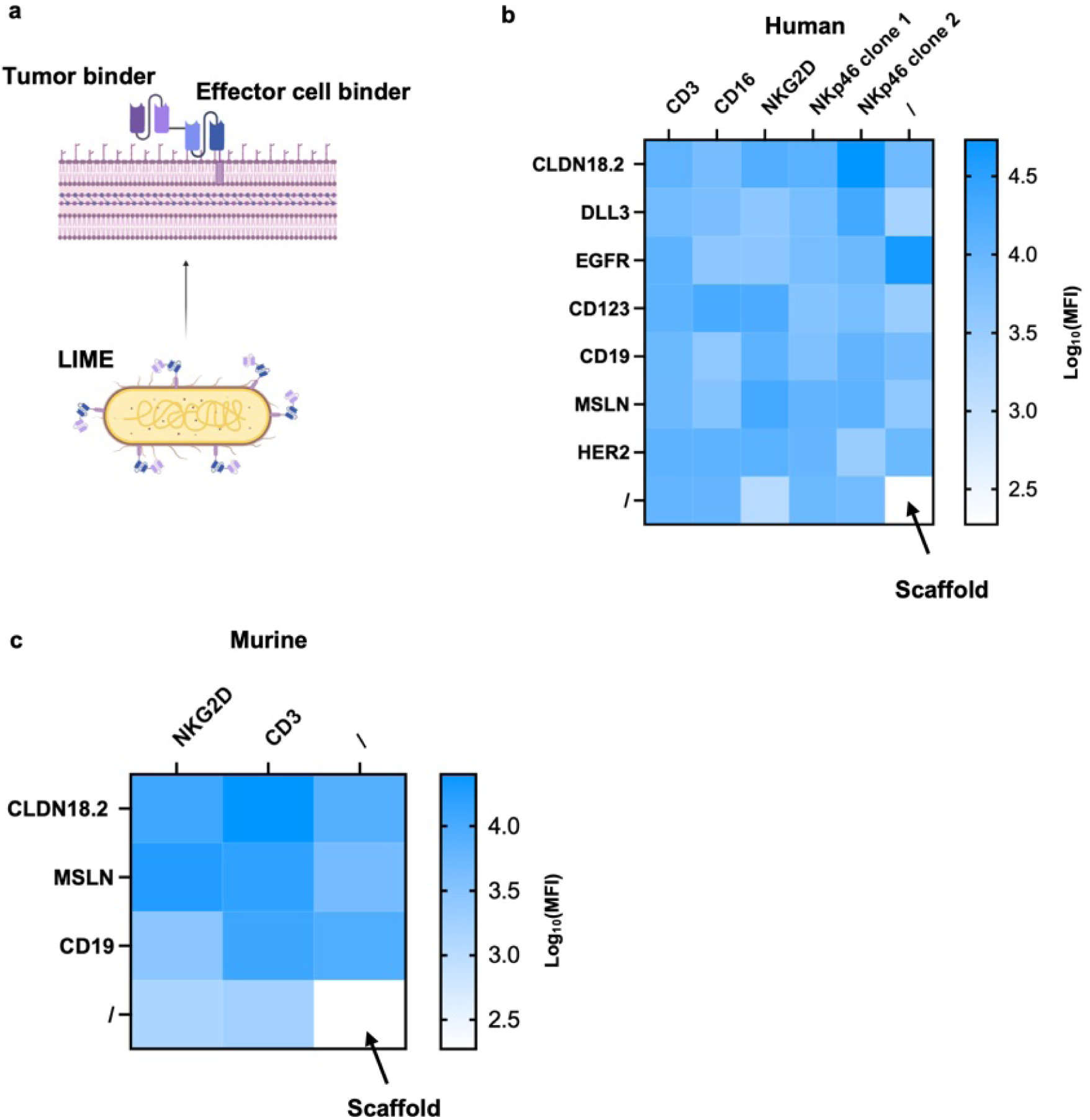
Neae can display diverse bi-specific scFvs on the *E. coli* cell surface. **a**, Schematic figure of the LIME design: *E. coli* engineered to display bi-specific scFvs that simultaneously target tumor antigens and effector cells (T or NK cells). **b, c**, Heatmap summarizing surface display levels of bi-specific scFvs or single scFv for human (**b**) or mouse (**c**) targets, as assessed by flow cytometry. “/” or “\” represent single scFv. “/” and “\” represent scaffold only (negative control). The display level is illustrated by log10 (Median Fluorescence Intensity; MFI).

**Supplemental Fig. 2.**
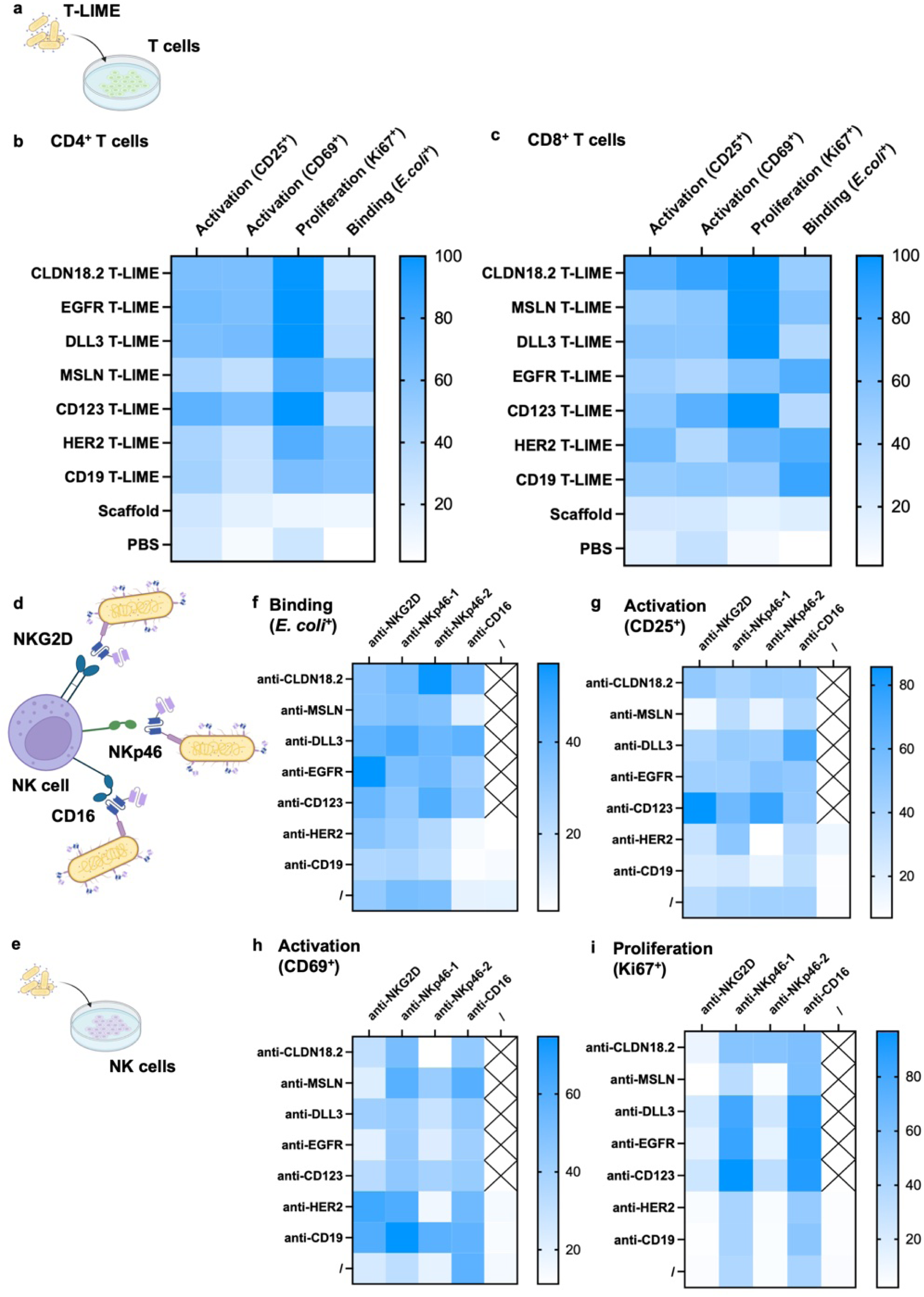
LIME can engage effector cells triggering activation and proliferation. **a**, Schematic figure of the co-culture assay used for functional T-LIME evaluation, as described in Methods. **b**, **c**, Heatmaps showing activation (CD25 and CD69), proliferation (Ki67) and binding (*E. coli*^+^) to CD4^+^ (**b**) and CD8^+^ T cells (**c**) following coculture with T-LIME targeting different tumor antigens or control conditions. **d, e**, Schematic figures describing N-LIME constructs targeting different surface markers of NK cells and the corresponding NK cell coculture assay. **f-i**, heatmaps depicting NK cell binding (**f**, *E. coli*⁺), activation (**g**, CD25; **h**, CD69), and proliferation (**i**, Ki67) after coculture with N-LIME or controls.

**Supplemental Fig. 3.**
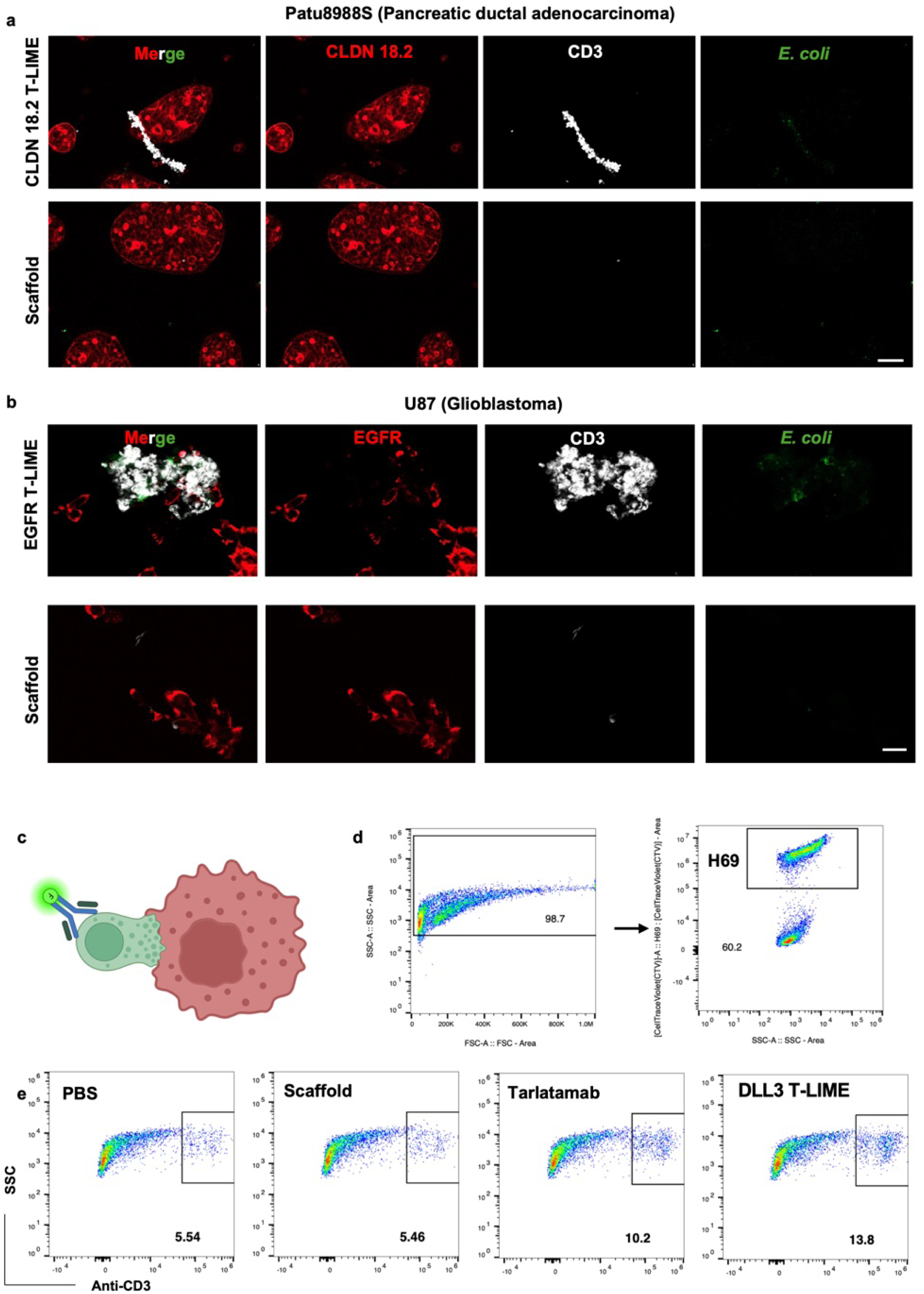
LIME-mediated immunological synapse formation between T cells and cancer cells. **a, b,** Representative confocal images of cocultures compromising a PDAC cell line Patu8988S (CLDN18.2^+^, **a**) or GBM cell line U87 (EGFR^+^, **b**), T cells and engineered *E. coli* (T-LIME or Scaffold control). Scale bar is 50μm (**a**), or 20μm (**b**). **c, d,** Schematic of the measurement for LIME mediated synapse formation by flow cytometer and its representative gating strategy**. e,** Synapse formation between human primary T cells and DLL3^+^ small cell lung cancer cells (H69) mediated by DLL3 targeting T-LIME, or control conditions (tarlatamab, scaffold only, and PBS). Synapse formation was quantified by detection of CD3 signal on CellTraceViolet (CTV)-labelled H69 cells.

**Supplemental Fig. 4.**
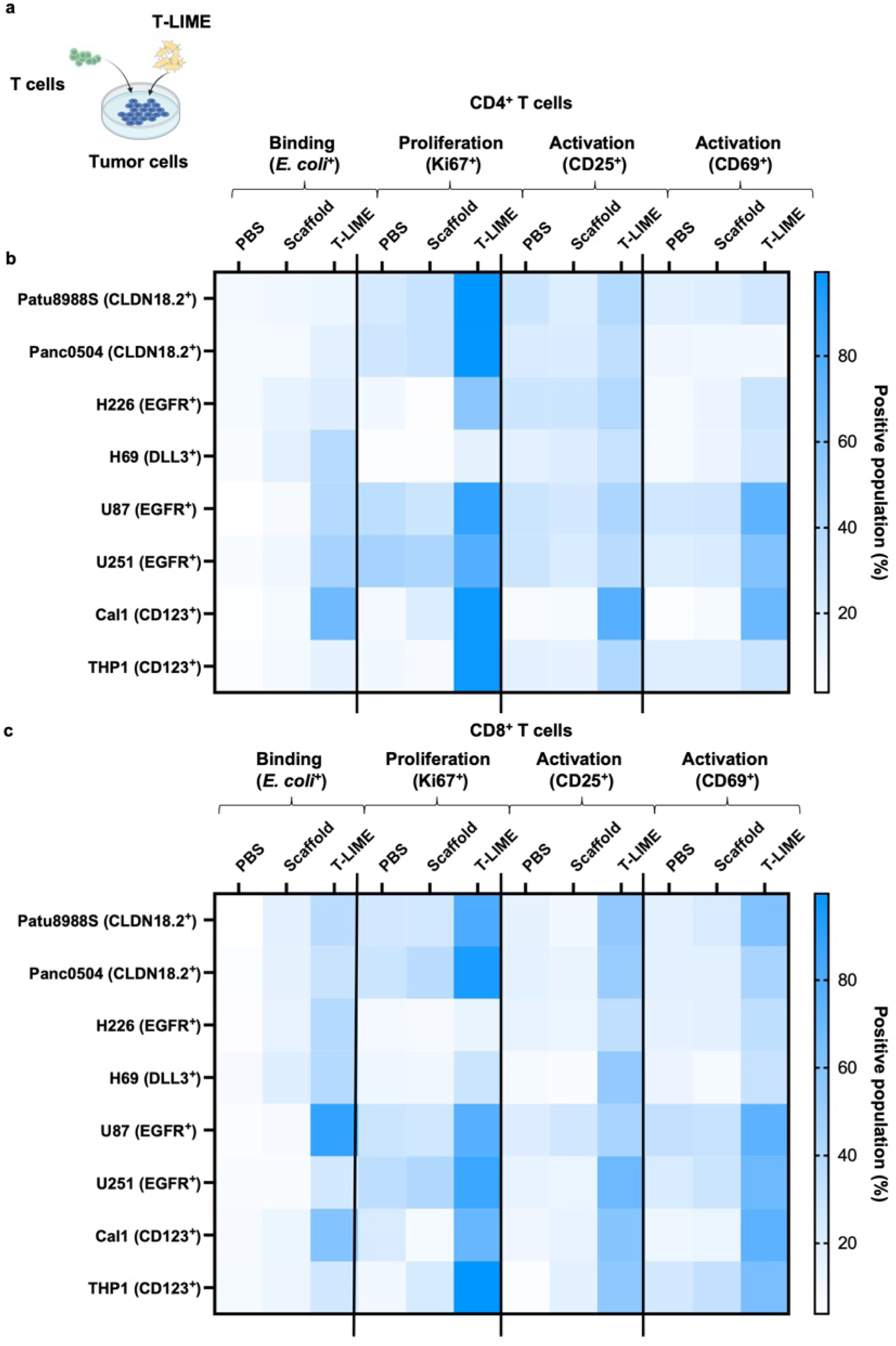
T-LIME bind to T cells and drive their activation and proliferation in the presence of tumor cells. **a,** Schematic figure of the T-lime, T- and tumor cell co-culture. **b, c**, Binding (of engineered *E. coli*⁺), activation (CD25, CD69), and proliferation (Ki67) of CD4⁺ (b) and CD8⁺ (c) T cells following coculture with T-LIMEs in the presence of tumor cells. Heatmap scale represent percentage of positive population.

**Supplemental Fig. 5.**
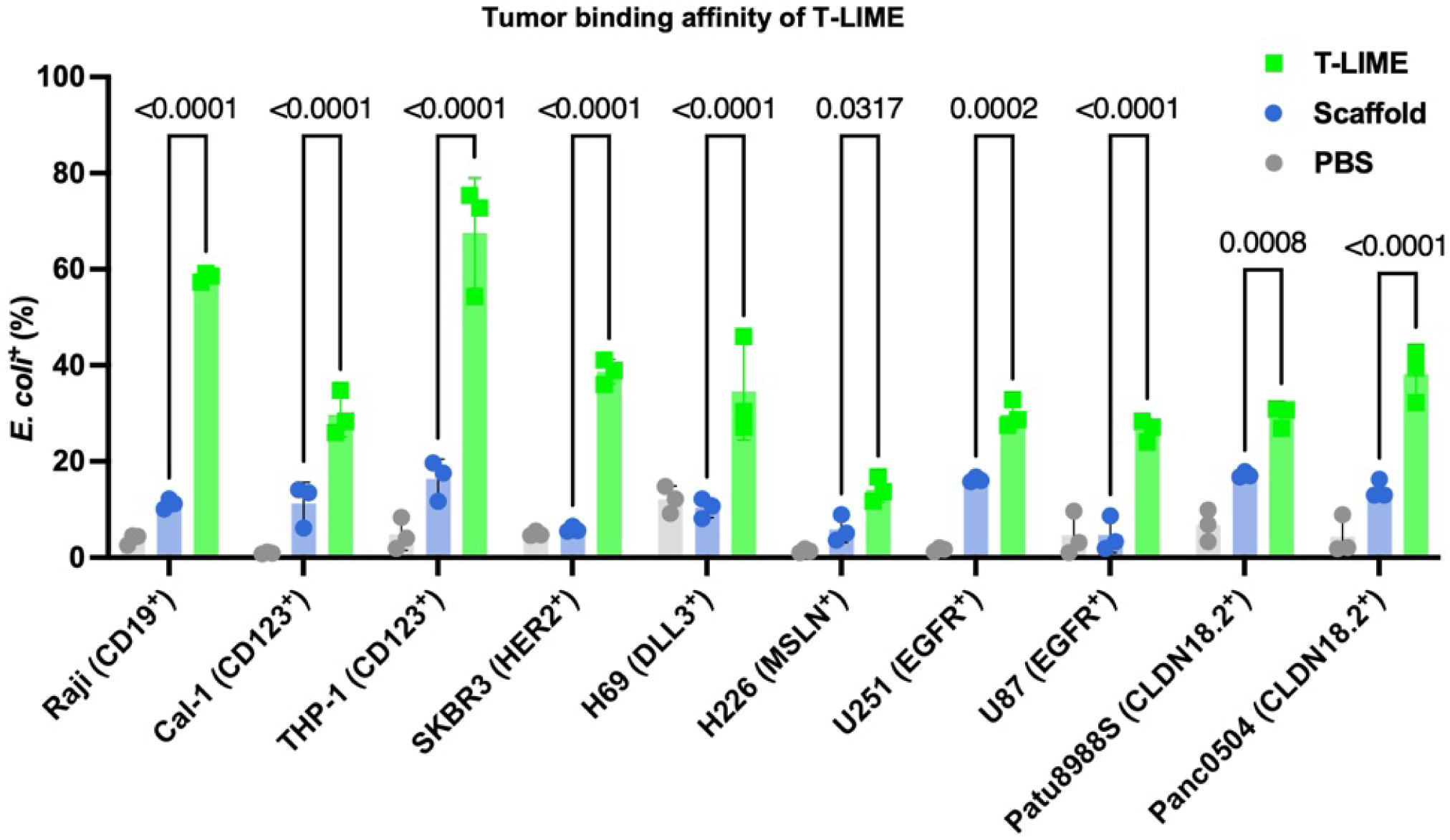
T-LIME have higher binding affinity to the tumor cells compared to controls (PBS and scaffold) as assessed by flow cytometry. Data are presented as mean ± s.d.

**Extended Data Fig. 2.**
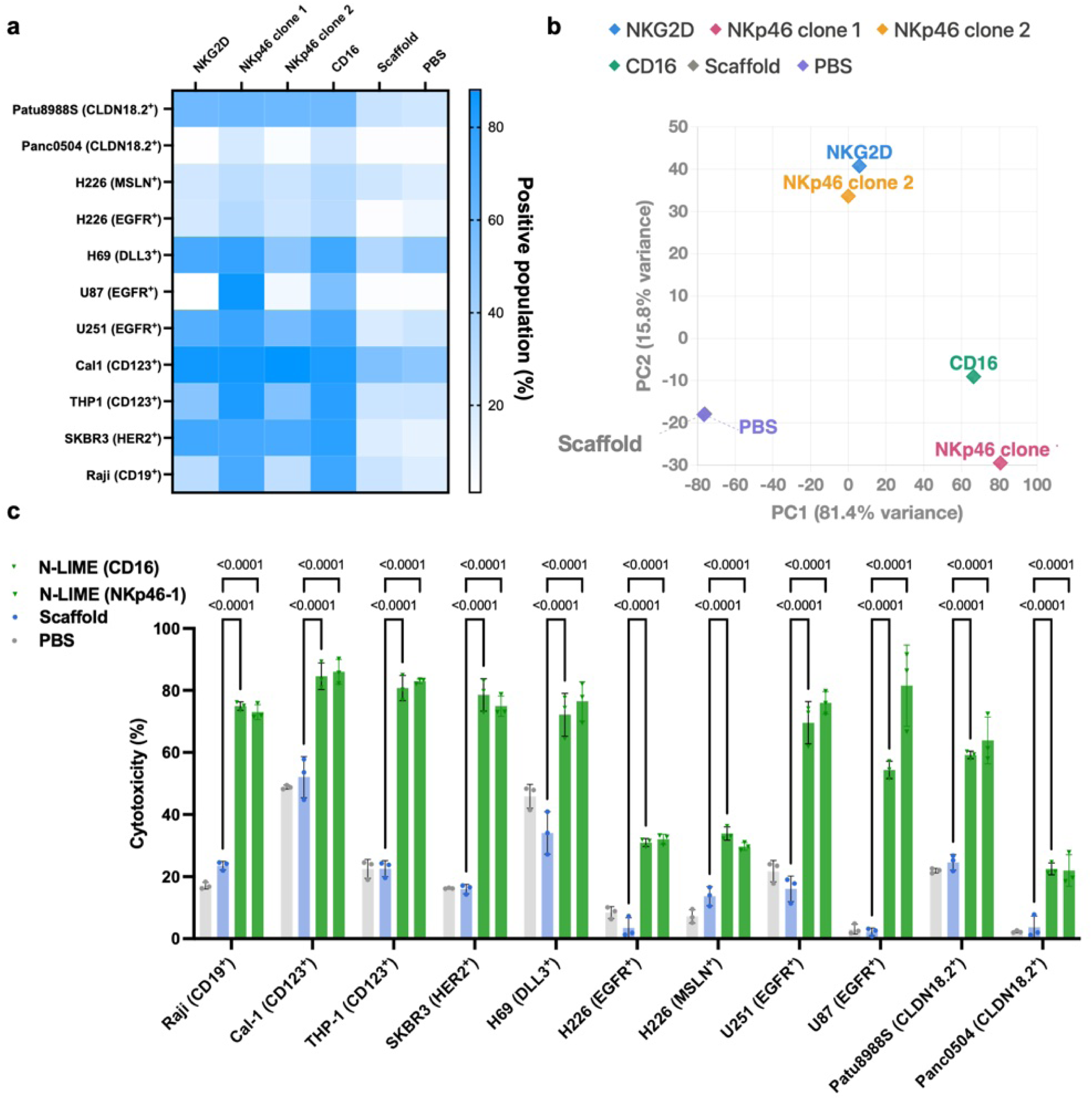
N-LIME enhance NK cell mediated anti-tumor responses. **a–c,** Cytotoxicity of human primary NK cells mediated by various N-LIME constructs (NKG2D, NKp46 or CD16 N-LIME) targeting a panel of cancer cell lines, shown as a heatmap (**a**), Principal Component Analysis (PCA, **b**) and bar graph (**c**). Heatmap scale represent percentage of positive population. Data are presented as mean ± (**c**).

**Extended Data Fig. 3.**
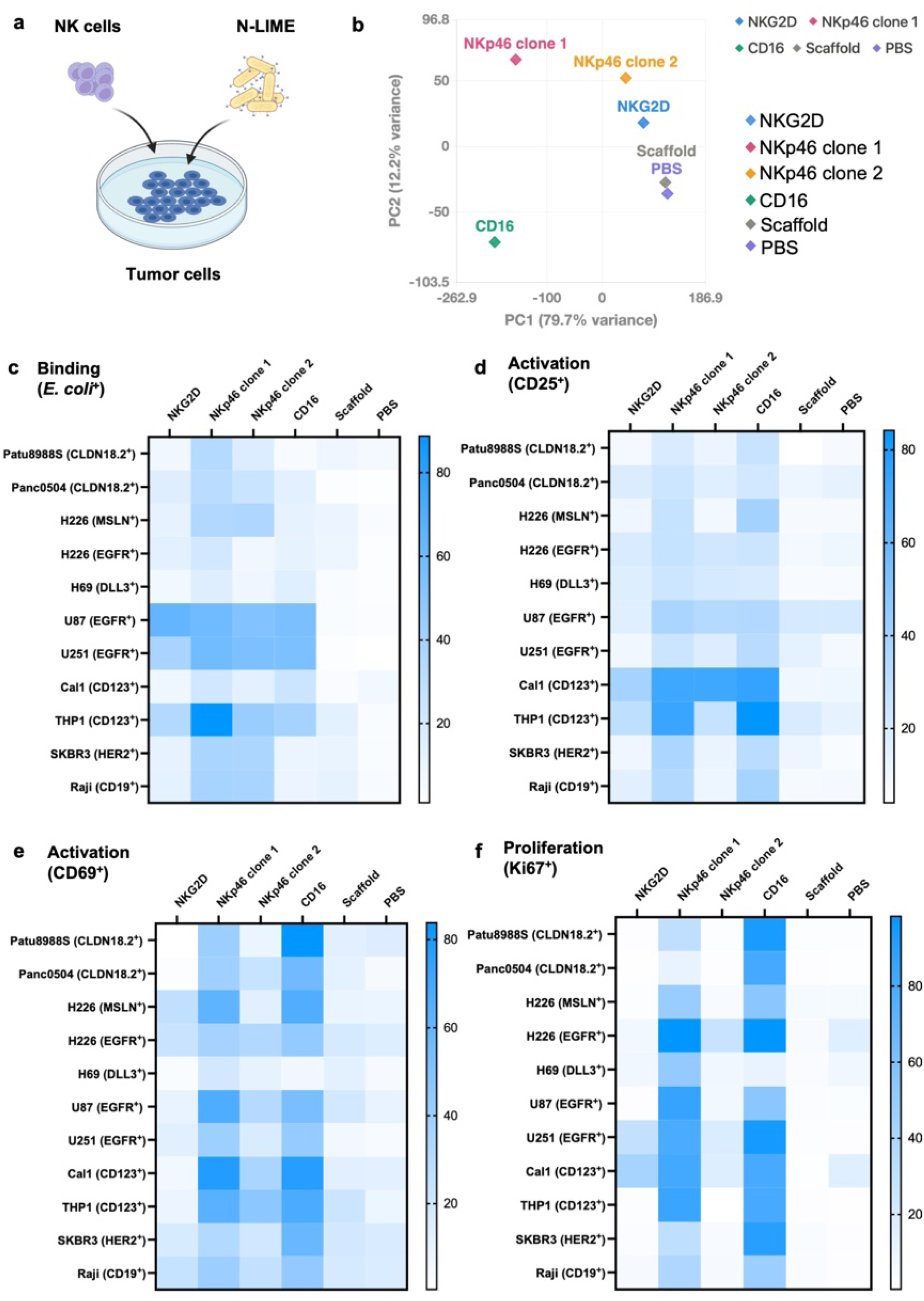
N-LIME engage NK-cells inducing their activation and proliferation in the presence of tumor cells. **a,** Schematic figure describing the co-culture assay involving NK cells, tumor cells and N-LIME. **b-f,** PCA (**b**) and heatmap illustrating NK cell binding affinity (*E. coli*, **c**), activation (CD25 and CD69, **d, e**) and proliferation (Ki67, **f**) following coculture with N-LIMEs or control conditions (PBS and scaffold). Heatmap scale represent percentage of positive population.

**Supplemental Fig. 6.**
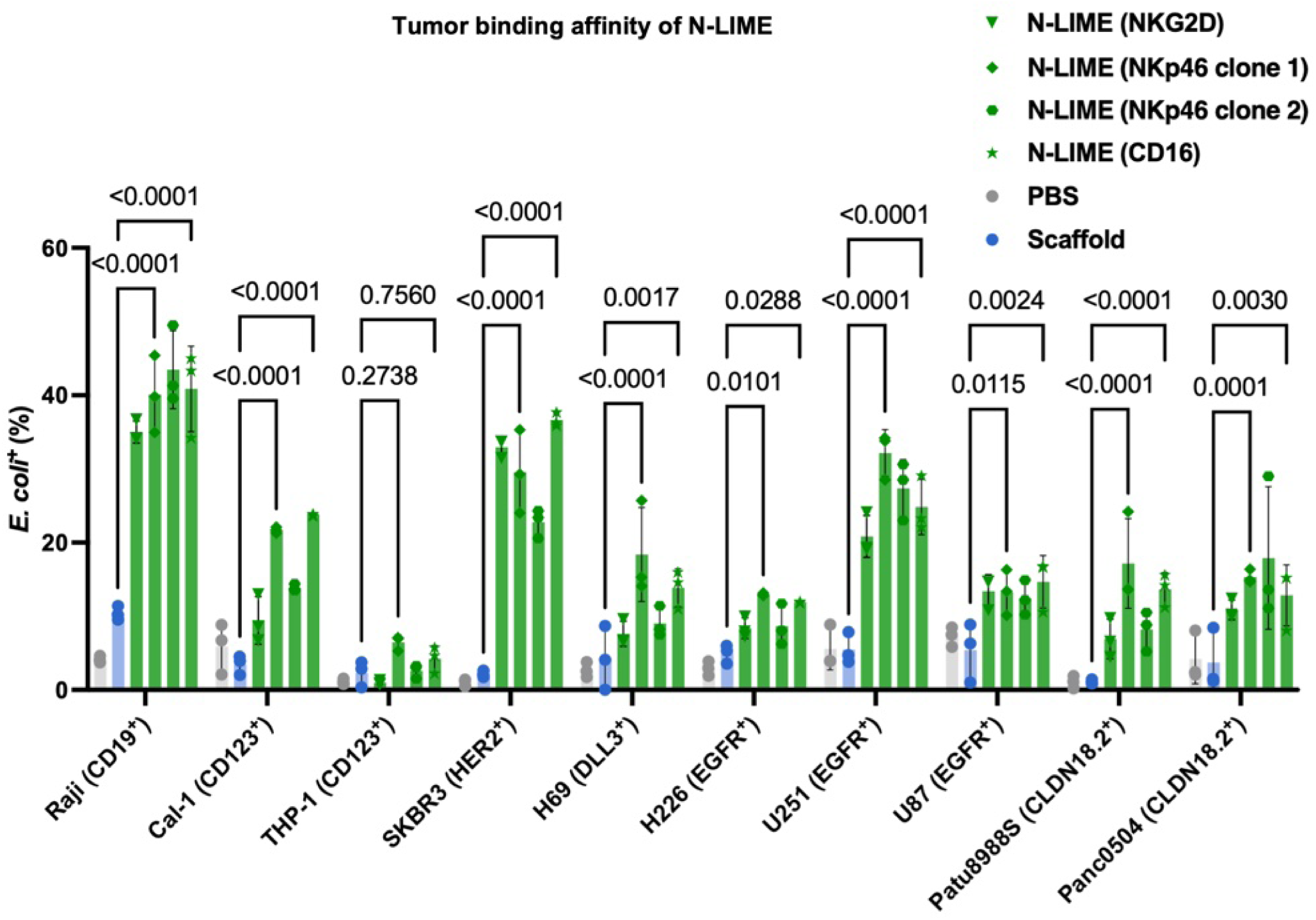
N-LIME exhibit enhanced binding affinity to the tumor cells compared to controls. Binding affinity of N-LIME to the various tumor cells relative to PBS and scaffold controls. Data are presented as mean ± s.d.

**Supplemental Fig. 7.**
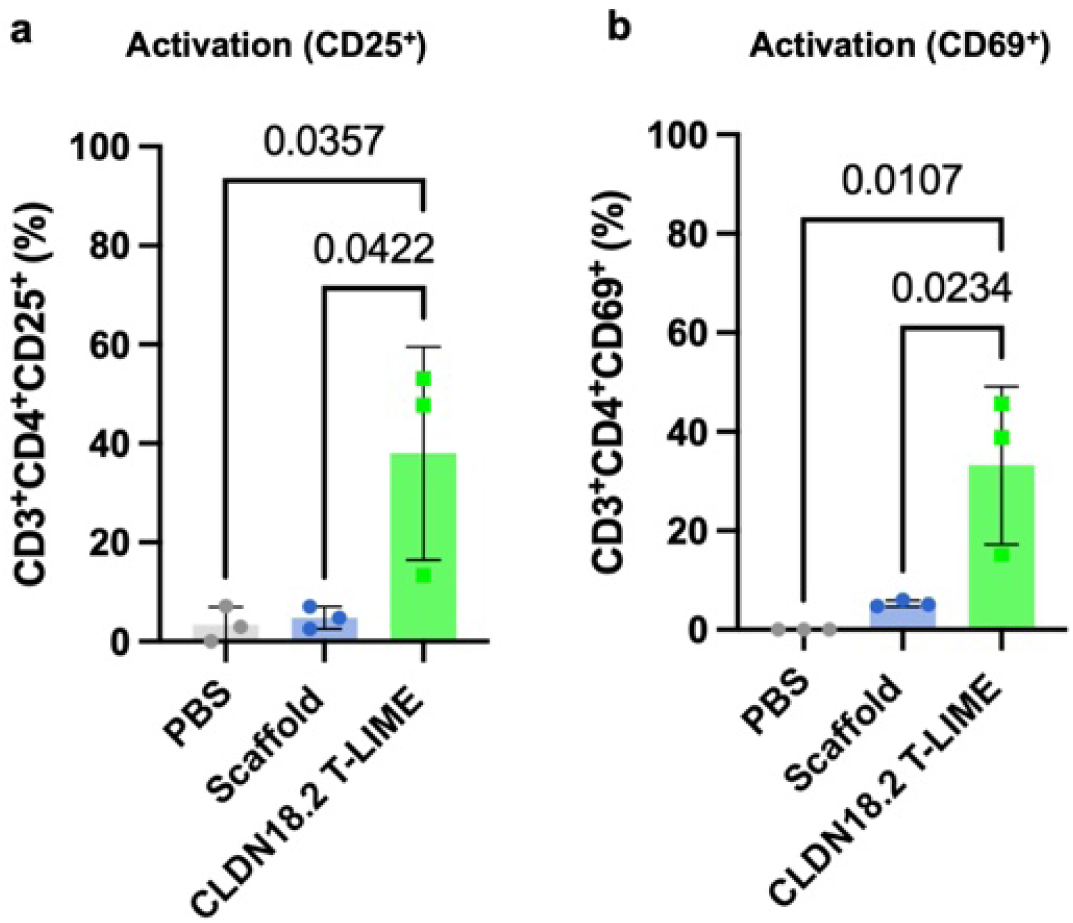
T-LIME induce activation of CD4^+^ T cells in the presence of patient derived CLDN18.2^+^ PDAC organoids. **a,** Activation stage of CD4^+^ T cells, assessed by CD25 (**a**) and CD69 (**b**) in the PDAC organoid co-culture assay, assessed by flow cytometry. Statistical significance was determined using one-way ANOVA test with Tukey’s post hoc test. Data represent mean ± s.d..

**Extended Data Fig. 4.**
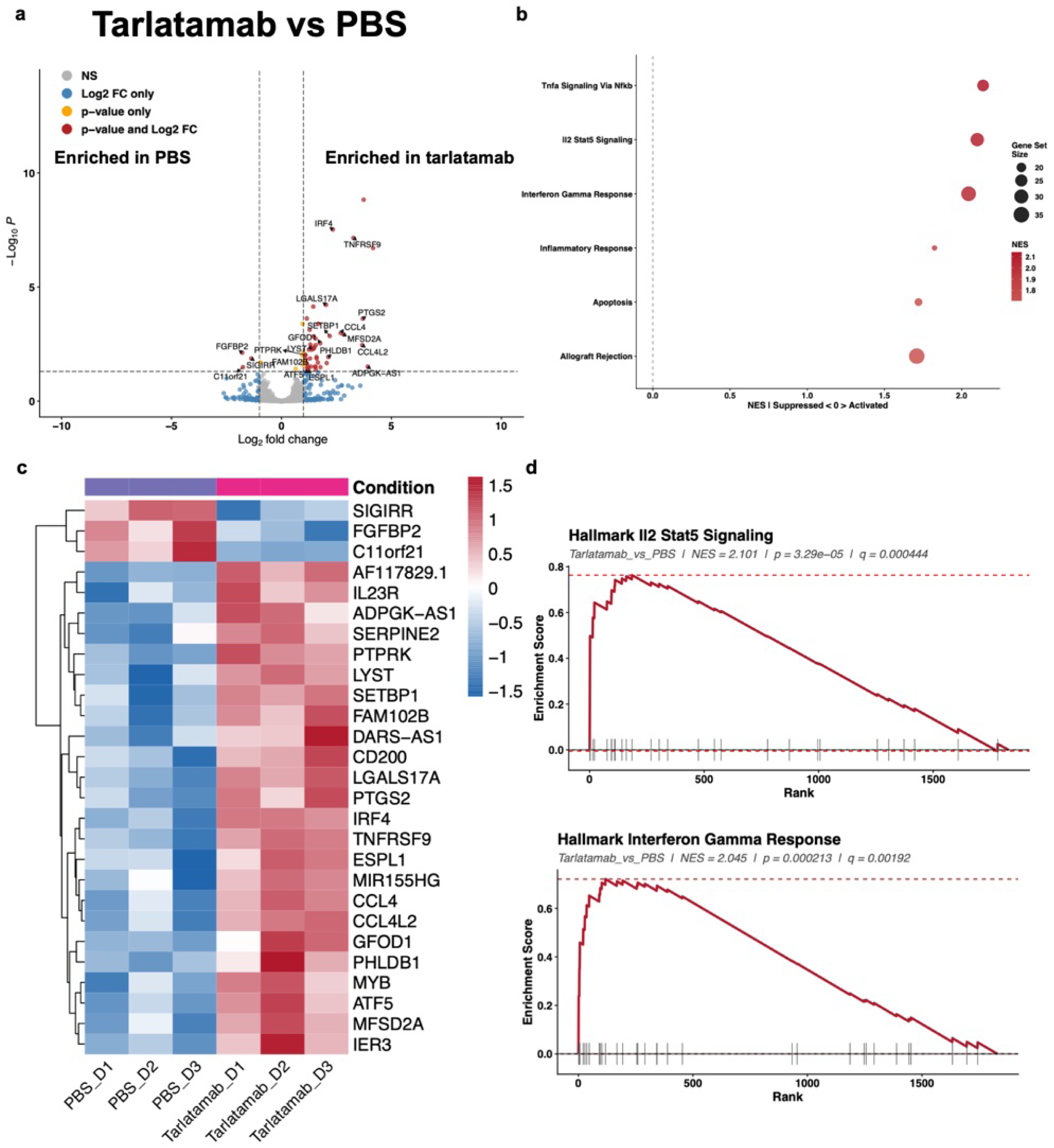
Transcriptional effects of tarlatamab treatment relative to PBS in T cells. Human T cells were treated with soluble DLL3-targeting T-cell engager (tarlatamab) or PBS and analyzed by bulk RNA-seq. **a,** Volcano plot showing differentially expressed genes between tarlatamab- and PBS-treated T cells. **b,** Heatmap of differentially expressed genes. **c, d,** Gene-set enrichment/pathway analyses showing transcriptional programs altered by tarlatamab.

**Extended Data Fig. 5.**
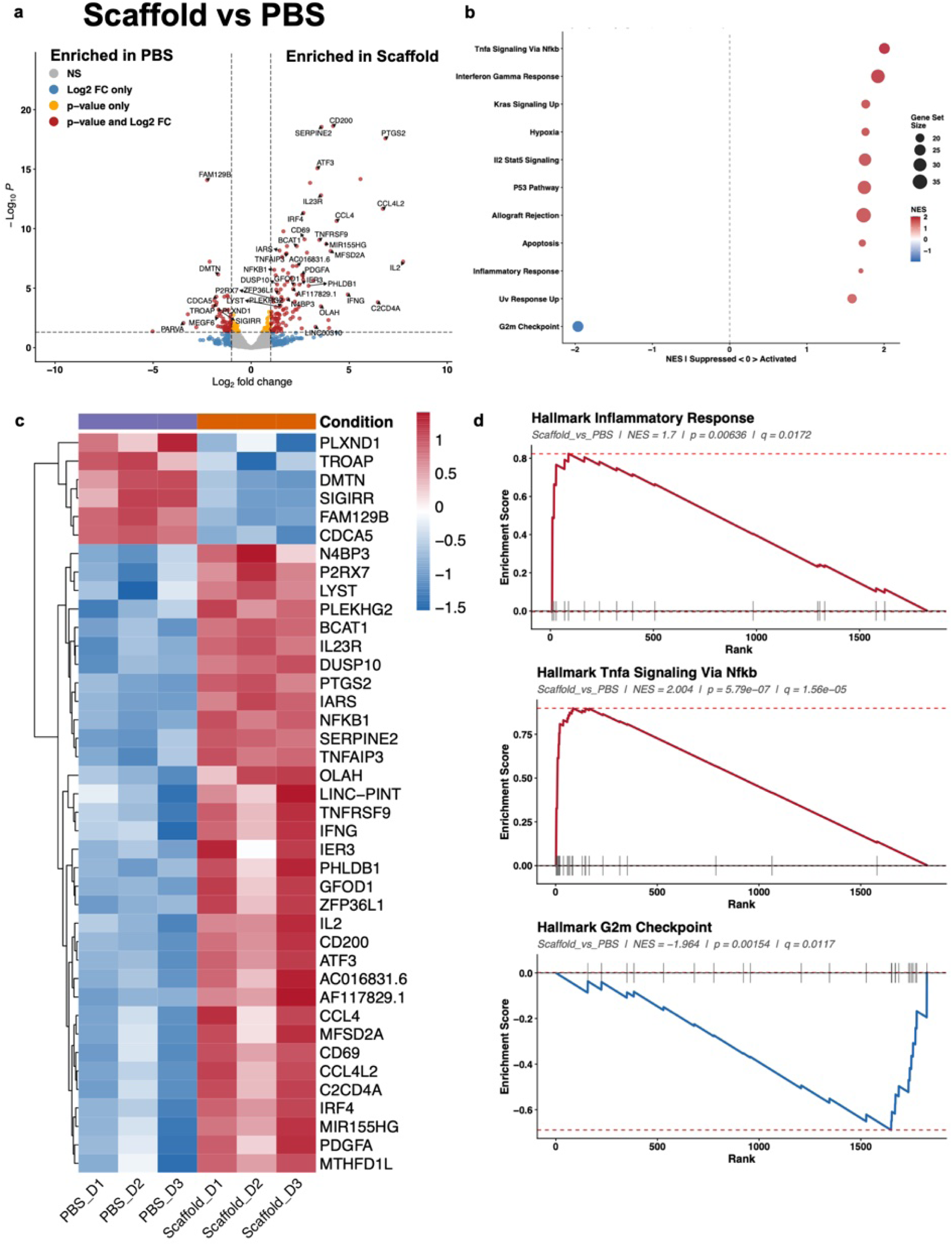
Transcriptional effects of scaffold bacteria relative to PBS in T cells. Human T cells were treated with scaffold bacteria or PBS and analyzed by bulk RNA-seq. **a,** Volcano plot showing differentially expressed genes between scaffold- and PBS-treated T cells. **b,** Heatmap of differentially expressed genes. **c, d,** Gene-set enrichment/pathway analyses showing pathways modulated by scaffold bacteria.

**Supplemental Fig. 8.**
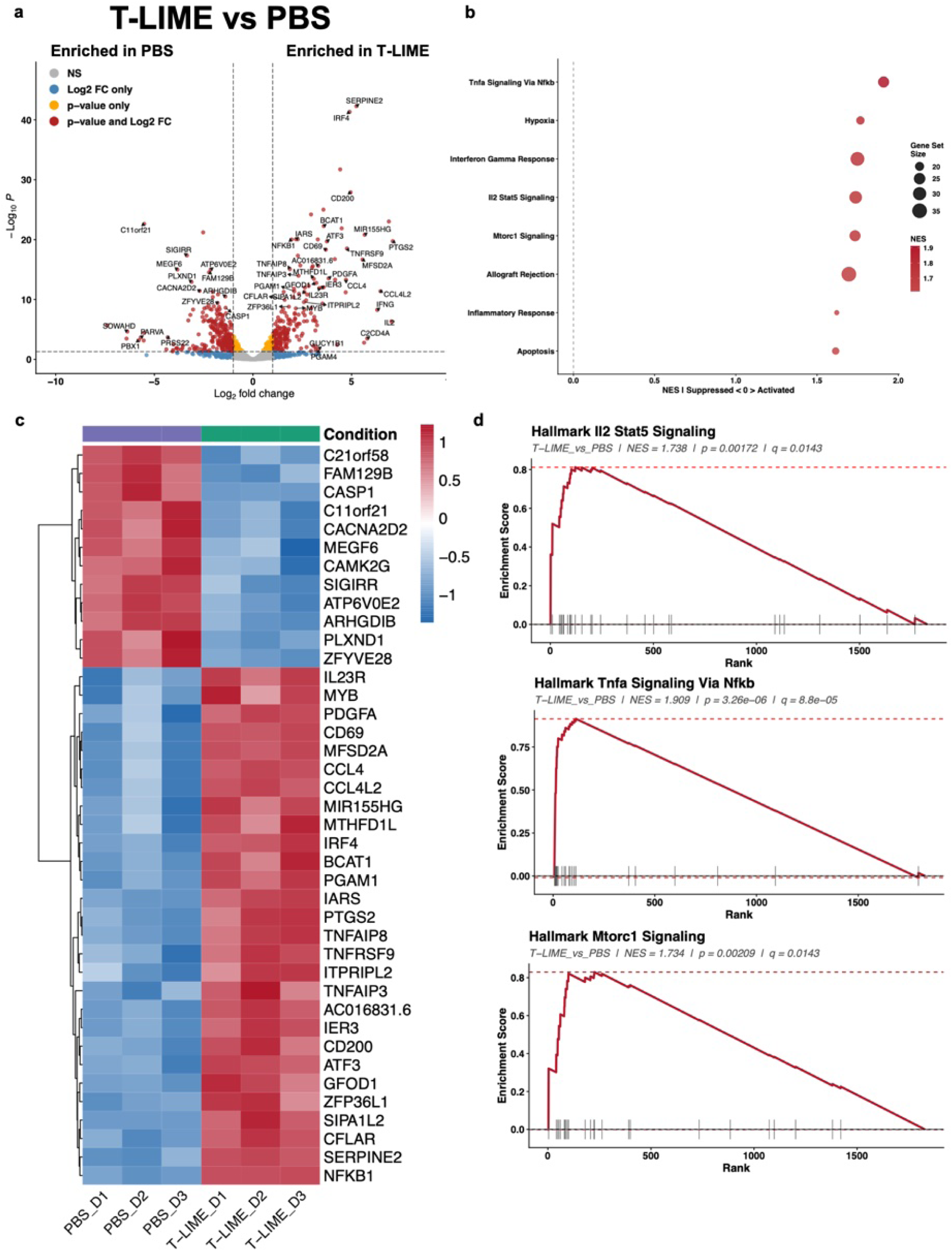
Transcriptional effects of DLL3 T-LIME relative to PBS in T cells. Human T cells were treated with DLL3 T-LIME or PBS and analyzed by bulk RNA-seq. **a,** Volcano plot showing differentially expressed genes between DLL3 T-LIME- and PBS-treated T cells. **b,** Heatmap of differentially expressed genes. **c, d,** Gene-set enrichment/pathway analyses showing pathways modulated by DLL3 T-LIME.

**Supplemental Fig. 9.**
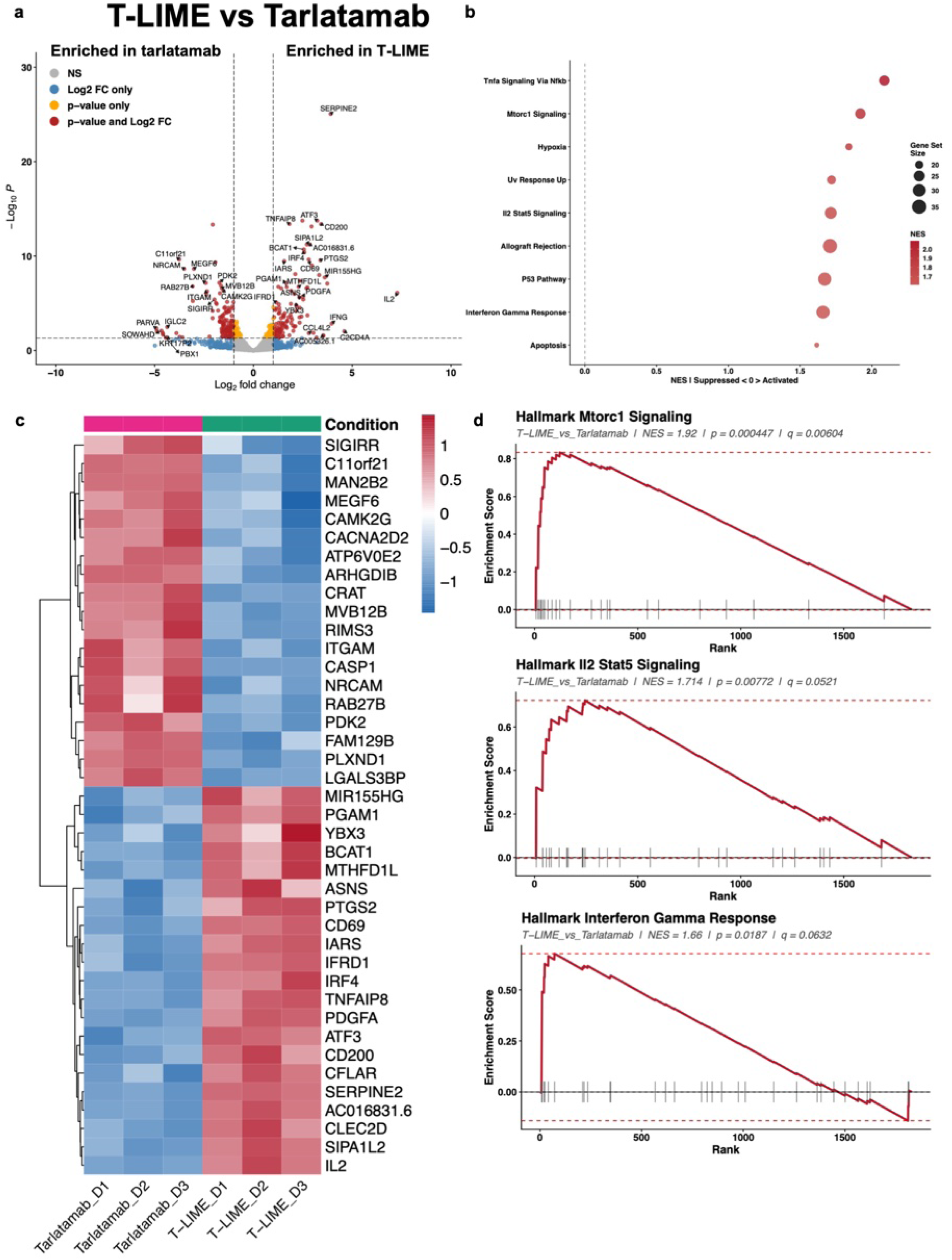
DLL3 T-LIME induces a transcriptional state distinct from tarlatamab in T cells. Human T cells treated with DLL3 T-LIME or soluble DLL3-targeting T-cell engager (tarlatamab) were compared by bulk RNA-seq. **a,** Volcano plot showing differentially expressed genes between T-LIME- and tarlatamab-treated T cells. **b,** Heatmap of differentially expressed genes. **c, d,** Gene-set enrichment/pathway analyses identifying transcriptional programs differentially regulated by DLL3 T-LIME treatment versus tarlatamab treatment.

**Supplemental Fig. 10.**
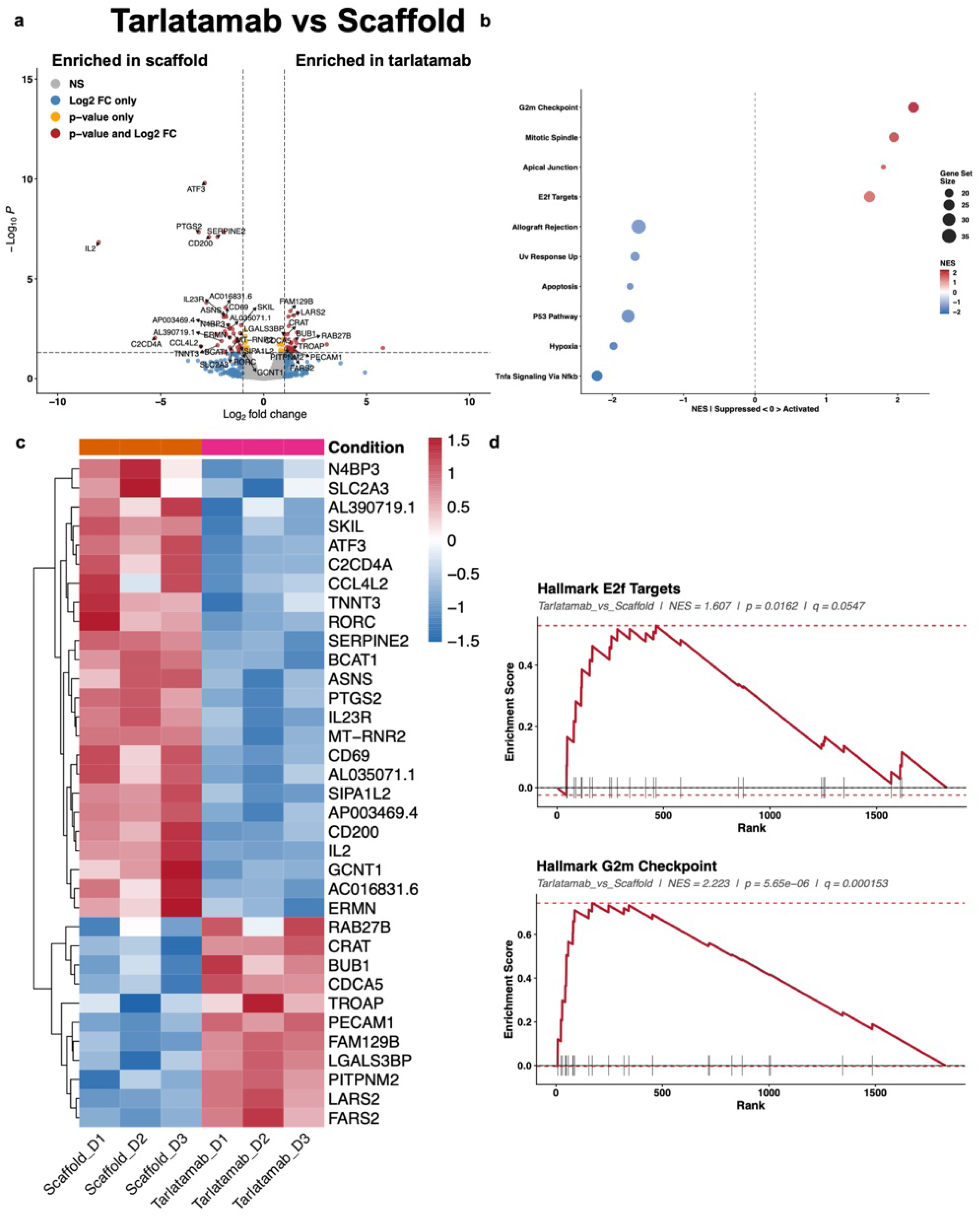
Tarlatamab and scaffold bacteria induce distinct T-cell transcriptional states. Human T cells treated with tarlatamab or scaffold bacteria were compared by bulk RNA-seq. **a,** Volcano plot showing differentially expressed genes between tarlatamab- and scaffold-treated T cells. **b,** Heatmap of differentially expressed genes. **c, d,** Gene-set enrichment/pathway analyses comparing immune pathways induced by soluble engager activity versus bacterial scaffold exposure.

**Supplemental Fig. 11.**
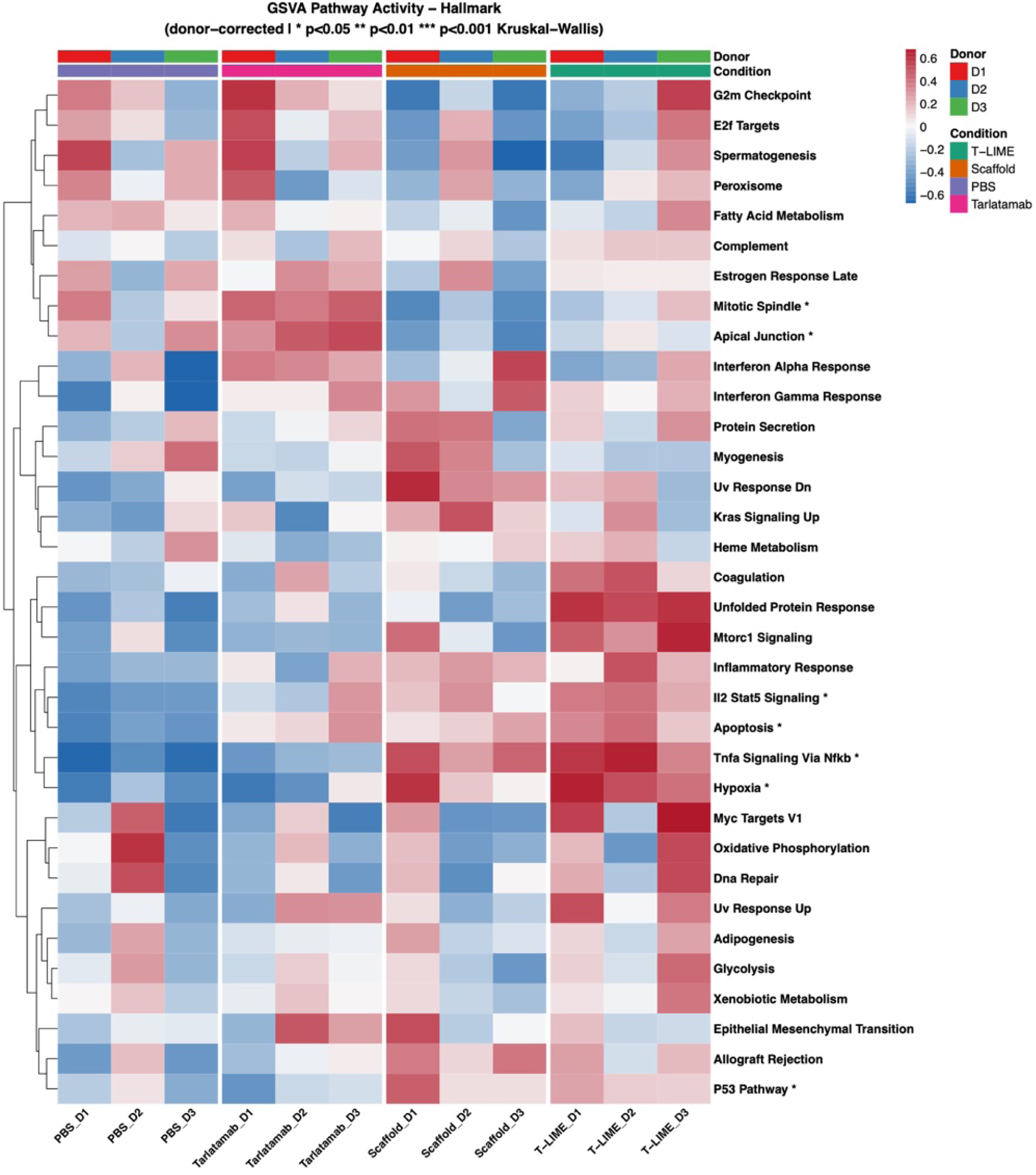
Hallmark GSVA analysis across treatment conditions. Gene set variation analysis (GSVA) of Hallmark pathways across human T cells treated with PBS, scaffold bacteria, tarlatamab or DLL3 T-LIME.

**Supplemental Fig. 12.**
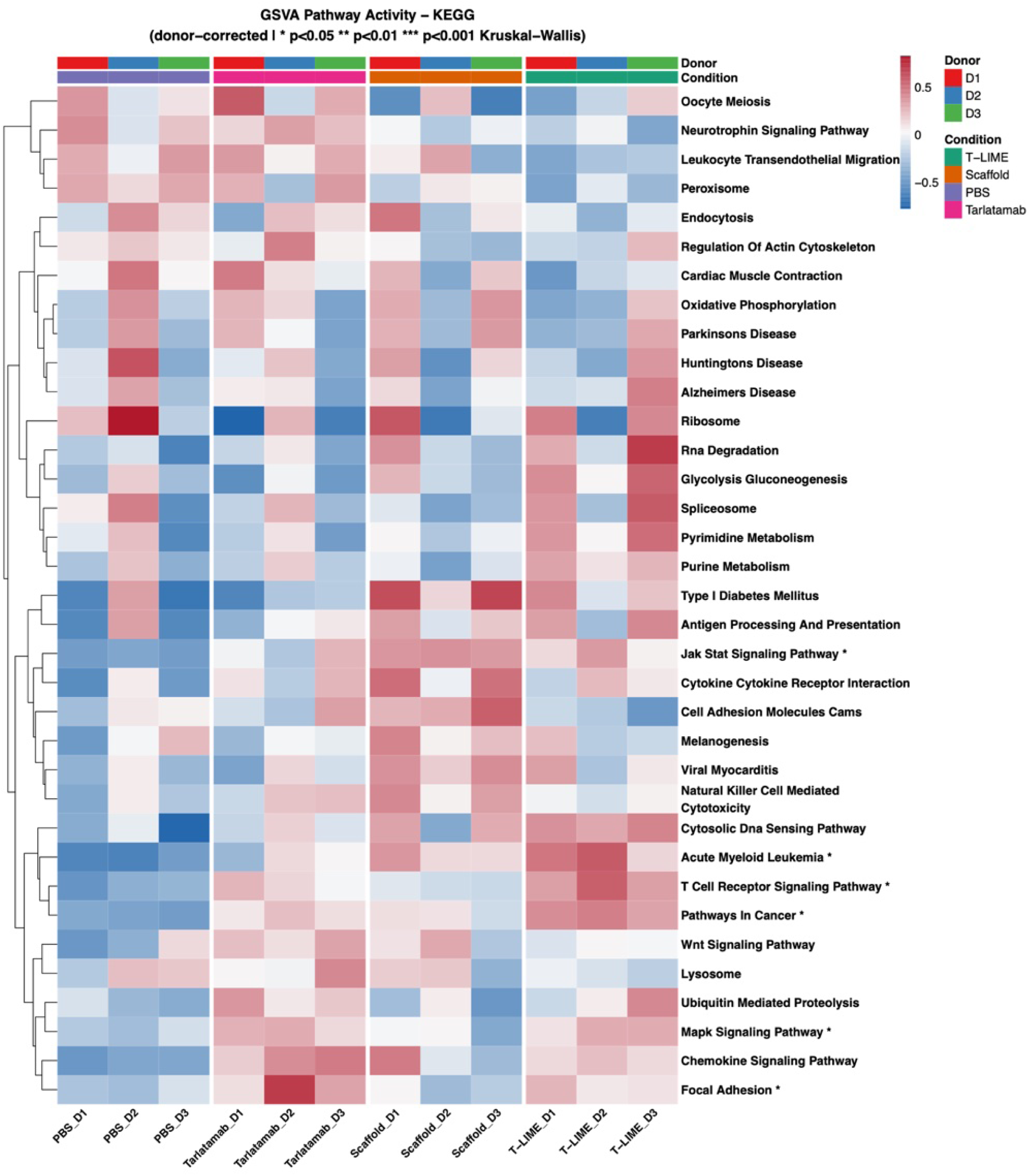
KEGG GSVA analysis across treatment conditions. Gene set variation analysis (GSVA) of KEGG pathways across human T cells treated with PBS, scaffold bacteria, tarlatamab or DLL3 T-LIME.

**Supplemental Fig. 13.**
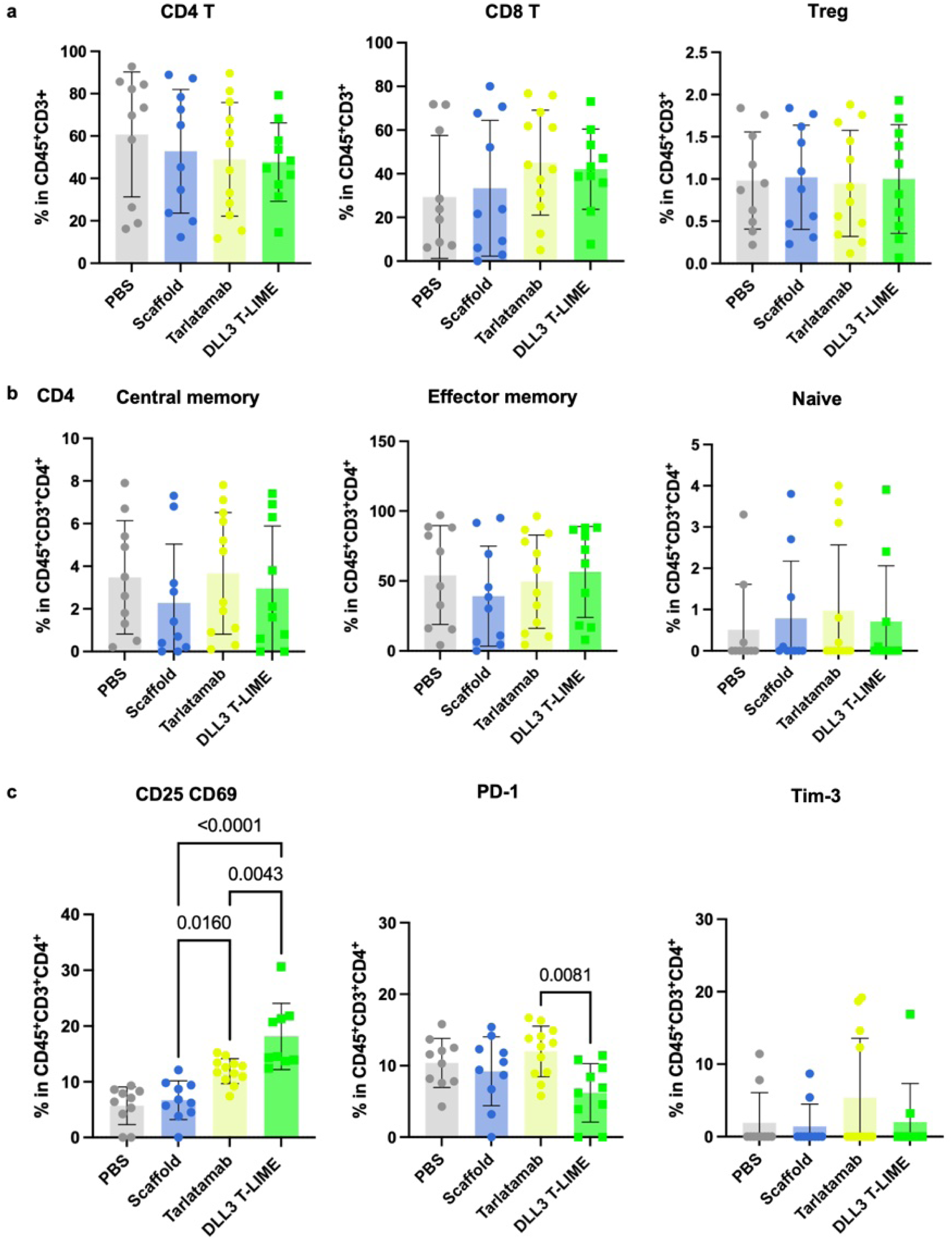
DLL3 T-LIME remodels the phenotype of tumor-infiltrating human CD4^+^ T cells. Human T cells from H69 subcutaneous xenografts described in Fig. 3d**-f** and analyzed by flow cytometry at the experimental endpoint. **a**, Frequencies of human CD4^+^ T cells, CD8^+^ T cells and regulatory T (Treg) cells within the tumor microenvironment (TME). **b**, Frequencies of naive, central memory and effector memory CD4^+^ T-cell subsets. **c**, Frequencies of CD25^+^, CD69^+^, PD-1^+^ and Tim-3^+^ populations among CD4^+^ T cells following treatment with PBS, scaffold bacteria, DLL3-targeting T-cell engager/tarlatamab or DLL3 T-LIME. Data are presented as mean ± s.d. Statistical significance was determined by one-way ANOVA.

**Supplemental Fig. 14.**
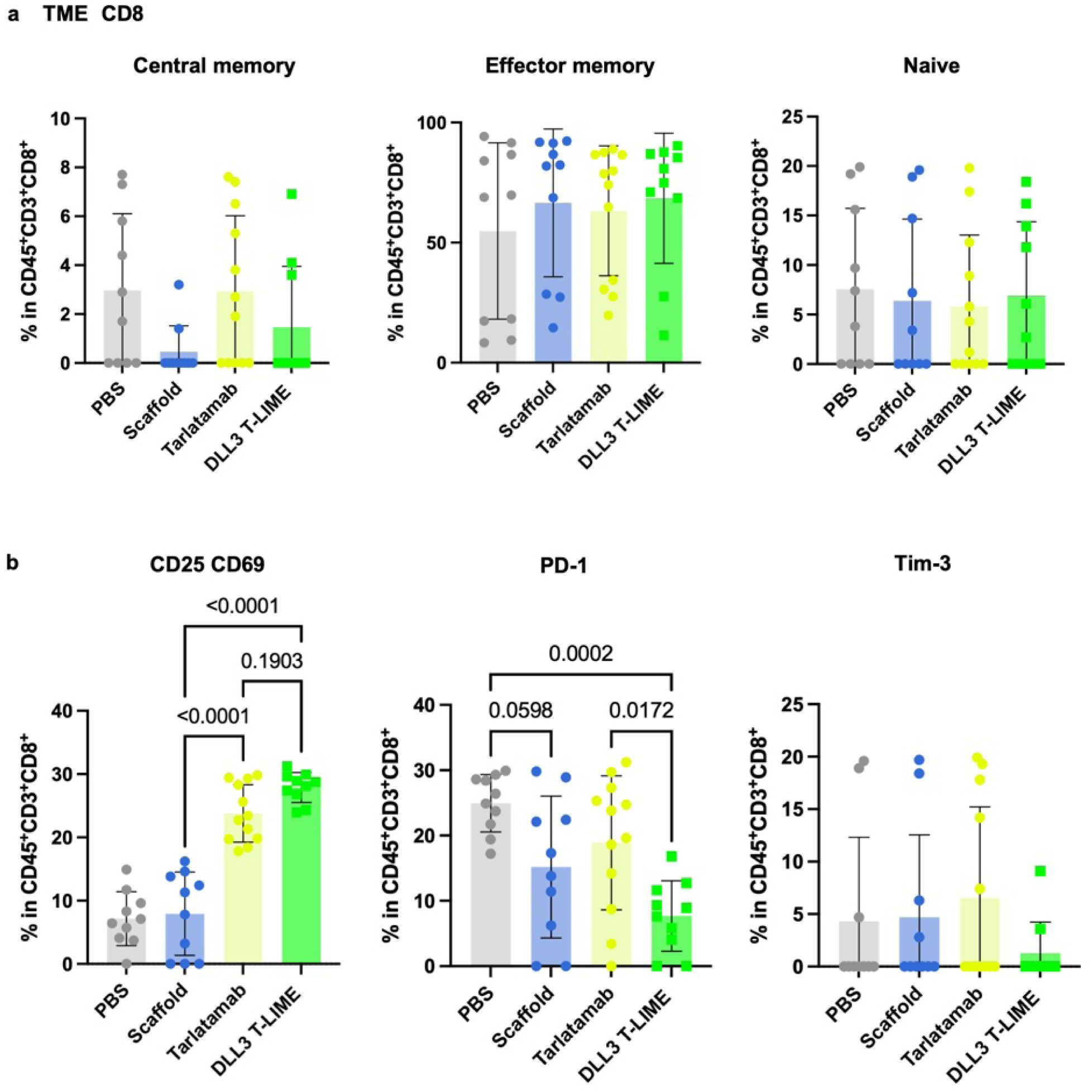
DLL3 T-LIME remodels the phenotype of tumor-infiltrating human CD8^+^ T cells. Human T cells from H69 subcutaneous xenografts described in Fig. 3d**-f** and analyzed by flow cytometry at the experimental endpoint. **a,** Frequencies of naive, central memory and effector memory CD8^+^ T-cell subsets. **b,** Frequencies of CD25^+^, CD69^+^, PD-1^+^ and Tim-3^+^ populations among CD8^+^ T cells following treatment with PBS, scaffold bacteria, the DLL3-targeting T-cell engager tarlatamab or DLL3 T-LIME. Data are presented as mean ± s.d. Statistical significance was determined by one-way ANOVA.

**Supplemental Fig. 15.**
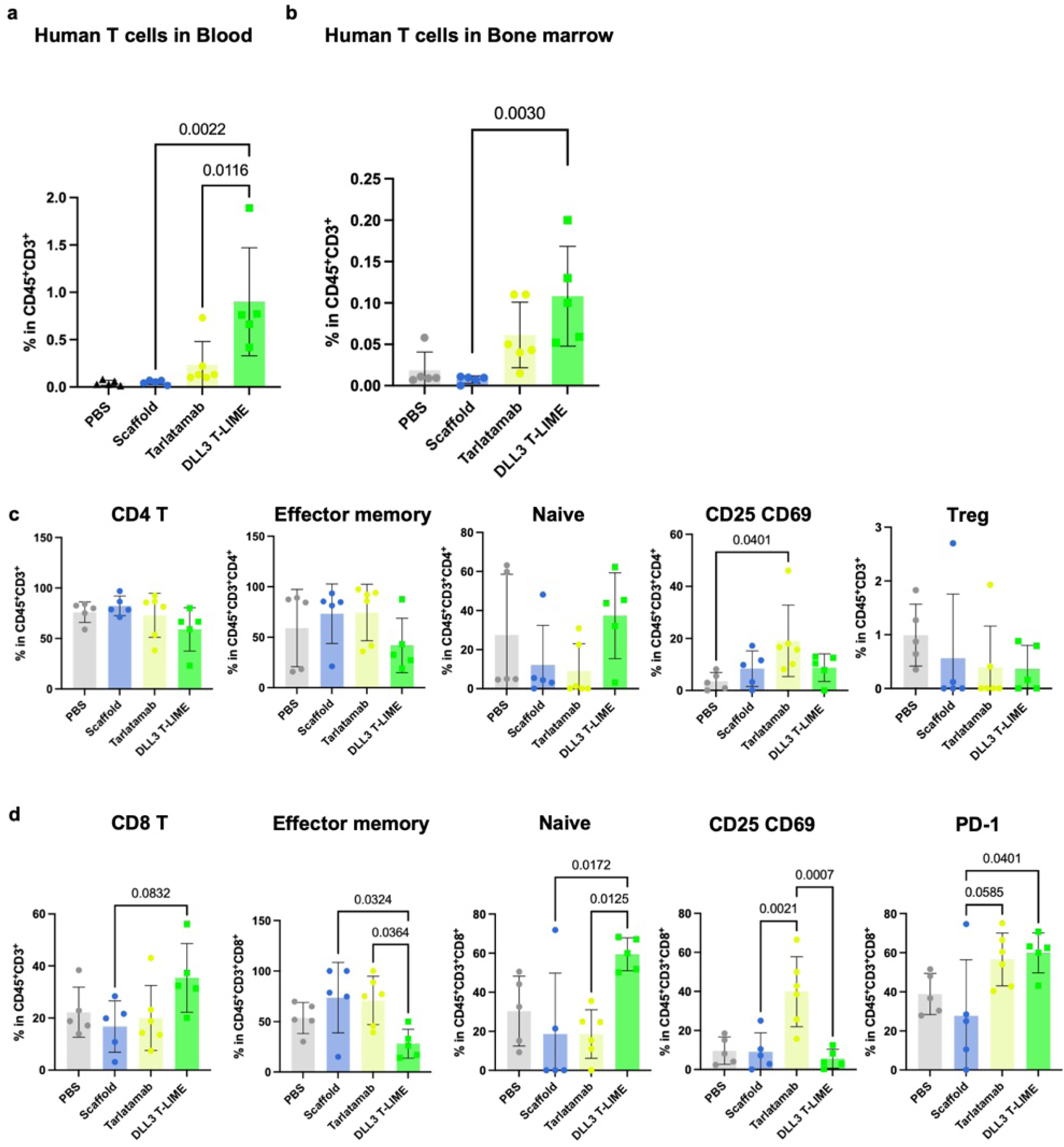
Frequency and phenotypic characterization of human T cells in peripheral blood and bone marrow. Human T cells from H69 subcutaneous xenograft-bearing mice described in Fig. 3d**–f** were analyzed by flow cytometry at the experimental endpoint. **a,** Frequency of human CD3^+^ T cells among peripheral blood mononuclear cells (PBMCs). **b,** Frequency of human CD3^+^ T cells in the bone marrow. **c,** Frequency of CD4^+^ T cells among human CD3^+^ T cells in the bone marrow and frequencies of naive, central memory and effector memory CD4^+^ T-cell subsets and CD25^+^, CD69^+^ and PD-1^+^ CD4+ T cells. **d,** Frequency of CD8^+^ T cells among human CD3^+^ T cells in the bone marrow and frequencies of naive, central memory and effector memory CD8^+^ T-cell subsets and CD25^+^, CD69^+^ and PD-1^+^ CD8^+^ T cells following treatment with PBS, scaffold bacteria, the DLL3-targeting T-cell engager tarlatamab or DLL3 T-LIME. Data are presented as mean ± s.d. Statistical significance was determined by one-way ANOVA.

**Supplemental Fig. 16.**
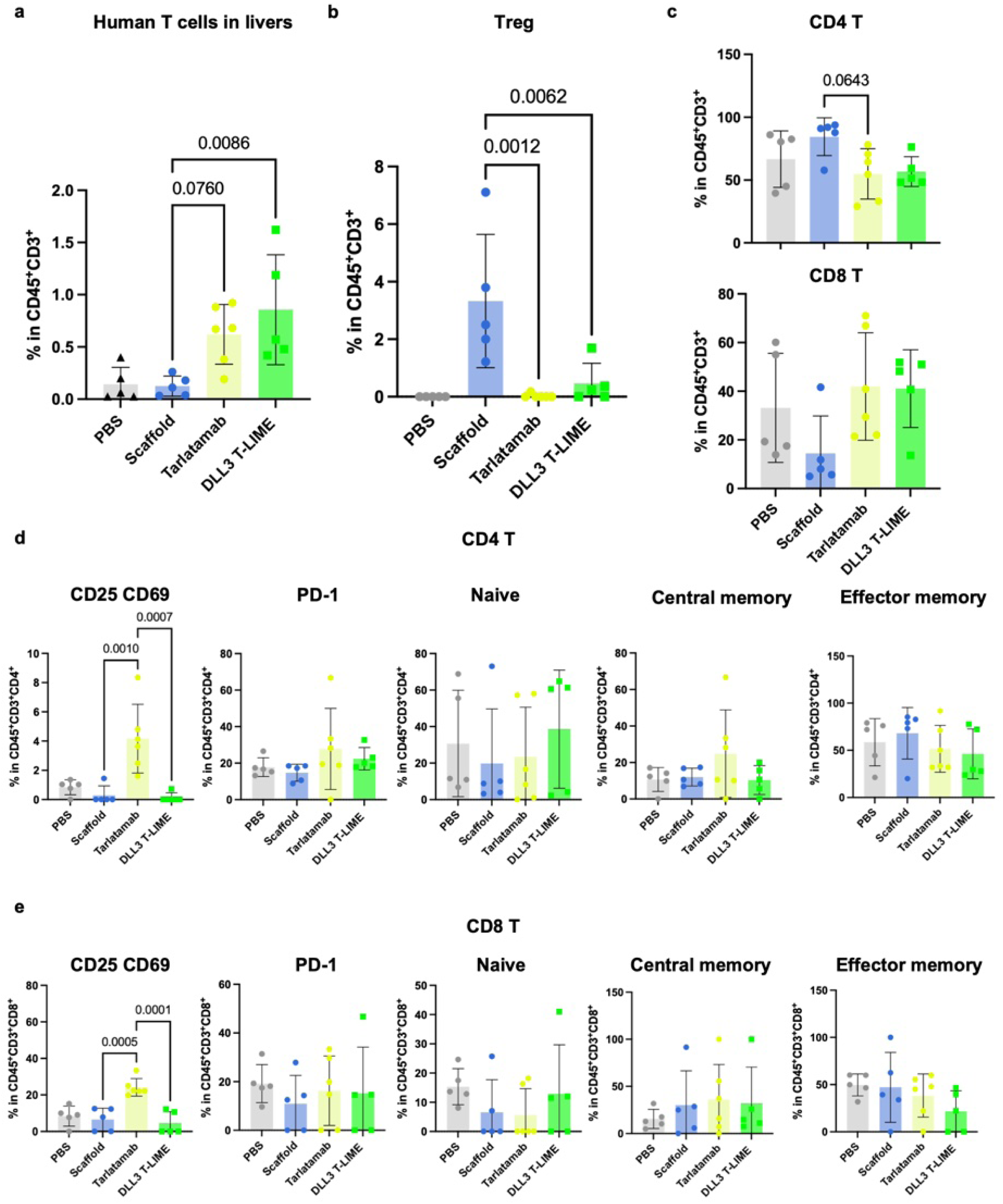
Frequency and phenotypic characterization of human T cells in the liver. Human T cells from H69 subcutaneous xenograft-bearing mice described in Fig. 3d**–f** were analyzed by flow cytometry at the experimental endpoint. **a,** Frequency of human CD3^+^ T cells in the liver. **b,** Frequency of regulatory T (Treg) cells among human CD3^+^ T cells in the liver. **c,** Frequency of CD4^+^ T and CD8^+^ T cells among human CD3^+^ T cells in the liver. **d, e,** Frequencies of naive, central memory and effector memory CD4^+^ T-cell subsets and CD25^+^, CD69^+^ and PD-1^+^ CD4^+^ T cells (**d**) and frequencies of naive, central memory and effector memory CD8^+^ T-cell subsets and CD25^+^, CD69^+^ and PD-1^+^ CD8^+^ T cells (**e**) Following treatment with PBS, scaffold bacteria, the DLL3-targeting T-cell engager tarlatamab or DLL3 T-LIME. Data are presented as mean ± s.d. Statistical significance was determined by one-way ANOVA.

**Supplemental Fig. 17.**
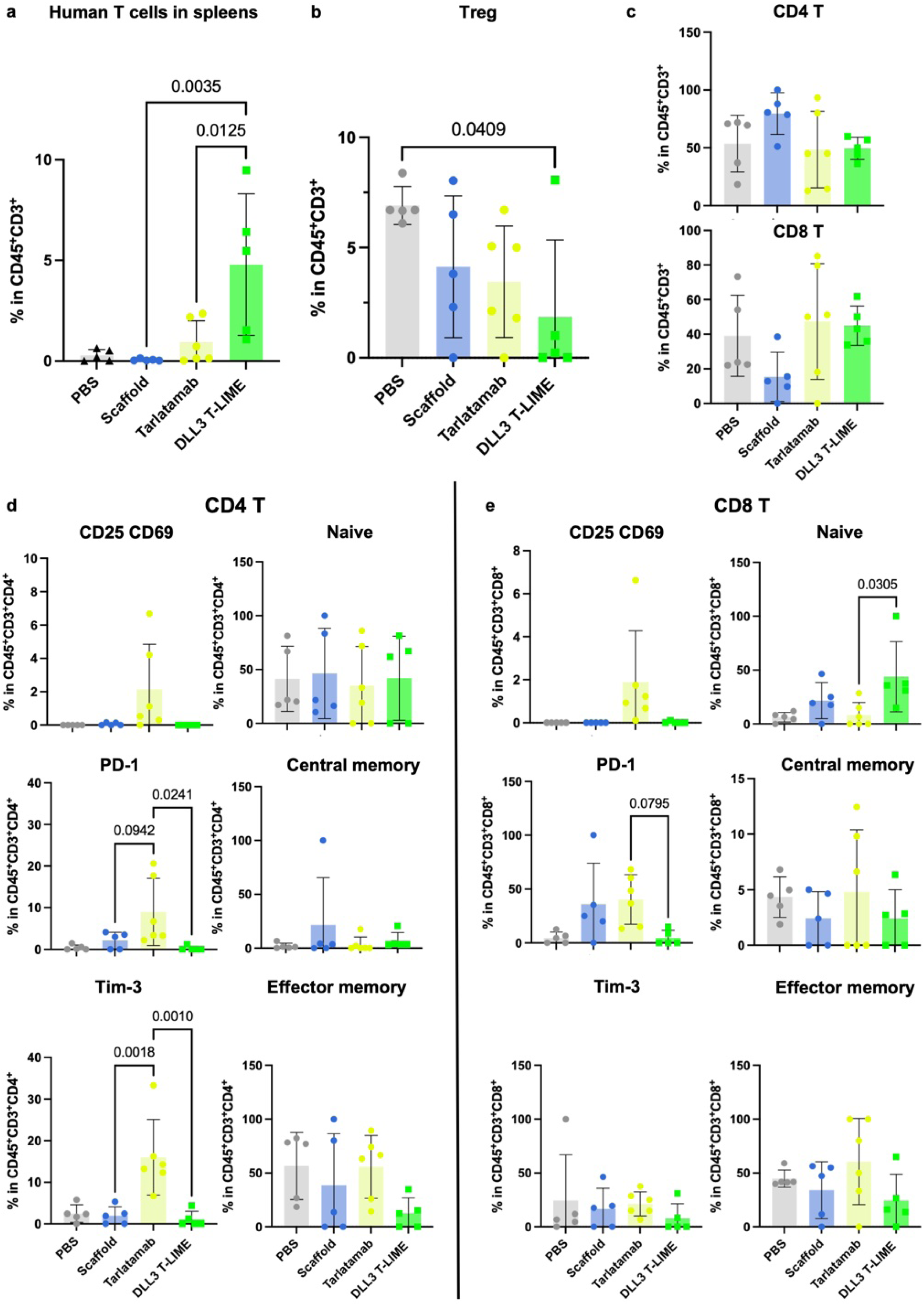
Frequency and phenotypic characterization of human T cells in the spleen. Human T cells from H69 subcutaneous xenograft-bearing mice described in Fig. 3d**–f** were analyzed by flow cytometry at the experimental endpoint. **a,** Frequency of human CD3^+^ T cells in the spleen. **b,** Frequency of regulatory T (Treg) cells among human CD3^+^ T cells in the spleen. **c,** frequencies of CD4^+^ and CD8^+^ T cells among human CD3^+^ T cells in the spleen. **d, e,** Frequencies of naive, central memory and effector memory CD4^+^ T-cell subsets and CD25^+^, CD69^+^, PD-1^+^ and Tim-3^+^ CD4^+^ T cells (**d**) and frequencies of naive, central memory and effector memory CD8^+^ T-cell subsets and CD25^+^, CD69^+^, PD-1^+^ and Tim-3^+^ CD8^+^ T cells (**e**) following treatment with PBS, scaffold bacteria, the DLL3-targeting T-cell engager tarlatamab or DLL3 T-LIME. Data are presented as mean ± s.d. Statistical significance was determined by one-way ANOVA.

**Supplemental Fig. 18.**
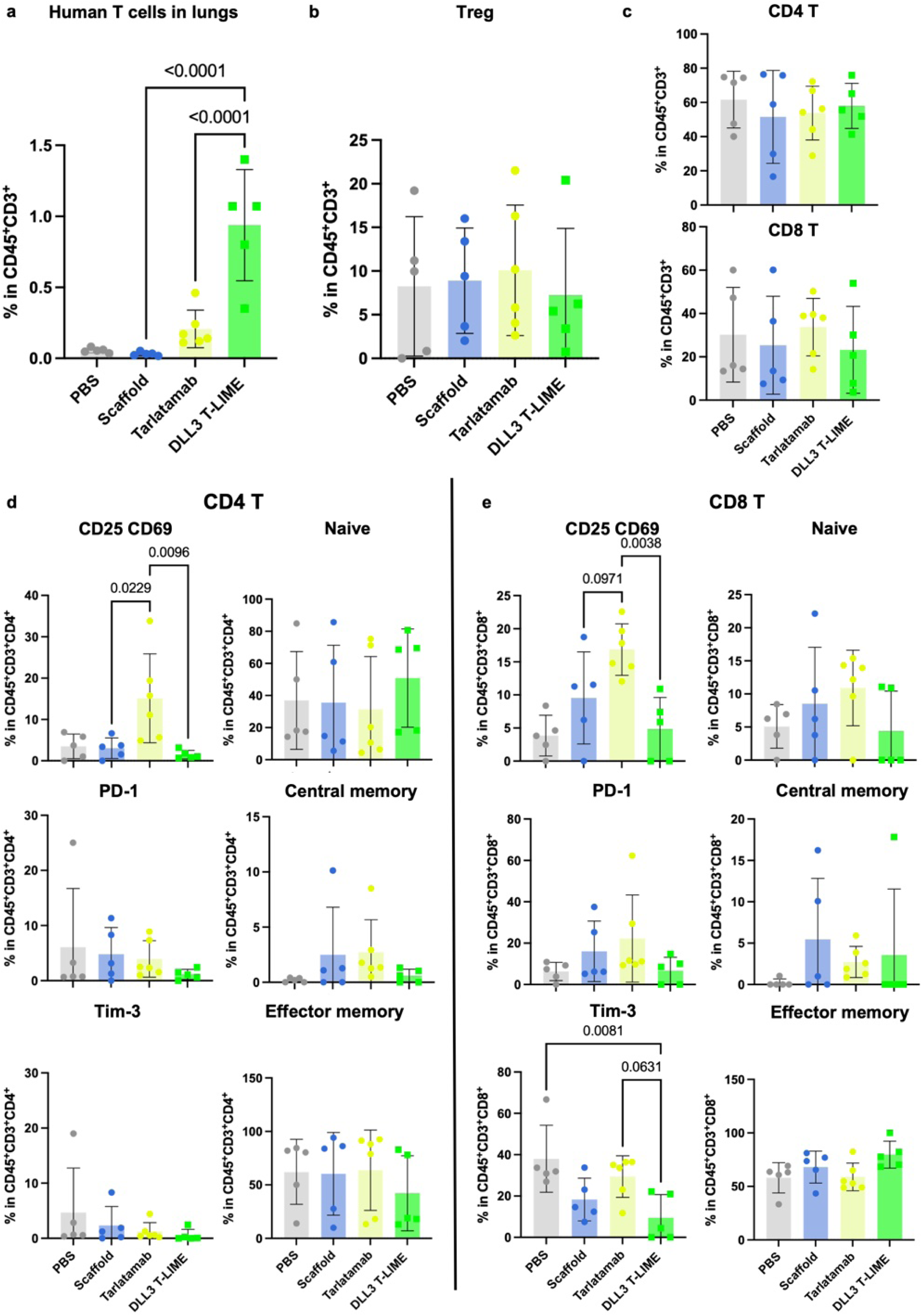
Frequency and phenotypic characterization of human T cells in the lung. Human T cells from H69 subcutaneous xenograft-bearing mice described in Fig. 3d**–f** were analyzed by flow cytometry at the experimental endpoint. **a,** Frequency of human CD3^+^ T cells in the lung. **b,** Frequency of regulatory T (Treg) cells among human CD3^+^ T cells in the lung. **c,** Frequencies of CD4^+^ and CD8^+^ T cells among human CD3^+^ T cells in the lung. **d, e,** Frequencies of naive, central memory and effector memory CD4^+^ T-cell subsets and CD25^+^, CD69^+^, PD-1^+^ and Tim-3^+^ CD4^+^ T cells (**d**) and frequencies of naive, central memory and effector memory CD8^+^ T-cell subsets and CD25^+^, CD69^+^, PD-1^+^ and Tim-3^+^ CD8^+^ T cells (**e**) Following treatment with PBS, scaffold bacteria, the DLL3-targeting T-cell engager tarlatamab or DLL3 T-LIME. Data are presented as mean ± s.d. Statistical significance was determined by one-way ANOVA.

**Supplemental Fig. 19.**
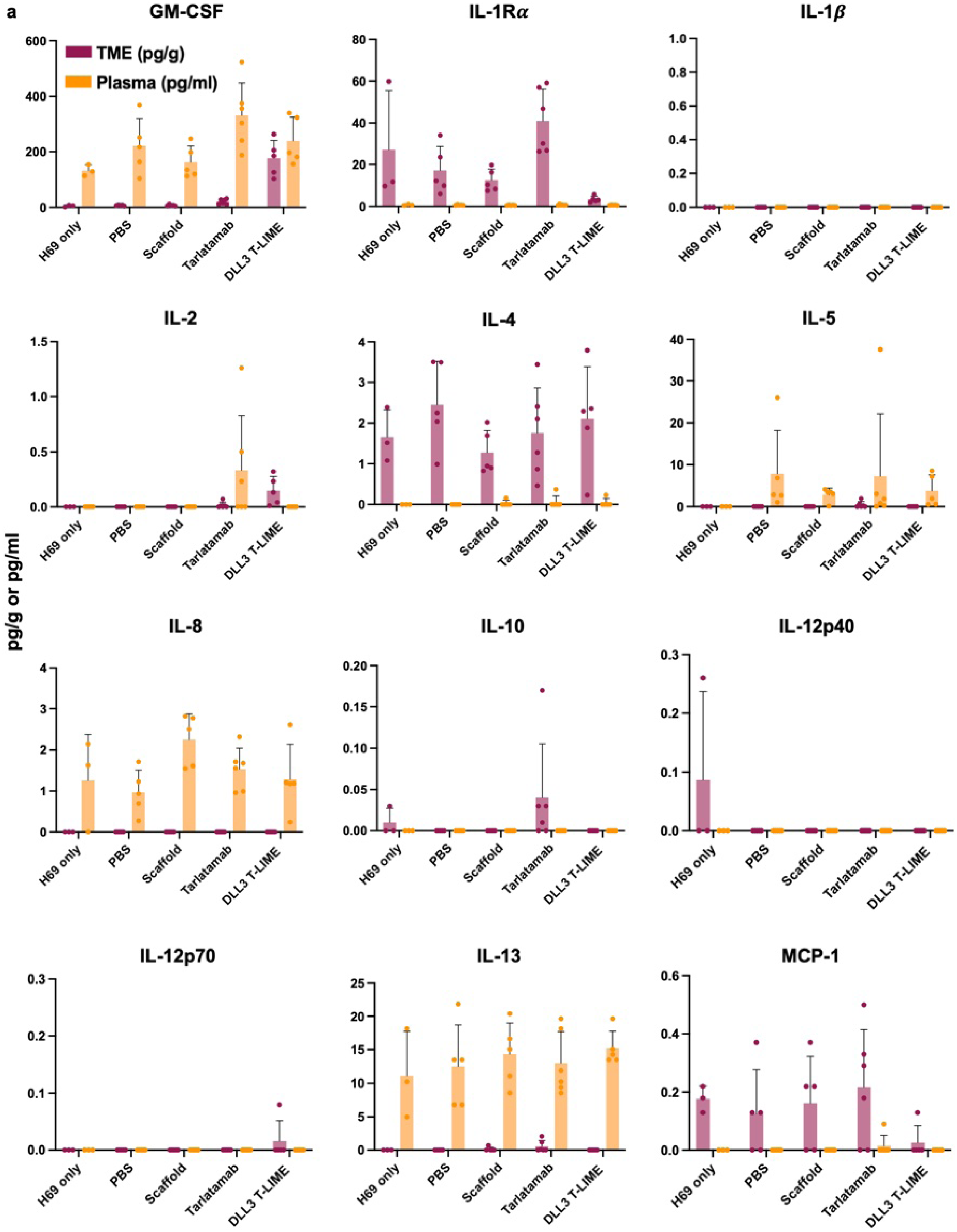
Extended cytokine profiles in the tumor microenvironment and plasma following treatment with DLL3 T-LIME. Cytokine concentrations in tumor tissue and plasma from H69 subcutaneous xenograft-bearing mice described in Fig. 3d**–f** were quantified at the experimental endpoint. **a,** Concentrations of GM-CSF, IL-1Rα, IL-1β, IL-2, IL-4, IL-5, IL-8, IL-10, IL-12p40, IL-12p70, IL-13 and MCP-1 in the tumor microenvironment and plasma following treatment with PBS, scaffold bacteria, the DLL3-targeting T-cell engager tarlatamab or DLL3 T-LIME. Data are presented as mean ± s.d.

**Supplemental Fig. 20.**
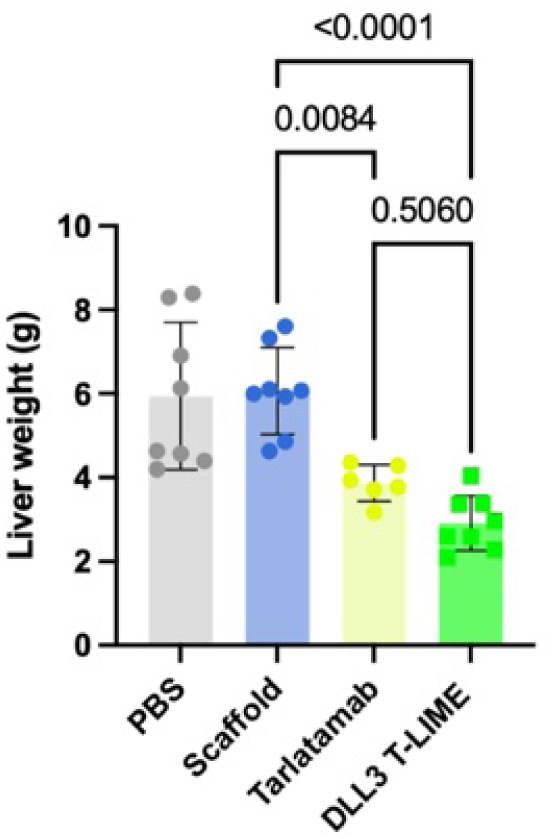
Liver weights following treatment in the H69 liver metastasis model. Liver weights were measured at the experimental endpoint in H69 liver metastasis-bearing mice treated with PBS, scaffold bacteria, the DLL3-targeting T-cell engager tarlatamab or DLL3 T-LIME. Data are presented as mean ± s.d. Statistical significance was determined by one-way ANOVA.

**Supplemental Fig. 21.**
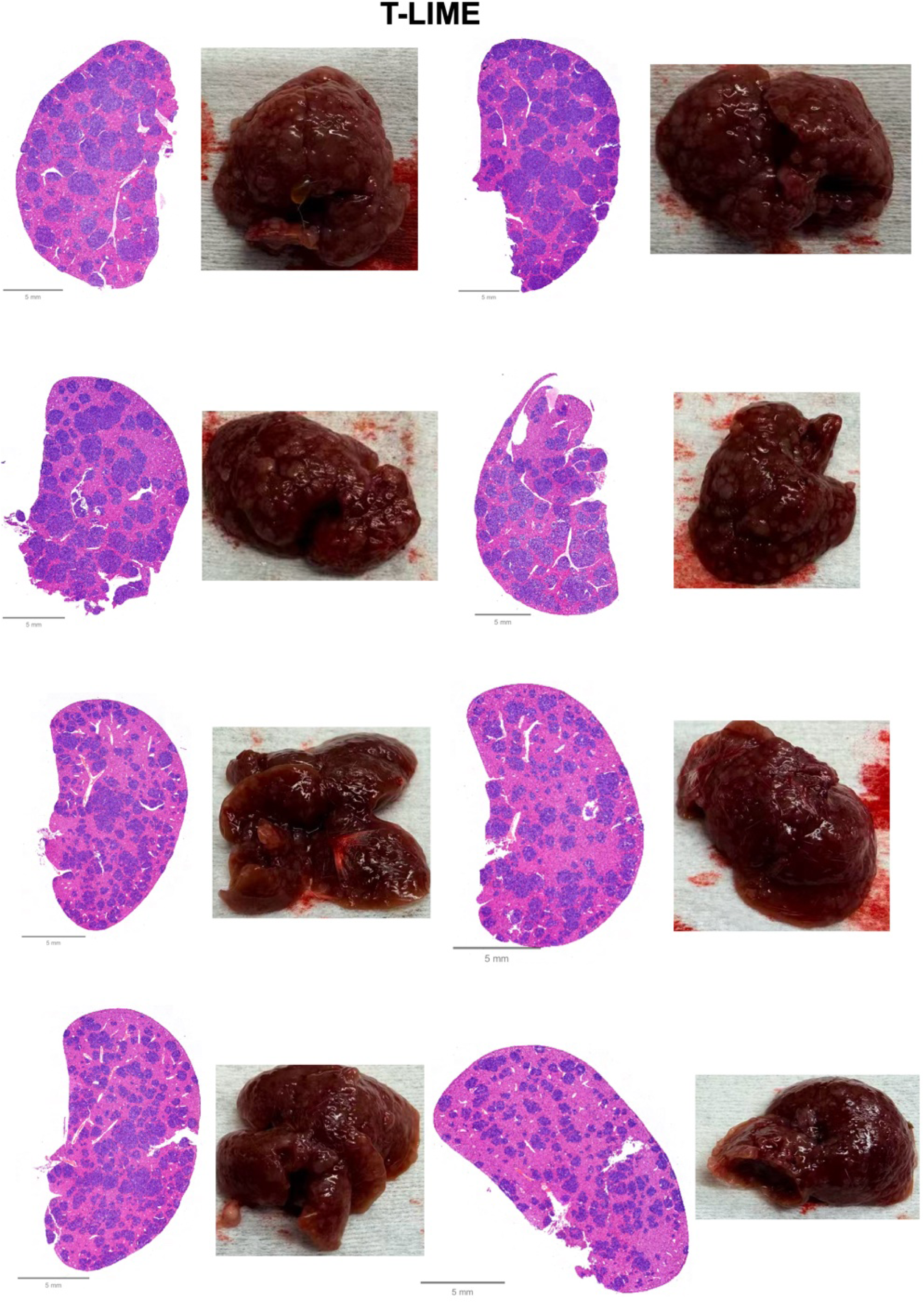
Representative liver histology and gross liver images from DLL3 T-LIME-treated mice. Representative whole-slide H&E-stained liver sections and corresponding gross liver images from H69 liver metastasis-bearing mice treated with DLL3 T-LIME. Scale bars: 5 mm.

**Supplemental Fig. 22.**
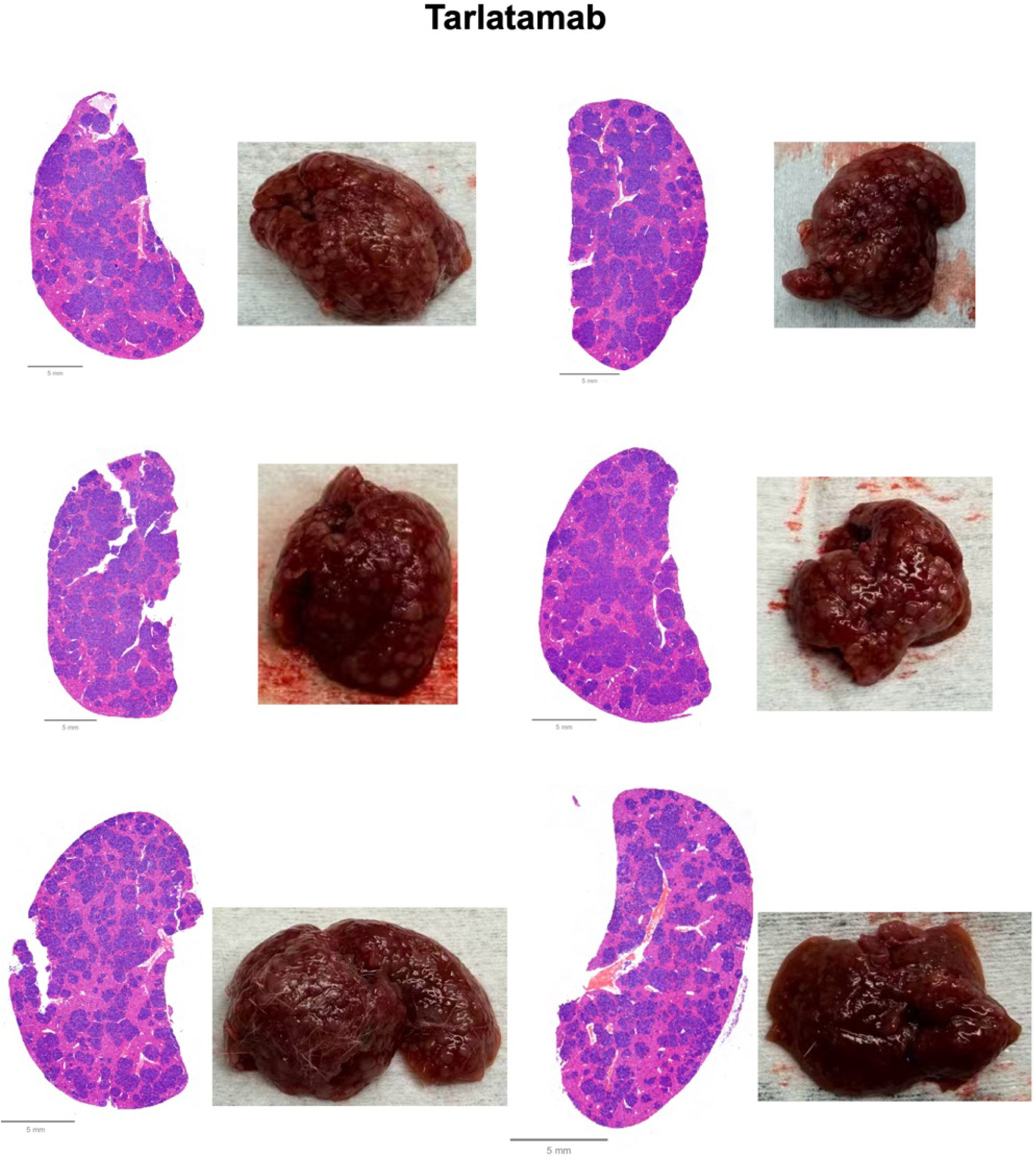
Representative liver histology and gross liver images from tarlatamab-treated mice. Representative whole-slide H&E-stained liver sections and corresponding gross liver images from H69 liver metastasis-bearing mice treated with the DLL3-targeting T-cell engager tarlatamab. Scale bars: 5 mm.

**Supplemental Fig. 23.**
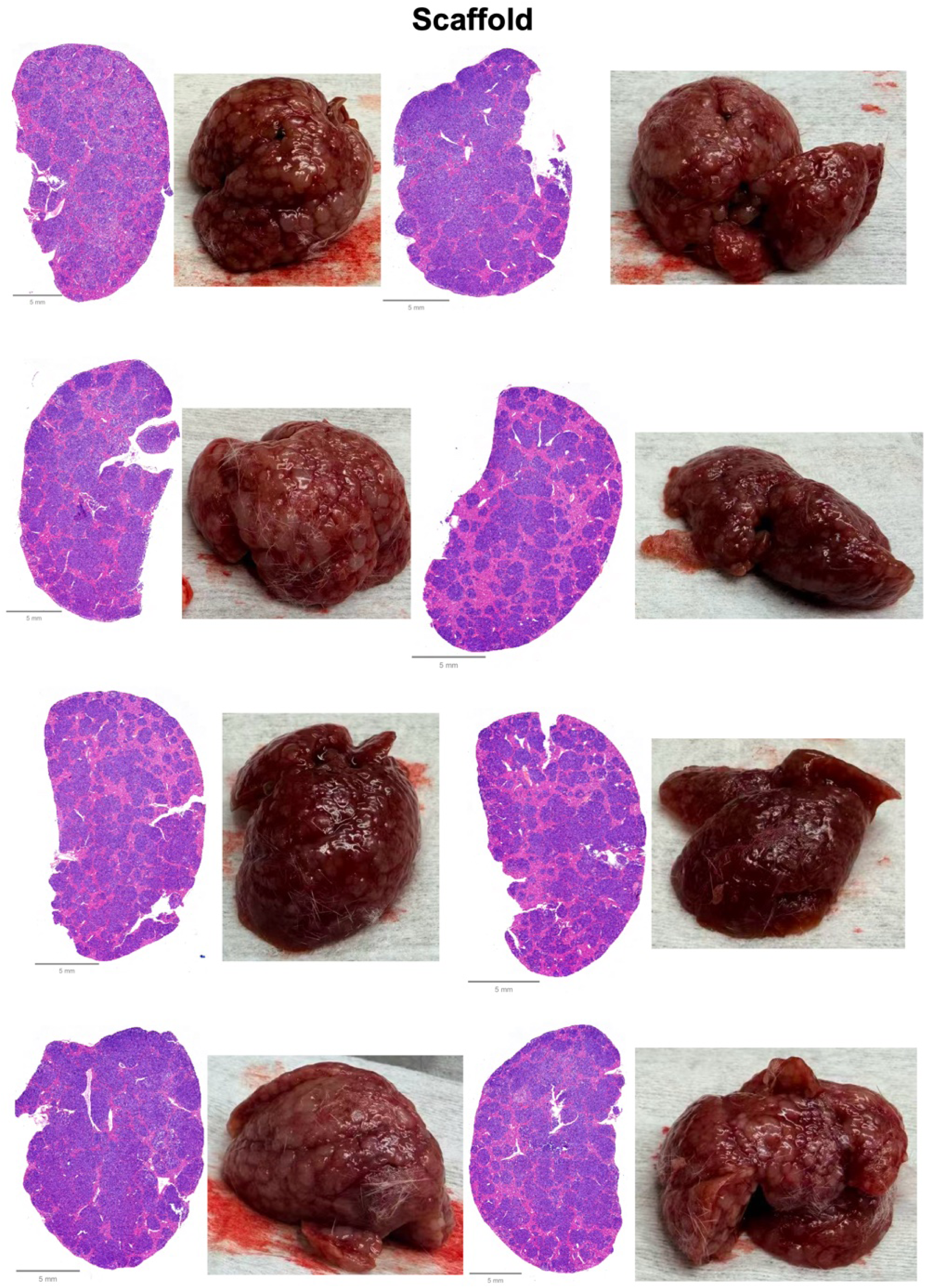
Representative liver histology and gross liver images from scaffold bacteria-treated mice. Representative whole-slide H&E-stained liver sections and corresponding gross liver images from H69 liver metastasis-bearing mice treated with scaffold bacteria. Scale bars: 5 mm.

**Supplemental Fig. 24.**
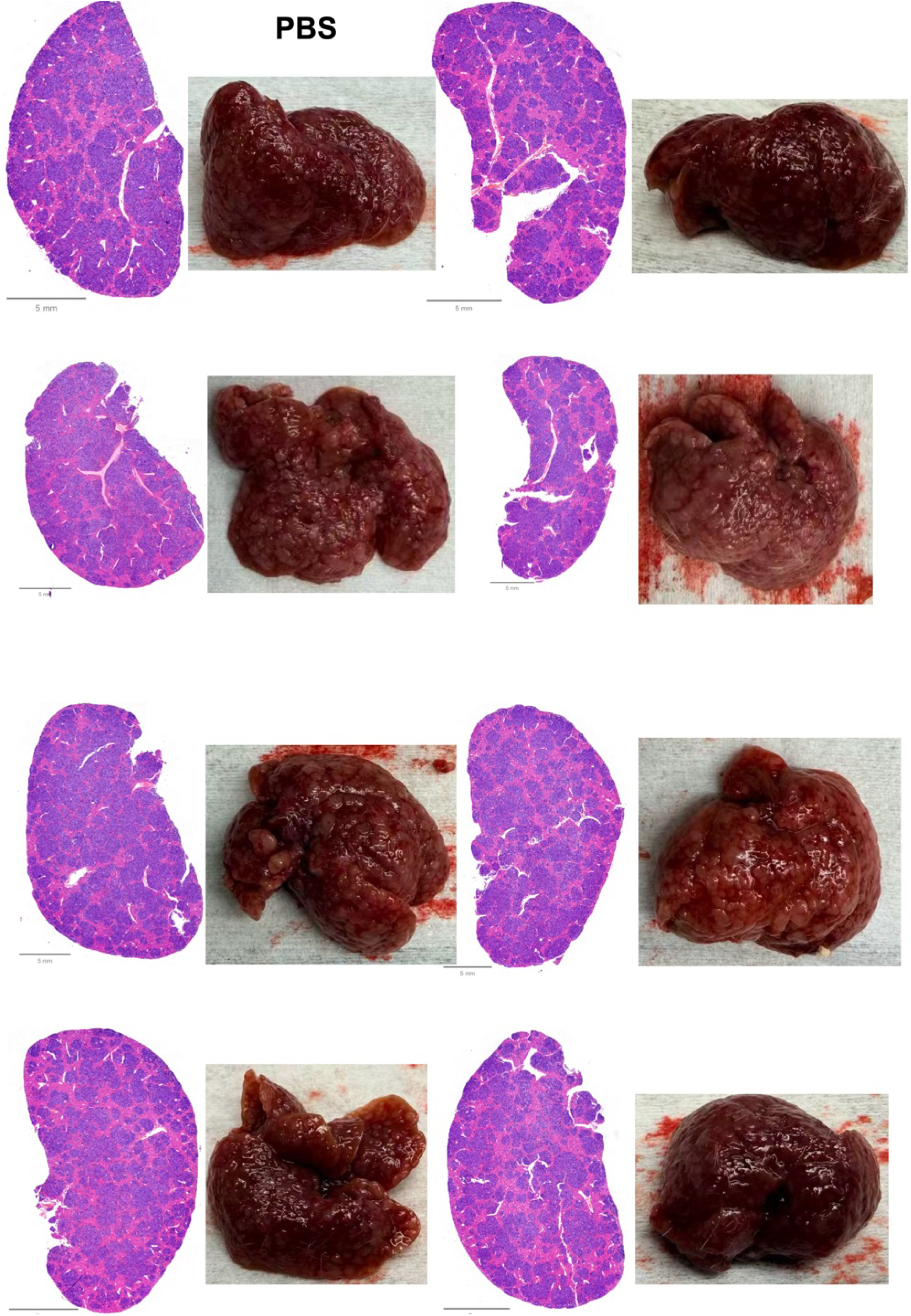
Representative liver histology and gross liver images from PBS-treated mice. Representative whole-slide H&E-stained liver sections and corresponding gross liver images from H69 liver metastasis-bearing mice treated with PBS. Scale bars: 5 mm.

**Supplemental Fig. 25.**
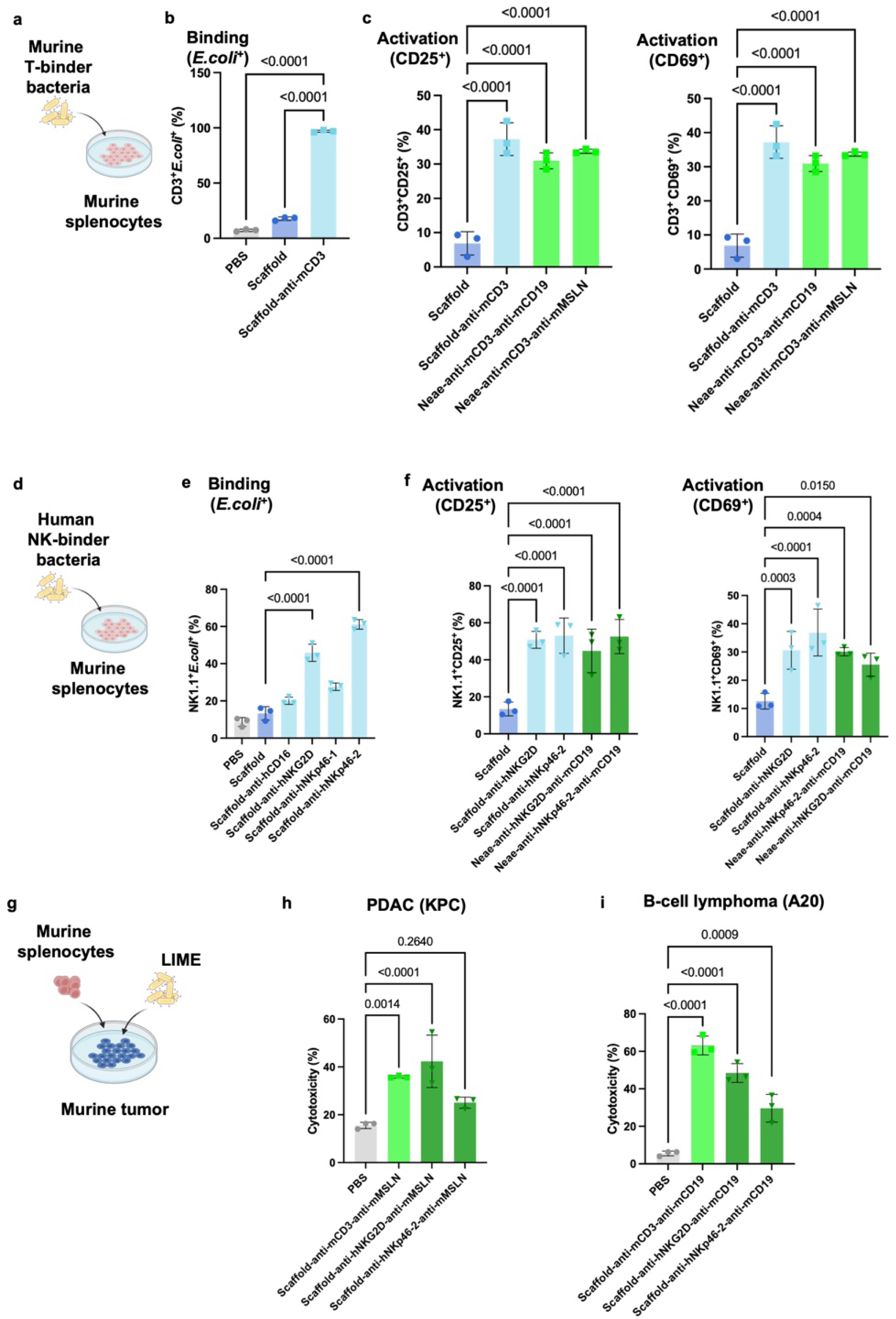
Evaluation of murine T-LIME and N-LIME. **a,** Schematic figure of the co-culture assay between *E. coli* K-12 displaying murine anti-CD3 scFv and murine splenocytes. **b,c,** Flow cytometry-based quantification of binding affinity (**b,** *E. coli*^+^) and activation (**c,** CD25 and CD69) of murine T cells. **d,** Schematic figure of co-culture assay between *E. coli* K-12 displaying scFvs binding to murine NK cell surface markers and murine splenocytes. **e,f,** Flow cytometry-based quantification of binding affinity (**e,** *E. coli*^+^) and activation (**f,** CD25 and CD69) of murine NK cells. **g,** Schematic figure of co-culture experiments involving murine cancer cell lines, splenocytes and leading LIME constructs. **h,i,** Flow cytometry-based assessment of cytotoxicity against KPC (**h**) and A20 (**i**) tumor cells. Statistical significance was determined using one-way ANOVA test with Tukey’s post hoc test (**b,c,e,f,h,i**). Data represent mean ± s.d.

**Supplemental Fig. 26.**
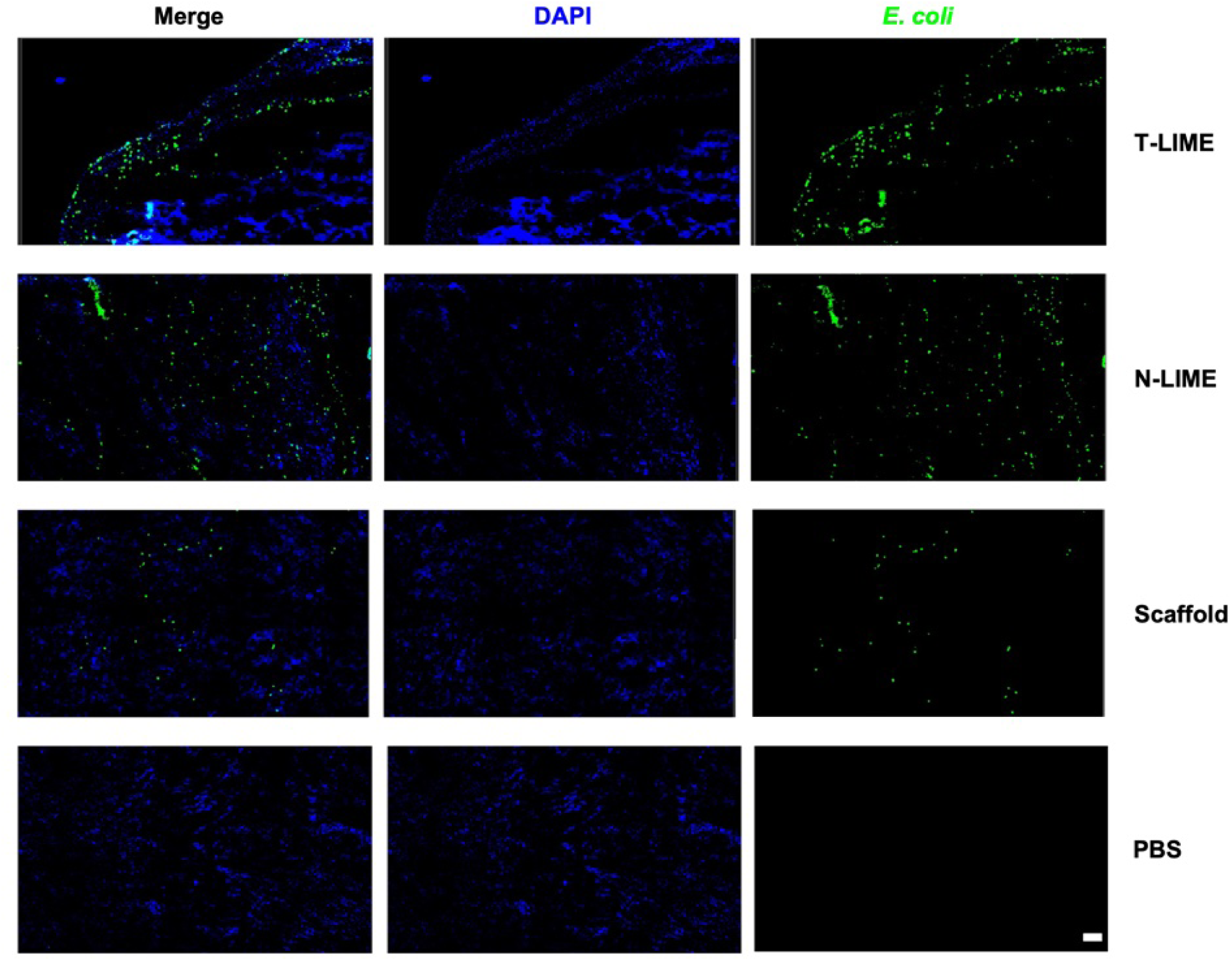
Intratumoral distribution of bacteria following LIME treatment in the A20 lymphoma model. Representative confocal images of tumor tissues collected from A20 tumor-bearing mice following treatment with CD19 T-LIME, CD19 N-LIME, Scaffold, or PBS. Tumor cryosections were stained with antibodies against *E. coli* (green) and DAPI (blue) with pseudocolor applied. Images were acquired using a Leica THUNDER Imager. Scale bar represents 100 μm.

**Supplemental Fig. 27.**
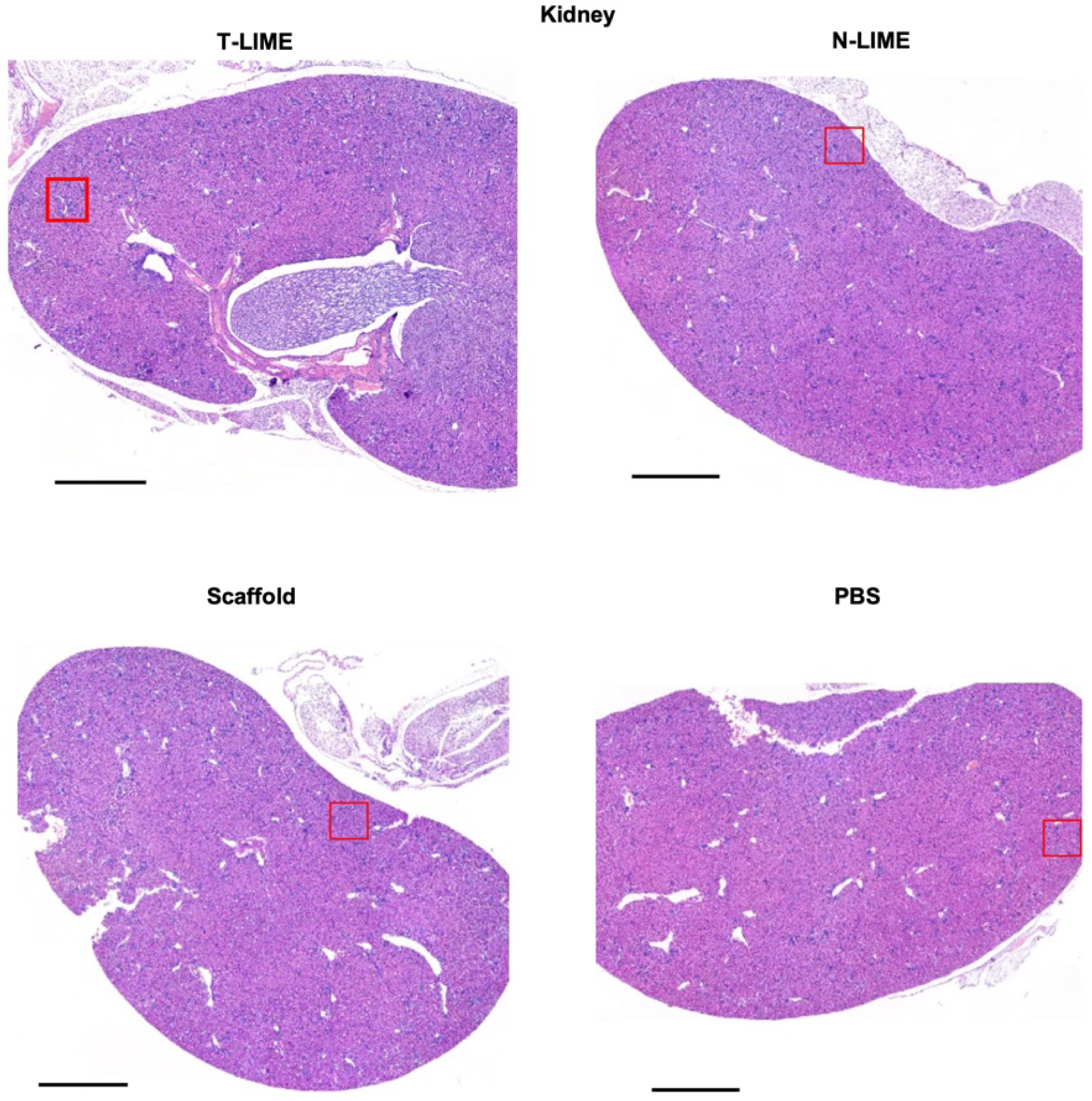
Impact of LIME treatment on kidney histology in the A20 lymphoma model. Representative hematoxylin and eosin (H&E)-stained sections of kidney from A20 tumor-bearing mice treated with CD19 T-LIME, CD19 N-LIME, Scaffold, or PBS. Scale bar represents 100 μm.

**Supplemental Fig. 28.**
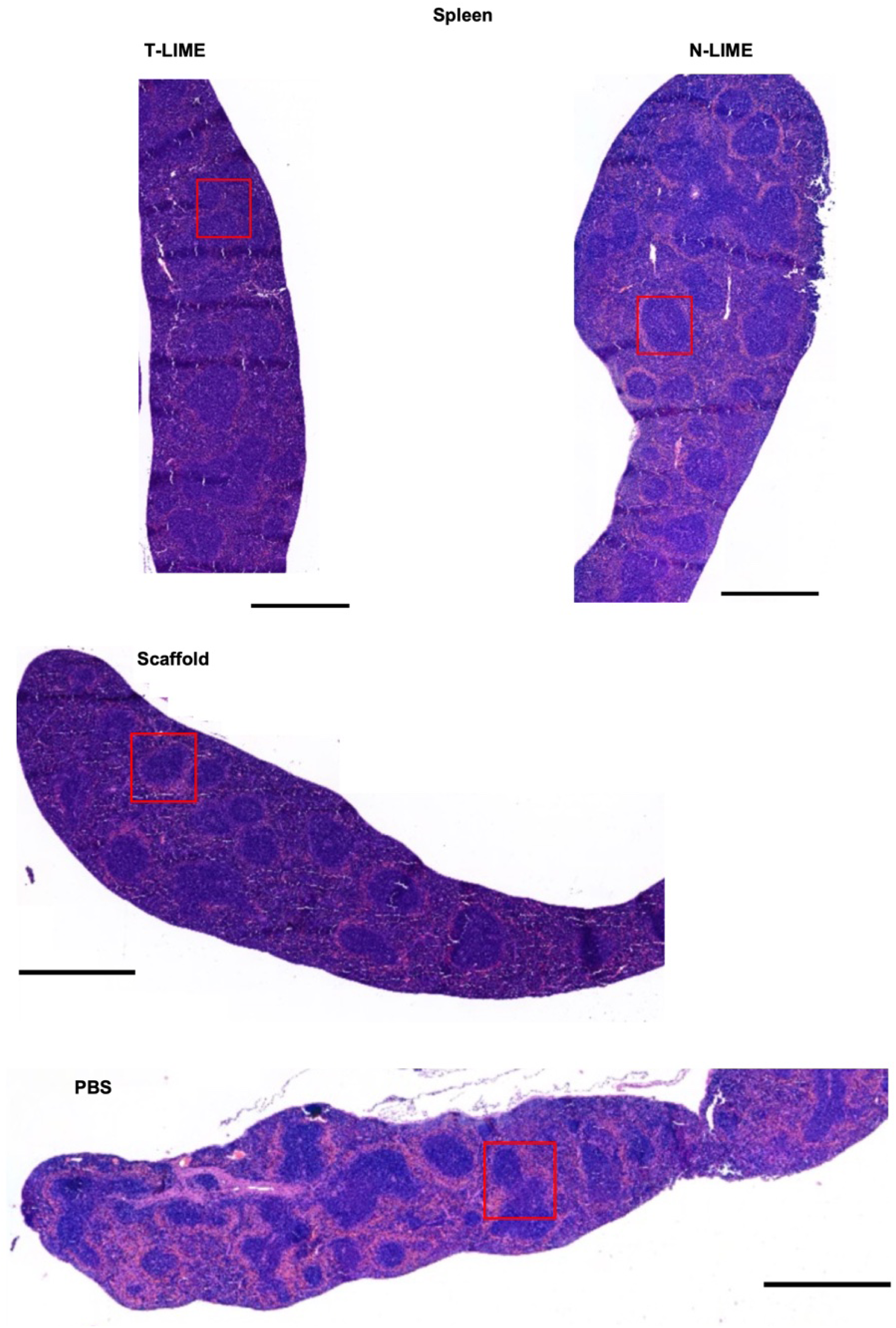
Impact of LIME treatment on spleen histology in the A20 lymphoma model. Representative hematoxylin and eosin (H&E)-stained sections of spleen from A20 tumor-bearing mice treated with CD19 T-LIME, CD19 N-LIME, Scaffold, or PBS. Scale bar represents 100 μm.

**Supplemental Fig. 29.**
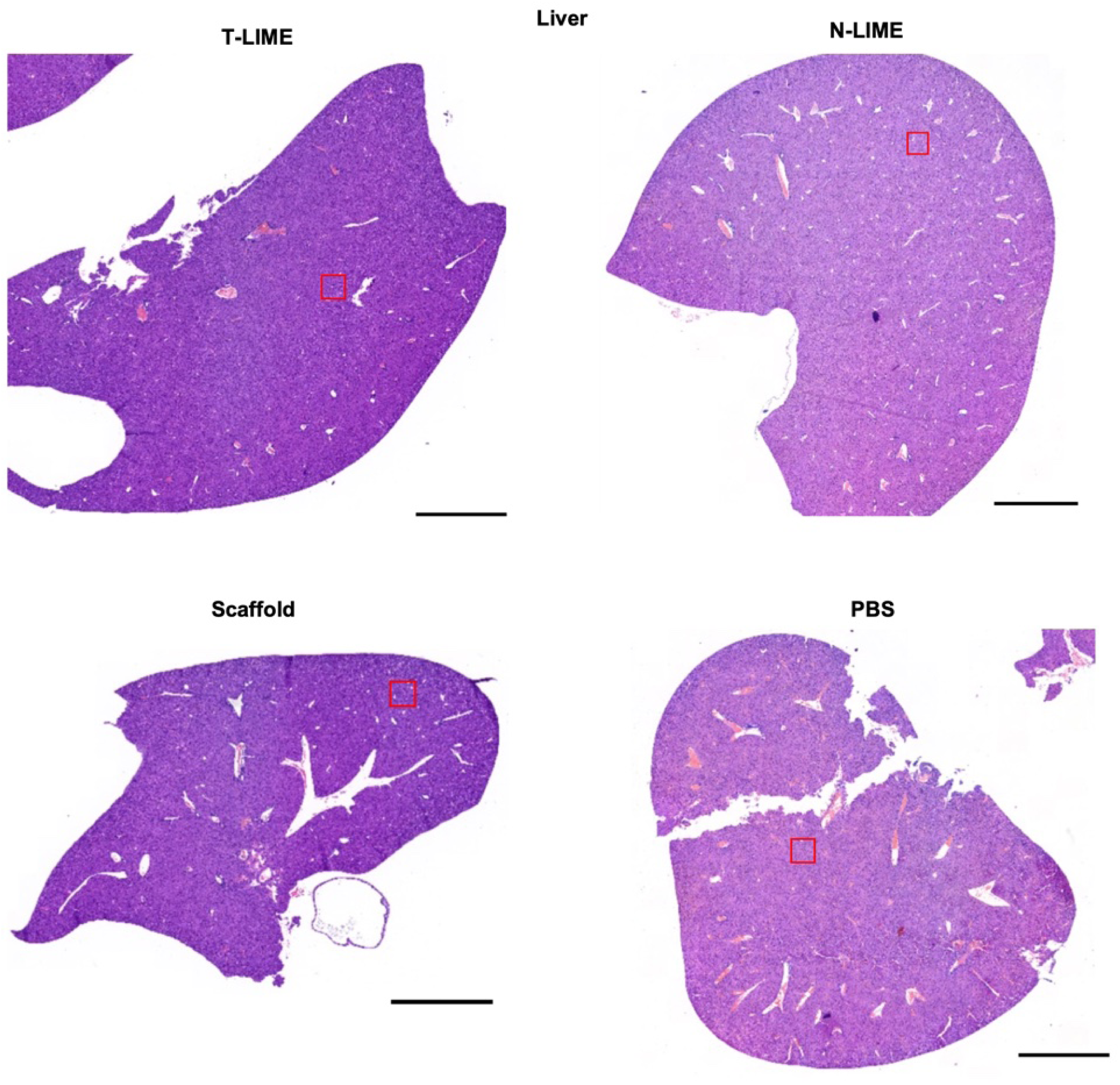
Impact of LIME treatment on liver histology in the A20 lymphoma model. Representative hematoxylin and eosin (H&E)-stained sections of liver from A20 tumor-bearing mice treated with CD19 T-LIME, CD19 N-LIME, Scaffold, or PBS. Scale bar represents 100 μm.

**Supplemental Fig. 30.**
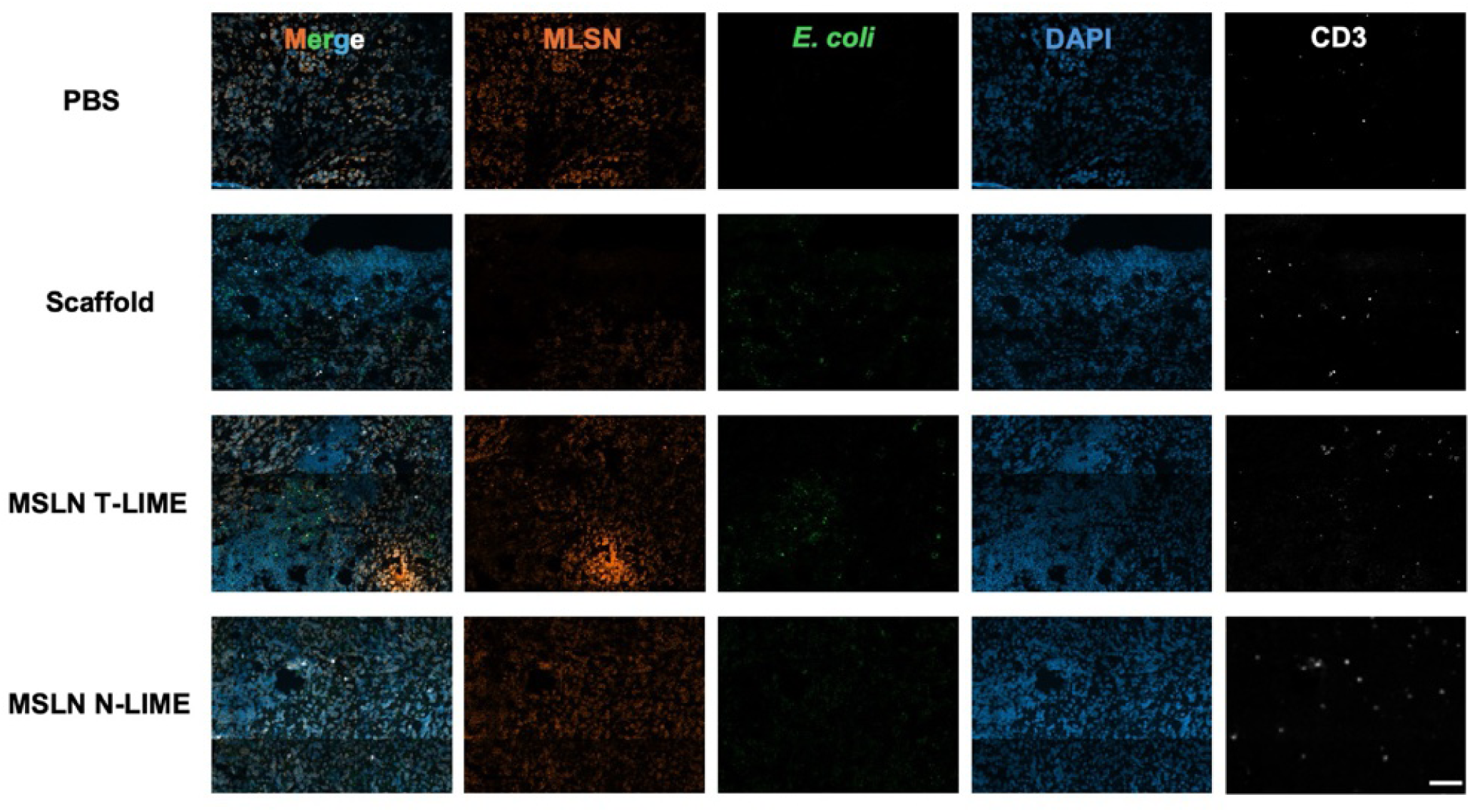
Intratumoral distribution of bacteria and T cells following LIME treatment. Representative confocal images of tumor tissue collected from KPC tumor-bearing mice on day 23 following treatment with PBS, Scaffold, MSLN T-LIME or MSLN N-LIME. Tumor cryosections were stained with antibodies for *E. coli* (green), CD3 (white), mesothelin (MSLN, orange), and DAPI (blue). Images were acquired using a ZEISS LSM 980 confocal microscope and displayed using pseudo color rendering. Scale bar represents 100μm.

**Supplemental Fig. 31.**
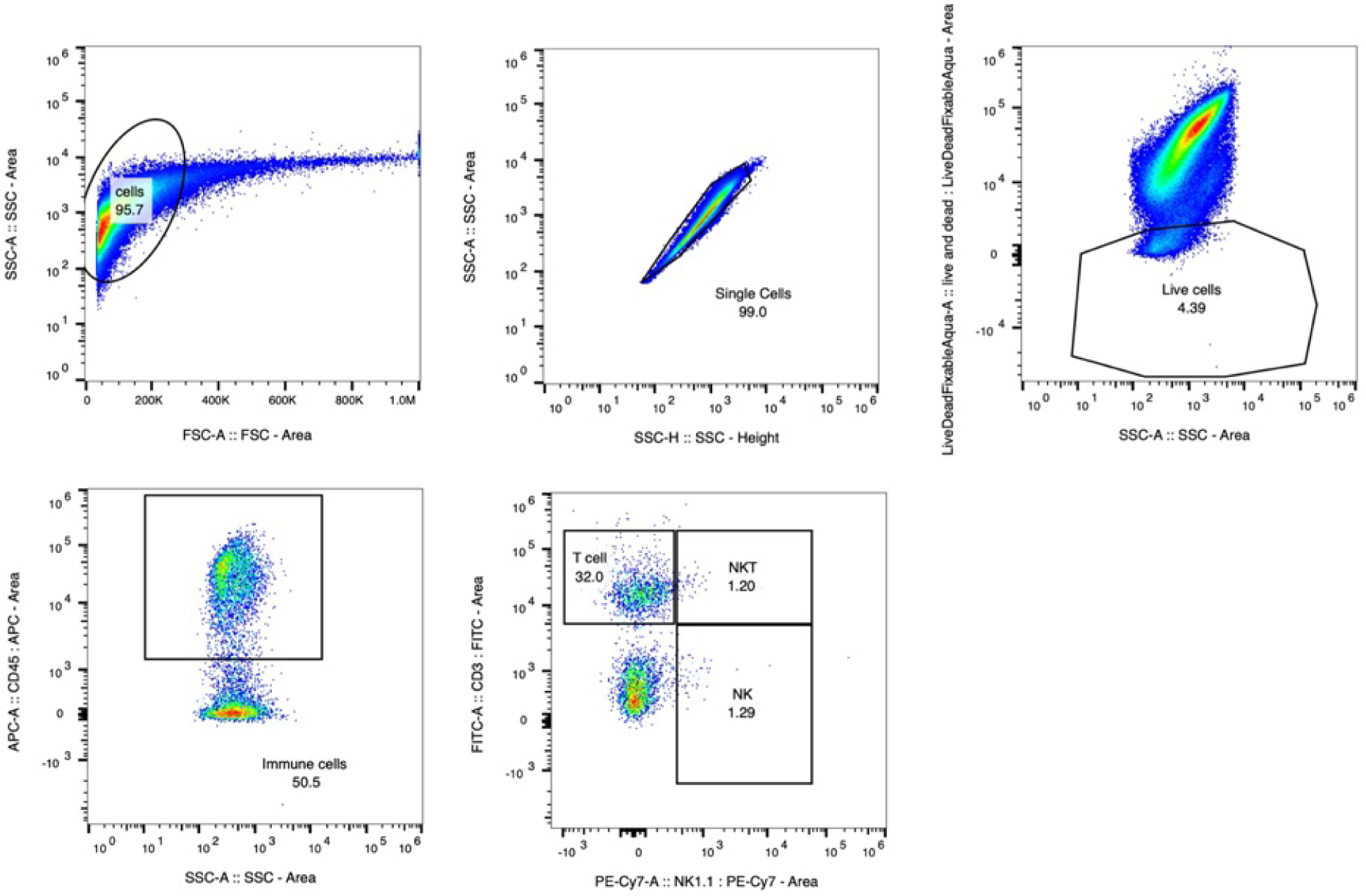
Characterization of NK and T cell populations in the tumor microenvironment (TME). Representative gating strategy used to identify NK and T cell populations in the TME.

**Extended Data Fig. 6.**
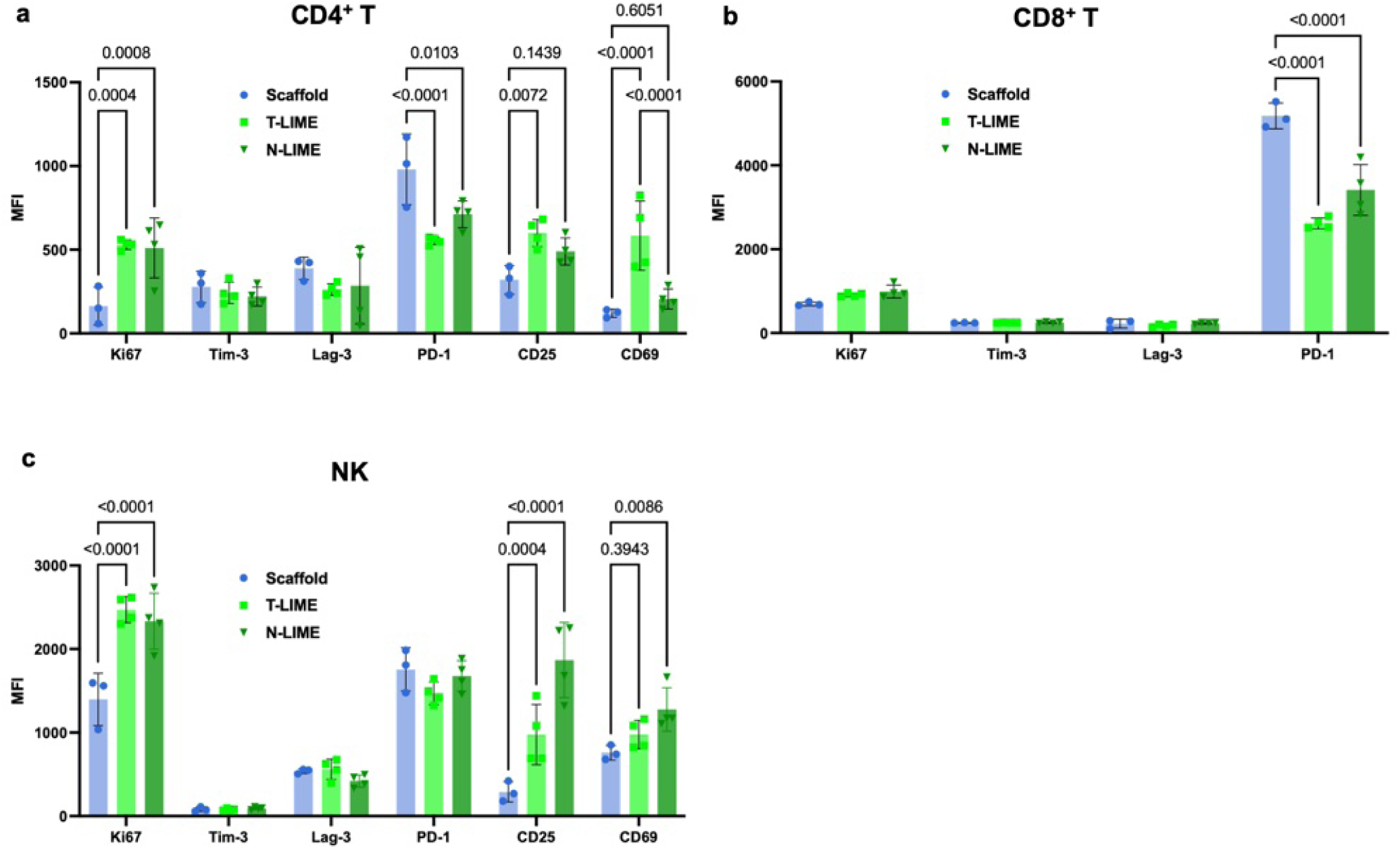
Phenotypic characterization of T and NK cells in the TME following LIME treatment. **a-c,** Expression of activation markers (CD25 and CD69), proliferation marker (Ki67) and inhibitory/exhaustion markers (Tim-3, Lag-3, and PD-1) in CD4^+^ T cells (**a**), CD8^+^ T cells (CD25 and CD69 shown in Figure 3h) (**b**), and NK cells (**c**). Statistical significance was determined using two-way ANOVA with Tukey’s post hoc test. Data represent median ± s.d. (**a-c**).

**Supplemental Fig. 32.**
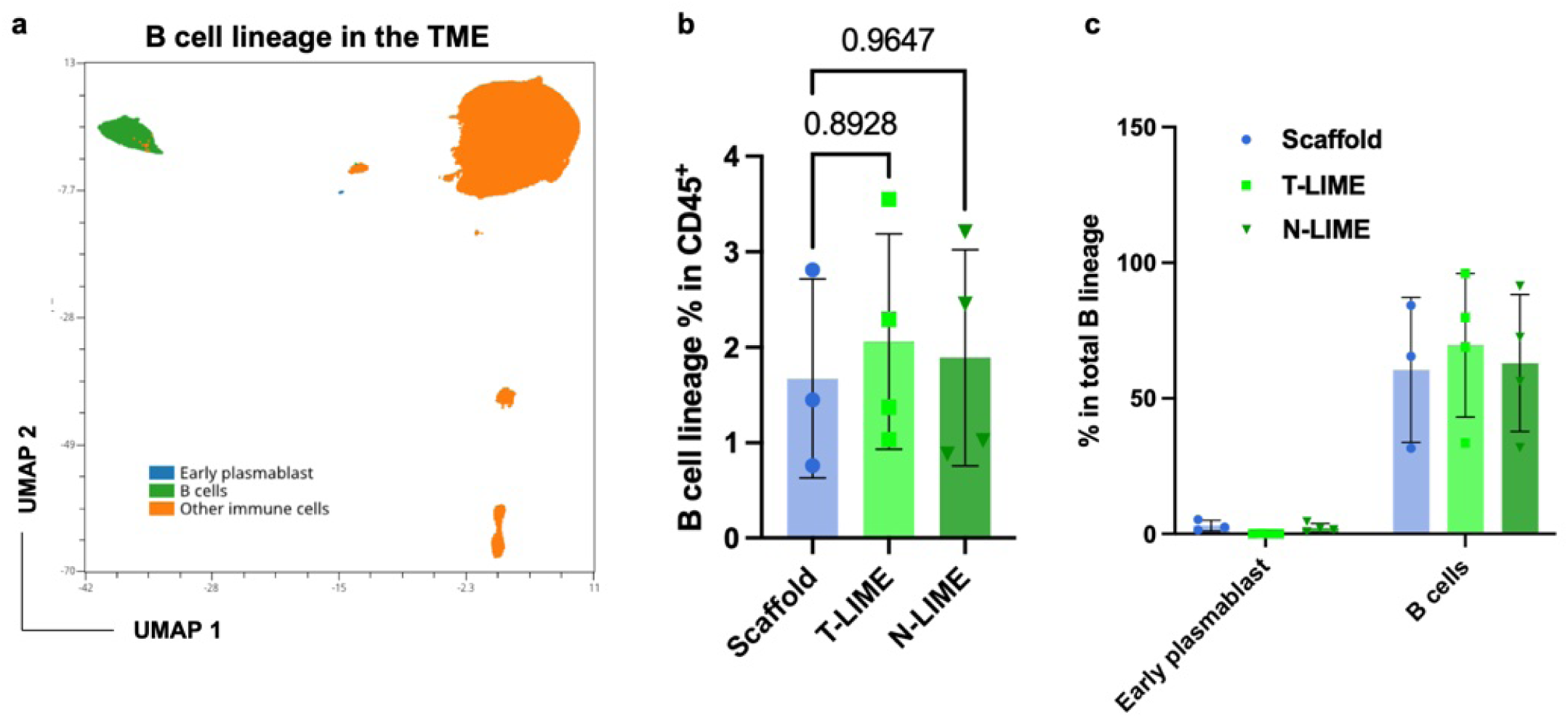
Impact of LIME treatment on B cell populations within the TME. **a,** UMAP visualization of B cell populations and other immune cell subsets within the TME. **b,** Frequences of B cell lineage populations (BCMA^+^, CXCR4^+^, B220^+^ and/or, CD19^+^) among CD45^+^ cells across treatment groups. **c,** Frequences of B cells (CD45^+^B220^+^CD19^+^) and early plasmablasts (CD45^+^BCMA^+^CXCR4^+^CD19^+^B220^-^) within the B cell lineage compartment. Statistical significance was determined using two-way ANOVA with Tukey’s post hoc test (**b**, **c**). Data represent mean ± s.d. (**b, c**).

**Supplemental Fig. 33.**
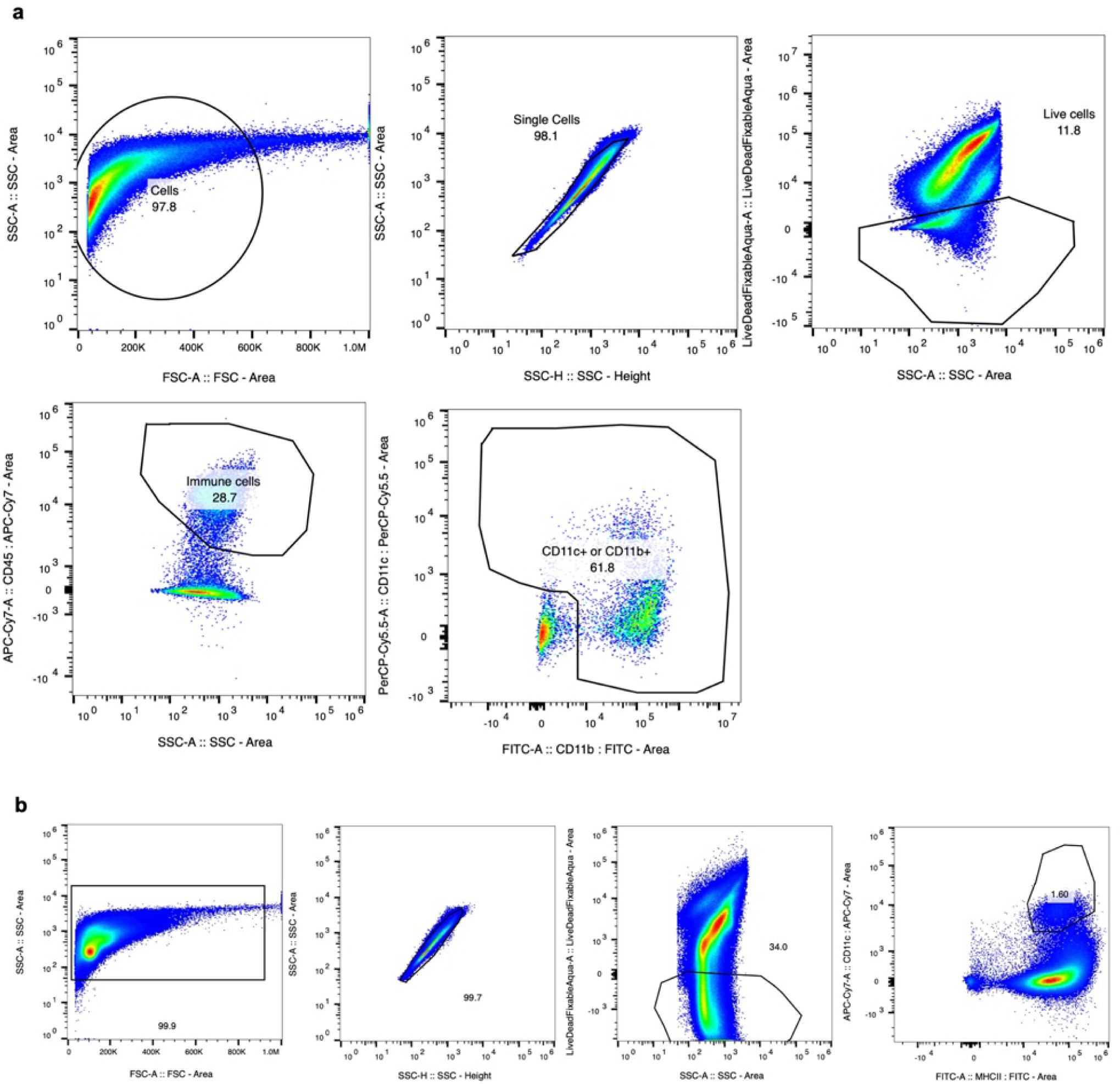
Impact of LIME treatment on myeloid cell populations in the TME. **a,** Representative flow cytometry plots illustrating the gating strategy used to identify myeloid cells in the TME. **b**, The representative gating strategy used to identify dendritic cell (DC) populations in the TME.

**Supplemental Fig. 34.**
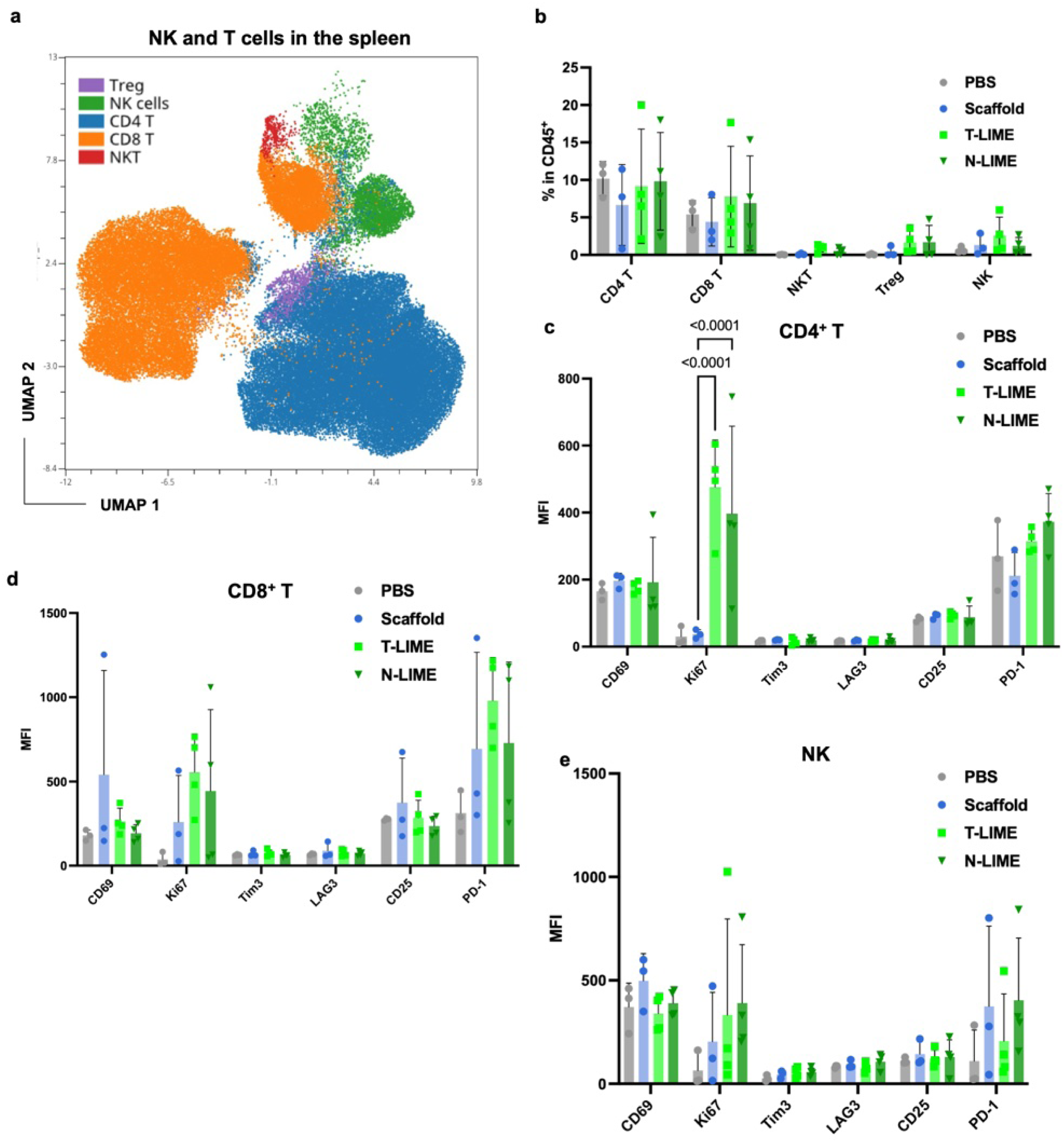
Characterization of NK and T cell populations in the spleen following LIME treatment. **a,** UMAP visualization of CD4^+^ T cells, CD8^+^ T cells, regulatory T (Treg) cells, NKT cells, and NK cells in the spleen. CD4^+^ T cells were defined as CD3^+^CD4^+^CD8^-^NK1.1^-^FoxP3^-^, CD8^+^ T cells as CD3^+^CD8^+^CD4^-^NK1.1^-^FoxP3^-^, Treg cells as CD3^+^CD8^-^CD4^+^NK1.1^-^FoxP3^+^. NK cells as CD4^-^CD8^-^NK1.1^+^, and NKT cells as CD3^+^CD4^-^CD8^-^NK1.1^+^. **b**, Frequencies of CD4^+^ T cells, CD8^+^ T cells, Treg cells, NKT cells, and NK cells among CD45^+^ cells in the spleen. **c-e,** Expression of activation markers (CD25 and CD69), proliferation marker (Ki67) and inhibitory/exhaustion markers (Tim-3, Lag-3, and PD-1) in CD4^+^ T cells (**c**), in CD8^+^ T cells (**d**), and NK cells (**e**). Statistical significance was determined using two-way ANOVA with Tukey’s post hoc test (**c**). Data represent mean ± s.d. (**b, c, d, e).**

**Supplemental Fig. 35.**
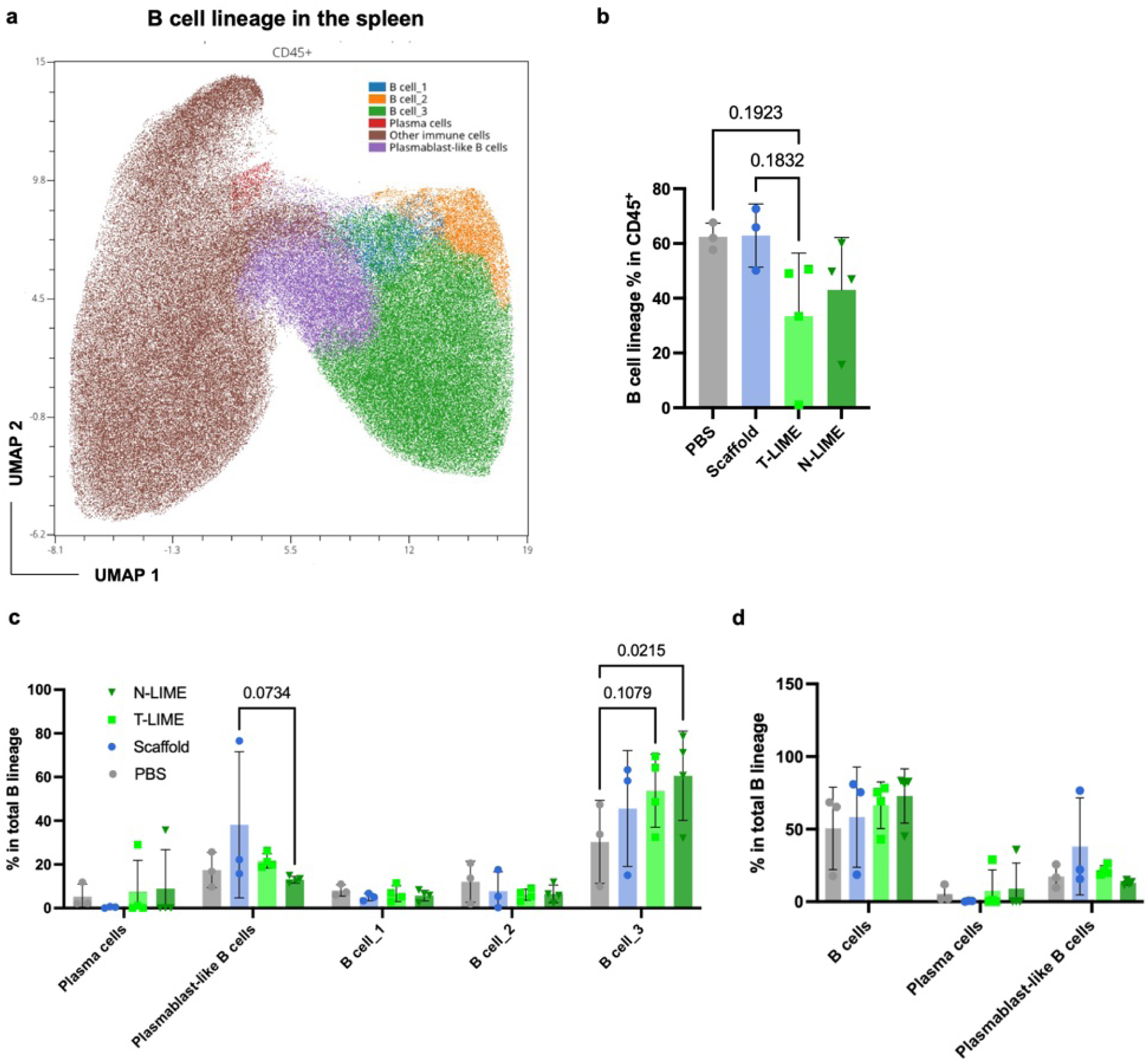
Impact of LIME treatment on B cell lineage populations in the spleen. **a**, UMAP visualization of B cell clusters, plasmablast-like B cells, plasma cells, and other immune cell populations in the spleen. Plasma cells were defined as BCMA^+^CXCR4^+^B220^-^CD19^-^IgM^-^IgD^-^, plasmablast-like B cells as BCMA^-^CXCR4^-^B220^-/low^CD19^-/low^IgM^-^IgD^-^, B cells_1 as BCMA^-^CXCR4^-^B220^+^CD19^+^IgM^+^IgD^-^, B cells_2 as BCMA^-^CXCR4^-^B220^+^CD19^+^IgM^+^IgD^+^, B cells_3 as BCMA^-^CXCR4^-^B220^+^CD19^+^IgM^-^IgD^+^, and other immune cells as BCMA^-^CXCR4^-^B220^-^CD19^-^IgM^-^IgD^-^. **b,** Frequencies of total B cells among CD45^+^ cells in the spleen. **c**, **d**, Frequences of distinct B cell lineage cell types (**c**) and clusters (**d**) within the B cell compartment in the spleen. Statistical significance was determined using two-way ANOVA with Tukey’s post hoc test (**b-d**). Data represent mean ± s.d. (**b**-**d**).

**Supplemental Fig. 36.**
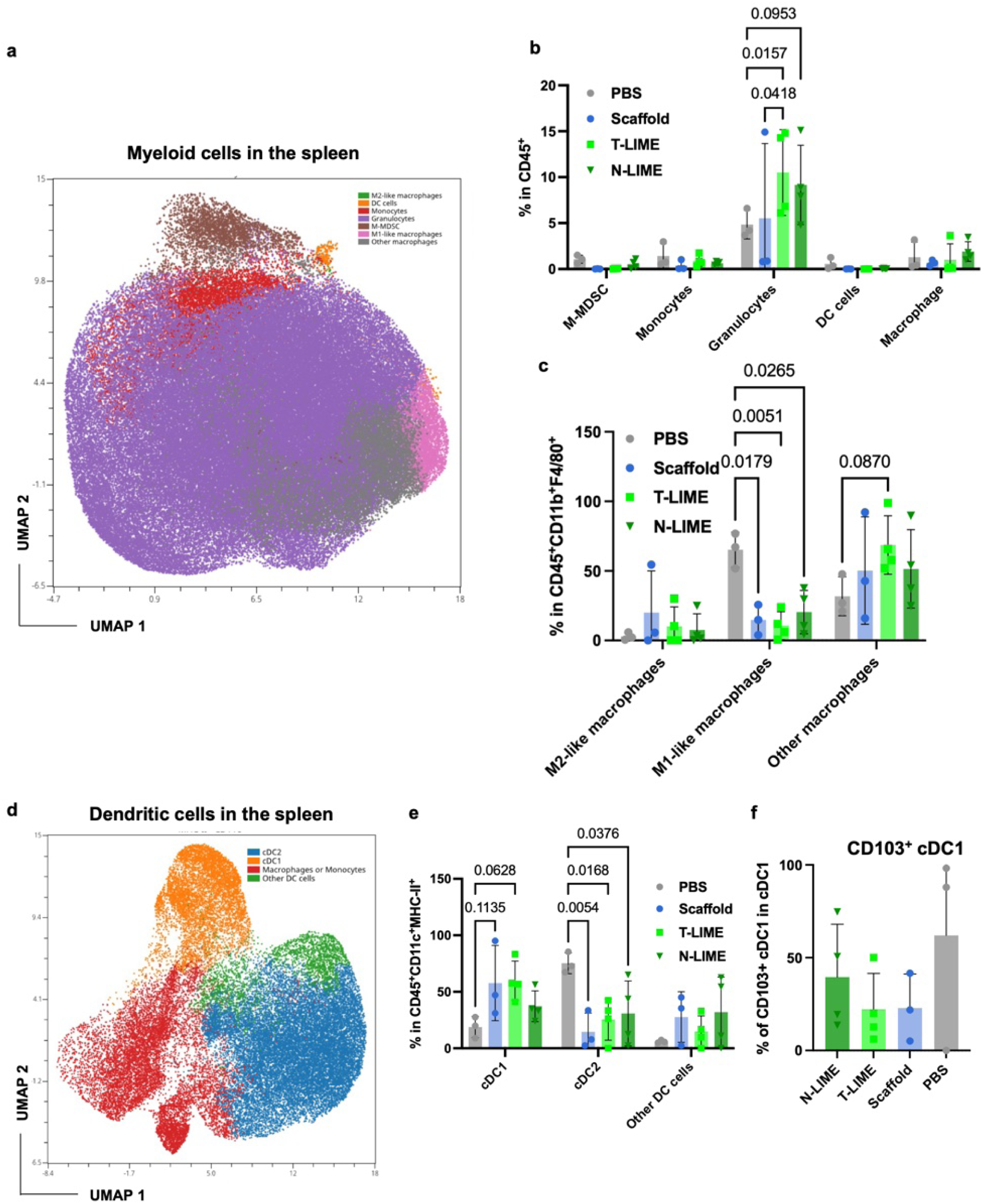
Impact of LIME treatment on myeloid cell populations in the spleen. **a,** UMAP visualization of myeloid cell populations in the spleen, including monocytes, granulocytes, dendritic cells, monocytic myeloid-derived suppressor cells (M-MDSC), M1-like macrophages, M2-like macrophages and other macrophages. M-MDSCs were defined as CD11b^+^F4/80^-^Ly6C^+^Ly6G^-^, monocytes as CD11b^+^F4/80^-^Ly6C^med^Ly6G^-^, granulocytes as CD11b^+^F4/80^-^Ly6G^+^Ly6C^+^, DCs as CD11c^+^MHC-II^+^F4/80^-^, M1-like macrophages as CD11b^+^F4/80^+^CD206^-^MHC-II^+^, M2-like macrophages as CD11b^+^F4/80^+^CD206^+^MHC-II^-^, and other macrophages as CD11b^+^F4/80^+^CD206^-^MHC-II^-^. **b**, Frequences of myeloid cell populations among CD45^+^ cells in the spleen. **c**, Frequences of macrophages subsets among total macrophages (CD11b^+^F4/80^+^) in the spleen. **d**, UMAP visualization of monocytes/macrophage and dendritic cells in the spleen. Conventional Dendritic cell 1 (cDC1) were defined as CD11c^+^MHC-II^+^CD64^-^CD24^+^XCR1^+^CD11b^-^, cDC2 as CD11c^+^MHC-II^+^CD64^-^CD24^-^XCR1^-^CD11b^+^, other DC cells as CD11c^+^MHC-II^+^CD64^-^CD24^-^XCR1^-^CD11b^-^, and macrophages/monocytes as CD11c^+^MHC-II^+^CD64^+^. **e**, Frequencies of key DC subsets among total DCs (CD11c^+^MHC-II^+^CD64^-^) in the spleen. **f**, Frequences of migratory cDC1 (CD11c^+^MHC-II^+^CD64^-^CD24^+^XCR1^+^CD11b^-^CD103^+^) among total cDC1 cells (CD11c^+^MHC-II^+^CD64^-^CD24^+^XCR1^+^CD11b^-^). Statistical significance was determined using two-way ANOVA with Tukey’s post hoc test (**b**, **c**, **e**). Data represent mean ± s.d. (**b, c, e**, **f**).

**Supplemental Fig. 37.**
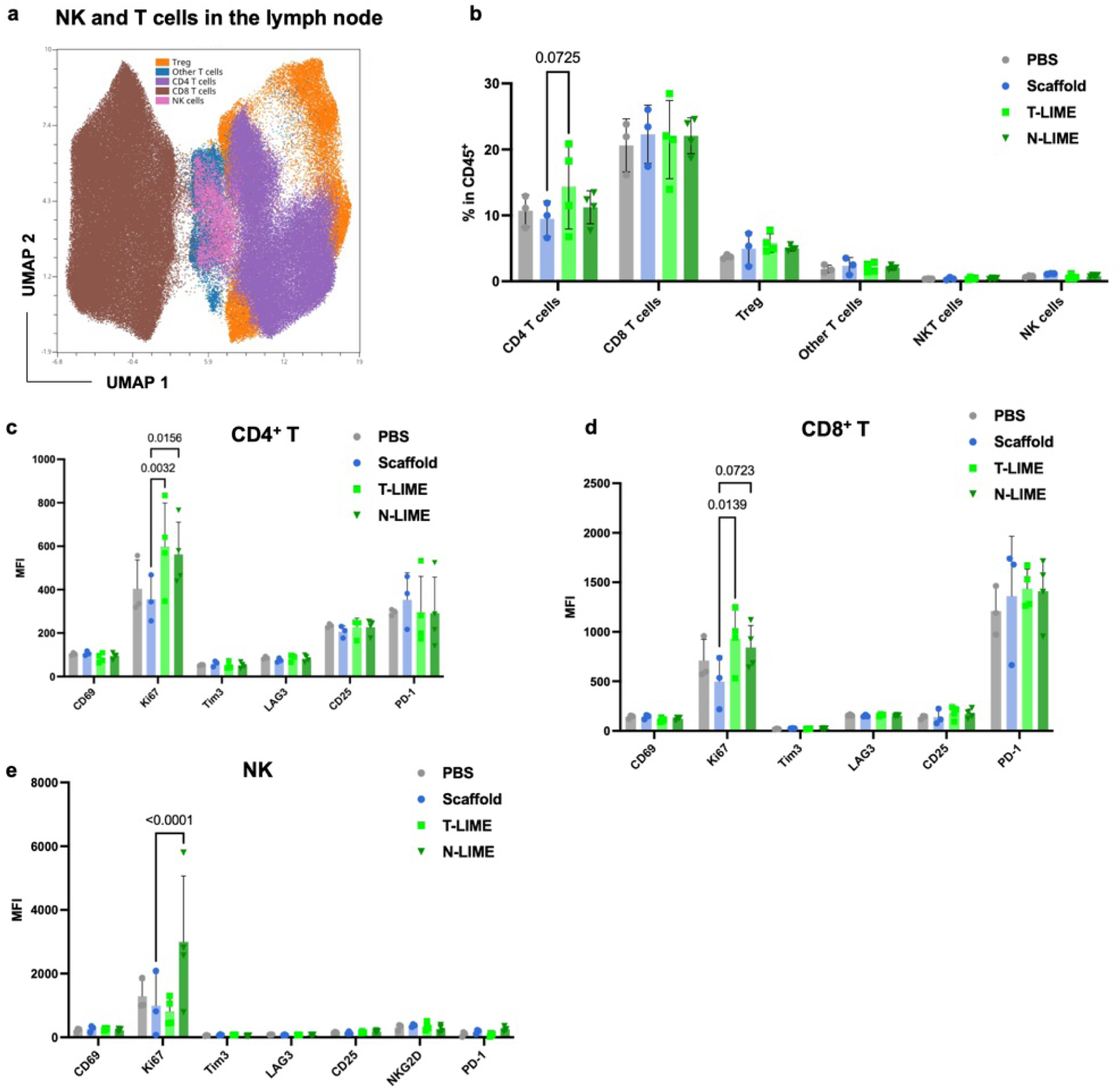
Impact of Lime treatment on NK and T cell populations in the tumor draining lymph nodes (TDLN). **a,** UMAP visualization of CD4^+^ T cells, CD8^+^ T cells, Treg cells, NKT cells, NK cells and other T cells in the TDLN. CD4^+^ T cells were defined as CD3^+^CD4^+^CD8^-^NK1.1^-^FoxP3^-^, CD8^+^ T cells as CD3^+^CD8^+^CD4^-^NK1.1^-^FoxP3^-^, Treg cells as CD3^+^CD8^-^CD4^+^NK1.1^-^FoxP3^+^CD25^+^CD127^-^, NK cells as CD3^-^CD4^-^CD8^-^NK1.1^+^, NKT cells as CD3^+^CD4^-^CD8^-^NK1.1^+^, and other T cells: CD3^+^CD4^-^CD8^-^NK1.1^-^. **b**, Frequences of CD4^+^ T, CD8^+^ T, Treg, NKT, NK and other T cells among CD45^+^ cells in the TDLN. **c-e,** Expression of activation (CD25 and CD69), proliferation (Ki67) and inhibitory/exhaustion (Tim-3, Lag-3, and PD-1) markers in CD4^+^ T (**c**), CD8^+^ T (**d**), and NK cells (**e**). Statistical significance was determined using two-way ANOVA with Tukey’s post hoc test. Data represent mean ± s.d. (**b-e).**

**Supplemental Fig. 38.**
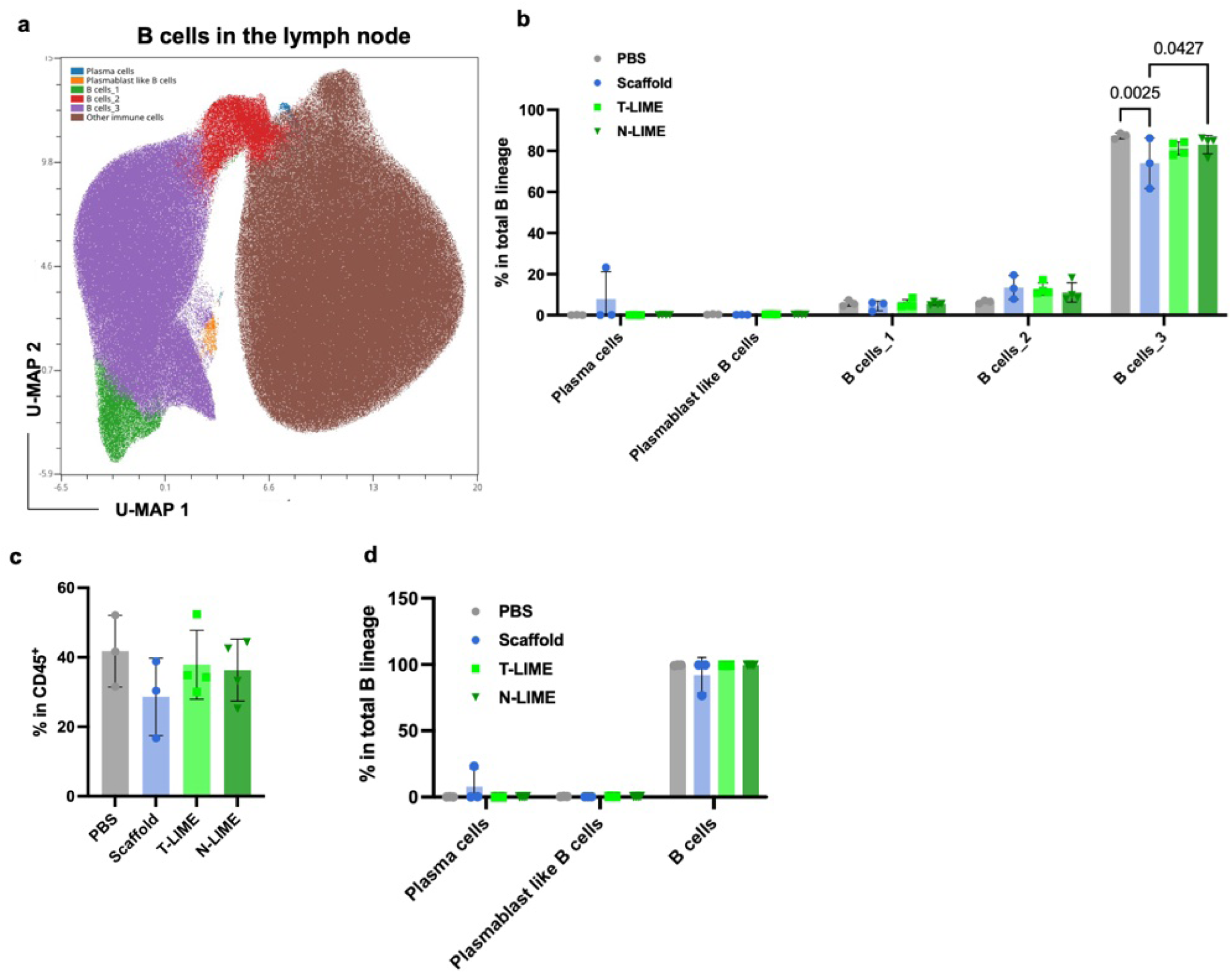
Impact of LIME treatment on B cell populations in the tumor draining lymph nodes (TDLN). **a**, UMAP visualization of B cell clusters, plasmablast-like B cells, plasma cells, and other immune cell populations in the TDLN. Plasma cells were defined as BCMA^+^CXCR4^+^B220^-^CD19^-^IgM^-^IgD^-^, plasmablast-like B cells as BCMA^+^CXCR4^+^B220^+^CD19^+^IgM^-^IgD^+^, B cells_1 as BCMA^-^CXCR4^-^B220^+^CD19^+^IgM^+^IgD^+^, B cells_2 as BCMA^-^CXCR4^-^B220^+^CD19^+^IgM^-^IgD^-^, B cells_3 as BCMA^-^CXCR4^-^B220^+^CD19^+^IgM^-^IgD^+^, and other immune cells as BCMA^-^ CXCR4^-^B220^-^CD19^-^IgM^-^IgD^-^. **b,** Frequences of total B cell lineage populations among CD45^+^ cells in the TDLN. **c**, **d**, Frequences of individual B cell clusters (**c**) and subsets (**d**) within the B cell lineage compartment in the TDLN. Statistical significance was determined using two-way ANOVA with Tukey’s post hoc test (**b**). Data represent mean ± s.d. (**b**-**d**).

**Supplemental Fig. 39.**
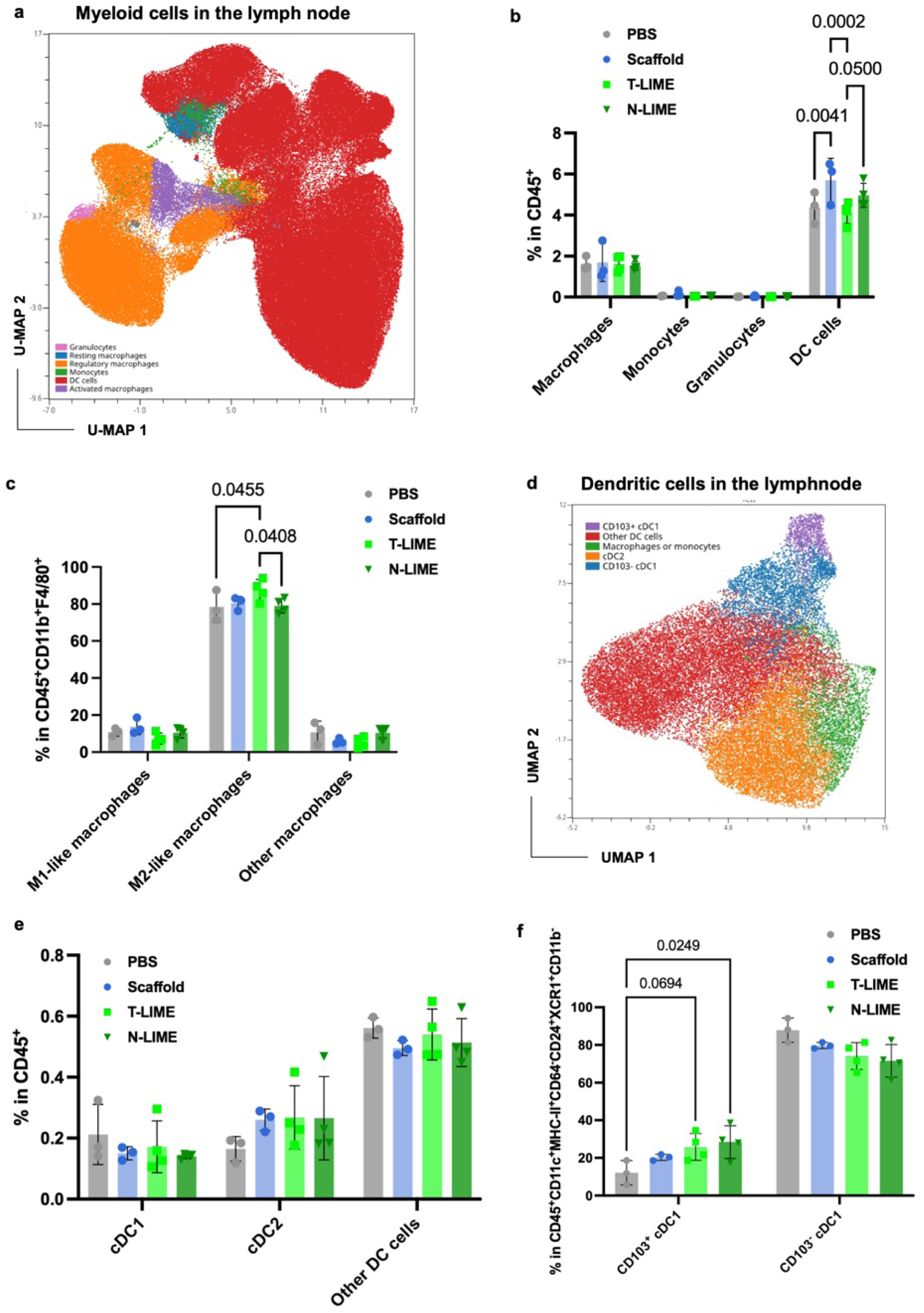
Characterization of myeloid cell populations in the tumor draining lymph nodes (TDLN) following LIME treatment. **a,** UMAP visualization of key myeloid cell populations (monocytes, granulocytes, dendritic cells, M1-like macrophages, M2-like macrophages and other macrophages subsets) in the TDLN. Monocytes were defined as CD11b^+^F4/80^-^Ly6C^+^Ly6G^-^, Granulocytes as CD11b^+^F4/80^-^Ly6G^+^Ly6C^+^, DC cells as CD11c^+^MHC-II^+^F4/80^-^, M1-like macrophages as CD11b^+^F4/80^+^CD206^-^MHC-II^+^, M2-like macrophages as CD11b^+^F4/80^+^CD206^+^MHC-II^-^, and other macrophages as CD11b^+^F4/80^+^CD206^+^MHC-II^+^. **b**, Frequences of total myeloid cell populations among CD45^+^ cells in the TDLN. **c**, Frequences of distinct macrophages subsets among total macrophages (CD11b^+^F4/80^+^) in the TDLN. **d**, UMAP visualization of selected myeloid cell populations in the TDLN including monocyte/macrophage and dendritic cells. CD103^+^ cDC1 cells were defined as CD11c^+^MHC-II^+^CD64^-^CD24^+^XCR1^+^CD11b^-^CD103^+^, CD103^-^ cDC1 cells as CD11c^+^MHC-II^+^CD64^-^CD24^+^XCR1^+^CD11b^-^ CD103^-^, cDC2 cells as CD11c^+^MHC-II^+^CD64^-^CD24^-^XCR1^-^CD11b^+^, other DC cells as CD11c^+^MHC-II^+^CD64^-^CD24^-^XCR1^-^CD11b^-^, and macrophages /monocytes as CD11c^+^MHC-II^+^CD64^+^. **e**, Frequencies of different DC subsets among total DCs (CD11c^+^MHC-II^+^CD64^-^) in the TDLN. **d**, Frequencies of CD103^+^ cDC1 or CD103^-^ cDC1 subsets among total cDC1 cells (CD11c^+^MHC-II^+^CD64^-^CD24^+^XCR1^+^CD11b^-^). Statistical significance was determined using two-way ANOVA with Tukey’s post hoc test (**b, c, f**). Data represent mean ± s.d. (**b, c, e, f**).

**Supplemental Fig. 40.**
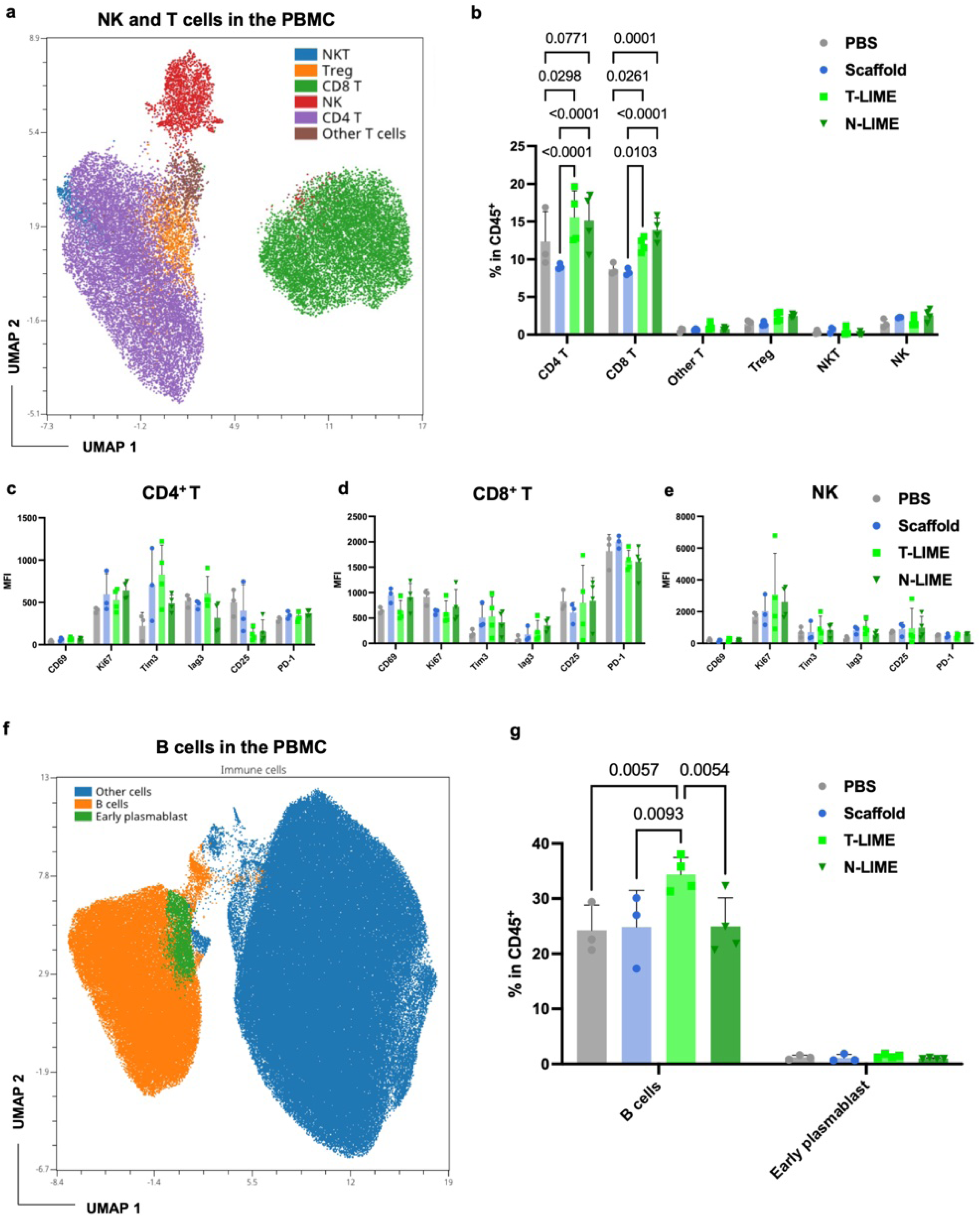
Impact of LIME treatment on peripheral blood lymphocyte populations. **a,** UMAP visualization of CD4^+^ T cells, CD8^+^ T cells, Treg cells, NKT cells, NK cells and other T cell populations within peripheral blood mononuclear cells (PBMCs). CD4^+^ T cells were defined as CD3^+^CD4^+^CD8^-^NK1.1^-^FoxP3^-^, CD8^+^ T cells as CD3^+^CD8^+^CD4^-^NK1.1^-^FoxP3^-^, Treg cells as CD3^+^CD8^-^CD4^+^NK1.1^-^FoxP3^+^CD25^+^CD127^-^, NK cells as CD3^-^CD4^-^CD8^-^NK1.1^+^, NKT cells as CD3^+^CD4^-^CD8^-^NK1.1^+^, and other T cells as CD3^+^CD4^-^CD8^-^NK1.1^-^. **b**, Frequences of CD4^+^ T cells, CD8^+^ T cells, Treg cells, NKT cells, NK cells and other T cell populations among CD45^+^ in PBMCs. **c-e,** Expression of activation markers (CD25 and CD69), proliferation marker (Ki67) and inhibitory/exhaustion markers (Tim-3, Lag-3, and PD-1) in CD4^+^ T cells (**c**), in CD8^+^ T cells (**d**), and in NK cells (**e**). **f**, UMAP visualization of B cells, early-plasmablast, and other immune cell populations within PBMCs. Early plasmablasts were defined as BCMA^-^CXCR4^+^B220^low^CD19^+^, B cells as BCMA^-^CXCR4^-^B220^+^CD19^+^IgM^-^IgD^+^, and other immune cells as BCMA^-^CXCR4^-^B220^-^CD19^-^IgM^-^IgD^-^. **g,** Frequences of B cells among CD45^+^ cells in PBMCs. Statistical significance was determined using two-way ANOVA with Tukey’s post hoc test (**b, g**). Data represent mean ± s.d. (**b**-**e, g**).

**Supplemental Fig. 41.**
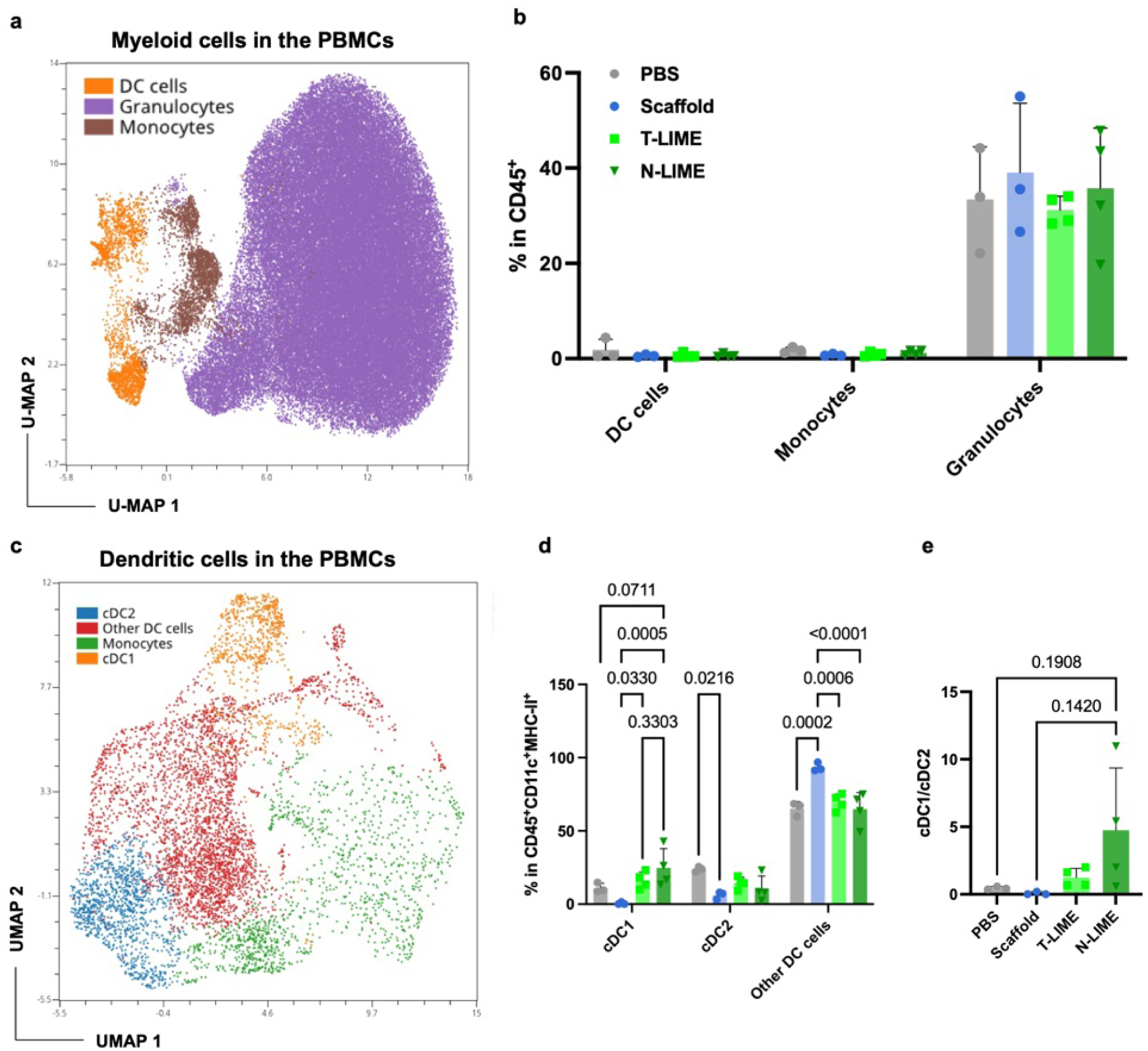
Impact of LIME treatment on peripheral blood myeloid cells. **a,** UMAP visualization of myeloid cell populations (monocytes, granulocytes and dendritic cells) in PBMCs. Monocytes were defined as CD11b^+^F4/80^-^ Ly6C^+^Ly6G^-^, granulocytes as CD11b^+^F4/80^-^Ly6G^+^Ly6C^+^, and DC cells as CD11c^+^MHC-II^+^F4/80^-^. **b**, Frequences of myeloid cells among CD45^+^ cells in PBMCs. **c**, UMAP visualization of selected myeloid cell populations, including monocytes and DC subsets within PBMCs. cDC1 were defined as CD11c^+^MHC-II^+^CD64^-^CD24^+^XCR1^+^CD11b^-^CD103^+^, cDC2 cells as CD11c^+^MHC-II^+^CD64^-^CD24^-^XCR1^-^CD11b^+^, other DC cells as CD11c^+^MHC-II^+^CD64^-^CD24^-^XCR1^-^CD11b^-^, and monocytes as CD11c^+^MHC-II^+^CD64^+^. **d**, Frequences of distinct DC subsets among total DCs (CD11c^+^MHC-II^+^CD64^-^) in PBMCs. **e**, Ratio of cDC1 to cCD2 populations in PBMCs. Statistical significance was determined using two-way ANOVA with Tukey’s post hoc test (**d**, **e**). Data represent mean ± s.d. (**b**, **d**, **e**).

**Supplemental Fig. 42.**
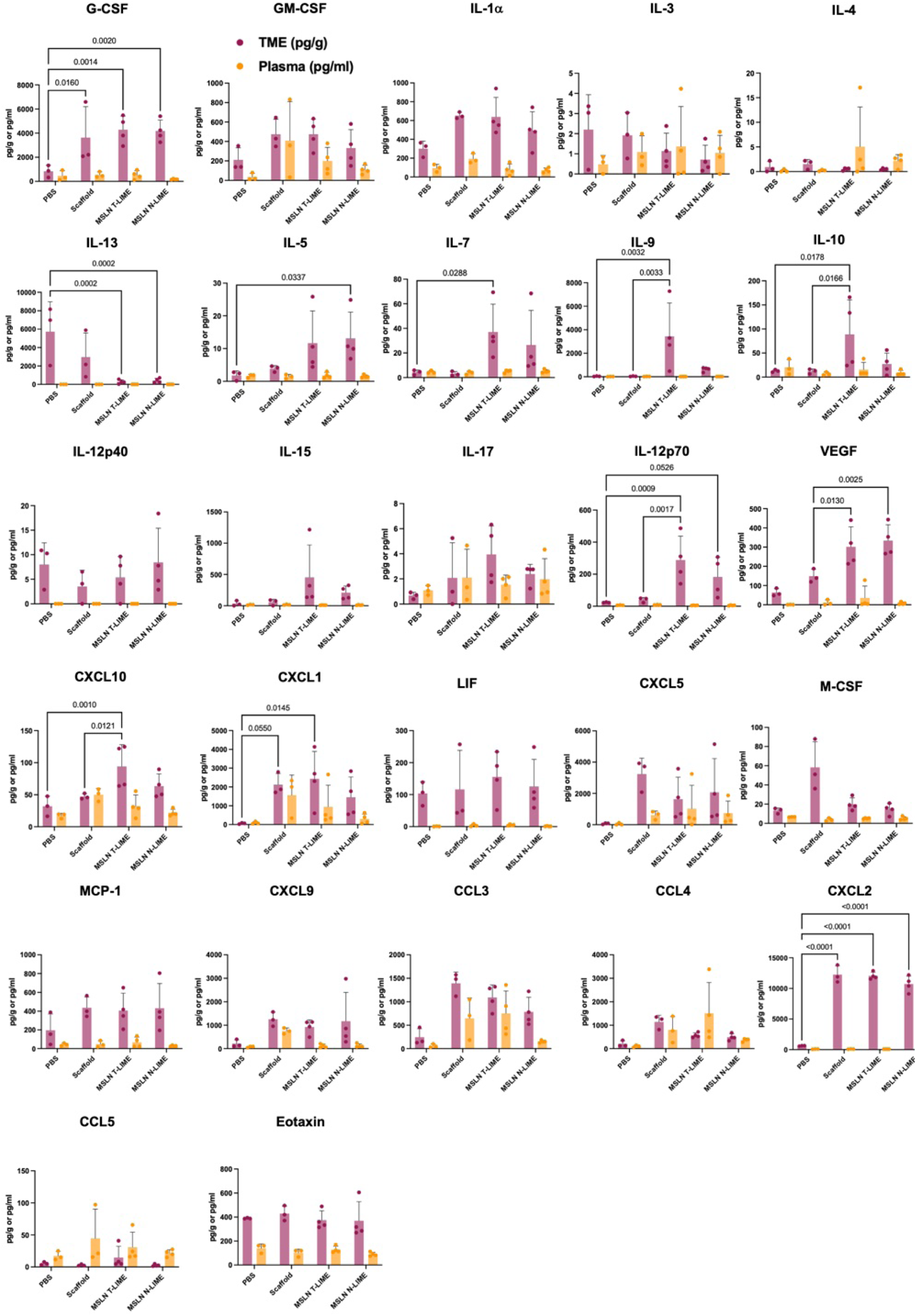
Luminex-based analysis of immune-relevant soluble factors in tumor lysates versus plasma collected from KPC C2 tumor-bearing mice at the experimental end point (day 23). Statistical significance was determined using two-way ANOVA with Tukey’s post hoc test. Data are presented as mean ± s.d.

**Extended Data Fig. 7.**
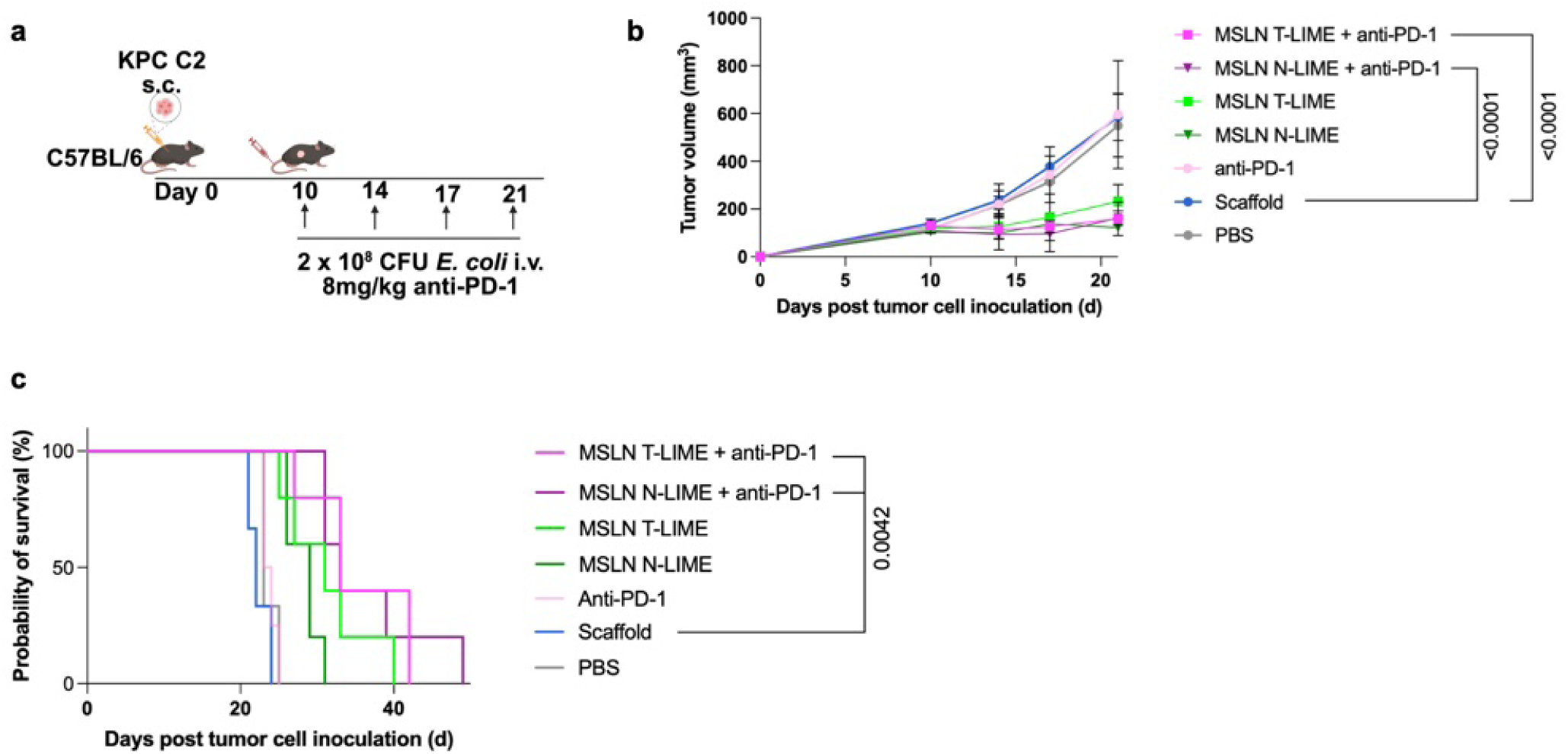
MSLN-targeting LIME combined with anti-PD-1 did not improve antitumor activity in the KPC C2 pancreatic cancer model. **a**, Experimental schema. C57BL/6 mice were subcutaneously implanted with KPC C2 pancreatic cancer cells and treated intravenously with 2 × 10⁸ CFU *E. coli* displaying MSLN-targeting T-LIME or N-LIME, scaffold control, or PBS, with or without anti-PD-1 antibody (8 mg/kg) on the indicated days. **b**, Tumor growth kinetics of the indicated treatment groups. **c**, Kaplan–Meier survival analysis of the indicated treatment groups. Statistical significance was determined using two-way ANOVA with Tukey’s post hoc test (**b**) and Logrank (Mantel-Cox) test (**c**). Data are presented as mean ± s.d. (**c**).

**Supplemental Fig. 43.**
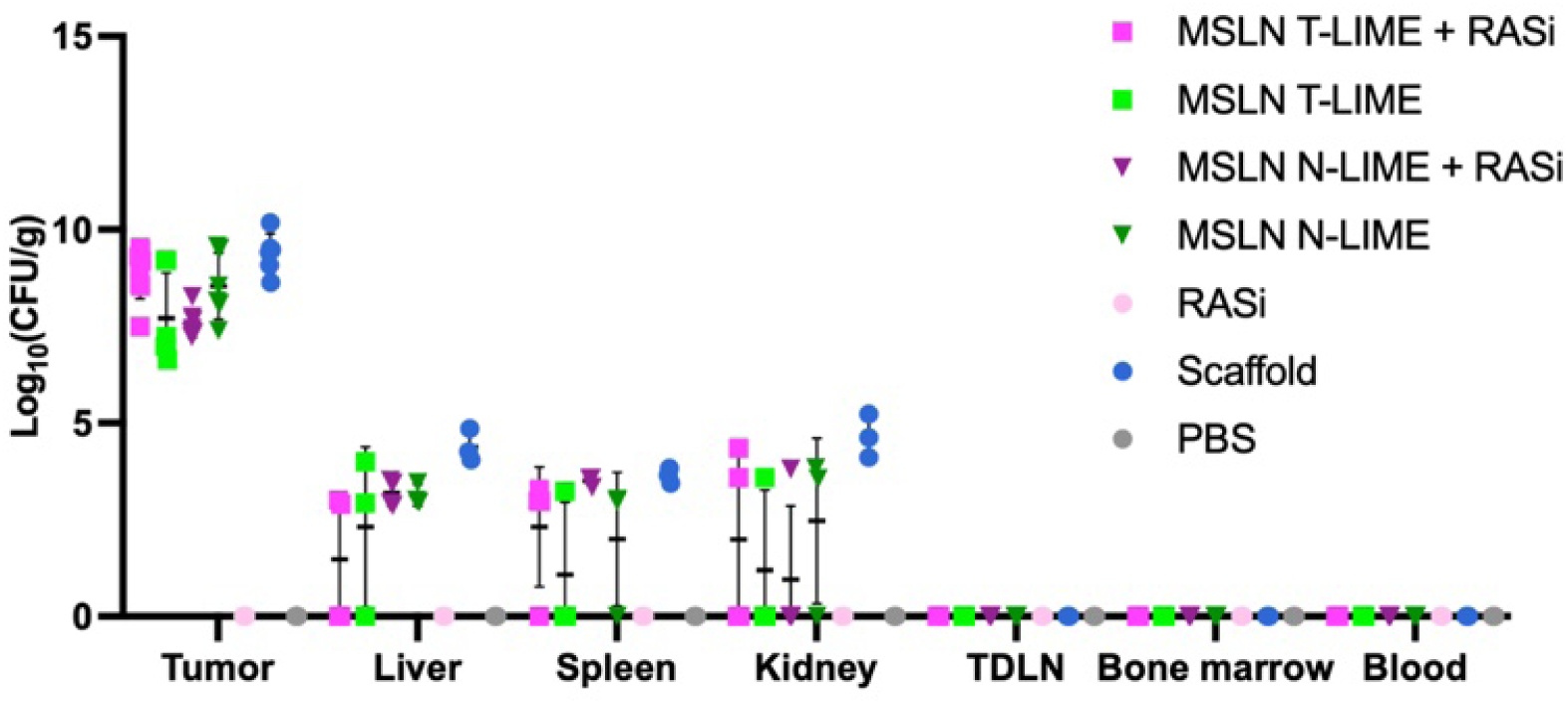
Biodistribution of MSLN-targeting LIME following treatment in the KPC C2 pancreatic cancer model. Bacterial burdens in tumor, liver, spleen, kidney, tumor-draining lymph node (TDLN), bone marrow and blood following treatment with MSLN T-LIME, MSLN N-LIME, scaffold control, PBS, RAS inhibitor (RASi), or the indicated LIME plus RASi combinations. Bacterial abundance is shown as log₁₀(CFU g⁻¹). Data are presented as mean ± s.d.

**Extended Data Fig. 8.**
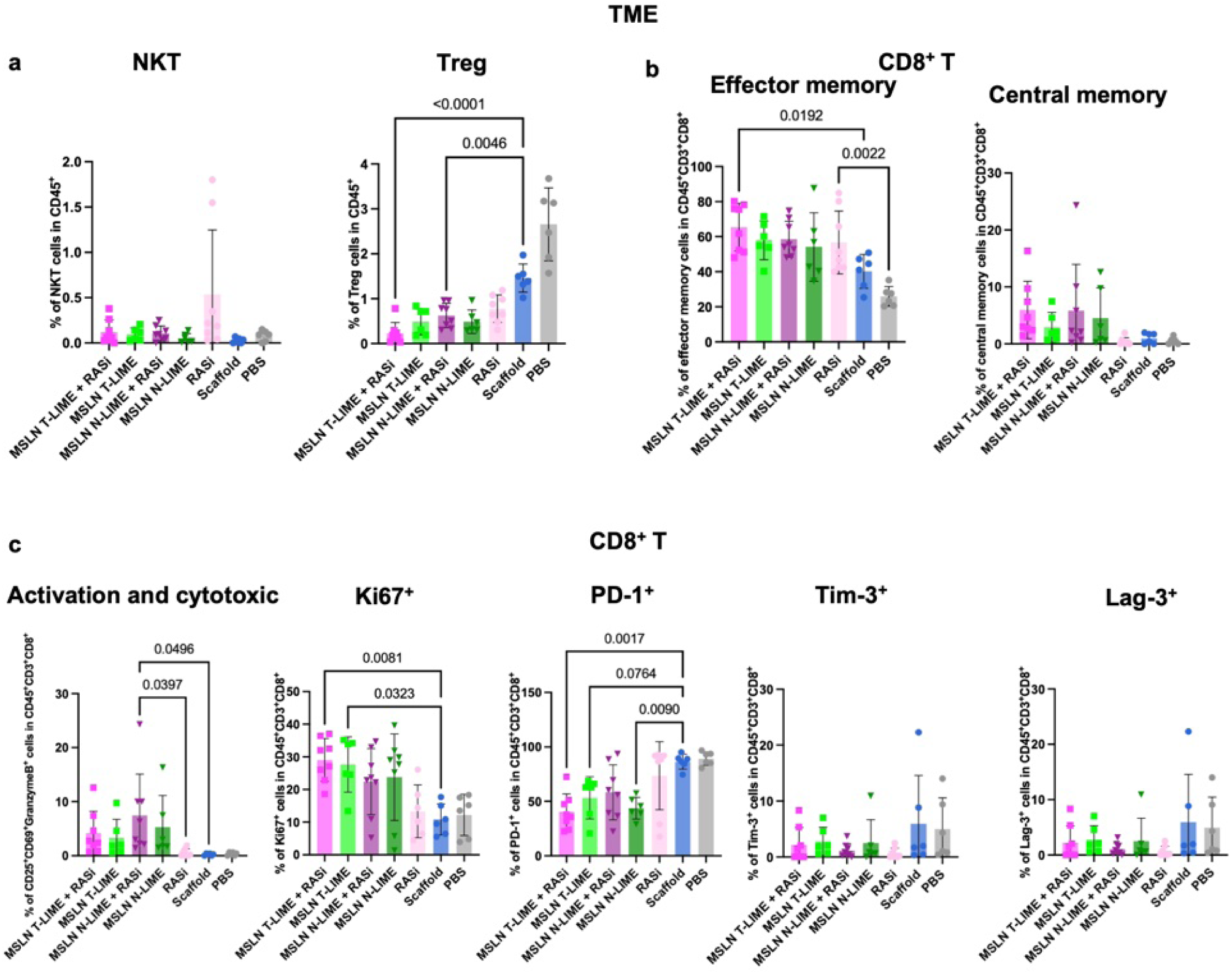
Phenotypic characterization of tumor-infiltrating NKT, Treg and CD8^+^ T cells following LIME and RASi treatment. **a**, Frequencies of NKT cells and regulatory T cells (Treg) among CD45^+^ cells in the tumor microenvironment (TME). **b**, Frequencies of effector memory and central memory CD8^+^ T cells. **c**, Frequencies of activation, proliferation, cytotoxicity and inhibitory-marker populations among CD8^+^ T cells, including CD25^+^CD69^+^Granzyme B^+^, Ki67^+^, PD-1^+^, Tim-3^+^ and Lag-3^+^ cells. Statistical significance was determined using one-way ANOVA with Tukey’s post hoc test (**a-c**). Data are presented as mean ± s.d. (**a-c**).

**Extended Data Fig. 9.**
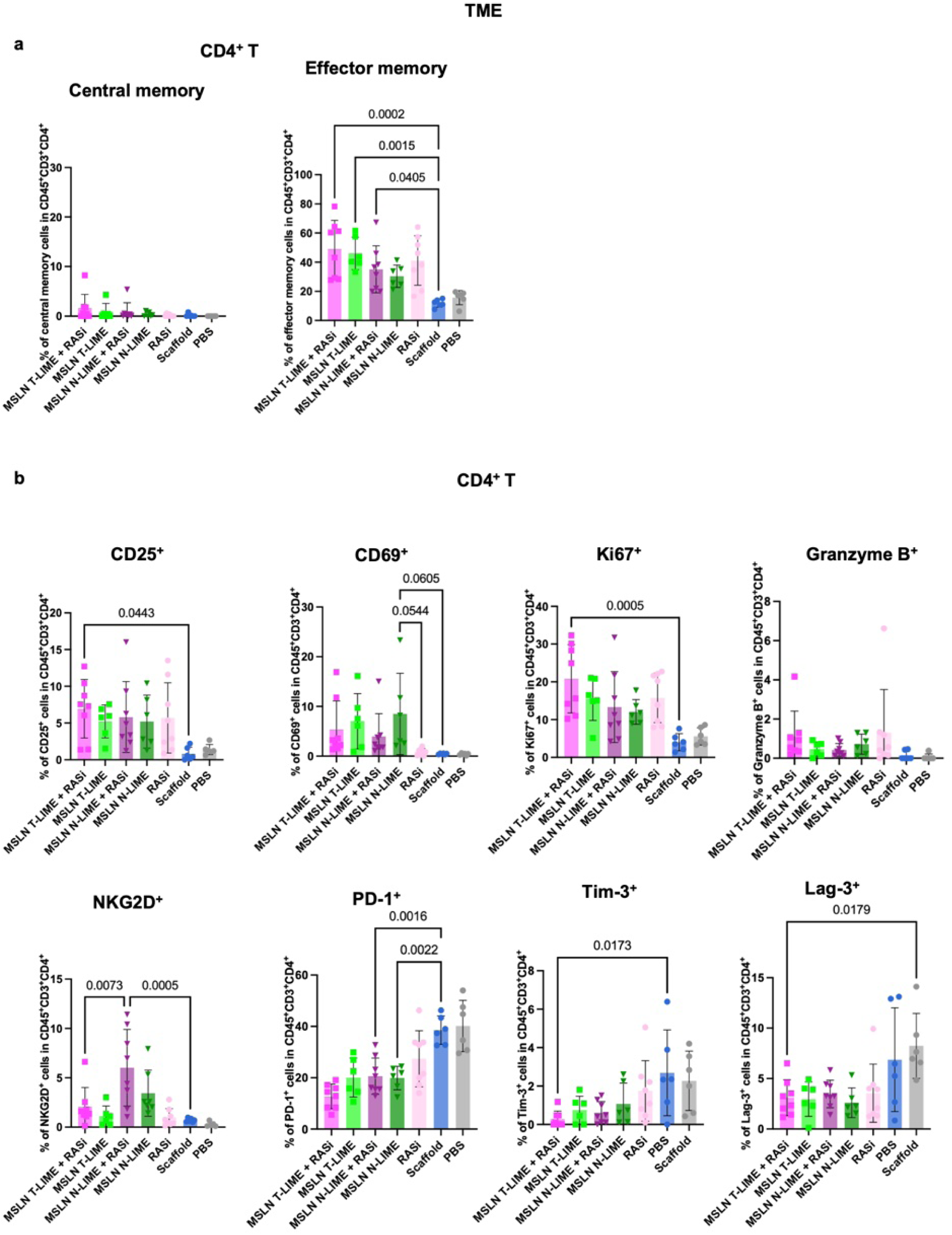
Phenotypic characterization of tumor-infiltrating CD4^+^ T cells following LIME and RASi treatment. **a**, Frequencies of central memory and effector memory CD4^+^ T cells in the TME. **b**, Frequencies of CD25^+^, CD69^+^, Ki67^+^, Granzyme B^+^, NKG2D^+^, PD-1^+^, Tim-3^+^ and Lag-3^+^ populations among CD4^+^ T cells. Statistical significance was determined using one-way ANOVA with Tukey’s post hoc test (**a**, **b**). Data are presented as mean ± s.d. (**a**, **b**).

**Supplemental Fig. 44.**
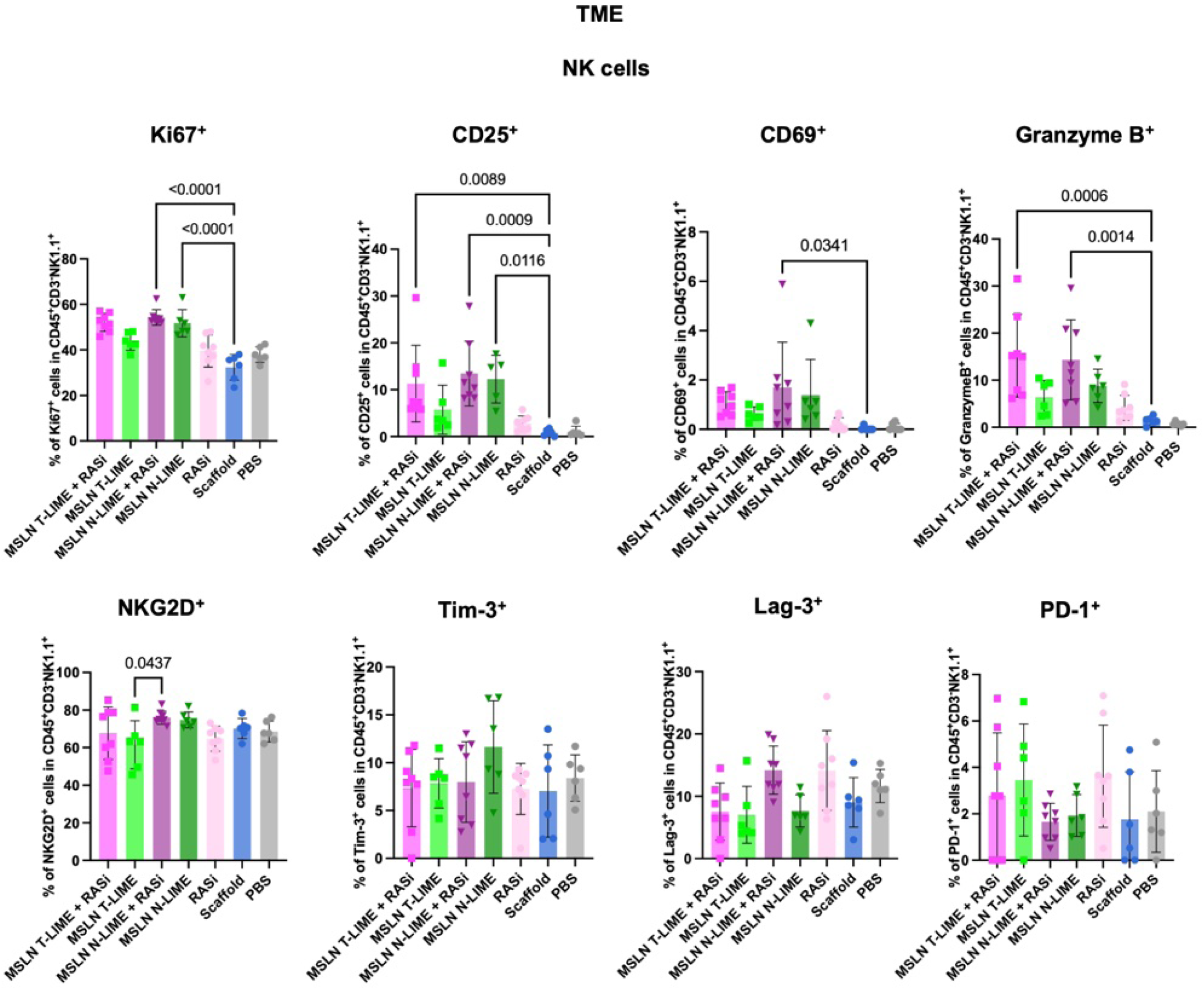
Phenotypic characterization of tumor-infiltrating NK cells following LIME and RASi treatment. Frequencies of CD25^+^, CD69^+^, Granzyme B^+^, Ki67^+^, NKG2D^+^, Tim-3^+^, Lag-3^+^ and PD-1^+^ populations among NK cells in the TME. Statistical significance was determined using one-way ANOVA with Tukey’s post hoc test. Data are presented as mean ± s.d.

**Supplemental Fig. 45.**
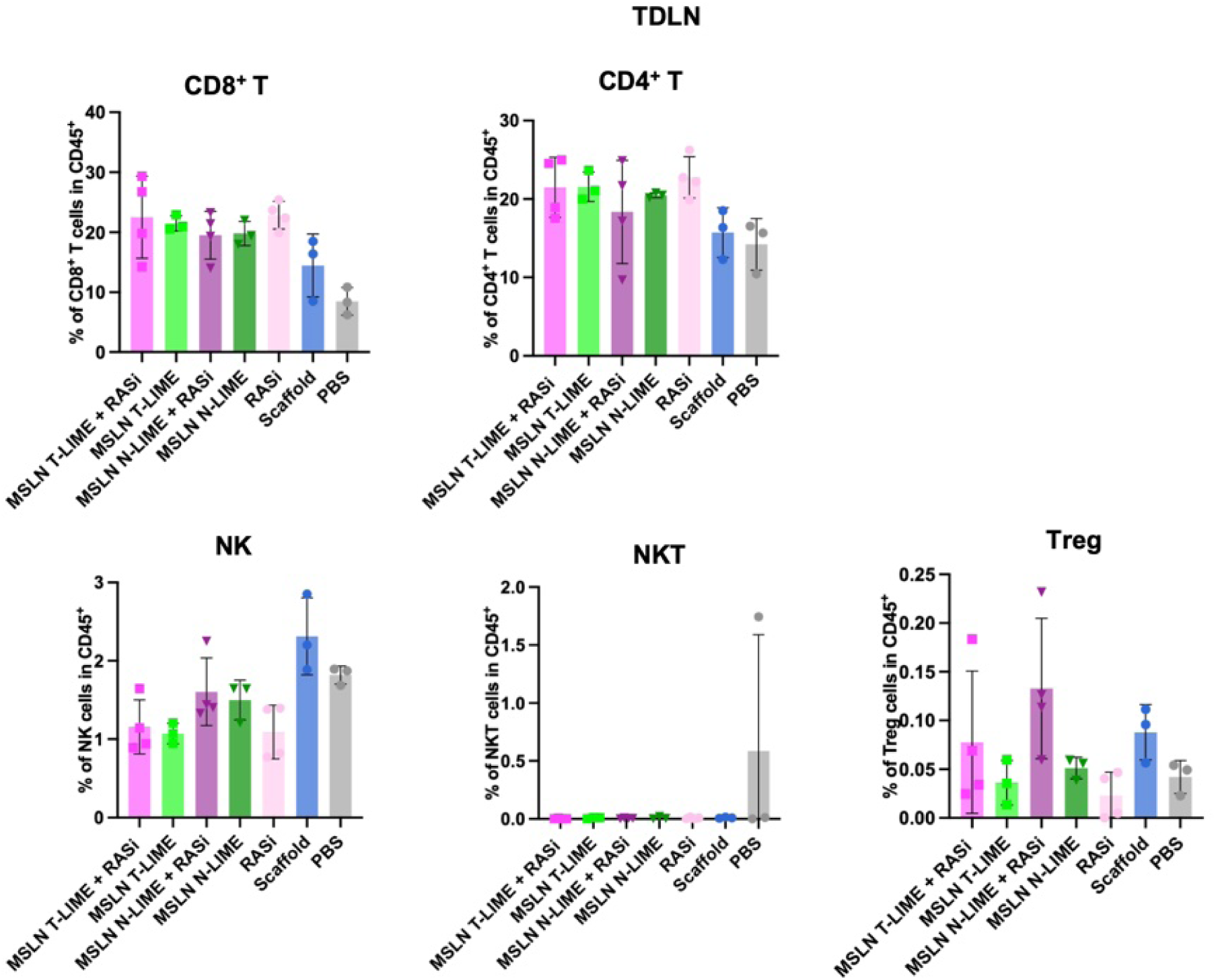
Lymphocyte composition in tumor-draining lymph nodes following LIME and RASi treatment. Frequencies of CD8^+^ T cells, CD4^+^ T cells, NK cells, NKT cells and Treg cells among CD45^+^ cells in the TDLN. Data are presented as mean ± s.d.

**Supplemental Fig. 46.**
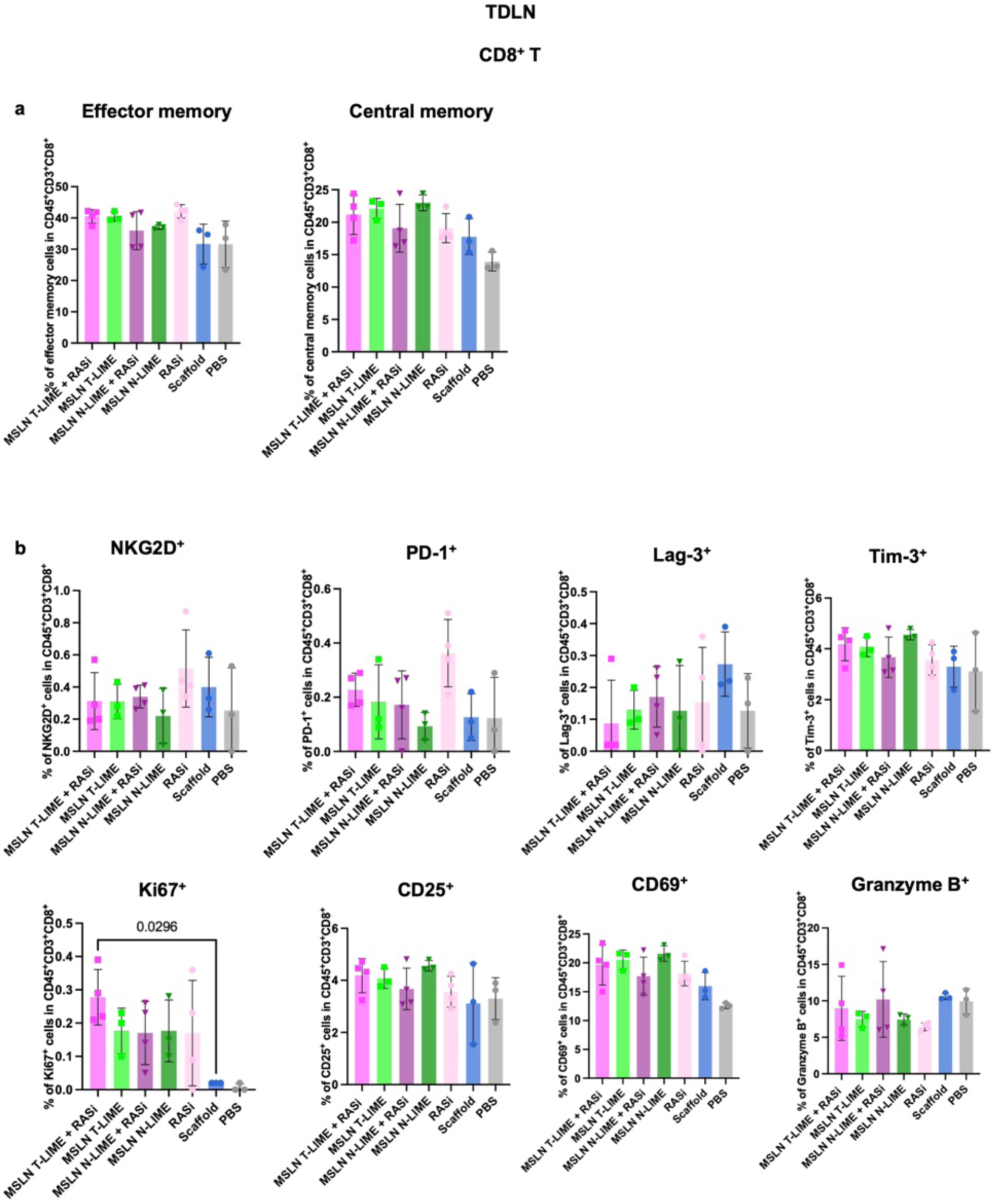
Phenotypic characterization of CD8^+^ T cells in tumor-draining lymph nodes following LIME and RASi treatment. **a**, Frequencies of effector memory and central memory CD8^+^ T cells in the TDLN. **b**, Frequencies of Ki67^+^, CD25^+^, CD69^+^, Granzyme B^+^, NKG2D^+^, PD-1^+^, Lag-3^+^ and Tim-3^+^ populations among CD8^+^ T cells. Statistical significance was determined using one-way ANOVA with Tukey’s post hoc test. Data are presented as mean ± s.d. (**a**, **b**).

**Supplemental Fig. 47.**
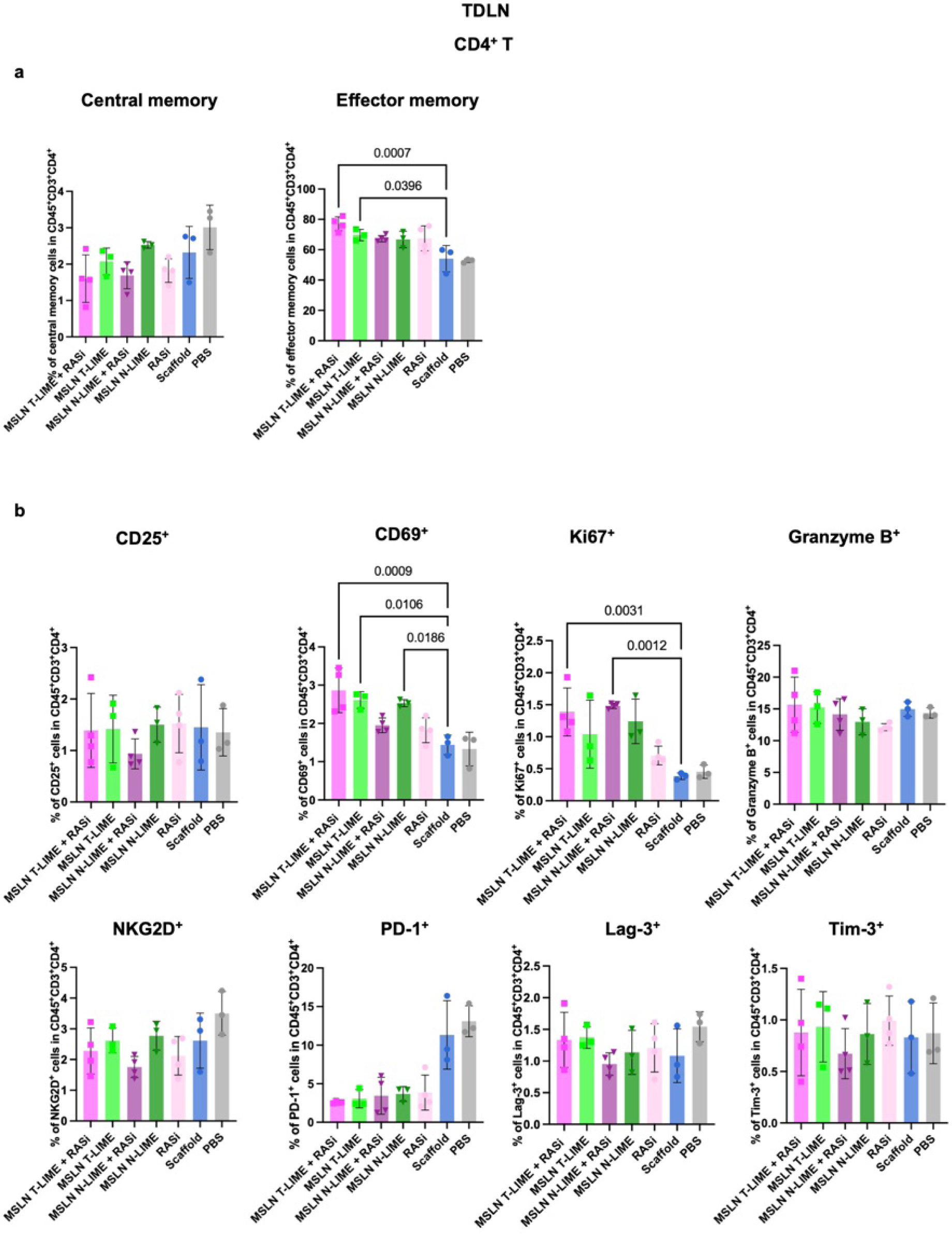
Phenotypic characterization of CD4^+^ T cells in tumor-draining lymph nodes following LIME and RASi treatment. **a**, Frequencies of central memory and effector memory CD4^+^ T cells in the TDLN. **b**, Frequencies of CD25^+^, CD69^+^, Ki67^+^, Granzyme B^+^, NKG2D^+^, PD-1^+^, Lag-3^+^ and Tim-3^+^ populations among CD4^+^ T cells. Statistical significance was determined using one-way ANOVA with Tukey’s post hoc test. Data are presented as mean ± s.d. (**a**, **b**).

**Supplemental Fig. 48.**
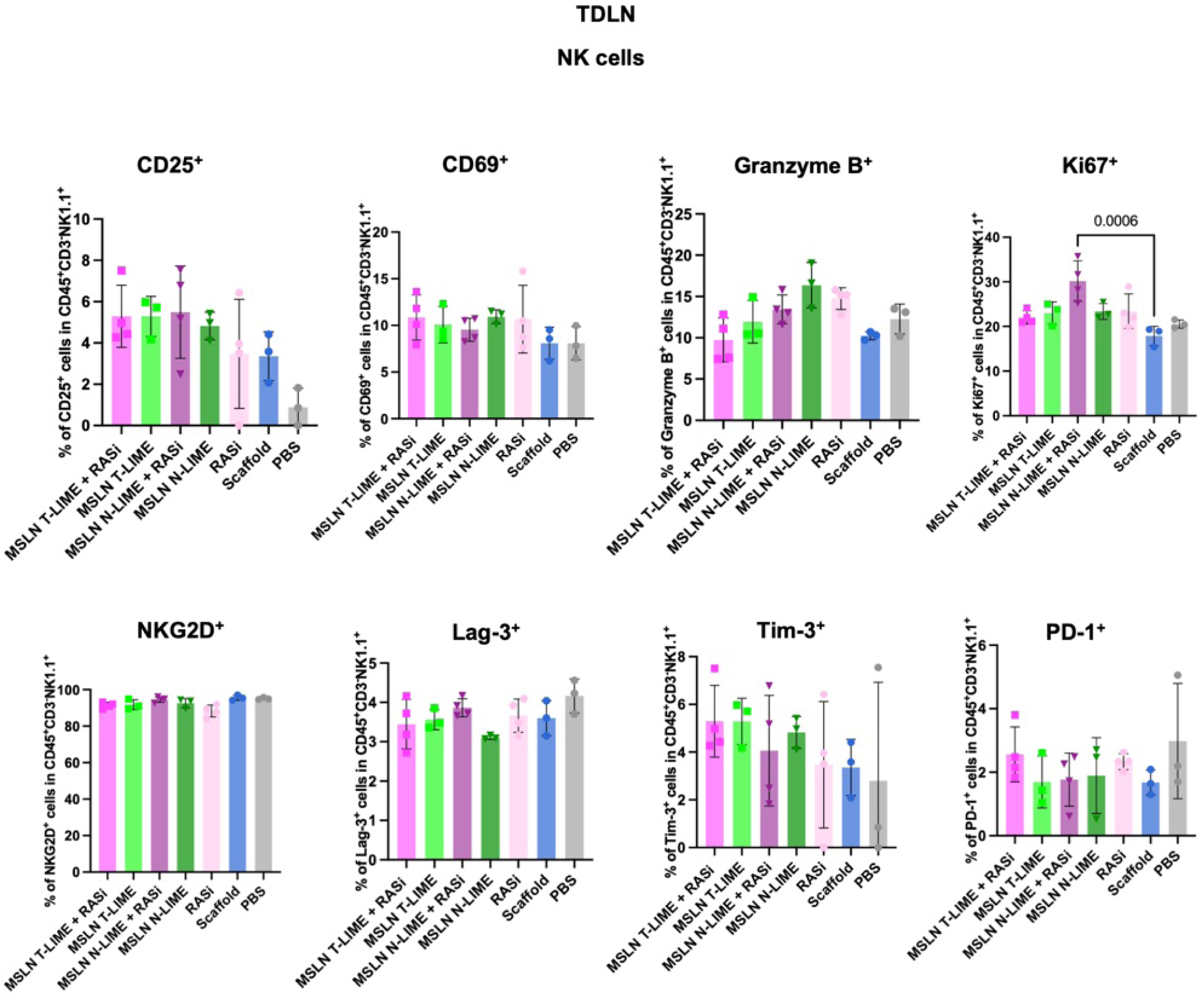
Phenotypic characterization of NK cells in tumor-draining lymph nodes following LIME and RASi treatment. Frequencies of CD25^+^, CD69^+^, Granzyme B^+^, Ki67^+^, NKG2D^+^, Lag-3^+^, Tim-3^+^ and PD-1^+^ populations among NK cells in the TDLN. Statistical significance was determined using one-way ANOVA with Tukey’s post hoc test. Data are presented as mean ± s.d.

**Supplemental Fig. 49.**
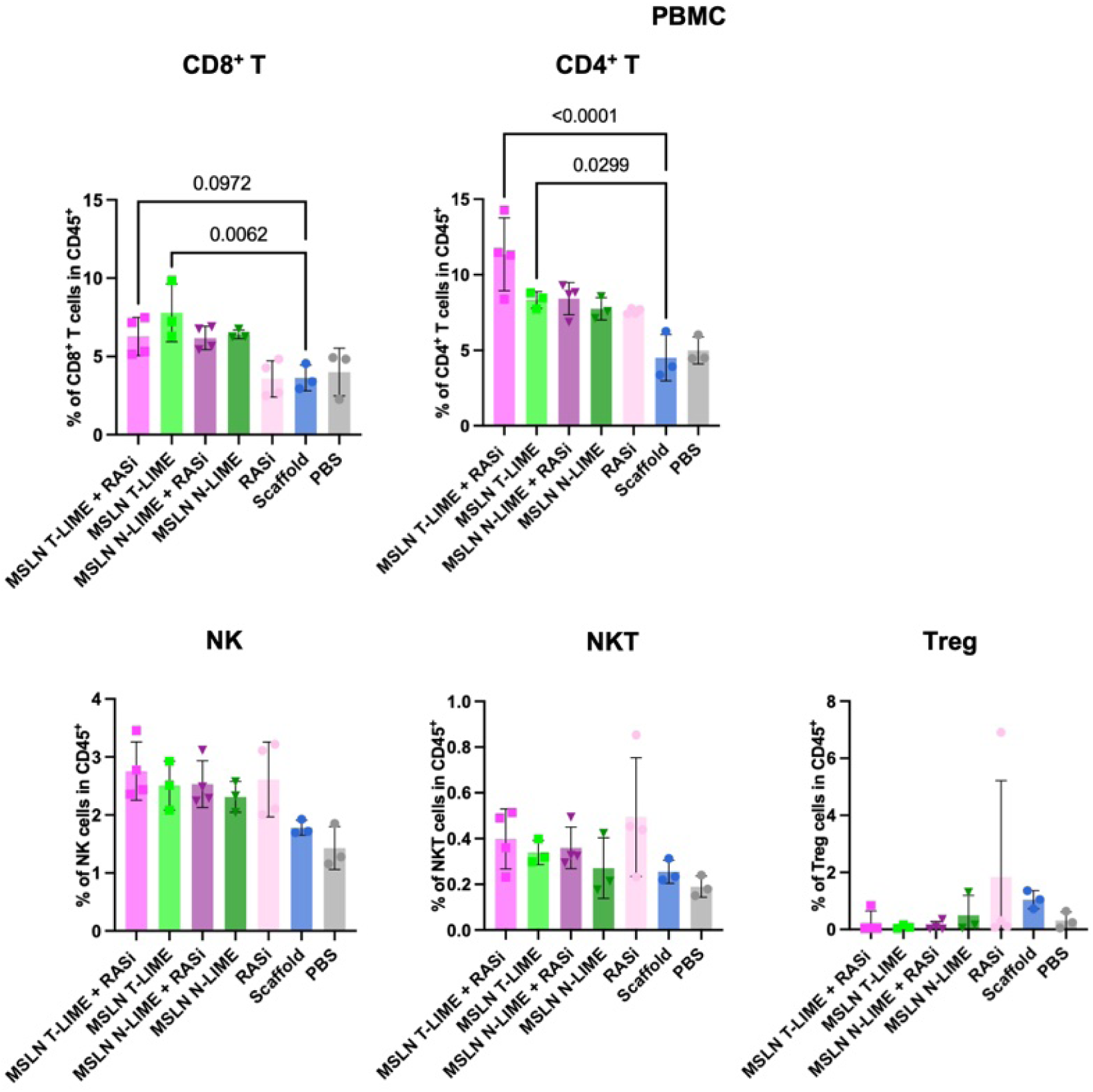
Peripheral blood lymphocyte composition following LIME and RASi treatment. Frequencies of CD8^+^ T cells, CD4^+^ T cells, NK cells, NKT cells and Treg cells among CD45^+^ cells in peripheral blood mononuclear cells (PBMCs). Statistical significance was determined using one-way ANOVA with Tukey’s post hoc test. Data are presented as mean ± s.d.

**Supplemental Fig. 50.**
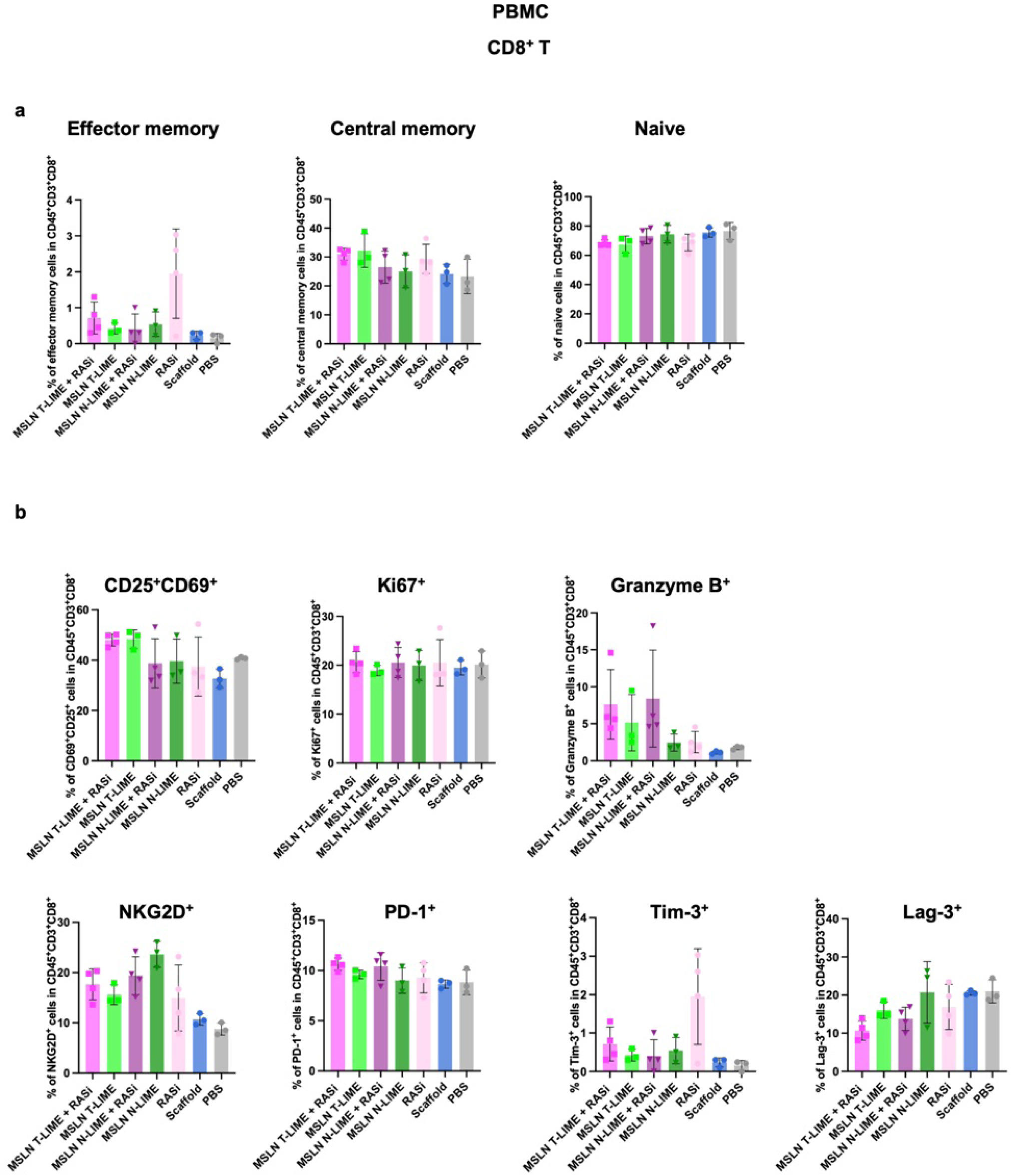
Phenotypic characterization of peripheral blood CD8^+^ T cells following LIME and RASi treatment. **a**, Frequencies of naive, central memory and effector memory CD8^+^ T cells in PBMCs. **b**, Frequencies of CD25^+^, CD69^+^, Ki67^+^, Granzyme B^+^, NKG2D^+^, PD-1^+^, Tim-3^+^ and Lag-3^+^ populations among CD8^+^ T cells. Data are presented as mean ± s.d. (**a**, **b**).

**Supplemental Fig. 51.**
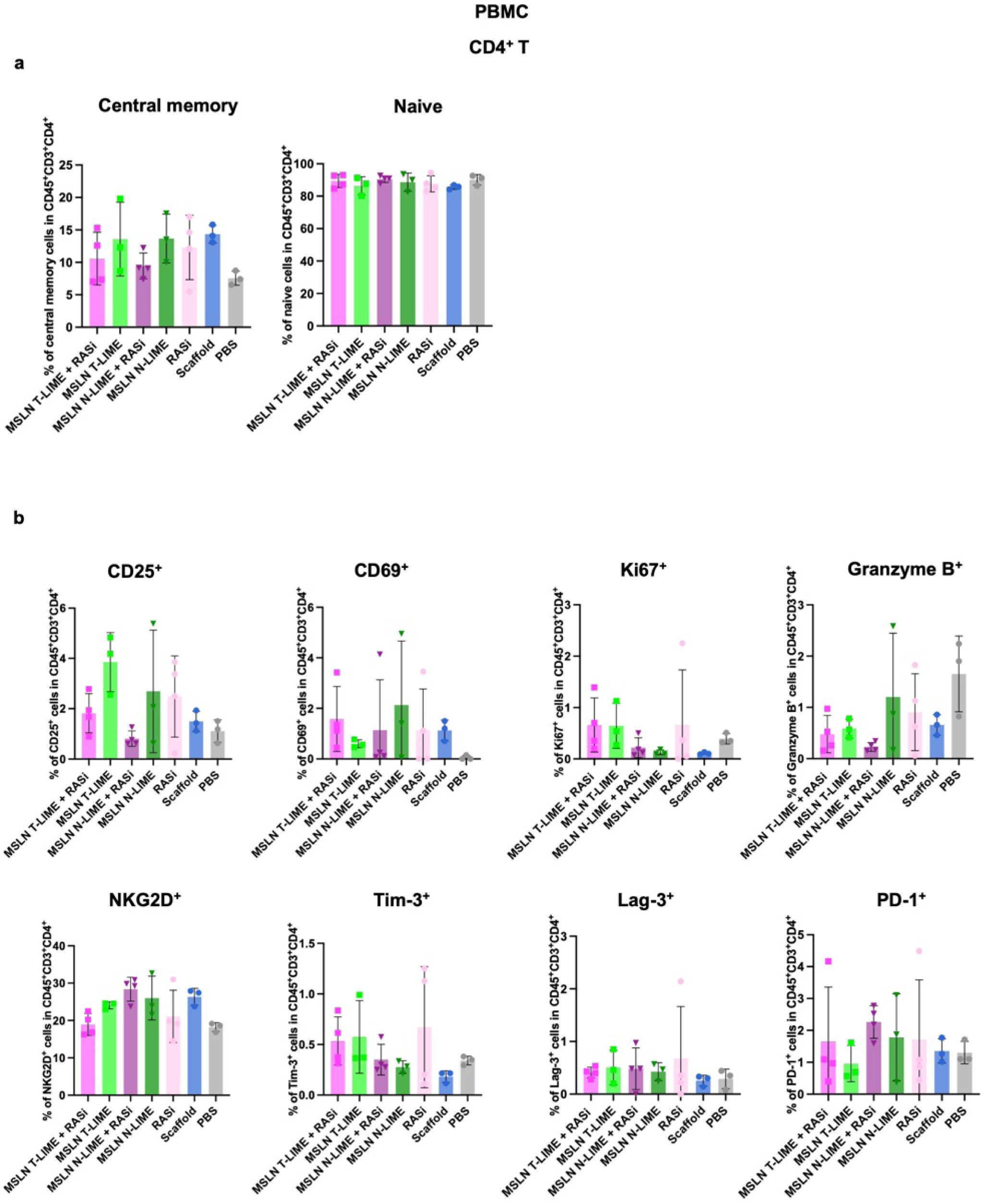
Phenotypic characterization of peripheral blood CD4^+^ T cells following LIME and RASi treatment. **a**, Frequencies of naive and central memory CD4^+^ T cells in PBMCs. **b**, Frequencies of Ki67^+^, CD25^+^, CD69^+^, Tim-3^+^, Lag-3^+^, PD-1^+^, NKG2D^+^ and Granzyme B^+^ populations among CD4^+^ T cells. Data are presented as mean ± s.d. (**a**, **b**).

**Supplemental Fig. 52.**
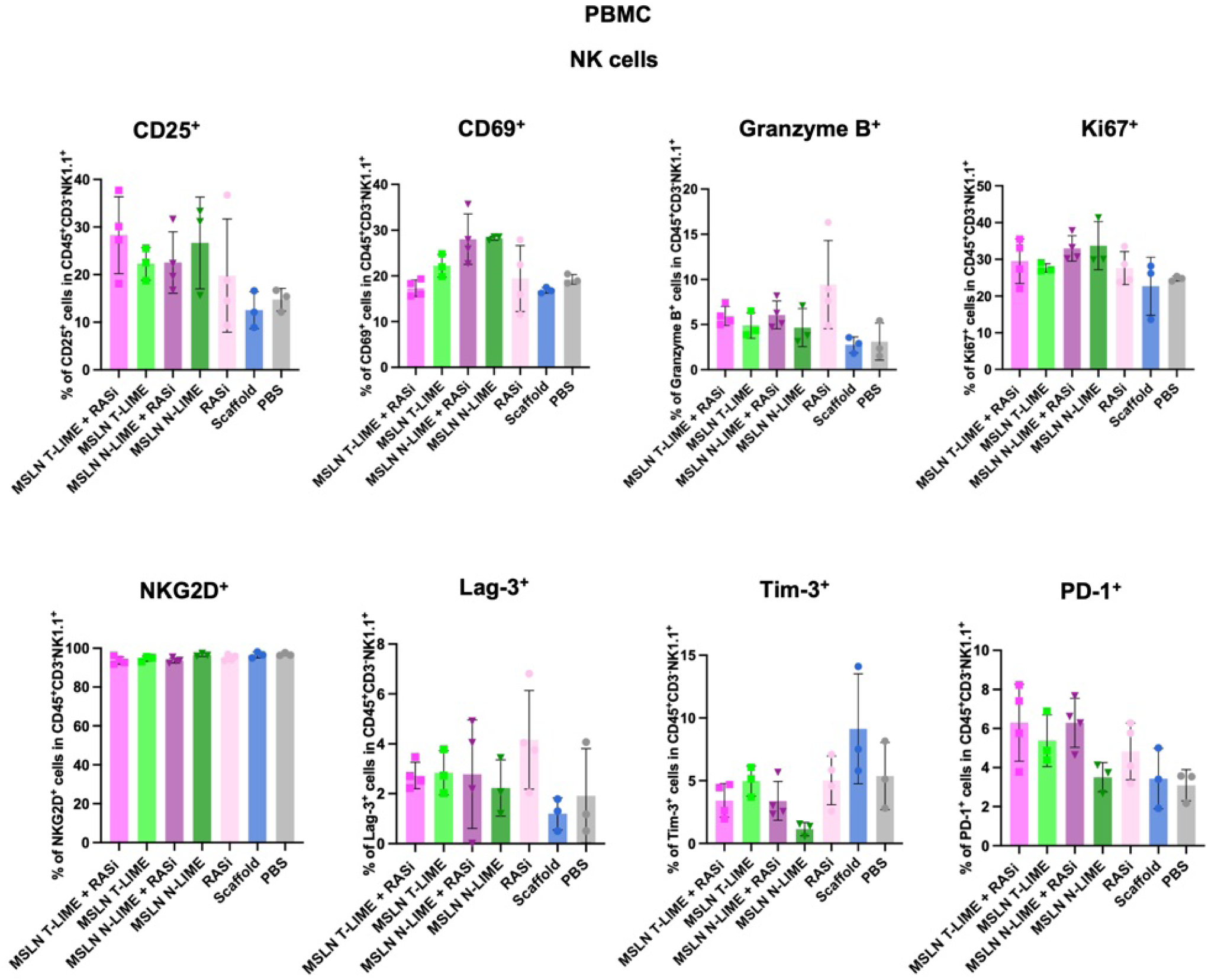
Phenotypic characterization of peripheral blood NK cells following LIME and RASi treatment. Frequencies of CD25^+^, CD69^+^, Granzyme B^+^, Ki67^+^, NKG2D^+^, Lag-3^+^, Tim-3^+^ and PD-1^+^ populations among NK cells in PBMCs. Data are presented as mean ± s.d.

**Supplemental Fig. 53.**
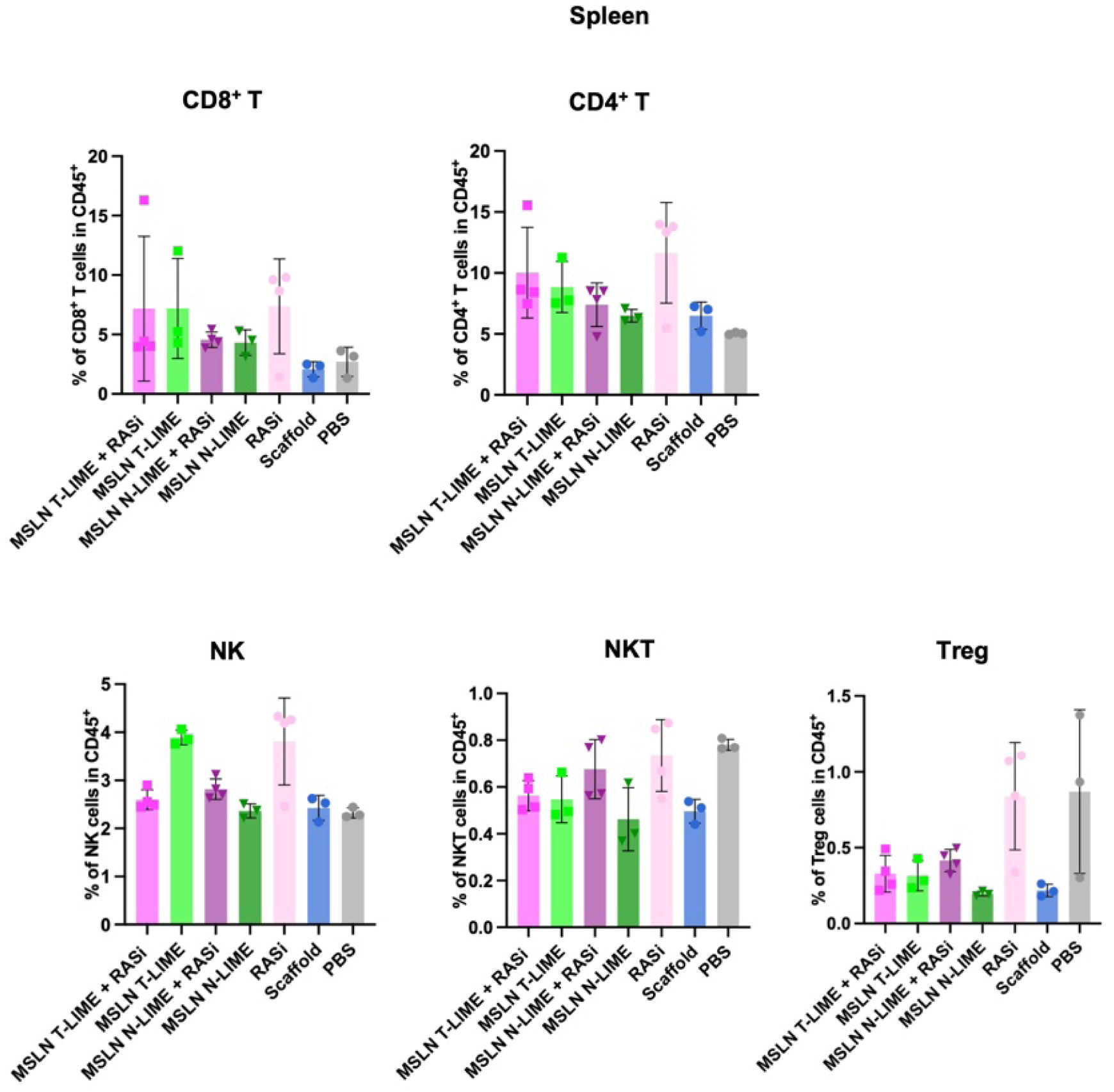
Splenic lymphocyte composition following LIME and RASi treatment. Frequencies of CD8^+^ T cells, CD4^+^ T cells, NK cells, NKT cells and Treg cells among CD45^+^ cells in the spleen. Data are presented as mean ± s.d.

**Supplemental Fig. 54.**
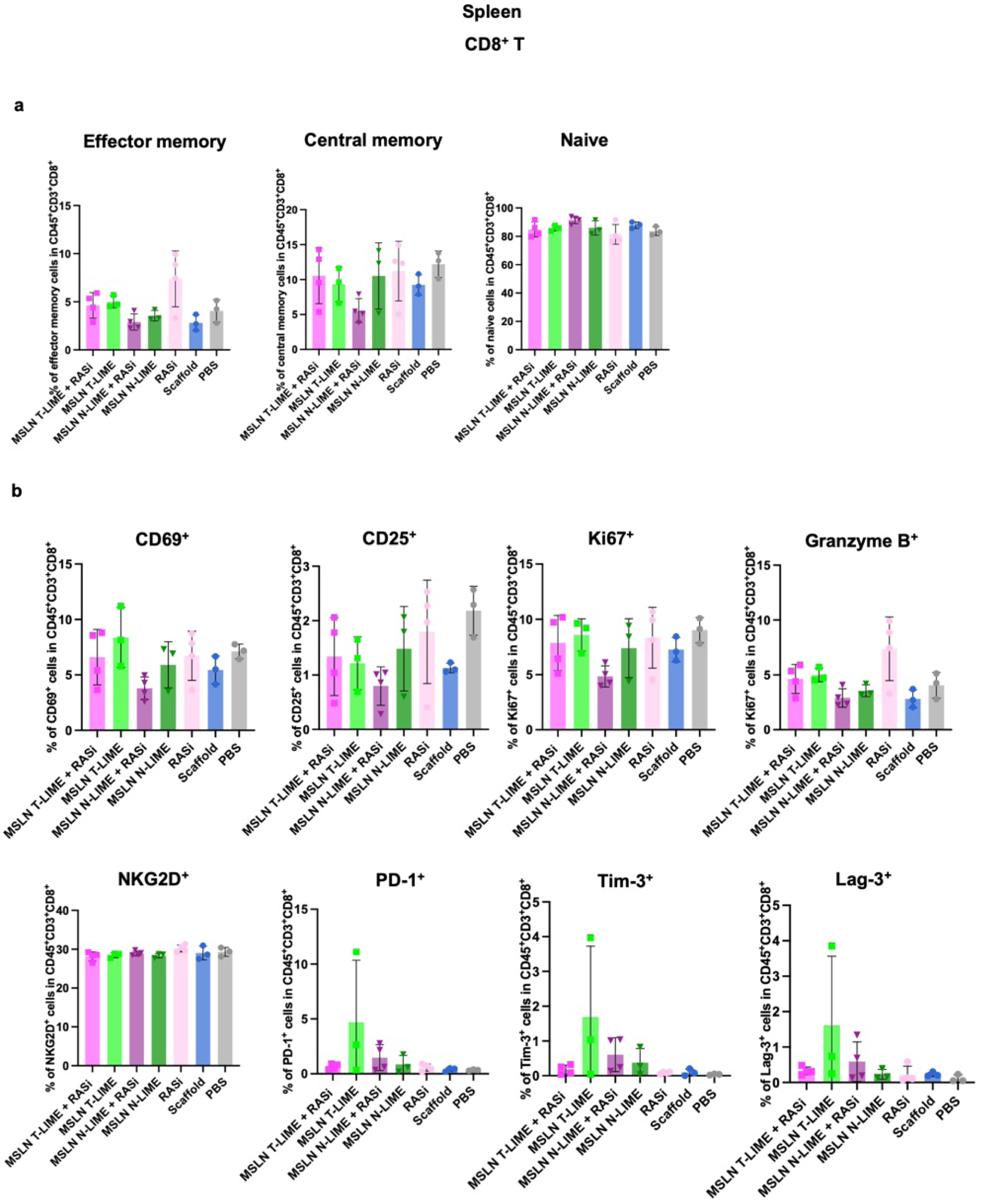
Phenotypic characterization of splenic CD8^+^ T cells following LIME and RASi treatment. **a**, Frequencies of naive, central memory and effector memory CD8^+^ T cells in the spleen. **b**, Frequencies of CD69^+^, Ki67^+^, Granzyme B^+^, NKG2D^+^, PD-1^+^, Tim-3^+^, Lag-3^+^ and CD25^+^ populations among CD8^+^ T cells. Data are presented as mean ± s.d. (**a**, **b**).

**Supplemental Fig. 55.**
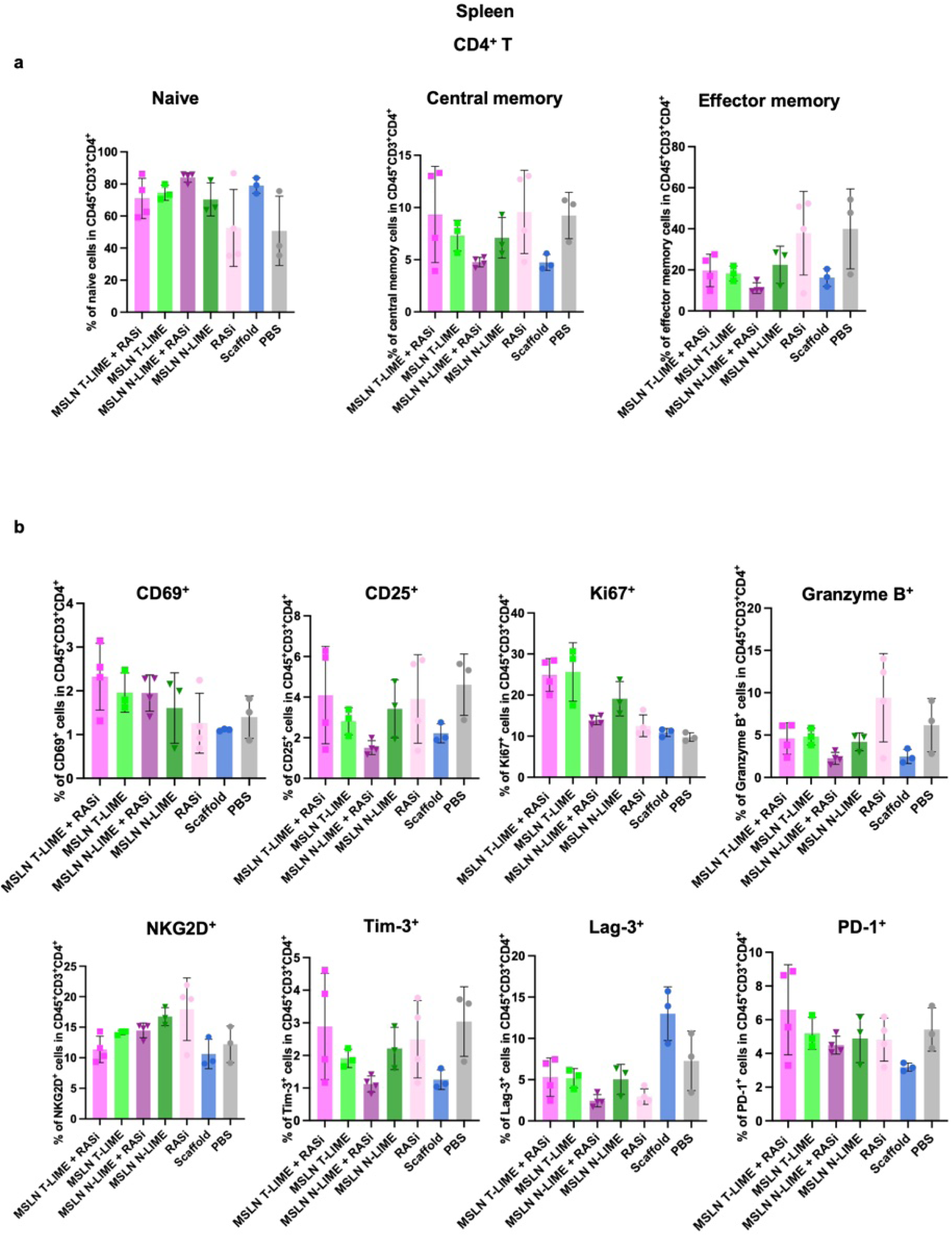
Phenotypic characterization of splenic CD4^+^ T cells following LIME and RASi treatment. **a**, Frequencies of naive, central memory and effector memory CD4^+^ T cells in the spleen. **b**, Frequencies of Ki67^+^, CD25^+^, CD69^+^, Tim-3^+^, Lag-3^+^, PD-1^+^, NKG2D^+^ and Granzyme B^+^ populations among CD4^+^ T cells. Data are presented as mean ± s.d. (**a**, **b**).

**Supplemental Fig. 56.**
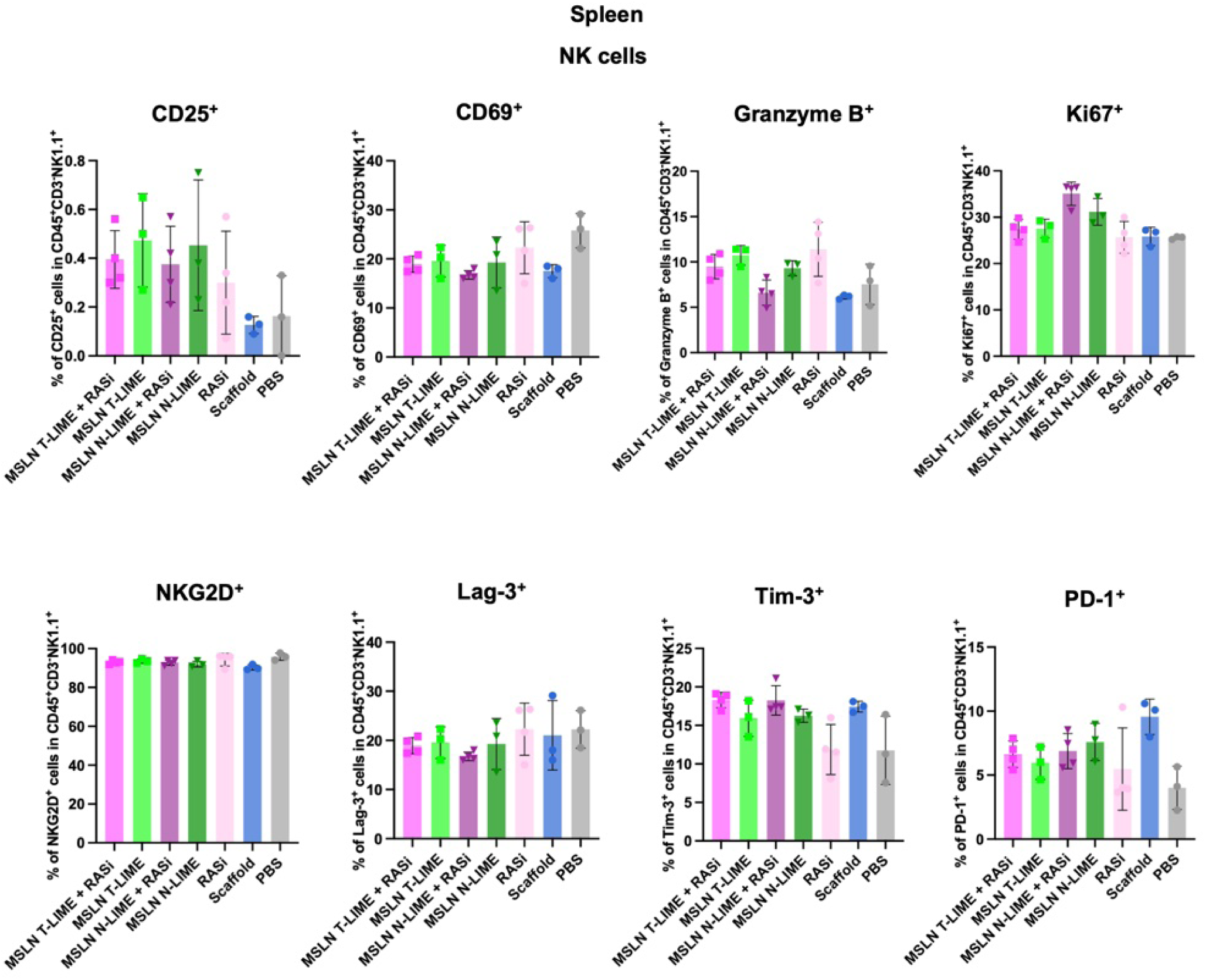
Phenotypic characterization of splenic NK cells following LIME and RASi treatment. Frequencies of CD25^+^, CD69^+^, Granzyme B^+^, Ki67^+^, NKG2D^+^, Lag-3^+^, Tim-3^+^ and PD-1^+^ populations among NK cells in the spleen. Data are presented as mean ± s.d.

**Supplemental Fig. 57.**
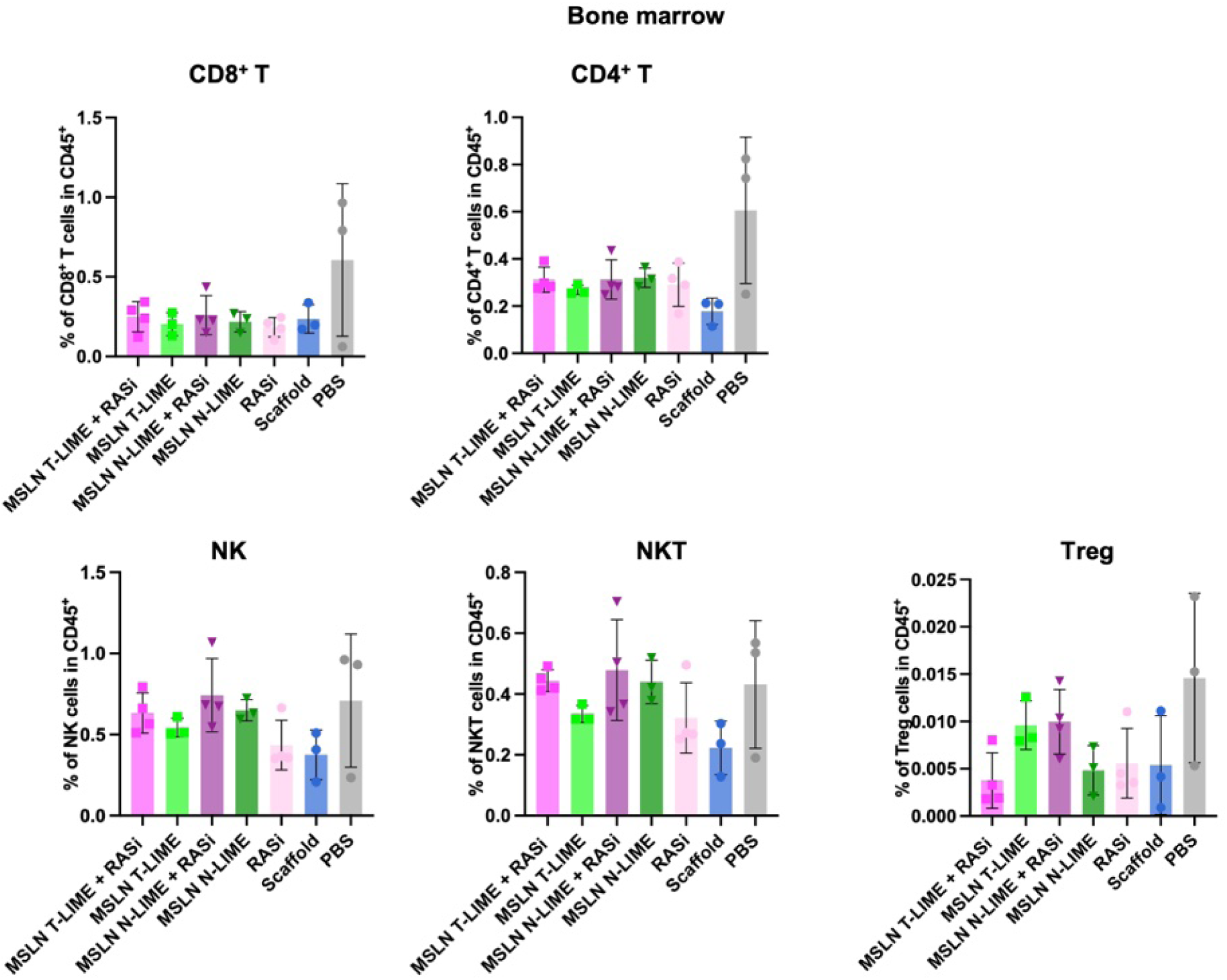
Bone marrow lymphocyte composition following LIME and RASi treatment. Frequencies of CD8^+^ T cells, CD4^+^ T cells, NK cells, NKT cells and Treg cells among CD45^+^ cells in bone marrow. Data are presented as mean ± s.d.

**Supplemental Fig. 58.**
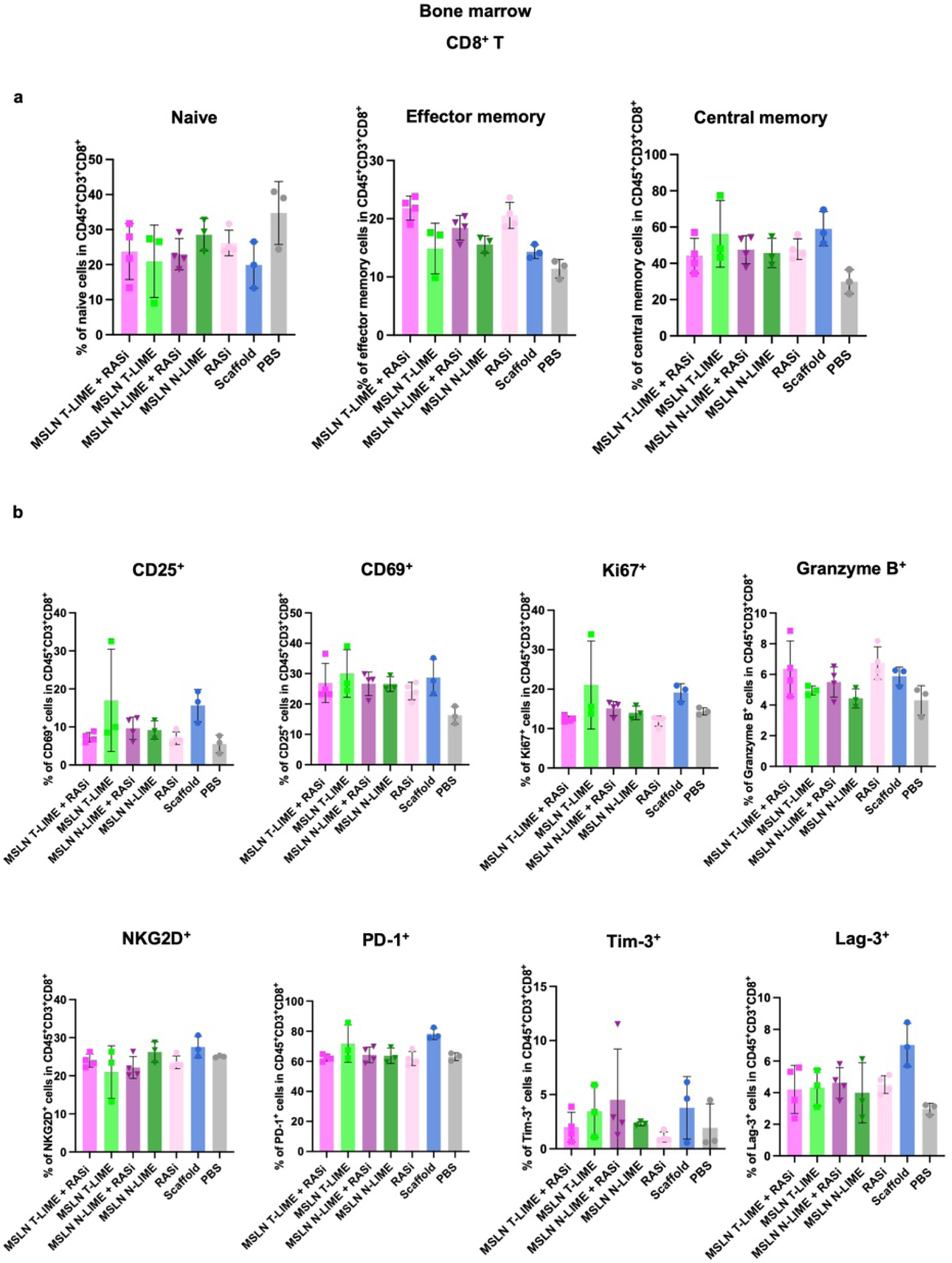
Phenotypic characterization of bone marrow CD8^+^ T cells following LIME and RASi treatment. **a**, Frequencies of naive, central memory and effector memory CD8^+^ T cells in bone marrow. **b**, Frequencies of CD69^+^, Ki67^+^, NKG2D^+^, PD-1^+^, Tim-3^+^, CD25^+^, Granzyme B^+^ and Lag-3^+^ populations among CD8^+^ T cells. Data are presented as mean ± s.d. (**a**, **b**).

**Supplemental Fig. 59.**
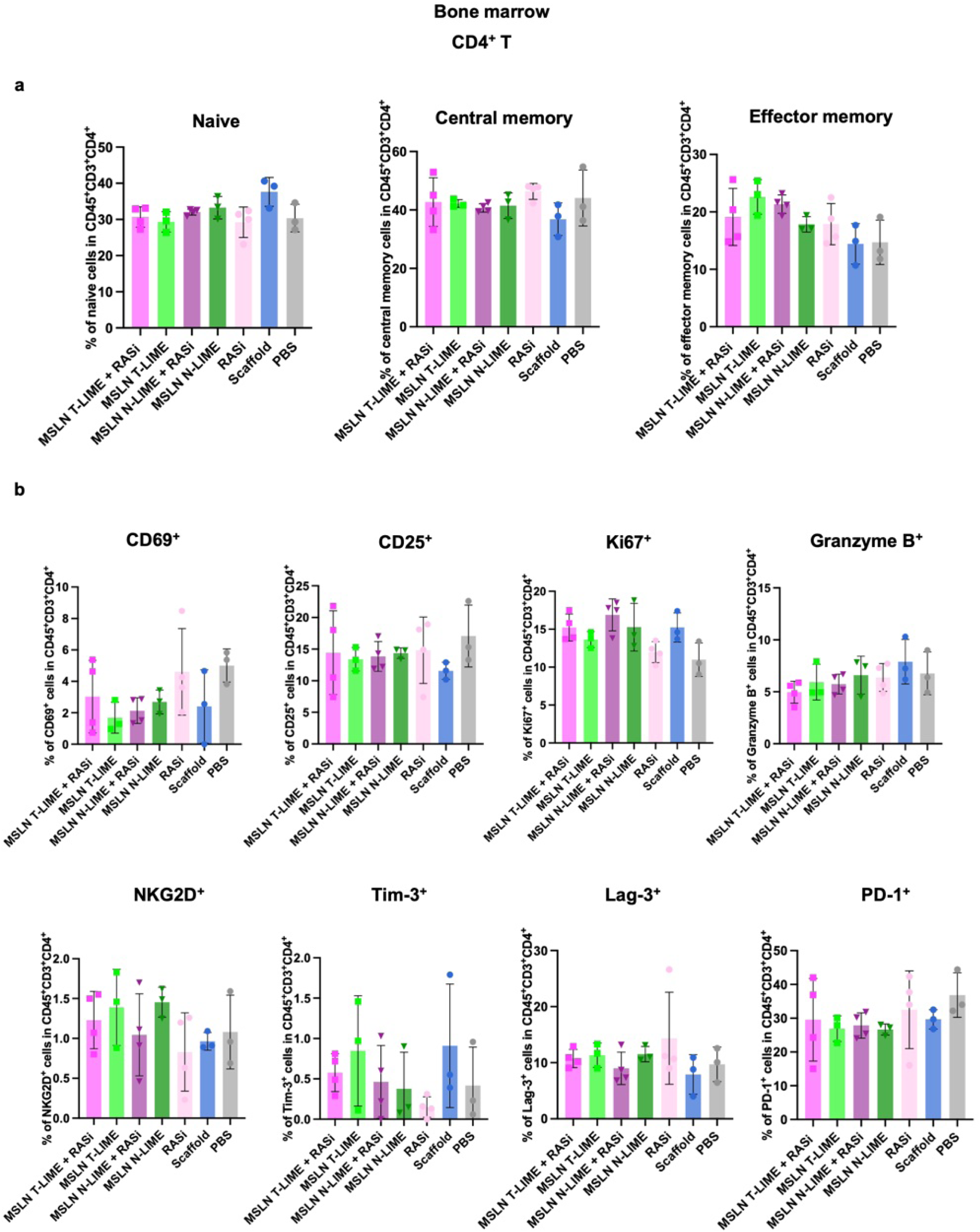
Phenotypic characterization of bone marrow CD4^+^ T cells following LIME and RASi treatment. **a**, Frequencies of naive, central memory and effector memory CD4^+^ T cells in bone marrow. **b**, Frequencies of Ki67^+^, CD25^+^, CD69^+^, Tim-3^+^, Lag-3^+^, PD-1^+^, NKG2D^+^ and Granzyme B^+^ populations among CD4^+^ T cells. Data are presented as mean ± s.d. (**a**, **b**).

**Supplemental Fig. 60.**
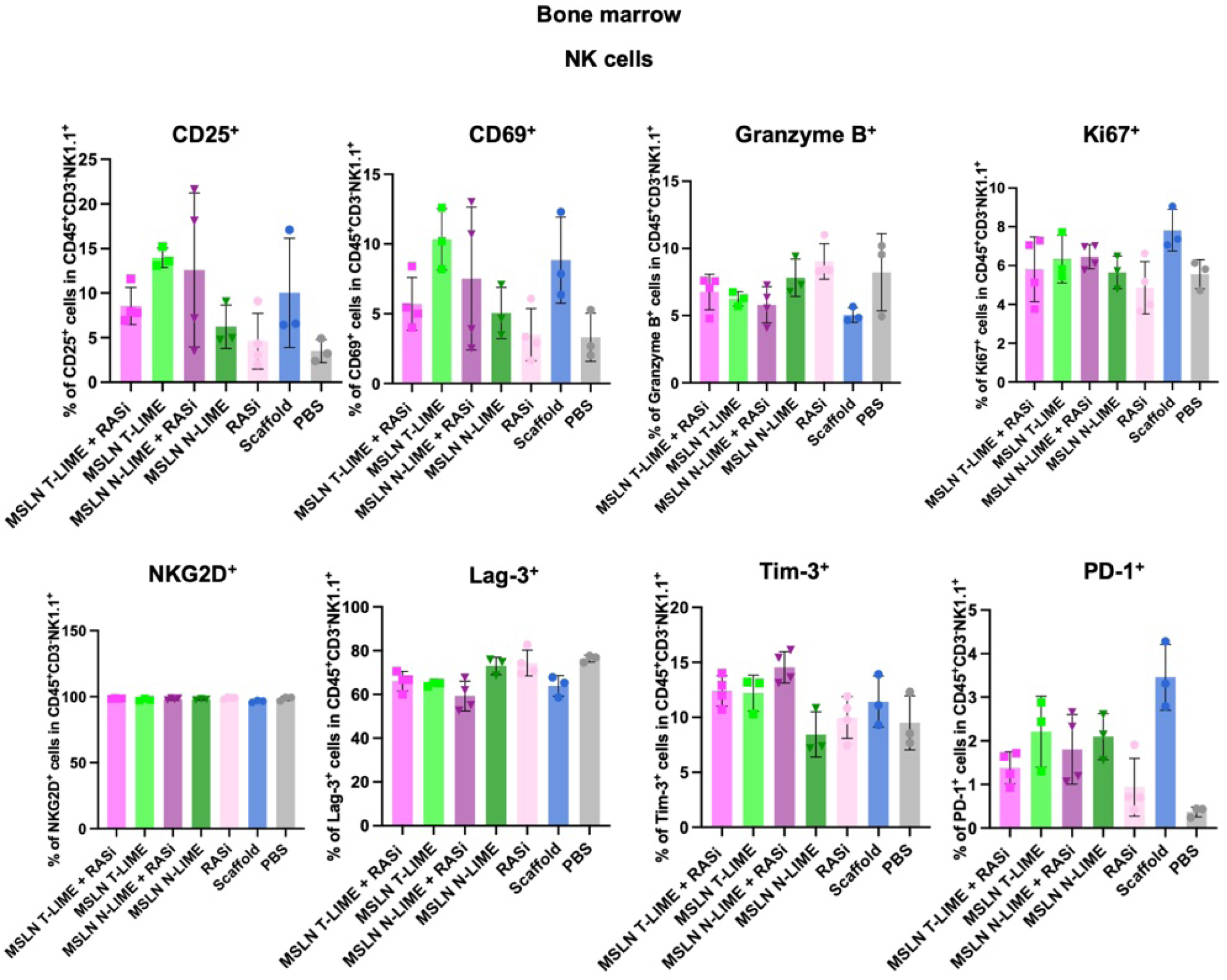
Phenotypic characterization of bone marrow NK cells following LIME and RASi treatment. Frequencies of CD25^+^, CD69^+^, Granzyme B^+^, Ki67^+^, NKG2D^+^, Lag-3^+^, Tim-3^+^ and PD-1^+^ populations among NK cells in bone marrow. Data are presented as mean ± s.d.

**Supplemental Fig. 61.**
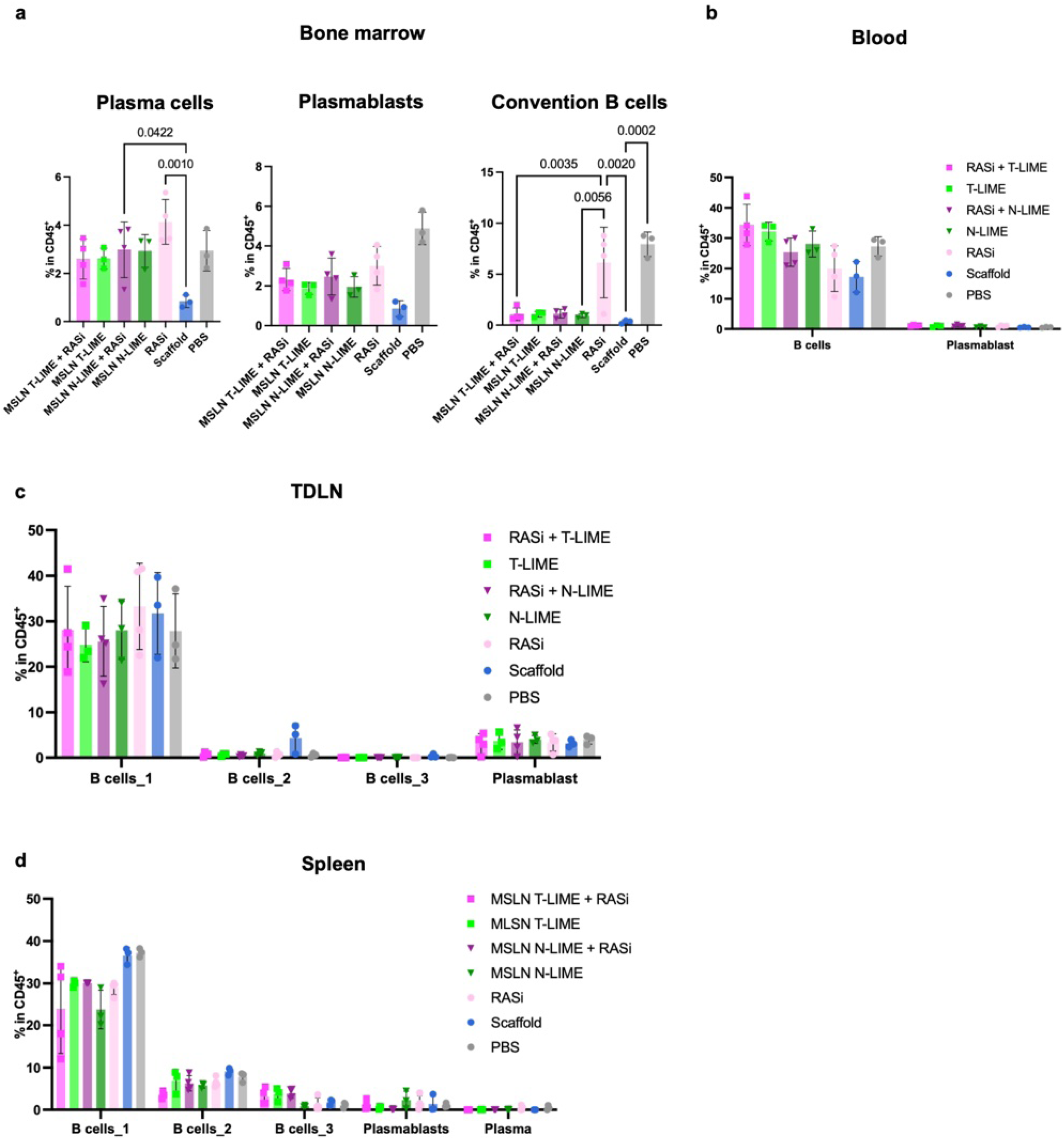
B-cell lineage populations across bone marrow, blood, TDLN and spleen following LIME and RASi treatment. **a**, Frequencies of plasma cells (CD45^+^CD38^+^CD138^+^), plasmablasts and conventional B cells (CD45^+^B220^+^CD19^+^) among CD45^+^ cells in bone marrow. **b**, Frequencies of B cells and plasmablasts among CD45^+^ cells in blood. **c**, Frequencies of B-cell subsets and plasmablasts among CD45^+^ cells in the TDLN. **d**, Frequencies of B-cell subsets, plasmablasts and plasma cells among CD45^+^ cells in the spleen. B cells_1: CD45^+^B220^+^CD19^+^IgD^high^IgM^low^, B cells_2: CD45^+^B220^+^CD19^+^IgD^low^IgM^low^, B cells_3: CD45^+^B220^+^CD19^+^IgD^low^IgM^high^. Statistical significance was determined using one-way ANOVA with Tukey’s post hoc test (**a**). Data are presented as mean ± s.d. (**a-d**).

**Extended Data Fig. 10.**
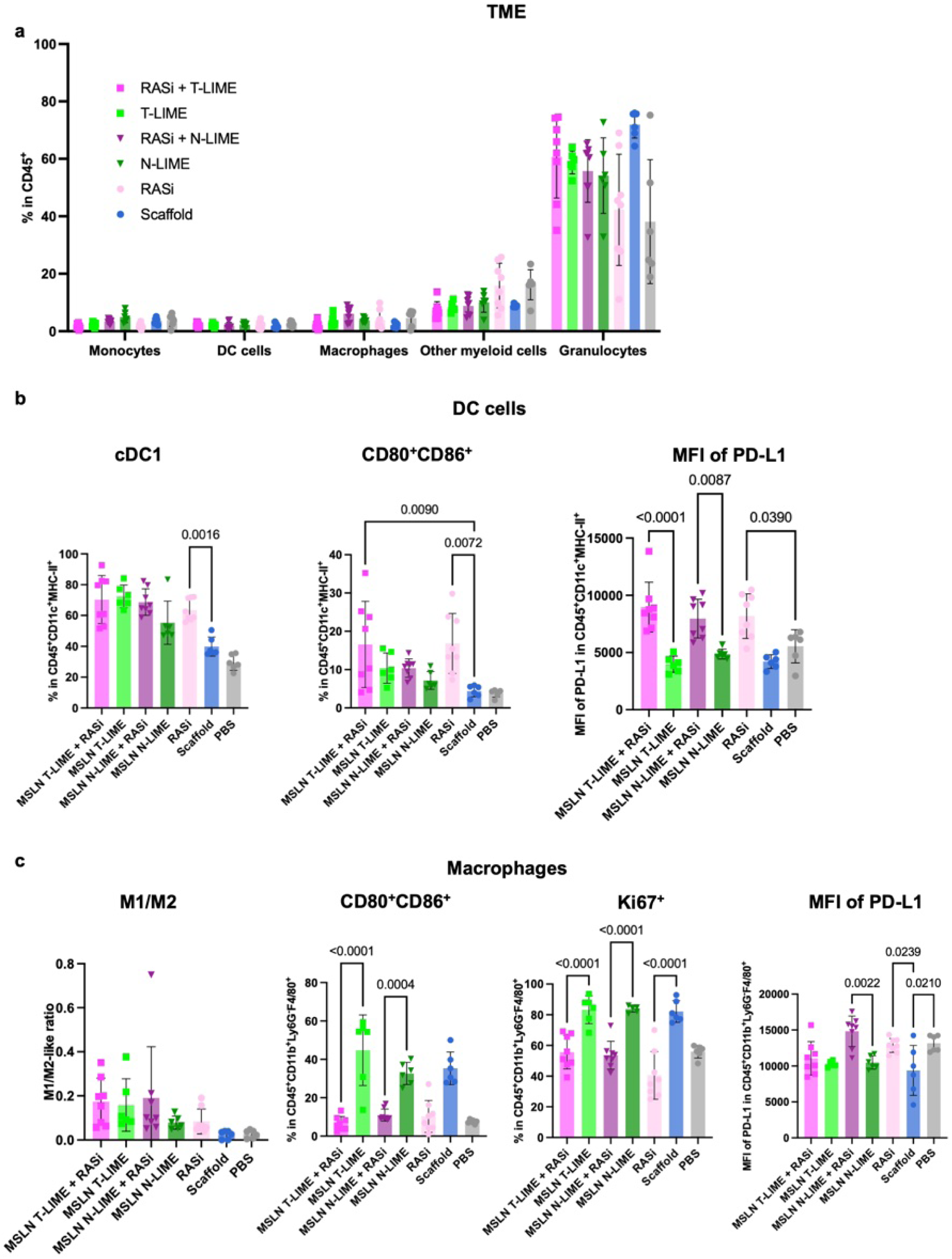
Myeloid-cell remodeling in the tumor microenvironment following LIME and RASi treatment. **a**, Frequencies of monocytes, dendritic cells (DCs), macrophages, other myeloid cells and granulocytes among CD45^+^ cells in the TME. **b**, Frequencies of cDC1 and CD80^+^CD86^+^ DCs and PD-L1 expression represented by median fluorescence intensity (MFI) on DCs. **c**, Macrophage M1/M2-like ratio and frequencies of CD80^+^CD86^+^ and Ki67+ macrophages, with PD-L1 expression on macrophages. Statistical significance was determined using one-way ANOVA with Tukey’s post hoc test (**b, c**). Data are presented as mean ± s.d. (**a**-**c**).

**Supplemental Fig. 62.**
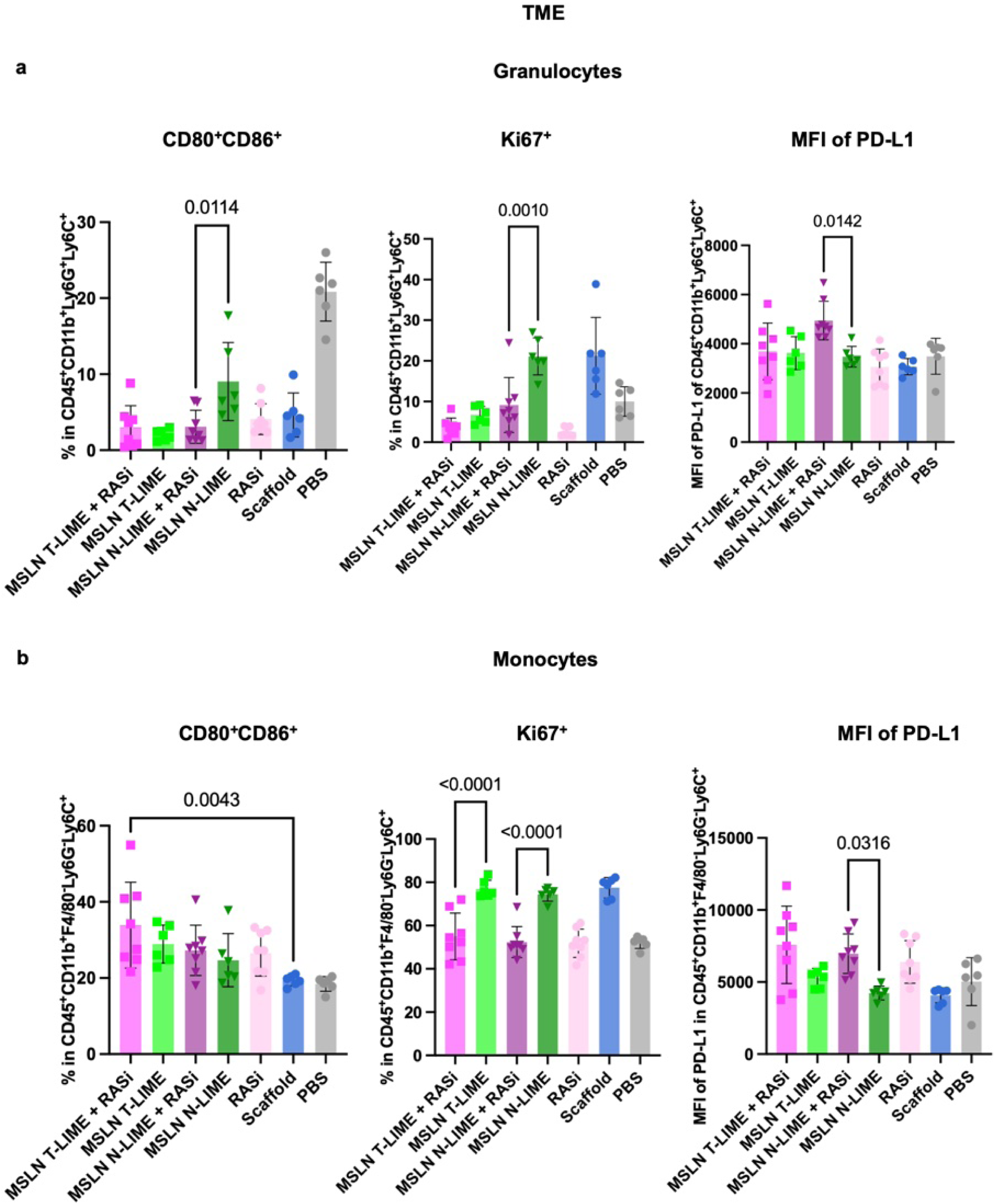
Granulocyte and monocyte phenotypes in the tumor microenvironment following LIME and RASi treatment. **a**, Frequencies of CD80^+^CD86^+^, Ki67^+^ and PD-L1^+^ granulocytes in the TME. **b**, Frequencies of CD80^+^CD86^+^, Ki67^+^ and PD-L1^+^ monocytes in the TME. Statistical significance was determined using one-way ANOVA with Tukey’s post hoc test (**a**, **b**). Data are presented as mean ± s.d. (**a**, **b**).

**Supplemental Fig. 63.**
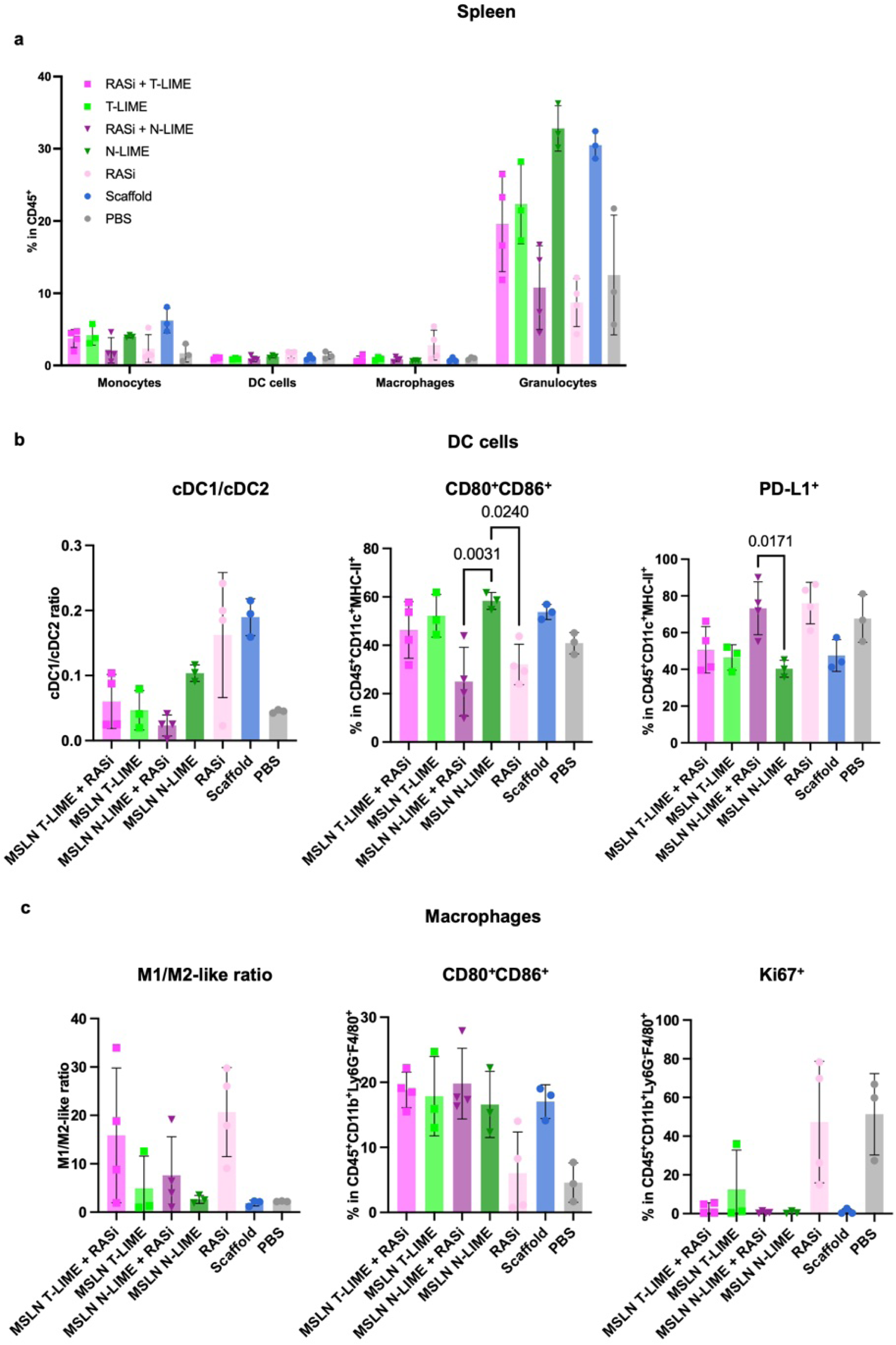
Splenic dendritic-cell and macrophage phenotypes following LIME and RASi treatment. **a**, Ratio of cDC1 to cDC2 cells and frequencies of DC cells, CD80^+^CD86^+^ DCs and PD-L1^+^ DCs in the spleen. **b**, Frequencies of CD80^+^CD86^+^ and Ki67^+^ macrophages. **c**, M1/M2-like macrophage ratio in the spleen. Statistical significance was determined using one-way ANOVA with Tukey’s post hoc test (**b**). Data are presented as mean ± s.d. (**a**-**c**).

**Supplemental Fig. 64.**
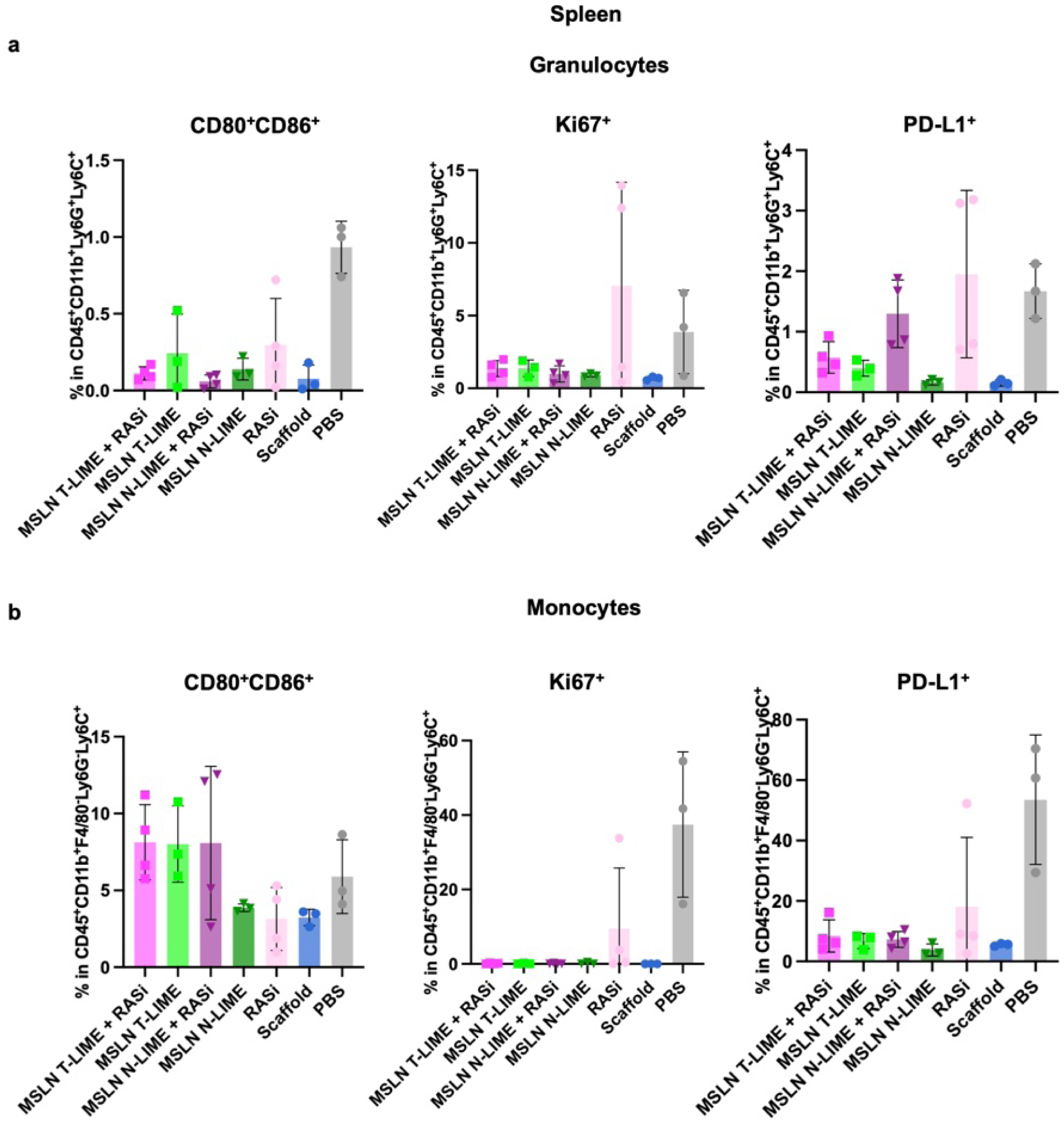
Splenic granulocyte and monocyte phenotypes following LIME and RASi treatment. **a**, Frequencies of CD80^+^CD86^+^, Ki67^+^ and PD-L1^+^ granulocytes in the spleen. **b**, Frequencies of CD80^+^CD86^+^, Ki67^+^ and PD-L1^+^ monocytes in the spleen. Data are presented as mean ± s.d. (**a**, **b**).

**Supplemental Fig. 65.**
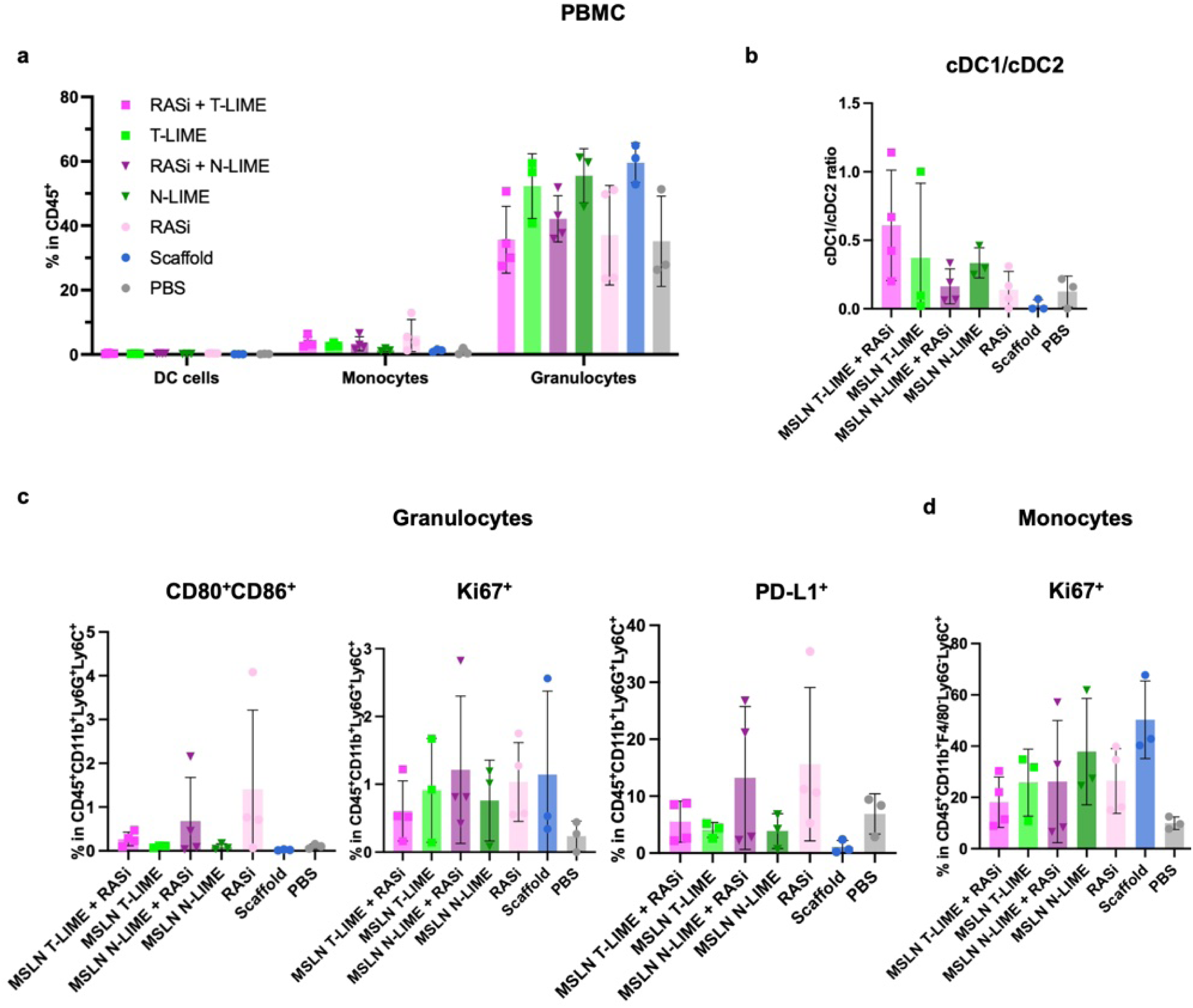
Peripheral blood myeloid-cell phenotypes following LIME and RASi treatment. **a**, Frequencies of myeloid-cell subsets among CD45^+^ PBMCs. **b**, cDC1/cDC2 ratio in PBMCs. **c**, Frequencies of CD80^+^CD86^+^, Ki67^+^ and PD-L1^+^ granulocytes. **d**, Frequencies of Ki67^+^ monocytes in PBMCs. Data are presented as mean ± s.d. (**a**-**d**).

**Supplemental Fig. 66.**
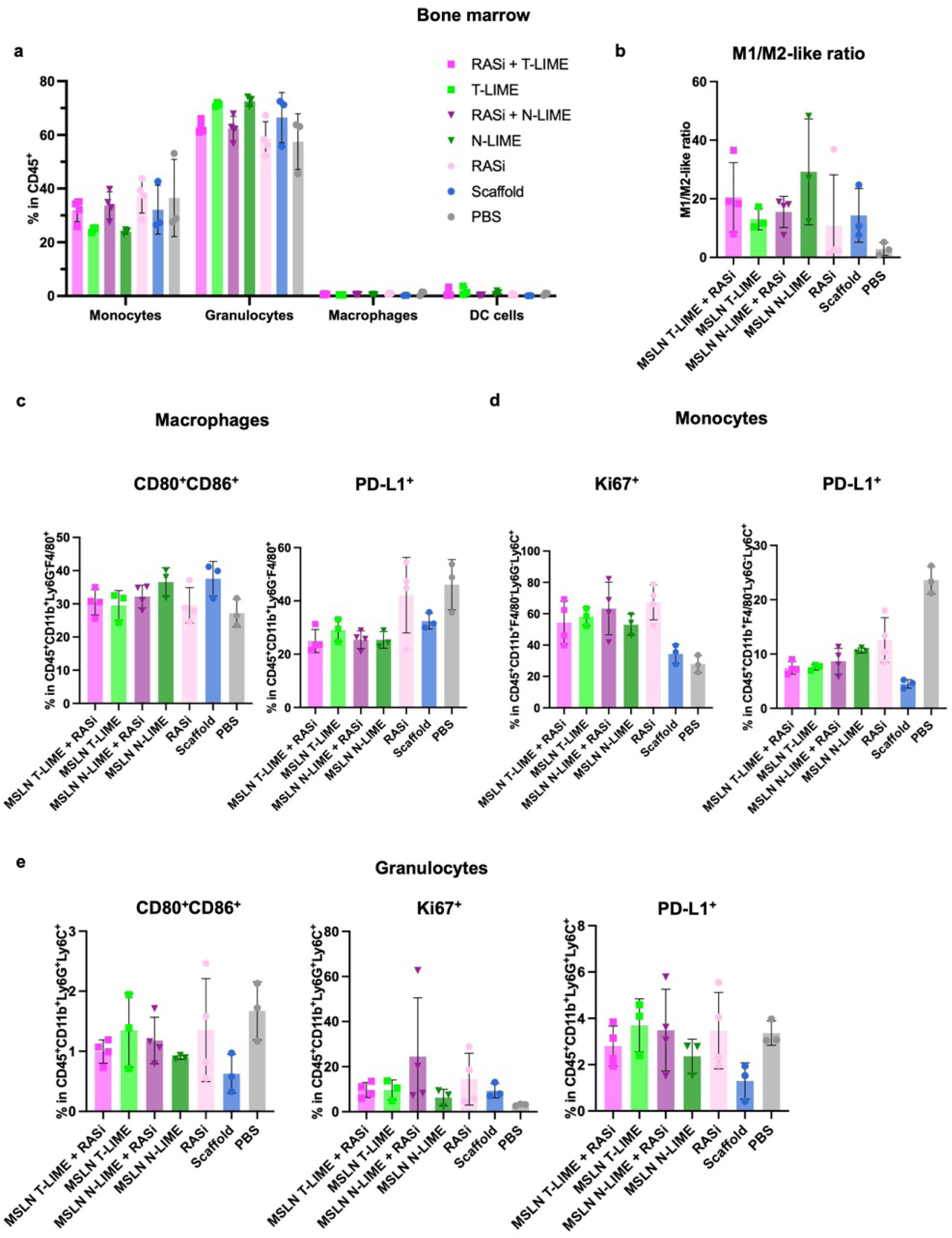
Bone marrow monocyte, macrophage and granulocyte phenotypes following LIME and RASi treatment. **a**, Frequencies of monocytes, macrophages and granulocytes among CD45^+^ bone marrow cells. **b**, M1/M2-like macrophage ratio. **c**, Frequencies of CD80^+^CD86^+^ and PD-L1^+^ monocytes. **d**, Frequencies of PD-L1^+^ and Ki67^+^ macrophages. **e**, Frequencies of CD80^+^CD86^+^, Ki67^+^ and PD-L1^+^ granulocytes. Data are presented as mean ± s.d. (**a**-**e**).

**Supplemental Fig. 67.**
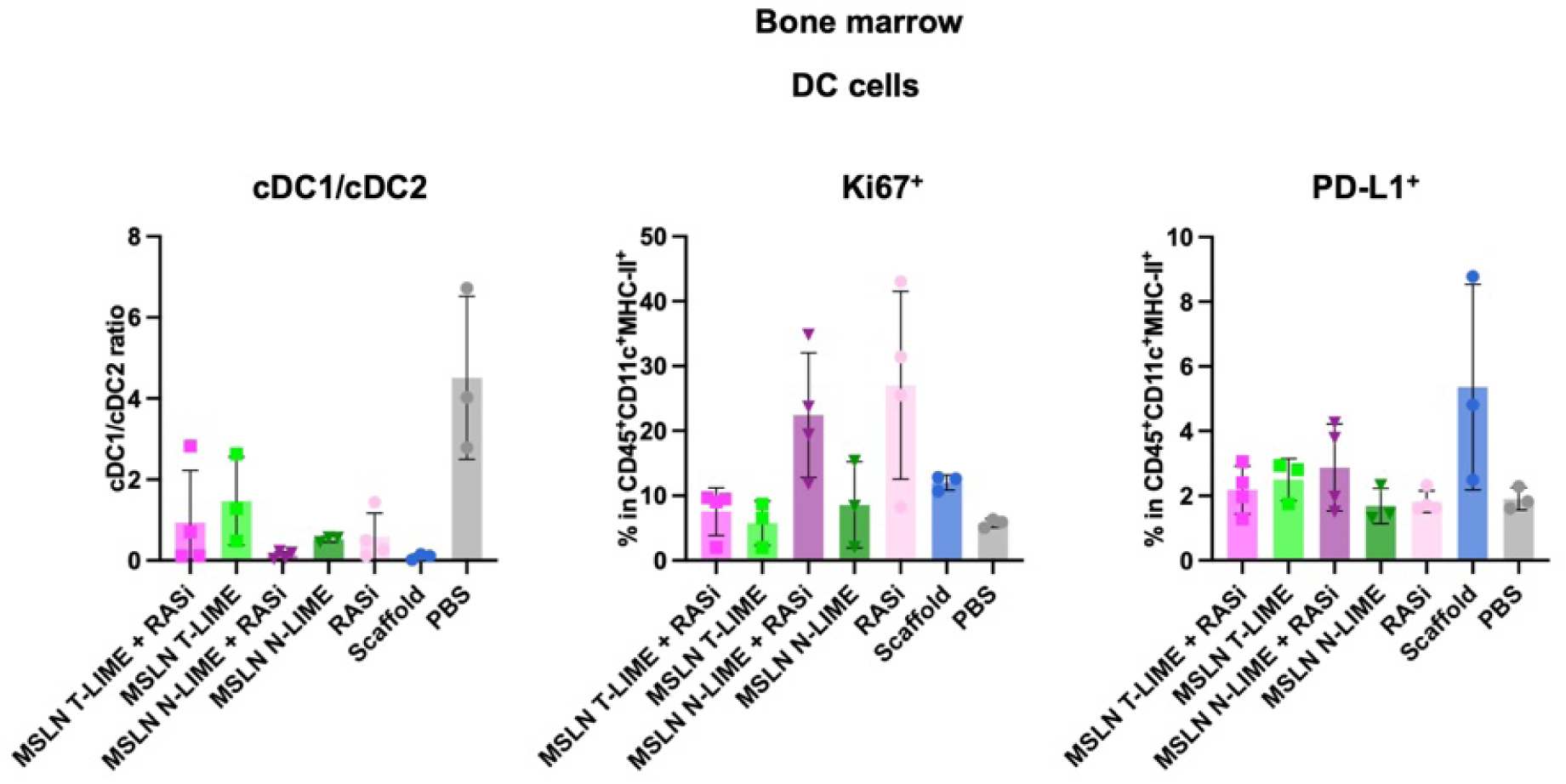
Bone marrow dendritic-cell phenotypes following LIME and RASi treatment. Frequencies of DC cells, cDC1/cDC2 ratio and frequencies of Ki67^+^ and PD-L1^+^ DCs in bone marrow. Data are presented as mean ± s.d.

**Supplemental Fig. 68.**
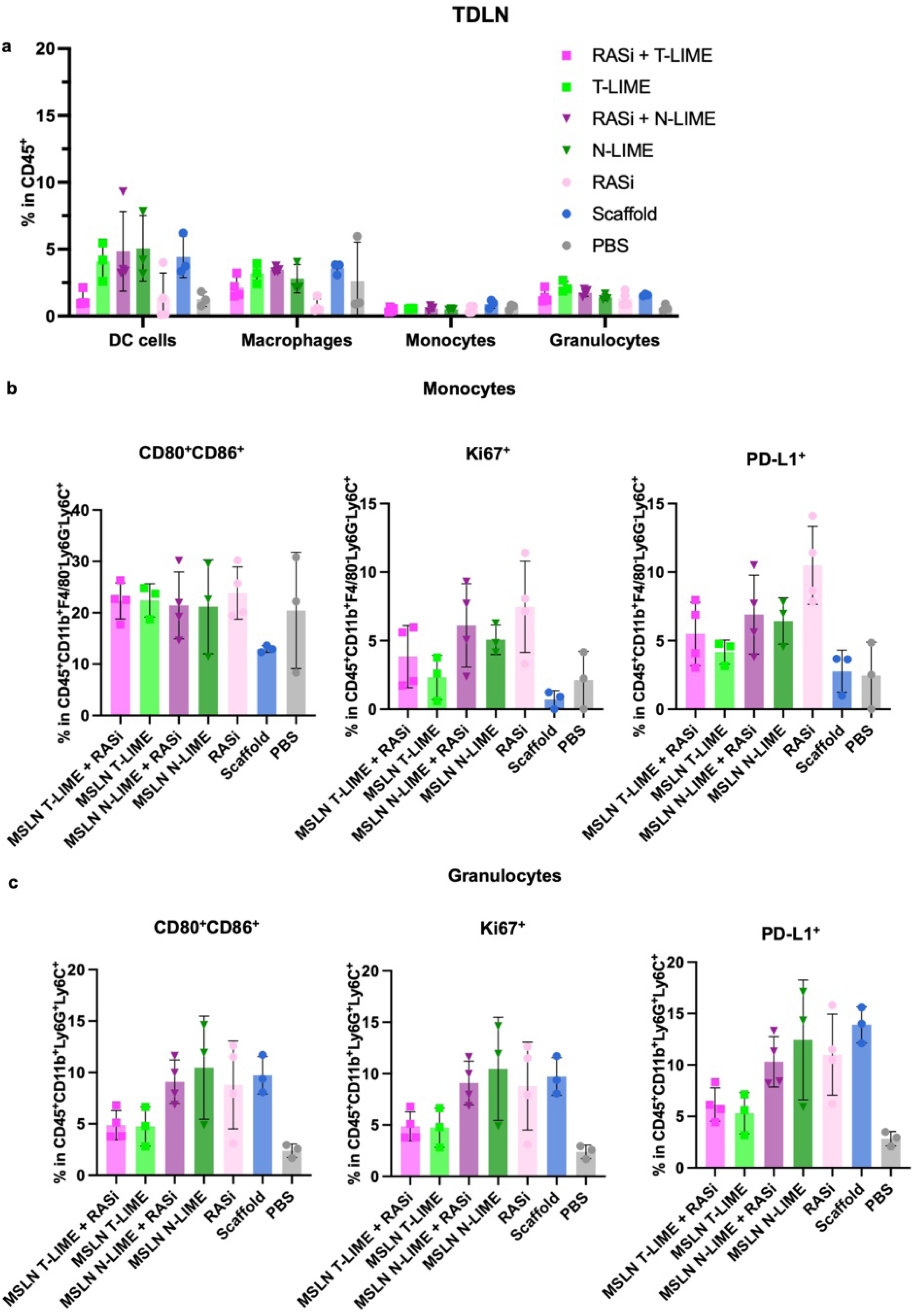
Tumor-draining lymph node granulocyte and monocyte phenotypes following LIME and RASi treatment. **a**, Frequencies of CD80^+^CD86^+^, Ki67^+^ and PD-L1^+^ granulocytes in the TDLN. **b**, Frequencies of CD80^+^CD86^+^, Ki67^+^ and PD-L1^+^ monocytes in the TDLN. **c**, Frequencies of granulocytes and monocytes among CD45^+^ cells in the TDLN. Data are presented as mean ± s.d. (**a**-**c**).

**Supplemental Fig. 69.**
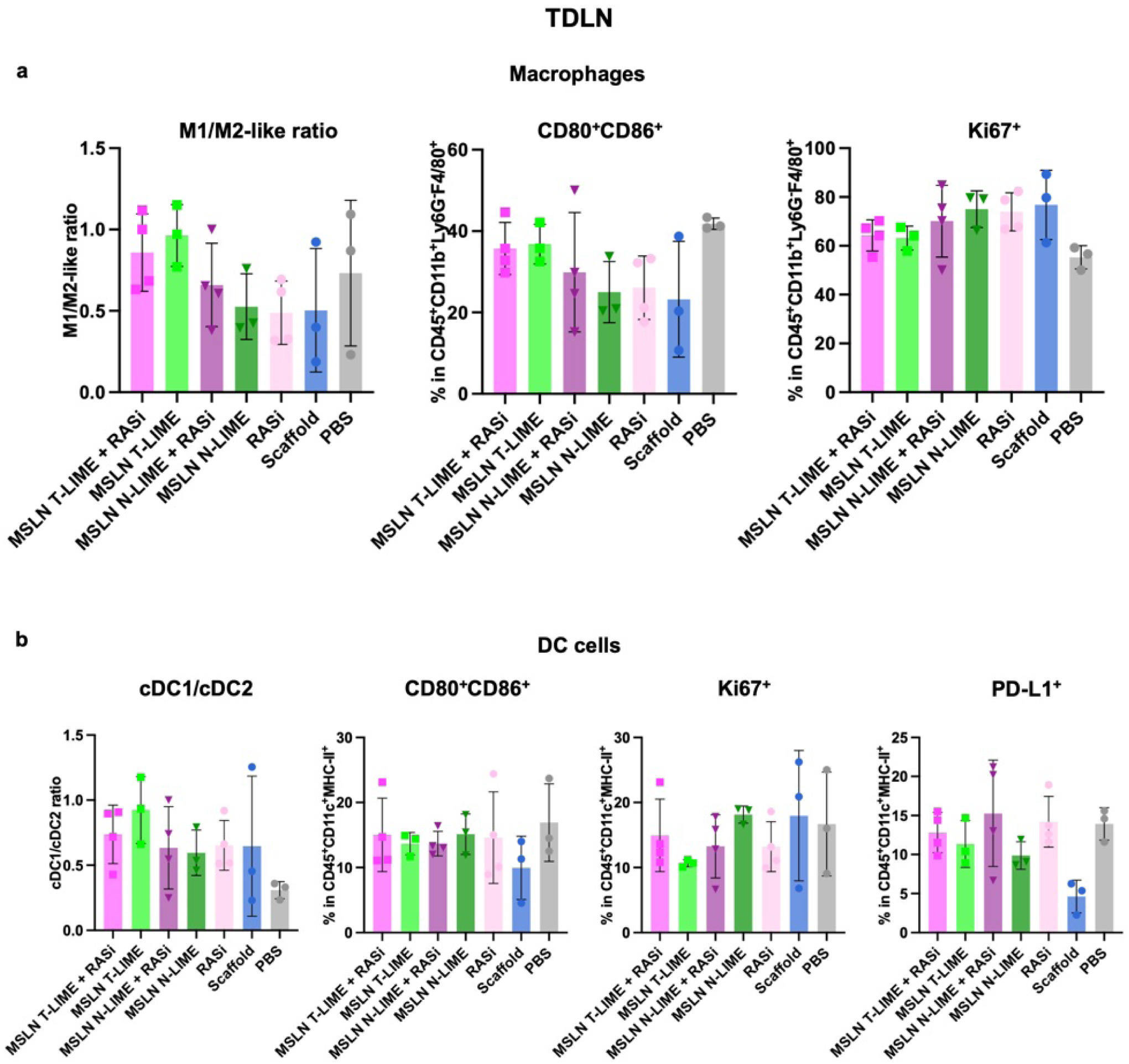
Tumor-draining lymph node macrophage and dendritic-cell phenotypes following LIME and RASi treatment. **a**, Frequencies of CD80^+^CD86^+^ and Ki67^+^ macrophages and M1/M2-like macrophage ratio in the TDLN. **b**, Frequencies of DC cells, cDC1/cDC2 ratio and frequencies of CD80^+^CD86^+^, Ki67^+^ and PD-L1^+^ DCs in the TDLN. Data are presented as mean ± s.d. (**a**, **b**).

**Supplemental Fig. 70.**
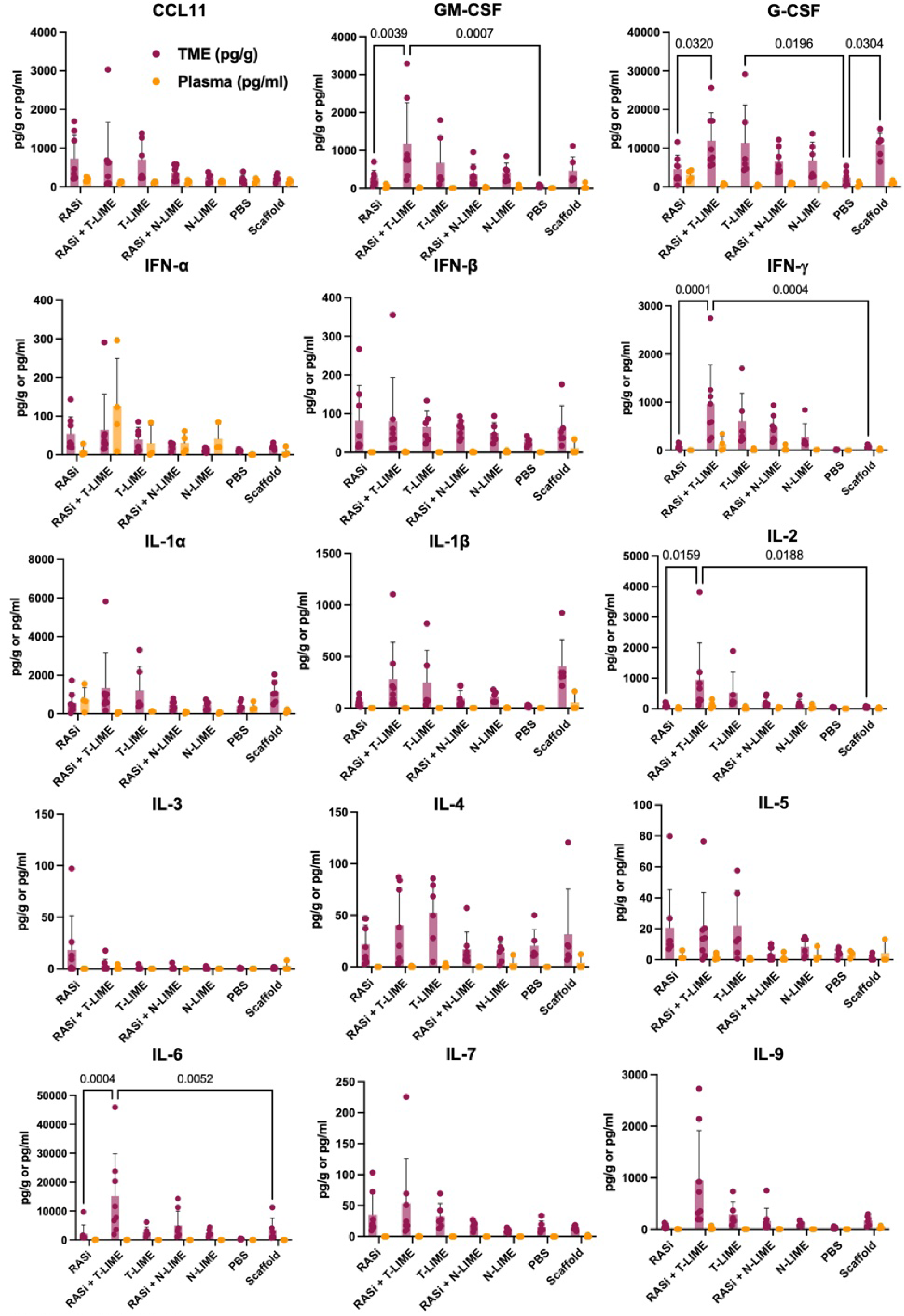
Soluble immune-factor profiling in tumor tissue and plasma following LIME and RASi treatment. Luminex-based quantification of CCL11, GM-CSF, G-CSF, IFN-α, IFN-β, IFN-γ, IL-1α, IL-1β, IL-2, IL-3, IL-4, IL-5, IL-6, IL-7 and IL-9 in tumor lysates and plasma collected from KPC C2 tumor-bearing mice after the indicated treatments. Data are presented as mean ± s.d.

**Supplemental Fig. 71.**
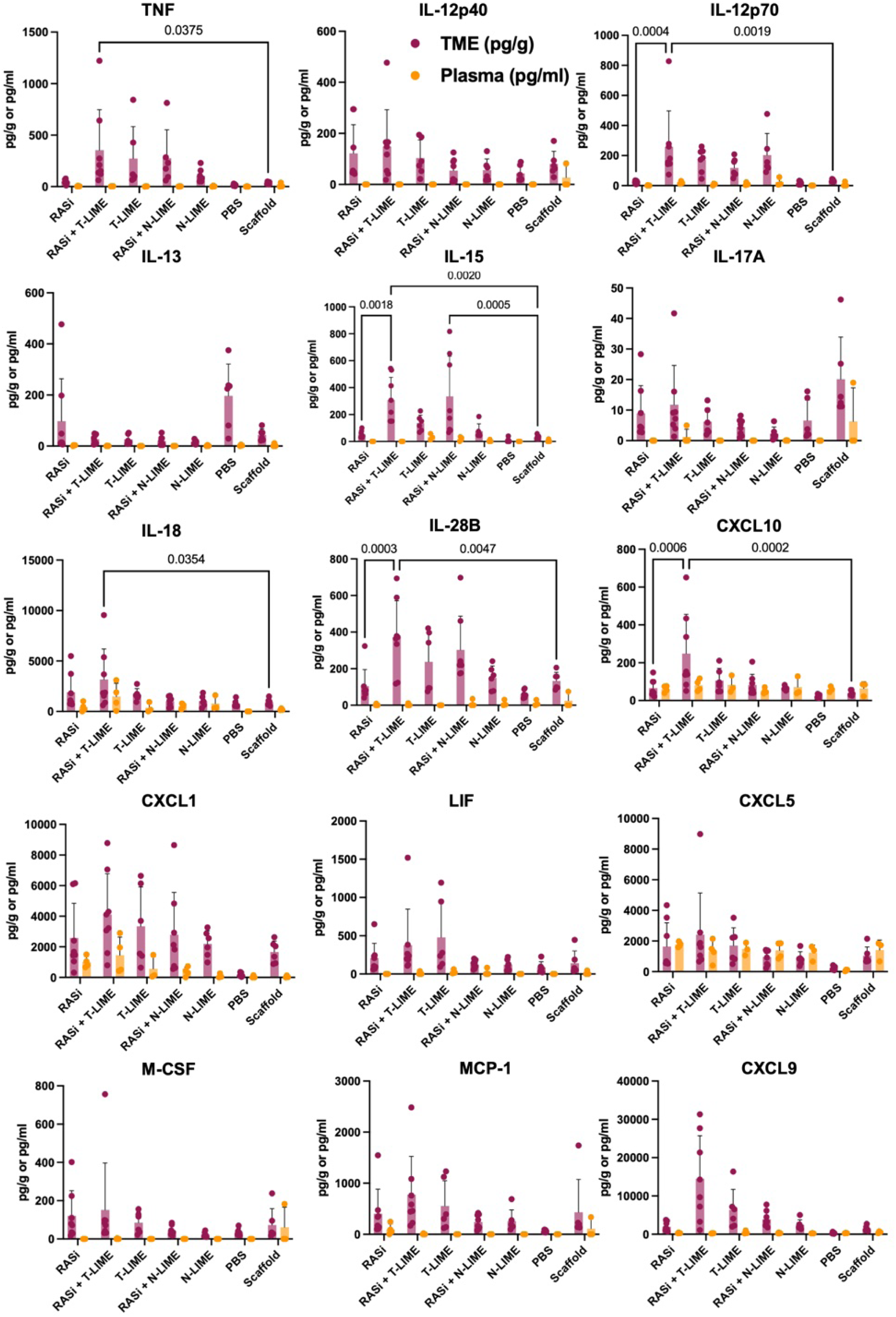
Extended cytokine and chemokine profiling in tumor tissue and plasma following LIME and RASi treatment. Luminex-based quantification of TNF, IL-12p40, IL-12p70, IL-13, IL-15, IL-17A, IL-18, IL-28B, CXCL-10, CXCL-1, LIF, CXCL-5, M-CSF, MCP-1 and CXCL-9 in tumor lysates and plasma collected from KPC C2 tumor-bearing mice after the indicated treatments. Data are presented as mean ± s.d.

**Supplemental Fig. 72.**
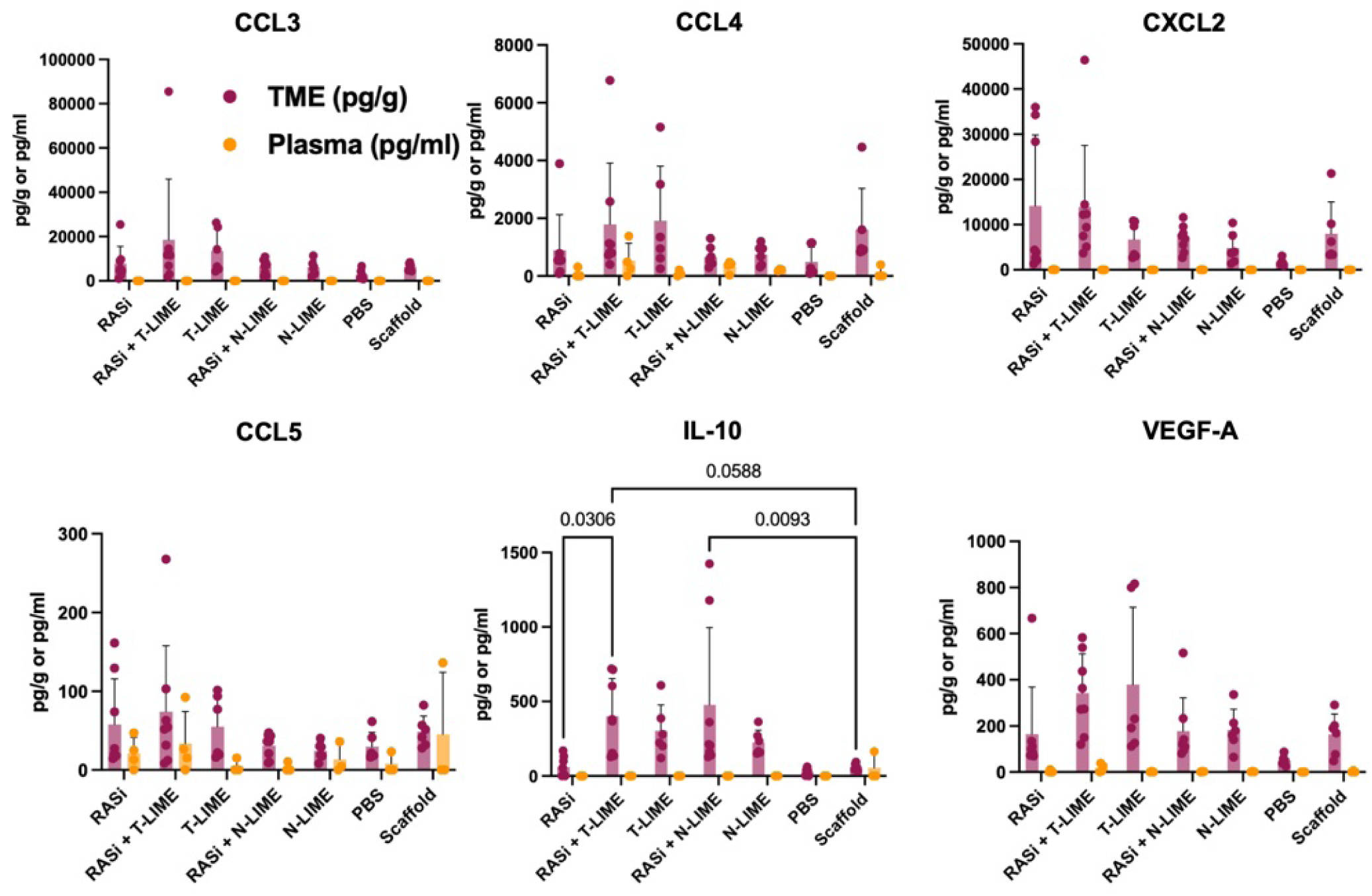
Additional soluble immune-factor profiling in tumor tissue and plasma following LIME and RASi treatment. Luminex-based quantification of CCL-3, CCL-4, CXCL-2, CCL-5, IL-10 and VEGF-A in tumor lysates and plasma collected from KPC C2 tumor-bearing mice after the indicated treatments. Data are presented as mean ± s.d.

## Reference

1. Fenis A, Demaria O, Gauthier L, Vivier E, Narni-Mancinelli E. New immune cell engagers for cancer immunotherapy. Nat Rev Immunol. 2024;24(7):471–86. Epub 20240125. doi: 10.1038/s41577-023-00982-7. PubMed PMID: 38273127.

2. Butterfield LH, Najjar YG. Immunotherapy combination approaches: mechanisms, biomarkers and clinical observations. Nat Rev Immunol. 2024;24(6):399–416. Epub 20231206. doi: 10.1038/s41577-023-00973-8. PubMed PMID: 38057451; PMCID: PMC11460566.

3. Blinatumomab (Blincyto): Indication: For the treatment of patients with Philadelphia chromosome-negative, CD19-positive B-cell precursor acute lymphoblastic leukemia in the consolidation phase of multiphase chemotherapy: Reimbursement Recommendation. Ottawa (ON) 2025.

4. Portuguese AJ, Davis JA, Raza S, Castaneda Puglianini O, Freeman C, Alsina M, Wright C, Blue BJ, Baz RC, Shain KH, Goel U, Khouri J, Anwer F, Ali HM, Pasvolsky O, Gaballa M, Patel K, Garcia-Pleitez H, Mikulski D, Fogel L, Zolotov E, Sannareddy A, Afrough A, Anderson LD, Jr., Julian K, Dima D, Banerjee R, Liang EC, Cassano Cassano R, Herr MM, Hassan H, Shune L, Kort J, Midha S, Nadeem O, Sidana S, Lin Y, Locke FL, Hansen DK, Sborov D, Biran N, Grajales-Cruz A. Real-world outcomes with elranatamab in multiple myeloma: a multicenter analysis from the U.S. Multiple Myeloma Immunotherapy Consortium. Blood Cancer J. 2026;16(1). Epub 20260327. doi: 10.1038/s41408-026-01477-z. PubMed PMID: 41896552; PMCID: PMC13039266.

5. Nathan P, Hassel JC, Rutkowski P, Baurain JF, Butler MO, Schlaak M, Sullivan RJ, Ochsenreither S, Dummer R, Kirkwood JM, Joshua AM, Sacco JJ, Shoushtari AN, Orloff M, Piulats JM, Milhem M, Salama AKS, Curti B, Demidov L, Gastaud L, Mauch C, Yushak M, Carvajal RD, Hamid O, Abdullah SE, Holland C, Goodall H, Piperno-Neumann S, Investigators IM-. Overall Survival Benefit with Tebentafusp in Metastatic Uveal Melanoma. N Engl J Med. 2021;385(13):1196–206. doi: 10.1056/NEJMoa2103485. PubMed PMID: 34551229.

6. Carlisle JW, O’Reilly K, Ji XA, Switchenko JM, Steuer CE, Ardeshir-Larijani F, Ramalingam SS, Leal T. Real-World Implementation of Tarlatamab-Dlle Therapy for Patients With Extensive-Stage Small Cell Lung Cancer and Other High-Grade Neuroendocrine Neoplasms. JCO Oncol Pract. 2026:OP2600074. Epub 20260708. doi: 10.1200/OP-26-00074. PubMed PMID: 42418753.

7. Goyal PK, Sangwan K. Tarlatamab-dlle: A New Hope for Patients with Extensive-Stage Small-Cell Lung Cancer. Curr Treat Options Oncol. 2024;25(11):1337–44. Epub 20241011. doi: 10.1007/s11864-024-01268-3. PubMed PMID: 39392556.

8. Cao L, Leclercq-Cohen G, Klein C, Sorrentino A, Bacac M. Mechanistic insights into resistance mechanisms to T cell engagers. Front Immunol. 2025;16:1583044. Epub 20250422. doi: 10.3389/fimmu.2025.1583044. PubMed PMID: 40330489; PMCID: PMC12053166.

9. Lucas AT, Moody A, Schorzman AN, Zamboni WC. Importance and Considerations of Antibody Engineering in Antibody-Drug Conjugates Development from a Clinical Pharmacologist’s Perspective. Antibodies (Basel). 2021;10(3). Epub 20210726. doi: 10.3390/antib10030030. PubMed PMID: 34449544; PMCID: PMC8395454.

10. Habibi MA, Nejati N, Najafi MB, Khodadadiyan A, Dashti M, Lorestani P, Karimizadeh Z, Ahmadpour M, Kalantari A, Jokar-Derisi A, Maghsood F, Robat-Jazi B, Ebrahimi E, Ahmadpour S, Tavakolpour S. Enhancing T cell infiltration in glioblastoma: a review article on challenges and therapeutic strategies. Cancer Treat Res Commun. 2025;45:100999. Epub 20250909. doi: 10.1016/j.ctarc.2025.100999. PubMed PMID: 40961824.

11. Lu G, Hickey JW, Haist M, Qin X, Zhao E, Naveed A, Forgo E, Baertsch MA, Mani L, Rovira-Clave X, Finegersh A, Goltsev Y, Caraccio C, van den Berg NS, Hom M, Colburg DR, Martin BA, Kong CS, Lui NS, Fisher GA, Colevas AD, West RB, Thurber GM, Poultsides GA, Nolan GP, Rosenthal EL. Single-cell spatial pharmacobiology identifies conserved stromal barriers to therapeutic antibody delivery in human solid tumors. Nat Biotechnol. 2026. Epub 20260603. doi: 10.1038/s41587-026-03152-x. PubMed PMID: 42236958; PMCID: PMC13390730.

12. Dewaele L, Fernandes RA. Bispecific T-cell engagers for the recruitment of T cells in solid tumors: a literature review. Immunother Adv. 2025;5(1):ltae005. Epub 20250127. doi: 10.1093/immadv/ltae005. PubMed PMID: 40083373; PMCID: PMC11904783.

13. Ahn MJ, Cho BC, Felip E, Korantzis I, Ohashi K, Majem M, Juan-Vidal O, Handzhiev S, Izumi H, Lee JS, Dziadziuszko R, Wolf J, Blackhall F, Reck M, Bustamante Alvarez J, Hummel HD, Dingemans AC, Sands J, Akamatsu H, Owonikoko TK, Ramalingam SS, Borghaei H, Johnson ML, Huang S, Mukherjee S, Minocha M, Jiang T, Martinez P, Anderson ES, Paz-Ares L, De L-I. Tarlatamab for Patients with Previously Treated Small-Cell Lung Cancer. N Engl J Med. 2023;389(22):2063–75. Epub 20231020. doi: 10.1056/NEJMoa2307980. PubMed PMID: 37861218.

14. Portell CA, Wenzell CM, Advani AS. Clinical and pharmacologic aspects of blinatumomab in the treatment of B-cell acute lymphoblastic leukemia. Clin Pharmacol. 2013;5(Suppl 1):5–11. Epub 20130412. doi: 10.2147/CPAA.S42689. PubMed PMID: 23671399; PMCID: PMC3650887.

15. Amash A, Volkers G, Farber P, Griffin D, Davison KS, Goodman A, Tonikian R, Yamniuk A, Barnhart B, Jacobs T. Developability considerations for bispecific and multispecific antibodies. MAbs. 2024;16(1):2394229. Epub 20240827. doi: 10.1080/19420862.2024.2394229. PubMed PMID: 39189686; PMCID: PMC11352713.

16. Westphal K, Leschner S, Jablonska J, Loessner H, Weiss S. Containment of tumor-colonizing bacteria by host neutrophils. Cancer Res. 2008;68(8):2952–60. doi: 10.1158/0008-5472.CAN-07-2984. PubMed PMID: 18413765.

17. Zheng LM, Luo X, Feng M, Li Z, Le T, Ittensohn M, Trailsmith M, Bermudes D, Lin SL, King IC. Tumor amplified protein expression therapy: Salmonella as a tumor-selective protein delivery vector. Oncol Res. 2000;12(3):127–35. doi: 10.3727/096504001108747602. PubMed PMID: 11216671.

18. St Jean AT, Zhang M, Forbes NS. Bacterial therapies: completing the cancer treatment toolbox. Curr Opin Biotechnol. 2008;19(5):511–7. Epub 20080918. doi: 10.1016/j.copbio.2008.08.004. PubMed PMID: 18760353; PMCID: PMC2600537.

19. Toso JF, Gill VJ, Hwu P, Marincola FM, Restifo NP, Schwartzentruber DJ, Sherry RM, Topalian SL, Yang JC, Stock F, Freezer LJ, Morton KE, Seipp C, Haworth L, Mavroukakis S, White D, MacDonald S, Mao J, Sznol M, Rosenberg SA. Phase I study of the intravenous administration of attenuated Salmonella typhimurium to patients with metastatic melanoma. J Clin Oncol. 2002;20(1):142–52. doi: 10.1200/JCO.2002.20.1.142. PubMed PMID: 11773163; PMCID: PMC2064865.

20. Janku F, Zhang HH, Pezeshki A, Goel S, Murthy R, Wang-Gillam A, Shepard DR, Helgason T, Masters T, Hong DS, Piha-Paul SA, Karp DD, Klang M, Huang SY, Sakamuri D, Raina A, Torrisi J, Solomon SB, Weissfeld A, Trevino E, DeCrescenzo G, Collins A, Miller M, Salstrom JL, Korn RL, Zhang L, Saha S, Leontovich AA, Tung D, Kreider B, Varterasian M, Khazaie K, Gounder MM. Intratumoral Injection of Clostridium novyi-NT Spores in Patients with Treatment-refractory Advanced Solid Tumors. Clin Cancer Res. 2021;27(1):96–106. Epub 20201012. doi: 10.1158/1078-0432.CCR-20-2065. PubMed PMID: 33046513.

21. Gniadek TJ, Augustin L, Schottel J, Leonard A, Saltzman D, Greeno E, Batist G. A Phase I, Dose Escalation, Single Dose Trial of Oral Attenuated Salmonella typhimurium Containing Human IL-2 in Patients With Metastatic Gastrointestinal Cancers. J Immunother. 2020;43(7):217–21. doi: 10.1097/CJI.0000000000000325. PubMed PMID: 32554977; PMCID: PMC7458080.

22. Batist G, Kavan P, Augustin L, Schottel J, Moradian J, Lee JT, Saltzman D. A phase 2 study of orally administered live biotherapeutic salmonella-IL2 with FOLFIRINOX for stage IV pancreatic cancer. Cancer Immunol Immunother. 2026;75(2):60. Epub 20260127. doi: 10.1007/s00262-026-04307-0. PubMed PMID: 41591505; PMCID: PMC12847481.

23. Luke JJ, Piha-Paul SA, Medina T, Verschraegen CF, Varterasian M, Brennan AM, Riese RJ, Sokolovska A, Strauss J, Hava DL, Janku F. Phase I Study of SYNB1891, an Engineered E. coli Nissle Strain Expressing STING Agonist, with and without Atezolizumab in Advanced Malignancies. Clin Cancer Res. 2023;29(13):2435–44. doi: 10.1158/1078-0432.CCR-23-0118. PubMed PMID: 37227176; PMCID: PMC11225568.

24. Bryant FR. Construction of a recombinase-deficient mutant recA protein that retains single-stranded DNA-dependent ATPase activity. J Biol Chem. 1988;263(18):8716–23. PubMed PMID: 2967815.

25. Scudamore RA, Beveridge TJ, Goldner M. Outer-membrane penetration barriers as components of intrinsic resistance to beta-lactam and other antibiotics in Escherichia coli K-12. Antimicrob Agents Chemother. 1979;15(2):182–9. doi: 10.1128/AAC.15.2.182. PubMed PMID: 106773; PMCID: PMC352630.

26. Stritzker J, Weibel S, Hill PJ, Oelschlaeger TA, Goebel W, Szalay AA. Tumor-specific colonization, tissue distribution, and gene induction by probiotic Escherichia coli Nissle 1917 in live mice. Int J Med Microbiol. 2007;297(3):151–62. Epub 20070419. doi: 10.1016/j.ijmm.2007.01.008. PubMed PMID: 17448724.

27. Yang S, Sheffer M, Kaplan IE, Wang Z, Tarannum M, Dinh K, Abdulhamid Y, Bobilev E, Shapiro R, Porter R, Soiffer R, Ritz J, Koreth J, Wei Y, Chen P, Zhang K, Marquez-Pellegrin V, Bonanno S, Joshi N, Guan M, Yang M, Li D, Bellini C, Liu F, Chen J, Wu CJ, Barbie D, Li J, Romee R. Non-pathogenic E. coli displaying decoy-resistant IL18 mutein boosts anti-tumor and CAR NK cell responses. Nat Biotechnol. 2025;43(8):1311–23. Epub 20241004. doi: 10.1038/s41587-024-02418-6. PubMed PMID: 39367093; PMCID: PMC12797303.

28. Yang S, Wang Z, Fang C, Yang M, Khawaled S, Bonanno S, Joshi NS, Wei Y, Zhang K, Marquez-Pellegrin V, Guan M, Zhang S, Bader AC, Ye N, Haley AE, Dame MK, Spence JR, He X, Fox JG, Yilmaz OH, Shah YM, Romee R, Li J. Surface expression of antitoxin on engineered bacteria neutralizes genotoxic colibactin in the gut. Nat Microbiol. 2026;11(1):53–66. Epub 20251208. doi: 10.1038/s41564-025-02177-3. PubMed PMID: 41361522; PMCID: PMC12833714.

29. Ding Z, Sun S, Yang X, Huang X, Hou X, Xie S, Liu A, Lu X. TCR-mimic bispecific nanobody-based T cell engager targeting intracellular tumor antigens for cancer immunotherapy. Signal Transduct Target Ther. 2026;11(1). Epub 20260625. doi: 10.1038/s41392-026-02745-x. PubMed PMID: 42342658; PMCID: PMC13294373.

30. Cao L, Coventry B, Goreshnik I, Huang B, Sheffler W, Park JS, Jude KM, Markovic I, Kadam RU, Verschueren KHG, Verstraete K, Walsh STR, Bennett N, Phal A, Yang A, Kozodoy L, DeWitt M, Picton L, Miller L, Strauch EM, DeBouver ND, Pires A, Bera AK, Halabiya S, Hammerson B, Yang W, Bernard S, Stewart L, Wilson IA, Ruohola-Baker H, Schlessinger J, Lee S, Savvides SN, Garcia KC, Baker D. Design of protein-binding proteins from the target structure alone. Nature. 2022;605(7910):551–60. Epub 20220324. doi: 10.1038/s41586-022-04654-9. PubMed PMID: 35332283; PMCID: PMC9117152.

31. Sauer T, Parikh K, Sharma S, Omer B, Sedloev D, Chen Q, Angenendt L, Schliemann C, Schmitt M, Muller-Tidow C, Gottschalk S, Rooney CM. CD70-specific CAR T cells have potent activity against acute myeloid leukemia without HSC toxicity. Blood. 2021;138(4):318–30. doi: 10.1182/blood.2020008221. PubMed PMID: 34323938; PMCID: PMC8323977.

32. Albayrak G, Wan PK, Fisher K, Seymour LW. T cell engagers: expanding horizons in oncology and beyond. Br J Cancer. 2025;133(9):1241–9. Epub 20250723. doi: 10.1038/s41416-025-03125-y. PubMed PMID: 40702106; PMCID: PMC12572156.

33. Ahn MJ, Cho BC, Ohashi K, Izumi H, Lee JS, Han JY, Chiang CL, Huang S, Hamidi A, Mukherjee S, Xu KL, Akamatsu H. Asian Subgroup Analysis of Patients in the Phase 2 DeLLphi-301 Study of Tarlatamab for Previously Treated Small Cell Lung Cancer. Oncol Ther. 2025;13(4):1041–54. Epub 20250904. doi: 10.1007/s40487-025-00372-0. PubMed PMID: 40908346; PMCID: PMC12647424.

34. Aladin F, Lautscham G, Humphries E, Coulson J, Blake N. Targeting tumour cells with defects in the MHC Class I antigen processing pathway with CD8+ T cells specific for hydrophobic TAP- and Tapasin-independent peptides: the requirement for directed access into the ER. Cancer Immunol Immunother. 2007;56(8):1143–52. Epub 20061202. doi: 10.1007/s00262-006-0263-2. PubMed PMID: 17143611; PMCID: PMC11031051.

35. Kyrysyuk O, Wucherpfennig KW. Designing Cancer Immunotherapies That Engage T Cells and NK Cells. Annu Rev Immunol. 2023;41:17–38. Epub 20221129. doi: 10.1146/annurev-immunol-101921-044122. PubMed PMID: 36446137; PMCID: PMC10159905.

36. Spinazzola A, Iannantuono GM, Gulley JL, Giudice E, Filetti M, Sganga S, Bianco FL, Floudas CS, Daniele G. Current landscape of T-cell engagers in early-phase clinical development in solid cancers. Front Immunol. 2025;16:1665838. Epub 20251006. doi: 10.3389/fimmu.2025.1665838. PubMed PMID: 41122165; PMCID: PMC12536262.

37. Herault A, Mak J, de la Cruz-Chuh J, Dillon MA, Ellerman D, Go M, Cosino E, Clark R, Carson E, Yeung S, Pichery M, Gador M, Chiang EY, Wu J, Liang Y, Modrusan Z, Gampa G, Sudhamsu J, Kemball CC, Cheung V, Nguyen TTT, Seshasayee D, Piskol R, Totpal K, Yu SF, Lee G, Kozak KR, Spiess C, Walsh KB. NKG2D-bispecific enhances NK and CD8+ T cell antitumor immunity. Cancer Immunol Immunother. 2024;73(10):209. Epub 20240808. doi: 10.1007/s00262-024-03795-2. PubMed PMID: 39112670; PMCID: PMC11306676.

38. Germann M, Zangger N, Sauvain MO, Sempoux C, Bowler AD, Wirapati P, Kandalaft LE, Delorenzi M, Tejpar S, Coukos G, Radtke F. Neutrophils suppress tumor-infiltrating T cells in colon cancer via matrix metalloproteinase-mediated activation of TGFbeta. EMBO Mol Med. 2020;12(1):e10681. Epub 20191202. doi: 10.15252/emmm.201910681. PubMed PMID: 31793740; PMCID: PMC6949488.

39. Wolpin BM, Park W, Garrido-Laguna I, Spira A, Starodub A, Sommerhalder D, Punekar SR, Barve M, Pelster M, Herzberg B, Azad NS, Hecht JR, Ou SHI, Lin T, Kar S, Tao L, Vora R, Hegde A, Aung K, Hong DS, Investigators RMC. Daraxonrasib in Previously Treated Advanced RAS-Mutated Pancreatic Cancer. N Engl J Med. 2026;394(18):1790–802. doi: 10.1056/NEJMoa2505783. PubMed PMID: 42090791.

40. Mahadevan KK, Maldonado AS, Li B, Bickert AA, Kacperczyk-Perdyan A, Kumbhar SV, Piya S, Sockwell AM, Morse SJ, Arian K, Sugimoto H, Shalapour S, Hong DS, Heffernan TP, Maitra A, Kalluri R. Oncogenic Kras targeting with MRTX1133 or Daraxonrasib specifically synergize with anti-CTLA4 to promote anti-tumor immunity in pancreatic cancer. Nature Communications. 2026. doi: 10.1038/s41467-026-75960-3.

41. Vincent RL, Gurbatri CR, Li F, Vardoshvili A, Coker C, Im J, Ballister ER, Rouanne M, Savage T, de los Santos-Alexis K, Redenti A, Brockmann L, Komaranchath M, Arpaia N, Danino T. Probiotic-guided CAR-T cells for solid tumor targeting. Science. 2023;382(6667):211–8. doi: 10.1126/science.add7034.

42. Gurbatri CR, Lia I, Vincent R, Coker C, Castro S, Treuting PM, Hinchliffe TE, Arpaia N, Danino T. Engineered probiotics for local tumor delivery of checkpoint blockade nanobodies. Science Translational Medicine. 2020;12(530). doi: 10.1126/scitranslmed.aax0876.

43. Chowdhury S, Castro S, Coker C, Hinchliffe TE, Arpaia N, Danino T. Programmable bacteria induce durable tumor regression and systemic antitumor immunity. Nature Medicine. 2019;25(7). doi: 10.1038/s41591-019-0498-z.

44. Ballister ER, Michels A, Vincent RL, Kreindler L, Chowdhury S, Upadhaya S, Saez-Ibanez AR, Tu T, Gottweis J, Danino T. The emerging landscape of engineered bacteria cancer therapies. Nat Biotechnol. 2025;43(5):672–6. doi: 10.1038/s41587-025-02623-x. PubMed PMID: 40169920.

45. Wu LY, Qiu JH, Qiao XY, Li L, Qiao LY, Li CY, Sun Y, Zhang SH, Du ZZ, Chang XY, Cheng C, Wang BH, Xiao YH, Lin L, Hua ZC. Octopus-inspired engineered bacteria with a plug-and-play surface display system achieves enhanced tumor-specific colonization and antitumor immunity. Mil Med Res. 2026;13(1):100030. Epub 20260427. doi: 10.1016/j.mmr.2026.100030. PubMed PMID: 42088059; PMCID: PMC13138155.

46. Leithner A, Staufer O, Mitra T, Liberta F, Valvo S, Kutuzov M, Dada H, Spaeth J, Zhou W, Schiele F, Reindl S, Nar H, Hoerer S, Crames M, Comeau S, Young D, Low S, Jenkins E, Davis SJ, Klenerman D, Nixon A, Pefaur N, Wyatt D, Dushek O, Kasturirangan S, Dustin ML. Solution structure and synaptic analyses reveal determinants of bispecific T cell engager potency. Proc Natl Acad Sci U S A. 2025;122(22):e2425781122. Epub 20250530. doi: 10.1073/pnas.2425781122. PubMed PMID: 40445758; PMCID: PMC12146755.

47. Hernandez-Rollan C, Falkenberg KB, Rennig M, Bertelsen AB, Ipsen JO, Brander S, Daley DO, Johansen KS, Norholm MHH. LyGo: A Platform for Rapid Screening of Lytic Polysaccharide Monooxygenase Production. ACS Synth Biol. 2021;10(4):897–906. Epub 20210402. doi: 10.1021/acssynbio.1c00034. PubMed PMID: 33797234.

48. Glass DS, Riedel-Kruse IH. A Synthetic Bacterial Cell-Cell Adhesion Toolbox for Programming Multicellular Morphologies and Patterns. Cell. 2018;174(3):649–58 e16. Epub 20180719. doi: 10.1016/j.cell.2018.06.041. PubMed PMID: 30033369.

49. Han MJ, Lee SH. An efficient bacterial surface display system based on a novel outer membrane anchoring element from the Escherichia coli protein YiaT. FEMS Microbiol Lett. 2015;362(1):1–7. Epub 20141204. doi: 10.1093/femsle/fnu002. PubMed PMID: 25790485.

50. Wierzchowski A, Wink DJ, Zhang H, Kambanis K, Robles JOR, Rosenhouse-Dantsker A. CoLab: A workshop-based undergraduate research experience for entering college students. J Chem Educ. 2022;99(12):4085–93. Epub 20220511. doi: 10.1021/acs.jchemed.1c01290. PubMed PMID: 37519308; PMCID: PMC10373424.

51. Kim D, Paggi JM, Park C, Bennett C, Salzberg SL. Graph-based genome alignment and genotyping with HISAT2 and HISAT-genotype. Nat Biotechnol. 2019;37(8):907–15. Epub 20190802. doi: 10.1038/s41587-019-0201-4. PubMed PMID: 31375807; PMCID: PMC7605509.

52. Liao Y, Smyth GK, Shi W. featureCounts: an efficient general purpose program for assigning sequence reads to genomic features. Bioinformatics. 2014;30(7):923–30. Epub 20131113. doi: 10.1093/bioinformatics/btt656. PubMed PMID: 24227677.

53. Love MI, Huber W, Anders S. Moderated estimation of fold change and dispersion for RNA-seq data with DESeq2. Genome Biol. 2014;15(12):550. doi: 10.1186/s13059-014-0550-8. PubMed PMID: 25516281; PMCID: PMC4302049.

54. Stephens M. False discovery rates: a new deal. Biostatistics. 2017;18(2):275–94. doi: 10.1093/biostatistics/kxw041. PubMed PMID: 27756721; PMCID: PMC5379932.

55. Ritchie ME, Phipson B, Wu D, Hu Y, Law CW, Shi W, Smyth GK. limma powers differential expression analyses for RNA-sequencing and microarray studies. Nucleic Acids Res. 2015;43(7):e47. Epub 20150120. doi: 10.1093/nar/gkv007. PubMed PMID: 25605792; PMCID: PMC4402510.

56. Subramanian A, Tamayo P, Mootha VK, Mukherjee S, Ebert BL, Gillette MA, Paulovich A, Pomeroy SL, Golub TR, Lander ES, Mesirov JP. Gene set enrichment analysis: a knowledge-based approach for interpreting genome-wide expression profiles. Proc Natl Acad Sci U S A. 2005;102(43):15545–50. Epub 20050930. doi: 10.1073/pnas.0506580102. PubMed PMID: 16199517; PMCID: PMC1239896.

57. Korotkevich G, Sukhov V, Budin N, Shpak B, Artyomov MN, Sergushichev A. Fast gene set enrichment analysis. bioRxiv. 2021:060012. doi: 10.1101/060012.

58. Liberzon A, Birger C, Thorvaldsdottir H, Ghandi M, Mesirov JP, Tamayo P. The Molecular Signatures Database (MSigDB) hallmark gene set collection. Cell Syst. 2015;1(6):417–25. doi: 10.1016/j.cels.2015.12.004. PubMed PMID: 26771021; PMCID: PMC4707969.

59. Hanzelmann S, Castelo R, Guinney J. GSVA: gene set variation analysis for microarray and RNA-seq data. BMC Bioinformatics. 2013;14:7. Epub 20130116. doi: 10.1186/1471-2105-14-7. PubMed PMID: 23323831; PMCID: PMC3618321.

60. Wasko UN, Jiang J, Dalton TC, Curiel-Garcia A, Edwards AC, Wang Y, Lee B, Orlen M, Tian S, Stalnecker CA, Drizyte-Miller K, Menard M, Dilly J, Sastra SA, Palermo CF, Hasselluhn MC, Decker-Farrell AR, Chang S, Jiang L, Wei X, Yang YC, Helland C, Courtney H, Gindin Y, Muonio K, Zhao R, Kemp SB, Clendenin C, Sor R, Vostrejs WP, Hibshman PS, Amparo AM, Hennessey C, Rees MG, Ronan MM, Roth JA, Brodbeck J, Tomassoni L, Bakir B, Socci ND, Herring LE, Barker NK, Wang J, Cleary JM, Wolpin BM, Chabot JA, Kluger MD, Manji GA, Tsai KY, Sekulic M, Lagana SM, Califano A, Quintana E, Wang Z, Smith JAM, Holderfield M, Wildes D, Lowe SW, Badgley MA, Aguirre AJ, Vonderheide RH, Stanger BZ, Baslan T, Der CJ, Singh M, Olive KP. Tumour-selective activity of RAS-GTP inhibition in pancreatic cancer. Nature. 2024;629(8013):927–36. Epub 20240408. doi: 10.1038/s41586-024-07379-z. PubMed PMID: 38588697; PMCID: PMC11111406.

